# Anticipatory metabolic reprogramming distinguishes caloric restriction from fasting-refeeding cycles

**DOI:** 10.64898/2026.03.15.711957

**Authors:** Nikkhil Velingkaar, Artem A Astafev, Archana Prabahar, Evelina Trokhimenko, Josefa-Marie B. Rom, Gena J Asi, Peng Jiang, Roman V Kondratov

**Affiliations:** Center for Gene Regulation in Health and Disease and Department of Biological Geological and Environmental Sciences, Cleveland State University, Cleveland, Ohio, 44115, USA

**Keywords:** aging, metabolism, caloric restriction, circadian rhythms, gene expression, fatty acid metabolism, transcription

## Abstract

Interest in fasting-based dietary interventions to improve metabolic health is growing. Caloric restriction (CR) with one meal per day includes an extended fasting component that contributes to its metabolic and longevity benefits, yet the specific role of fasting within CR remains unclear. Here, we compared mice under CR with those subjected to a fasting–refeeding–fasting (FRF) regimen while controlling pre-fasting food intake and fasting duration. Simultaneous comparison of diet induced changes in plasma insulin and free fatty acids, hepatic mTOR signaling and ketogenesis, total body metabolic rhythms with kinetics of food digestion suggested that gastric emptying served as a primary metabolic trigger in acute fasting. In contrast, in CR, fasting responses were actively regulated and suggested anticipatory mechanisms. At the transcriptomic level, CR enhanced circadian rhythmicity and metabolic gene coordination, whereas FRF disrupted it. In agreement with the expression data, CR improves glucose and fatty acid metabolism while fasting leads to glucose intolerance and fat accumulation in the liver induced glucose intolerance and hepatic steatosis. These findings reveal that CR engages clock-aligned, anticipatory metabolic control, while fasting–refeeding cycles rely on direct nutrient cues. This mechanistic distinction between active and passive metabolic regulation may underlie the superior metabolic and longevity outcomes of caloric restriction.

## Introduction

Disruption of metabolic homeostasis is a significant contributing factor in the development of many diseases including diabetes, cancer, cardiovascular and neurological diseases.^1^ Increased body weight and obesity have a profound negative impact on metabolism and are strongly associated with disease risk.^2^ Healthy lifestyle interventions such as dietary modifications and physical exercise, often in combination with pharmacological treatments, have been strongly recommended as strategies to prevent and manage metabolic diseases.^3–6^ Among various lifestyle interventions, dietary modification is an efficient non-pharmacological way to regulate body weight and improve metabolic health. Various dietary interventions, including fasting-based regimens such as time restricted fasting (TRF) and intermittent fasting (IF), are growing in popularity for their potential to effect metabolic benefits.^7–9^

Caloric restriction (CR) is a diet where food intake is chronically reduced without malnutrition. CR robustly improved metabolic health and increased lifespan in a variety of organisms, from yeast to mammals.^10–13^ In mammals, CR has been proven to improve insulin sensitivity, glucose and lipid metabolism and delay development of many diseases.^14–16^ The beneficial impact of CR on metabolism has also been demonstrated in humans, as shown by the CALERIE trials.^17,18^ At the molecular level, CR impacts multiple physiological and biochemical processes, such as insulin and mTOR signaling pathways, autophagy and protein homeostasis, glucose and fat metabolism, and circadian clocks and rhythms and these pathways are implicated in molecular mechanisms of CR.^15,19–23^

Rodents are popular models to study the mechanistic basis of CR-induced metabolic and longevity benefits. One of the most common ways to implement CR in rodents is to provide daily food intake as a single meal at a fixed time each day. Implementation of this feeding regimen results in mice being entrained to receive food at the same time of the day and them consuming all the provided food within two hours of the feeding window, thus imposing a self-implemented time-restricted eating, followed by a prolonged fasting period of 22 hours until the next meal.^21,22,24^ Contrary to that, ad libitum (AL) fed mice have constant access to unlimited food and eat multiple rounds of small meals throughout the day and night. This prolonged fasting component of CR has been indicated to contribute significantly to the longevity benefits of CR in rodents. Indeed, lifespan extension is dramatically attenuated when CR is implemented in rodents without the prolonged fasting component, either by providing small CR meals across the day or by using a low-calorie chow, compared to the one-meal-a-day CR regimen.^24–26^

A few recent studies have reported both fasting-dependent and fasting-independent contributions of caloric restriction to lifespan extension, thus arguing for better understanding the role of fasting in molecular mechanisms of CR.^15,26–28^ Several recent reports have directly compared the physiological and molecular changes induced by CR and fasting at either one or multiple time points across the day.^19,24,25,27,29^ This approach identified similar changes induced by both diets such as a reduction in blood glucose, an increase in blood NEFA, and reduced mTOR activity. At the same time, CR and fasting differentially impact circulating metabolites in the blood, insulin sensitivity, glucose tolerance, hepatic PPARα signaling, fatty acid metabolism, and ketogenesis.^21,25,29,30^ Thus, CR and fasting induce overlapping but distinct metabolic states; however, only a subset of metabolic benefits of CR can be attributed to the fasting component. One of the key differences between CR and fasting is the element of entrainment to periodic feeding- fasting cycle in CR and the involvement of the circadian clock.

Circadian clock is a hierarchical network of tissue oscillators that generate 24-hour circadian rhythms in gene expression and physiology and synchronizing metabolic pathways with environmental and nutritional cues.^31,32^ Peripheral clocks are coordinated by the central clock in SCN to synchronize behavior, feeding/fasting cycle, and metabolic processes with external environment. Molecular component of the circadian clock transcriptional factors CLOCK (encoded by *Clk* gene) and BMAL1 (encoded by *Arntl1* gene) directly regulate the expression of three *Period* (*Pers*) genes, two *Cryptochromes* (*Crys*) genes, *Nr1D1*, *Nr1D2* (encodes REV-ERB-beta and REV-ERB-alpha transcriptional factors) and *Ror*α and γ (encodes retinoid receptor) genes. In turn, PERs and CRYs inhibit CLOCK/BMAL1 complex activity, REV-ERBs inhibits, and ROR gamma induces *Bmal1* gene transcription, thus forming negative and positive feedback loops. Circadian clock and circadian rhythms have been demonstrated to be an integral component of CR-mediated metabolic benefits, and disruption of circadian rhythms attenuates the metabolic and longevity benefits of CR in mice.^19,22,33,34^ Together, these observations warrant the need for a refined experimental approach to bifurcate fasting-dependent effects from entrainment-dependent effects of CR.

Direct comparison of CR and fasting provides significant advantage to study mechanisms of CR. Recent reports compare CR and fasting implementing unanticipated food withdrawal from AL mice. However, this approach lacks information on the precise previous feeding time and the amount of food consumed by AL-fed animals before the food was withdrawn and fasting was initiated. These unknown components limit direct comparing with the most popular one-meal-a-day CR when a fixed amount of food is provided at a fixed time of the day. Thus, in the present study, we addressed this limitation by redesigning the fasting approach to better control feeding-fasting component and help directly compare with CR. We implemented a fasting-refeeding-fasting regimen (FRF), which better mimicked the CR feeding-fasting cycle, but did not involve the component of entrainment. We performed a high temporal resolution analysis of the liver transcriptomes at four-hour intervals over a 24-hour period in CR and FRF mice and assayed the impact of both feeding regimen on mTOR signaling, glucose metabolism, and lipid metabolism. We found that CR and FRF reprogrammed the liver transcriptome in overlapping but distinct ways, with the opposite effects on circadian gene expression. Consistent with these transcriptional changes, CR improved glucose and fatty acid metabolism, while FRF impaired the glucose tolerance and fatty acid metabolism. Most importantly, the metabolic transition from fed to fasting state was tightly coupled to gastric emptying in FRF mice, contrary to CR mice, where the transition, unexpectedly, was not directly connected with gastric emptying, suggesting that entrainment governs/regulates metabolic state transitions in CR. These observations indicate that CR engages regulatory mechanisms, preferably clock-aligned, anticipatory metabolic program that decouples from gastric emptying, unlike nutrient-based metabolic transition in FRF regimen.

## Results

### Effect of CR and unanticipated Fasting on glucose and fat metabolism

At 3 months of age, male and female mice were randomly assigned to Ad libitum (AL) and 30% calorie-restricted (CR) groups. CR mice received 70% of their AL food intake as a single meal once per day at ZT14, two hours after the light was turned off. After three months on the diets, the CR mice had significantly reduced body weight (Figure 1B) compared with the AL group. CR mice also entrained themselves to consume all provided food between ZT14 and ZT16, as was previously reported,^21,22^ and were without food for about 22 hours. To understand the contribution of this self-implemented fasting to CR metabolism, the food was unexpectedly removed from AL mice at ZT16, referred to as Fasting (Figure 1A).

**Figure 1.**
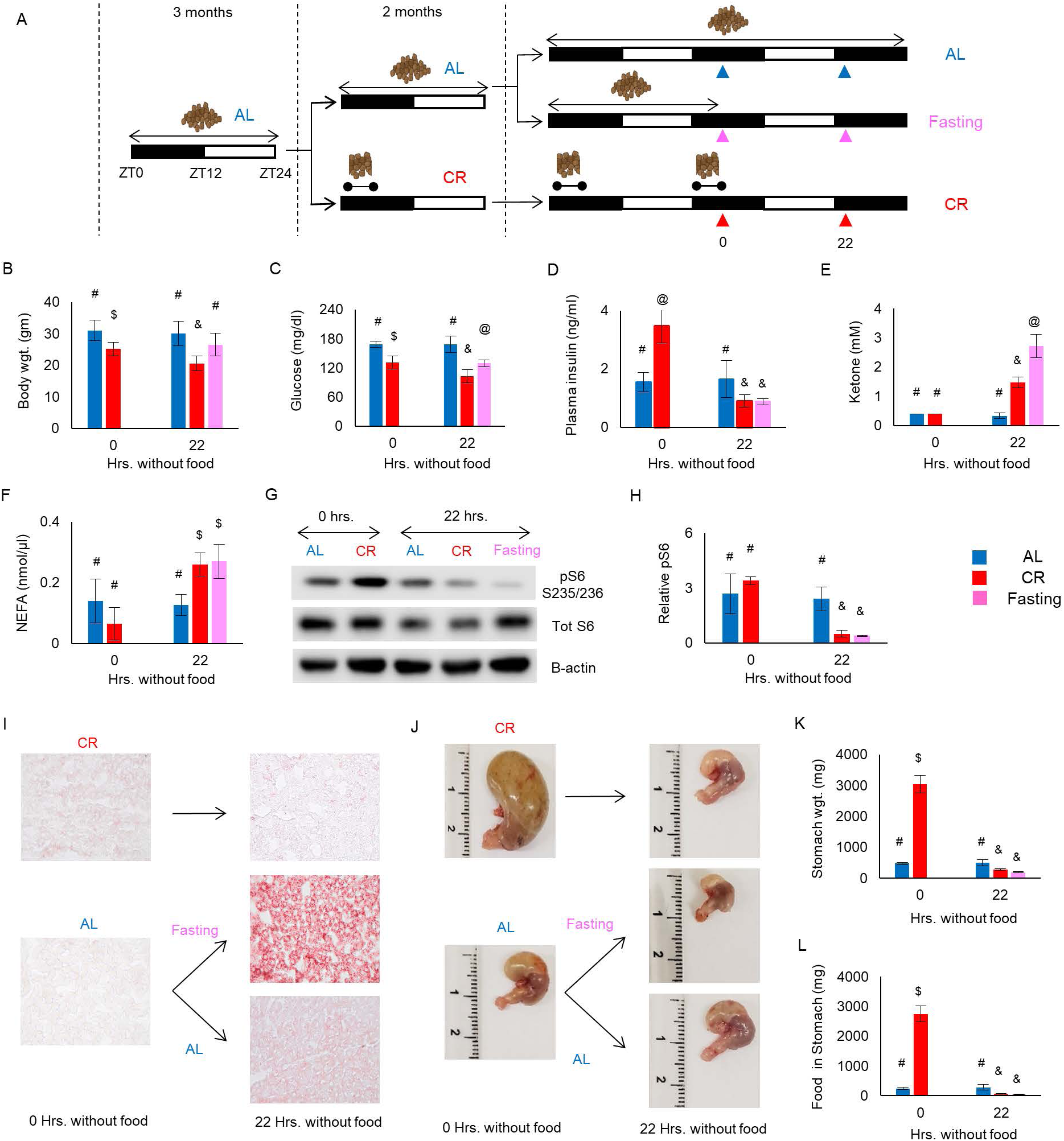
Caloric restriction induces a distinct metabolic state compared to acute fasting. (A) Scheme of feeding protocol. Mice were maintained on Ad libitum (AL) diet for 3 months after which they were randomly assigned into two groups: AL had unlimited access to the food around the clock, 30% calorie restriction (CR 30%). After 2 months, CR group continued receiving 30% CR food whereas AL group was further split into two groups: one group continued AL, another group had their food access withdrawn (Fasting – F) on a random day at ZT16 and fasted for 22 hours. Analysis was performed at two time points ZT16 (0 hours) and ZT14 (22 hours), and are indicated by colored arrows. (B) Absolute body weight (AL, CR, and Fasting (n=8 per diet)) measured at 0 hours and 22 hours without food. AL - blue bar; CR – red bar; Fasting 1 – pink bar. (C) Glucose (n=4) in the blood obtained from tail vein nick of mice subjected to AL, CR and Fasting regimen at 0 hours and 22 hours without food. (D) Plasma insulin levels in the blood of mice on AL, CR, and Fasting (n=4 per diet) at 0 hours and 22 hours without food. (E) Blood βOHB obtained from the tail of mice on AL, CR, and Fasting (n=4 per diet). (F) Serum NEFA levels measured in the tail blood of AL, CR, and Fasting mice (n=4 per diet). (G) Western blot, and (H) Quantification of ribosomal protein S6 in the liver of mice subjected to AL, CR, and Fasting (n=4 per time point per diet). (I) Oil Red O staining of mouse livers from each of the 3 diets (n=3 per time point). (J) Stomach weight, and (K-L) Quantification of stomach weight and amount of food retained in the stomach of mice maintained on all 3 diets (n=4 per diet per time point). All data represented as Mean ± SD. For comparison between diets at 0 hours without food, unpaired t-test with Welch’s correction was used. For comparison between diets at 22 hours without food, One-way ANOVA with Tukey’s correction for post-hoc analysis was used. For comparison within diets at 0 hours and 22 hours without food, unpaired t-test with Welch’s correction was used. Different symbols indicate statistically significant effect of the diet, whereas the same symbols indicate no significant effect of the diet. Statistical significance was set at p<0.05. Light was turned on at ZT0, and light was turned off at ZT12. Light and Dark bars indicate light and dark phases of the day. AL is represented by blue bar, CR is represented by red bar, and Fasting is represented by pink bar.

At the starting point, 0 hours without the food, the blood glucose was significantly lower (Figure 1C), and plasma insulin was significantly higher (Figure 1D) in CR compared with AL. The blood ketones (Figure 1E), blood NEFAs (Figure 1F), and liver TAGs, judged by Oil Red O staining (Figure 1I, Supplementary Figure 1), were comparable between AL and CR mice. After 22 hours without food, the blood glucose and liver TAGs did not significantly change; blood plasma insulin reduced, blood ketones and NEFAs increased in CR mice (Figure 1E-F, 1I, Supplementary Figure 1). Blood glucose and blood plasma insulin significantly reduced; blood ketones, blood NEFAs and liver TAGs increased after 22 hours of fasting (Figure 1E-F, 1I, Supplementary Figure 1). AL mice that were never fasted were also assayed at the same time points to account for possible circadian variability, and no significant changes were detected for any of the above parameters (Figure 1C, 1E-F). mTOR pathway is a nutrient sensing pathway and master regulator of metabolism, which regulates transition from anabolism in fed state to catabolism in fasting. In fasting, mTOR activity reduces, and increases upon refeeding.^35–37^ To estimate whether the fasting response was induced in both diets, we monitored mTOR activity in the liver of mice on the AL, CR, and F diets using phosphorylation of ribosomal protein S6 as a readout. As was expected, 22 hours without food resulted in a significant reduction of S6 phosphorylation in both CR and Fasting (Figure 1H). Thus, after 22 hours without food, some changes were similar between CR and F, while some changes were observed only in Fasting, with liver TAGs accumulation was the most striking difference between CR and F.

When tissues were harvested for the analysis, a significant difference in the stomach size was noted between CR and AL mice. The amount of food in the stomach was significantly higher in CR mice at time point 0 hours (ZT16) compared with AL (Figure 1K). There was no difference between CR and Fasting at 22 hours without food (ZT14), and it was significantly less in both CR and FRF compared with 0 hours or with AL at the same time. AL fed mice eat during dark phase of the day, but they also eat during the light phase, and they eat at random time points. Therefore, the amount of food consumed by AL mice and the exact time of feeding was not known before the food was removed at ZT16, which might contribute to the difference in metabolic response in CR and fasting.

### CR and FRF have similar responses to feeding-fasting cycles

The difference in the food, significantly less in AL compared with CR, and the stomach size at the starting time point of fasting complicated result interpretation. To address that, we designed the following experiment (Figure 2A). On one random day, the food was withdrawn from a group of AL mice at ZT16. After 22 hours without the food, the mice were provided with unlimited amount of food for 2 hours, between ZT14-16 (the same time when CR mice consumed their food), after which the food was removed again. These mice were designated as fasted-refed-fasted (FRF). Thus, CR mice and FRF mice started the meal at the same time, ZT14, and they finished the feeding at the same time, ZT16. ZT16 was set up as 0 hours without food for CR and FRF mice. Thus, the feeding time at ZT14 was set up as -2 hours. The food intake of AL mice was also measured during the same time between ZT14 and ZT16 and these animals consumed about 0.5 grams (about 15% of their daily food intake). Both CR and FRF mice consumed significantly more compared to AL mice. Importantly, during the refeeding phase, while FRF mice consumed slightly less than CR mice, both groups consumed a comparable amount of food (Figure 2B). Thus, in this design, the CR and FRF mice started fasting at the same time and ate a comparable amount of food. We monitored physiologic and metabolic changes in these groups of mice by collecting samples every 4 hours across the day (Figure 2A). AL mice that never fasted were collected at the same time points and served as a control for possible circadian variability.

**Figure 2.**
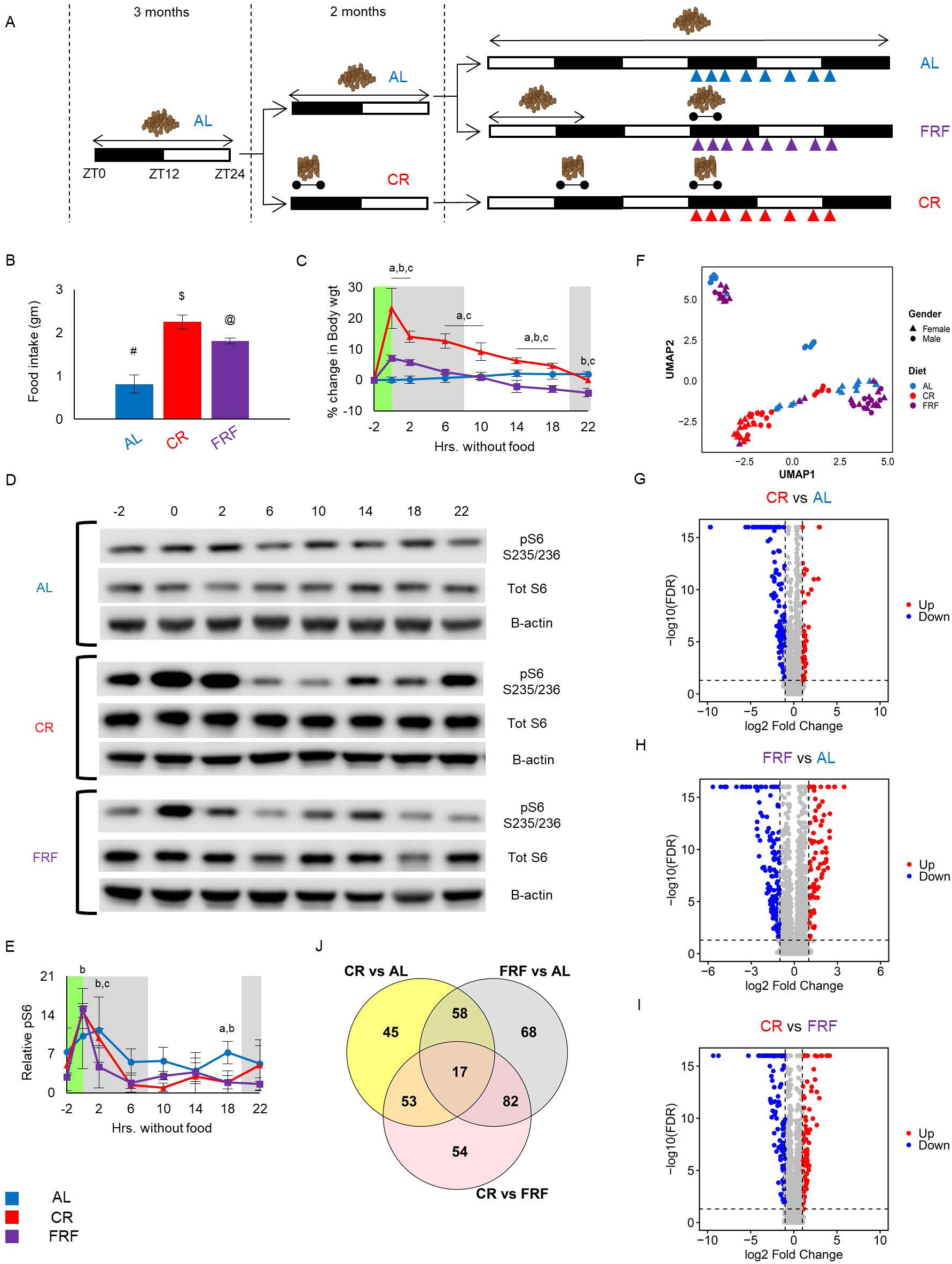
CR and FRF demonstrate comparable metabolic responses to controlled feeding. (A) Experimental design depicting applied diet. Mice in the AL group were randomly split into 2 groups, one group continued to receive AL diet, another group was assigned to Fasted-refed-fasted (FRF) diet. Mice in the FRF group had their food removed unexpectedly on a random day at ZT16 and fasted for 22 hours. FRF mice were then fed at the same time as CR group (ZT14) and 2 hours later (ZT16), had their food access removed again for 22 hours. Mice in the FRF group were then compared with the CR group. Mice in the AL group were used as controls. Blue, red, and purple triangles indicate time of blood and tissue collection. (B) Amount of food consumed by mice on AL, CR, and FRF (n=8 per diet) between ZT14 and ZT16. (C) Change in body weight (in %) measured every 4 hours without food in AL, CR, and FRF mice (n=8 per diet per time point). Body weight at ZT14 (-2 hours without food) was used as the starting point for all 3 diets to calculate % change in body weight with duration without food. (D) Representative western blotting and (E) quantification for mTORC1 signaling in the liver of mice on AL, CR, and FRF (n=4 per time point). For the western representative image, lysates from four individual mice were pooled together. (F) UMAP colored by diet and gender denoted by different symbols show that samples clustered together in a diet-dependent manner. (G-I) Volcano plot of changes in hepatic transcriptome in mice when compared between two diets. All time points were combined for this analysis. Dashed lines indicate cut-off for significance (p≤0.05) and fold change (log_2_ > 2 for upregulated genes, < 0.5 for downregulated genes). (J) Venn diagram analysis shows distribution of differentially expressed genes when all three diets (AL, CR, FRF) are compared. All time points are pooled for this analysis. Feeding period for CR and FRF mice is indicated by green bar. All data represented as Mean ± SD. One-way ANOVA with Tukey’s correction for multiple comparison was performed for food intake measurements. Two-way ANOVA with Tukey’s correction for multiple comparisons was performed for percent change in body weight and western blot quantification. Letters indicate statistically significant effect of the diet; a – AL versus CR, b – AL versus FRF, c – CR versus FRF. For food intake, different symbols indicate statistical significant effect of the diets whereas the same symbols indicate no statistical significant effect of the diets. Statistical significance was set at p<0.05. Light was turned on at ZT0 and light was turned off at ZT12. Grey bars indicate dark phase of the day. AL is represented by blue bars and blue solid lines, CR is represented by red bars and red solid lines, and FRF is represented by purple bars and purple solid lines.

First, we checked for changes in body weight across the day (Figure 2C). The body weight of AL mice did not significantly change across the day, most likely due to the feeding throughout the day. Both CR and FRF mice showed a significant increase in body weight between -2 hours and 0 hours without food, which corresponded to their feeding activity between ZT14-16. After that their body weight gradually decreased with duration of time without food. Because the activity of mTOR complex reduces with duration of fasting and increases upon feeding, we measured it in the liver of mice on all three diets to monitor a molecular transition between fed and fasting states. mTOR activity demonstrated low amplitude rhythms in the AL liver, high during the dark phase and low during the day in agreement with the nocturnal feeding pattern of these mice. In CR liver, the mTORC1 activity was low before the feeding (-2 hours); it rapidly increased after the feeding, peaked 4 hours after the feeding, at ZT18, and reduced with duration of the time without food. In the FRF liver, the mTOR activity was low at -2 hours, increased after refeeding, and decreased gradually with an increase in time without food (Figure 2E). Thus, changes in mTOR activity followed similar kinetics in both CR and FRF livers, suggesting a comparable transition from fed to fasting states.

### CR and FRF reprogramed liver transcriptome in overlapping but distinct ways

Diets, including CR, are known to impact gene expression. To understand global transcriptome effects of feeding regimens, we analyzed the liver transcriptome in mice subjected to AL, CR, and FRF. UMAP analysis revealed diet-dependent organization of the liver transcriptome (Figure 2F). AL samples were segregated into four different groups with clear separation of males and females. CR samples clustered as two groups: one group was males exclusively, and the other consisted of both males and females. FRF samples also clustered as two main groups, but contrary to CR, they clustered with two AL groups and there was no separation between male and females. While sexual dimorphism in the liver transcriptomes existed in a diet-specific manner, we decided to focus on the effects of diets independent of sex.

To identify time-of-the-day-independent effect of the diets, all time points were pooled together and results of differential expression between diets are presented as volcano plots in Figures 2G-J and the list of genes for each pairwise comparison are presented in Supplementary Table 1. 46 differentially expressed genes (DEGs) were upregulated and 127 DEGs were downregulated by CR compared to AL. 84 DEGs were upregulated and 141 DEGs downregulated by FRF compared to AL. 43% of DEGs (75 out 173) affected by CR were also affected by FRF. This significant overlap suggested similarity in reprogramming of the liver transcriptome by both diets, but significant number of DEGs were unique for each diet, thus, while both CR and FRF diets have a common fasting component, the metabolic reprogramming was not the same.

While pooled analysis revealed robust diet-dependent changes in transcriptome, it did not resolve temporal specificity. Hence, we next examined transcriptional responses at each individual time point across three diets. Volcano plots and pathway enrichment analyses for pairwise comparison between AL, CR, and FRF are presented in Supplementary Figure 2. Venn diagrams of DEGs from pairwise comparison between diets and representative examples are shown in Supplementary Figure 3A-C and the list of genes for each pairwise comparison are presented in Supplementary Tables 3-10. Several important observations were made from these comparisons the total number of DEGs in each pairwise comparison changed with the time without food, the lists of DEGs were different at each time points, many genes were differentially expressed only at one or two time points. Figure 3A-C (left panels) summarizes DEGs between the diets across the feeding cycle using Circos plots when DEGs for each time points are connected. The pathways enriched are summarized across time without food. Pathway analysis revealed that different metabolic processes were induced in CR and FRF with anabolic processes immediately after the refeeding were replaced by catabolic processes with duration of time without food. Interestingly, fatty acid metabolism pathways were highly enriched in FRF compared to AL and to CR, while carbohydrate metabolism pathways were enriched in CR compared with AL.

**Figure 3.**
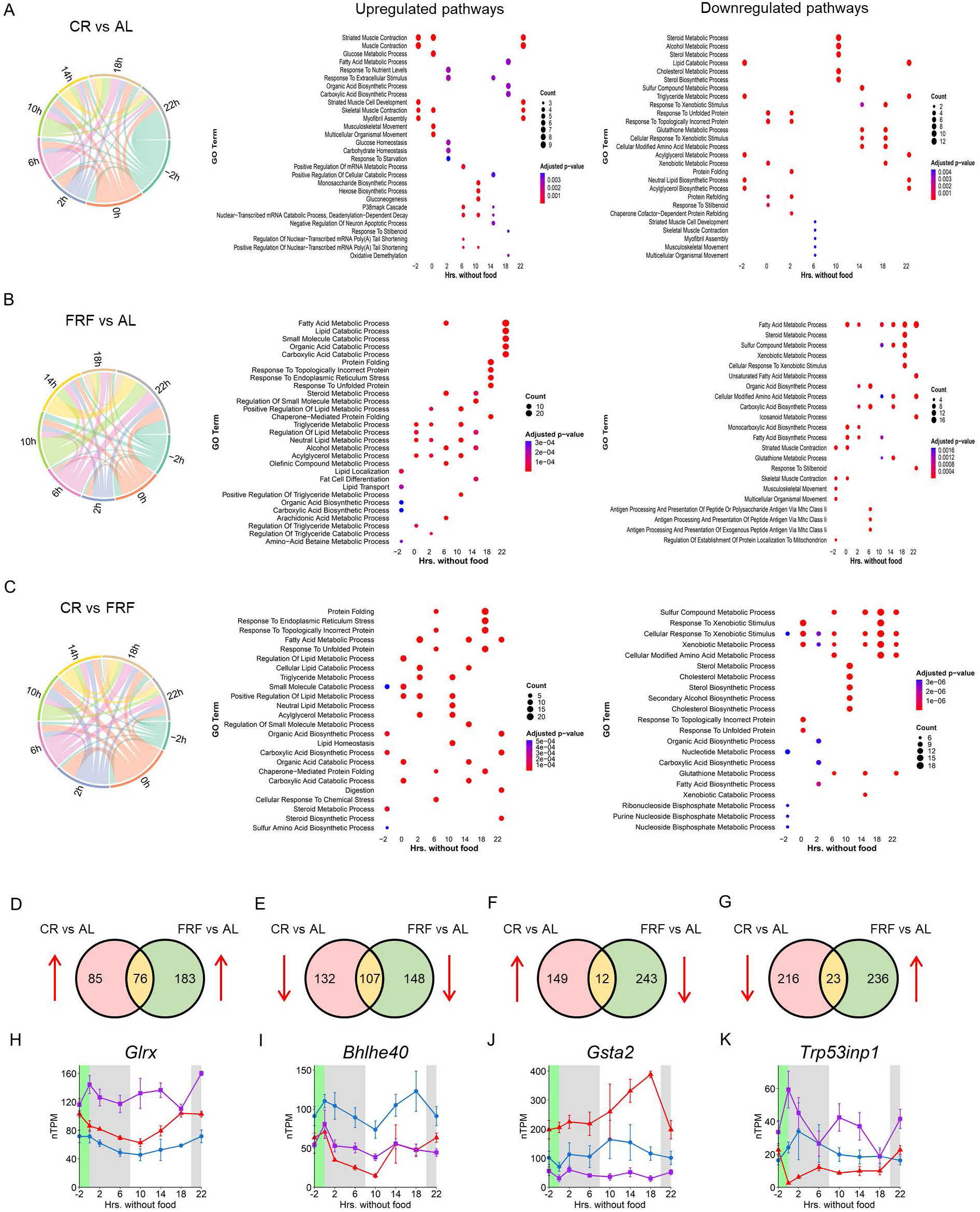
Distinct transcriptional states underlie CR and FRF. (A-C) Left panel indicates Circos plots that show connection of differentially expressed genes across time without food, when comparing any two diets. Right panel indicates GO pathway annotations that are either upregulated or downregulated with respect to duration without food. For each time point without food, only top 5 pathways were selected. Only those top 5 pathways whose adjusted p-value was <0.05 were selected for the plot. (D-G) Venn diagram indicating overlap of differentially expressed genes (DEGs) that were identified in pairwise comparison of CR vs AL and FRF vs AL. DEGs at the intersection are common genes that are either upregulated or downregulated in both CR and FRF or regulated in opposite direction compared to AL. Red arrows next to Venn diagrams indicate direction of change of these DEGs. (H-K) Representative example of a DEG that belongs to the intersection indicated in panel 3D-G. A complete list of DEGs in this shared subset is provided in Supplementary Table 2, with common DEGs being highlighted in green.

In total, 161 genes were up regulated in CR compared with AL and in 259 genes upregulated in FRF compared with AL, while 239 genes were down regulated in CR and 255 genes were down regulated in FRF when unique DEGs were counted for each time point and diet. We compared these lists for the genes that are differentially expressed at least at one time point. Results of these comparisons are represented as Venn diagrams in Figure 3D-G. In these Venn diagrams, red arrows indicate the direction of regulation relative to AL. Similar with the analysis in Figure 2F-J more genes were affected by FRF then by CR and about 50% DEGs were common between the diets (Figure 3D-G, Supplementary Table 2). Representative examples of common DEGs that are either upregulated, downregulated or regulated in opposite direction by CR and FRF are represented in Figure 3H-K. When focused on DEGs common between CR and FRF, 183 (76 upregulated and 107 downregulated) were regulated in the same direction, whereas only 35 (12 upregulated and 23 downregulated) exhibited an opposite regulation by diets. For some DEGs, the magnitude of the changes was also different between CR and FRF. Thus, the detailed time point analysis further confirmed overlapping but distinct reprogramming of the liver transcriptome by CR and FRF.

### CR enhanced while FRF disrupted circadian rhythms of liver transcriptome and circadian clock gene expression

Diets, including CR, are known to affect circadian rhythms in gene expression.^19,24,38,39^ Circadian rhythms in gene transcription are regulated on multiple levels including epigenetic changes and circadian clock plays an important role in this regulation.^40^ To investigate the effect of fasting-refeeding-fasting (FRF) intervention on circadian rhythms we directly compared the effect of the diets on the rhythms of liver transcriptomes (Figure 4A) and on the rhythms in the expression of core circadian clock genes (Figure 4B, C). Molecular circadian clock is negative and positive feedback loops formed by dozens of genes and their products: *Clk*, *Arntl1* (encodes BMAL1), three *Period* (*Pers*), two *Cryptochromes* (*Crys*), *Nr1D1*, *Nr1D2* (encodes REV-ERB-beta and REV-ERB-alpha) and *Ror*α and γ. In AL liver, the expression of circadian clock genes, with exception for *Rorα,* was rhythmic. In the CR liver, although several genes showed a modest one-hour phase shift, all clock genes were rhythmic. In FRF liver, the circadian rhythms of expression for all clock genes were disrupted (Figure 4B, C). Thus, even though fasting and feeding time windows were aligned between FRF and CR diet, CR and FRF had opposite effect on molecular circadian clock in the liver: CR increases robustness of the circadian clock gene oscillation in agreement with previous reports,^19,22,33^ while FRF disturbed the rhythms.

**Figure 4.**
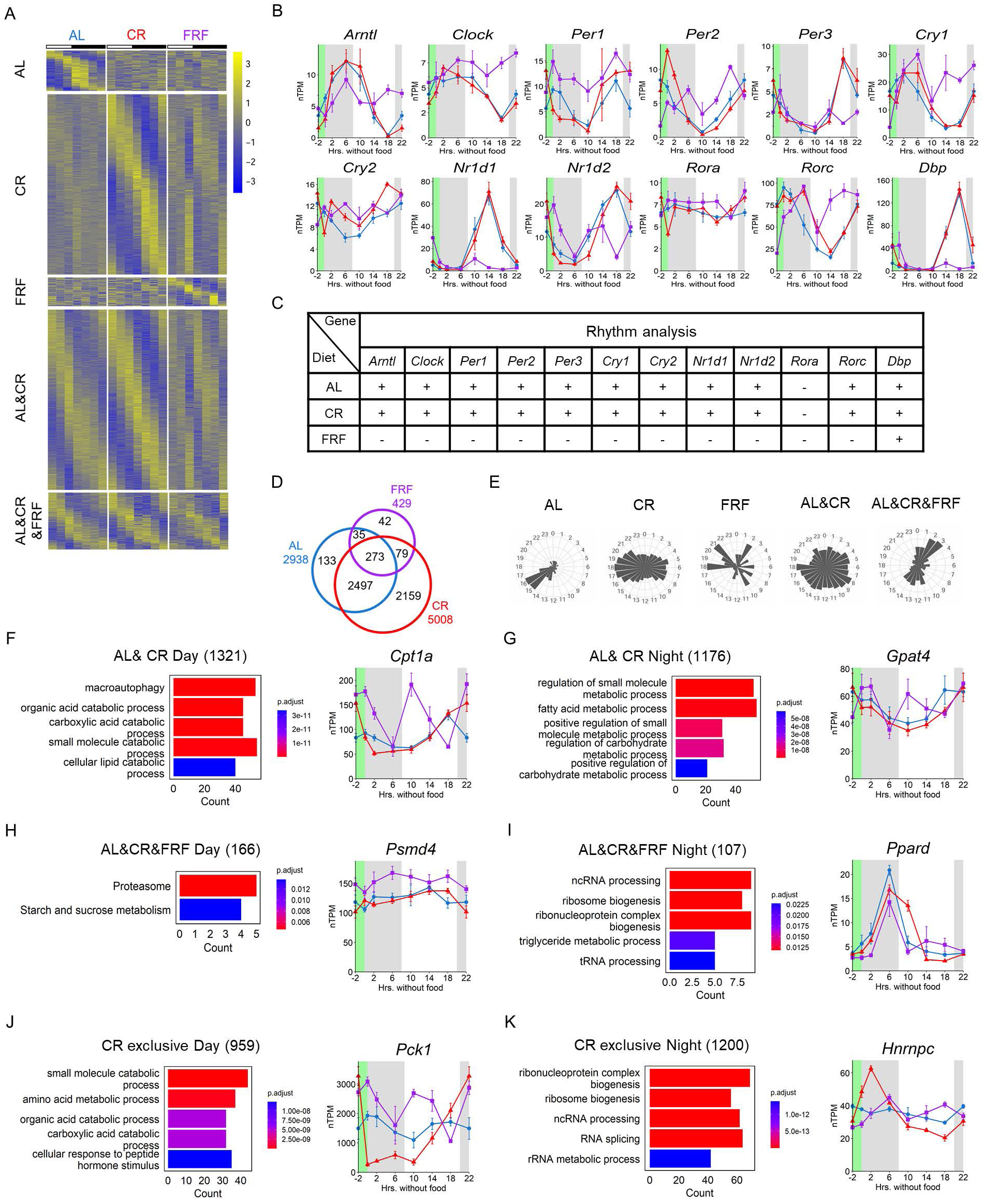
FRF disrupts hepatic circadian rhythms. (A) Phase-sorted heatmap of rhythmic transcripts from the groups shown above. Rhythmicity was determined using the R Studio (version 4.5.1) software in the DryR package (BICW > 0.4), and five cluster of genes were identified. (B) mRNA expression of core clock genes in the liver of mice on AL, CR, and FRF diets (n=4 per time point per diet). (C) DryR rhythmicity analysis of core clock genes in the liver of mice on AL, CR, and FRF diets. “+ and -” indicate genes are either rhythmic or non-rhythmic. (D) Venn diagram and (E) Phase distribution of genes that are rhythmic as determined by DryR analysis. (F-K) Pathway analysis using Gene Ontology and representative examples of genes that are rhythmic either during the day (rest phase) or night (active phase). Only those top 5 pathways whose adjusted p-value was <0.05 were selected for the plot.

As expected and previously reported by us^41^ and others,^24,42^ CR increased the number of rhythmic genes. FRF severely disrupted the 24h rhythms in the transcriptome and significantly reduced the number of rhythmic genes (Figure 4A, B). The largest group of rhythmic genes (2497) was rhythmic under both AL and CR, and under FRF this group of genes lost their rhythmicity (Figure 4A). This group of genes was enriched for pathways involved in autophagy and fatty acid catabolism (Figure 4F) for the day-peaking subset, and for pathways involved in fatty acid and carbohydrate metabolism (Figure 4G) for night-peaking subset. The large number of genes (2159; Figure 4A, D-E) became rhythmic under CR. Within this group, the day-peaking subset was enriched in genes involved in amino acid and fatty acid catabolism (Figure 4J), and the night-peaking genes in RNA processing and ribosome-related pathways (Figure 4K). Interestingly, a smaller group of genes (273; Figure 4A, D-E) stayed rhythmic on all diets – among those genes, the day-peaking were involved in proteasome and sucrose metabolism (Figure 4H), while the night-peaking genes were involved in triglyceride metabolism and tRNA processing (Figure 4I). Thus, CR and FRF have opposite impact on the circadian rhythms in liver gene expression and differential effect of CR and FRF on the rhythms of core circadian clock genes contributed to the differential reprograming of circadian liver transcriptome induced by the diets.

### Caloric Restriction and FRF have different effect on glucose homeostasis

GO pathway and rhythmic analysis identified glucose metabolism as a top pathway being affected by both CR and FRF. Figure 5A represents a schematic overview of glycolysis and gluconeogenesis, emphasizing the major rate-limiting enzymes. Enzymes whose transcripts displayed differential or rhythmic expressions in our dataset are highlighted. The heatmap in Figure 5B depicts the expression of all genes involved in both pathways. While both CR and FRF significantly impacted the expression of many glucose metabolism genes, the effects were different between the diets. Importantly, the number of rhythmic genes increased in CR liver, the expression *Gck, Pfkl, Adoa, AldoC* and *Pck1* became highly rhythmic in CR liver. At the same time the rhythms were disrupted in the FRF liver. All these changes led us to the hypothesis that glucose metabolism might be highly regulated and coordinated in CR but not in FRF and, therefore, glucose homeostasis will be different between CR and FRF.

**Figure 5.**
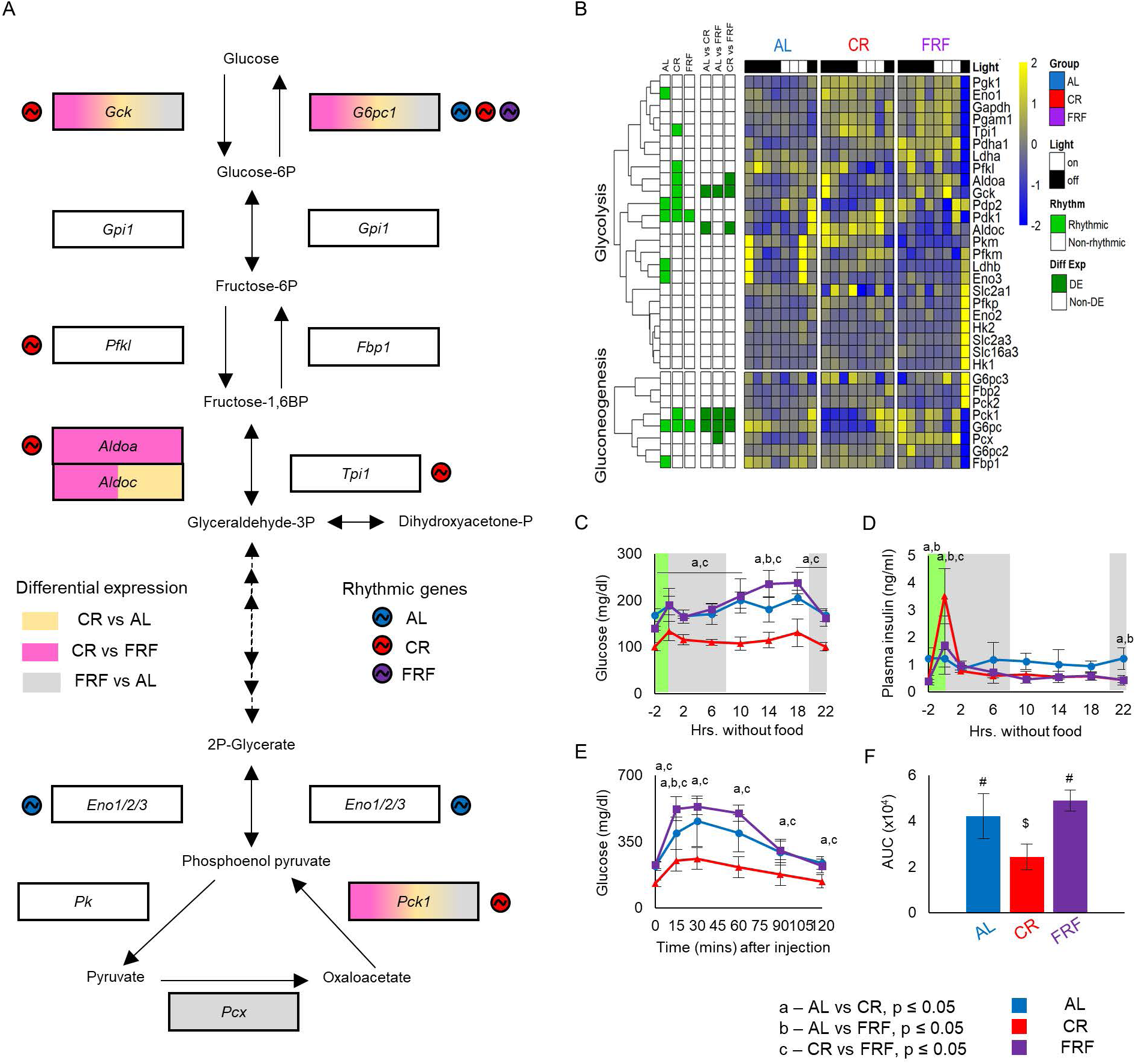
Distinct regulation of glucose homeostasis by CR and FRF. (A) Schematic representation of glycolysis and gluconeogenesis pathways. Enzymes that show differential expression or rhythmicity are highlighted. (B) Heatmap depicting the expression of all the genes involved in glycolysis and gluconeogenesis pathways in AL, CR, and FRF diets. White and black bars below the diet indicate light (rest) and dark (active) phase. (C) Blood glucose levels in mice subjected to AL, CR, and FRF diet (n=4 per time point). (D) Plasma insulin levels in the blood of mice on AL, CR, and FRF (n=3-4 per diet per time point) across time without food. (E) Intraperitoneal Glucose tolerance test (ip-GTT) and (F) Area under the curve in mice subjected to overnight fast on AL, CR and FRF regimen (n=4). In Figure 5F, same letters indicate no statistical significant difference, whereas different letters indicate statistical significant difference (p < 0.05). Feeding period for CR and FRF mice is indicated by green bar. All data represented as Mean ± SD. For Fig 5B-D, Two-way ANOVA with Tukey’s correction for multiple comparison was performed. For Fig 5E, One-way ANOVA with Tukey’s correction for multiple comparison was performed. Letters indicate statistically significant effect of the diet; a – AL versus CR, b – AL versus FRF, c – CR versus FRF. Statistical significance was set at p<0.05. Light was turned on at ZT0 and light was turned off at ZT12. Grey bars indicate dark phase of the day. AL is represented by blue bars and blue solid lines, CR is represented by red bars and red solid lines, and FRF is represented by purple bars and purple solid lines.

To test this hypothesis, we assayed blood glucose and plasma insulin in all three groups across the feeding cycle. As expected, AL mice showed low amplitude fluctuations in blood glucose across the day (Figure 5C). CR mice had significantly lower blood glucose compared to AL mice. The blood glucose in CR was remarkably stable across the day, it did not change with duration of fasting and was not affected by refeeding, which confirmed a strong control of glucose homeostasis in CR. FRF had a very distinct effect on blood glucose levels. Before the refeeding, at – 2 hours without food, the blood glucose was significantly lower compared with AL but higher than CR. After the refeeding, at 0 hours, blood glucose was significantly increased to the level comparable with AL. Surprisingly the blood glucose in FRF mice did not reduce with duration of fasting and was comparable or even above AL at all time points. Plasma insulin did not significantly oscillate in AL mice across the feeding cycle. CR and FRF mice showed similar kinetics of plasma insulin. Compared to AL, insulin levels were low before feeding, at -2 hours, in both CR and FRF. Insulin increased, upon the refeeding, to levels significantly higher compared with AL for a short period of time and dropped fast in the next 2 hours. Insulin levels in CR and FRF remained low compared with AL through the rest of the feeding cycle. There were some differences in insulin between FRF and CR diets: insulin peaked significantly higher in CR compared with FRF mice. Another notable observation was that insulin levels in FRF group reduced to CR levels at 2 hours without food. The kinetics of blood insulin (Figure 5D) are well correlated with kinetics of mTOR signaling (Figure 2D, E).

Finally, we assayed for glucose tolerance in mice on all three diets. Kinetics of blood glucose changes are shown in Figure 5E and calculated area under the curve (Figure 5F). Thus, CR significantly increased glucose tolerance compared with AL and FRF did not. The feeding schedule CR and FRF mice was aligned, mice in both groups were without food for comparable amount of time and had similar kinetics of blood insulin, however, only CR significantly improved glucose homeostasis judged by blood glucose and glucose tolerance. Improvement in glucose homeostasis correlated with increased rhythmic expression of glucose metabolism enzymes in line with our hypothesis.

### Caloric Restriction and FRF have different effect on liver lipid metabolism

Another major biochemical pathway affected by the diets was fatty acid metabolism. Figure 6A and Supplementary Figure 4 represent the schematics of rate-limiting enzymes involved in liver lipid metabolism, affected by CR and/or FRF and the heat map of expression for genes known to be involved in fatty acid biosynthesis, transport, storage and catabolism is shown in Figure 6B. Similar to the effect of the diets on glucose metabolism gene expression, both CR and FRF caused significant changes in the expression of multiple genes, including genes that encode the rate limiting and committed step enzymes. FRF induced effects occurred much earlier, were stronger and more genes were affected compared with CR. Another important difference between the diets was that CR increased, whereas FRF decreased, the number of rhythmically expressed genes (Figure 6A-B, Supplementary Fig 5). Thus, although the expression of fatty acid metabolism genes was affected by both the feeding regimen, the kinetics and overall outcome of these effects were distinct. We hypothesized that CR and FRF will differently affect liver fat metabolism.

**Figure 6.**
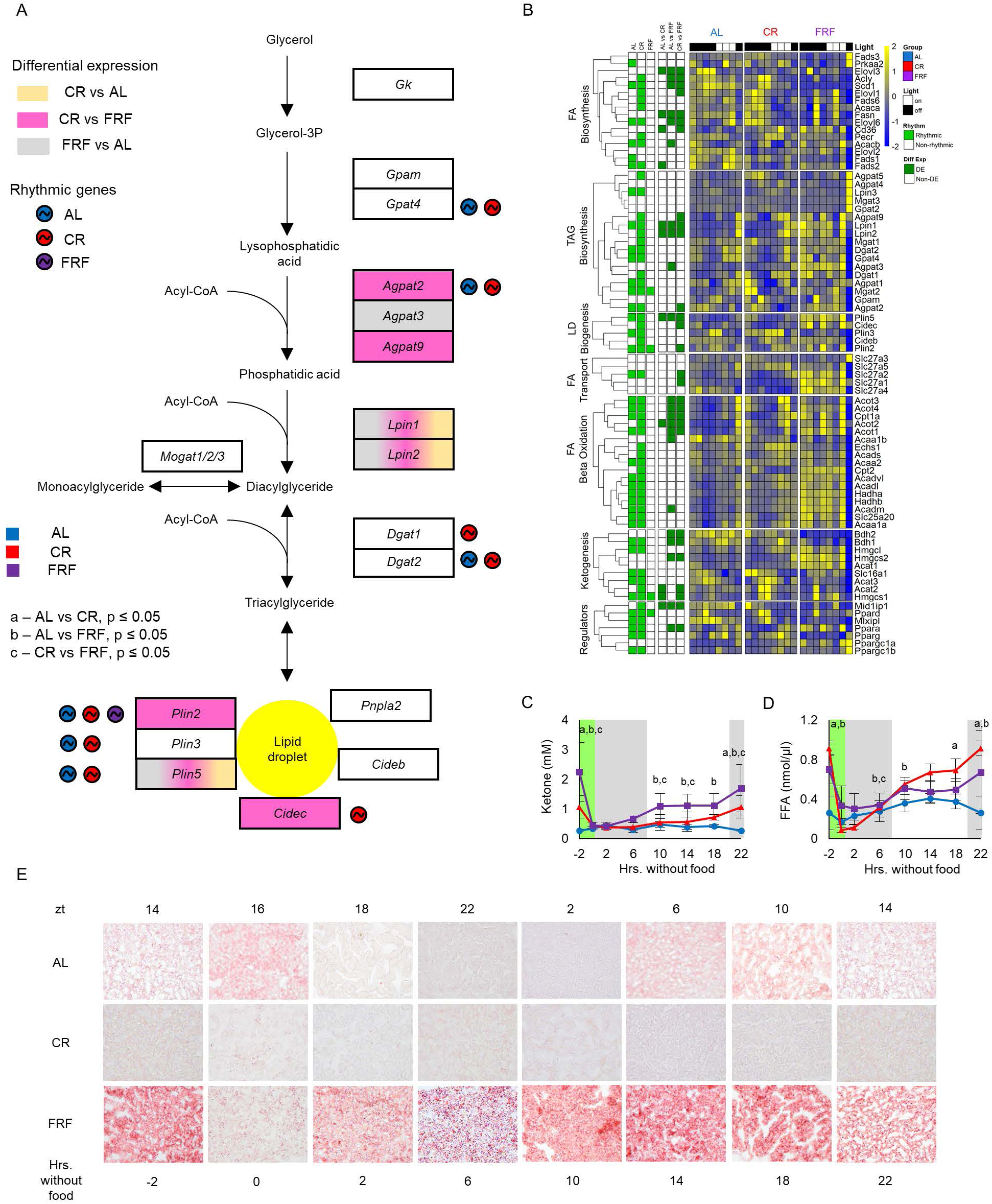
CR and FRF differentially remodel lipid homeostasis. (A) Schematic representation of lipid droplet synthesis pathway. Enzymes that show differential expression or rhythmicity are highlighted. (B) Heatmap depicting the expression of all the genes involved in lipid metabolism pathways in AL, CR, and FRF diets. White and black bars below the diet indicate light (rest) and dark (active) phase. (C) Blood βOHB obtained from the tail of mice subjected to AL, CR, and FRF diet (n=4 per time point). (D) Serum NEFA levels measured in the tail blood of AL, CR, and FRF mice (n=3-4 per time point). (E) Oil Red O staining of liver samples from mice on AL, CR, and FRF regimen. n=3 per time point per diet. Image magnification = 40x. Feeding period for CR and FRF mice is indicated by green bar. All data represented as Mean ± SD. For Fig 6C-D, Two-way ANOVA with Tukey’s correction for multiple comparisons was performed. Letters indicate statistically significant effect of the diet; a – AL versus CR, b – AL versus FRF, c – CR versus FRF. Statistical significance was set at p<0.05. Light was turned on at ZT0 and light was turned off at ZT12. Grey bars indicate dark phase of the day. AL is represented by blue solid lines, CR is represented by red solid lines, and FRF is represented by purple solid lines.

Next, we assayed for blood β-hydroxybutyrate (βOHB) and NEFA. In agreement with our expectations, CR and FRF blood βOHB and NEFAs were high compared to AL before the feeding, reduced after the feeding and increased again with time without food. Blood NEFAs were comparably induced in both diets, but in CR mice, NEFAs demonstrated a tendency to be induced with slightly faster kinetics (Figure 6D). Starting 6 hours after the refeeding, blood βOHB was significantly higher in FRF compared with AL and CR. In CR mice, the blood βOHB significantly increased compared to AL only after 22 hours without food (Figure 6C). Importantly, the difference in blood βOHB kinetics between diets correlated with the expression of genes in β-oxidation and ketogenesis pathways. Thus, while in FRF the kinetics of blood βOHB and NEFA levels closely followed each other, in CR mice blood NEFA and ketones were uncoupled.

We also performed Oil Red O staining of the liver for lipid droplets (LD) accumulation in on mice on all three feeding regimens (Figure 6E, Supplementary Fig 5). AL mice have moderate intensity LD staining across all time points. Mice in the CR group had a low level of LDs at all time points. Mice in FRF group had high LD staining compared with AL and CR before the feeding, at -2 hours without food, and the LD abundance decreased after the refeeding, at 0 hours without food. LDs re-accumulated in FRF liver as early as 10 hours without food, and the abundance continued to increase up to 18 hours without food until it reached a plateau. Again the LD accumulation in the liver was in correlation with changes induced in the expression of TAG biosynthesis and LD biogenesis genes, which were highly induced in FRF compared with AL and CR but lost their rhythmicity, while in CR the same genes were rhythmic but were not induced compared with AL. Altogether these data suggest that CR and FRF differently reprogramed liver lipid metabolism, with CR improved and FRF impaired liver fat metabolism, in agreement with the hypothesis.

### Effects of CR and FRFF on Whole-Body Metabolic rhythms

The observed differences in glucose and lipid homeostasis between CR and FRF prompted an investigation into their effects on whole-body metabolism using the Comprehensive Lab Animal Monitoring System (CLAMS). The respiratory exchange ratio (RER) was used to estimate the primary nutrient source utilized for energy production. In AL mice, RER exhibited low-amplitude daily oscillations, with values around 1.0 during the dark phase—indicating predominant carbohydrate oxidation—and approximately 0.9 during the light phase, suggesting oxidation of both fatty acids and carbohydrates (Figure 7A). Two hours before food availability, both CR and FRF mice had RER values near 0.78, reflecting predominant fatty acid oxidation. Upon refeeding, RER in FRF mice rapidly increased to ∼0.97, consistent with a switch to carbohydrate oxidation and to approximately 1.1, indicating active lipogenesis, in CR. RER started to decline 4 hours without food in FRF and CR, indicating a transition from carbohydrates to fat oxidation. FRF mice had switched to predominantly fatty acid oxidation after 8 hours after food withdrawal and CR mice after 14 hours without food. Thus, both diets induced a robust shift from carbohydrate to fat oxidation with duration of fasting, but this transition was faster in FRF mice compared with CR mice.

**Figure 7.**
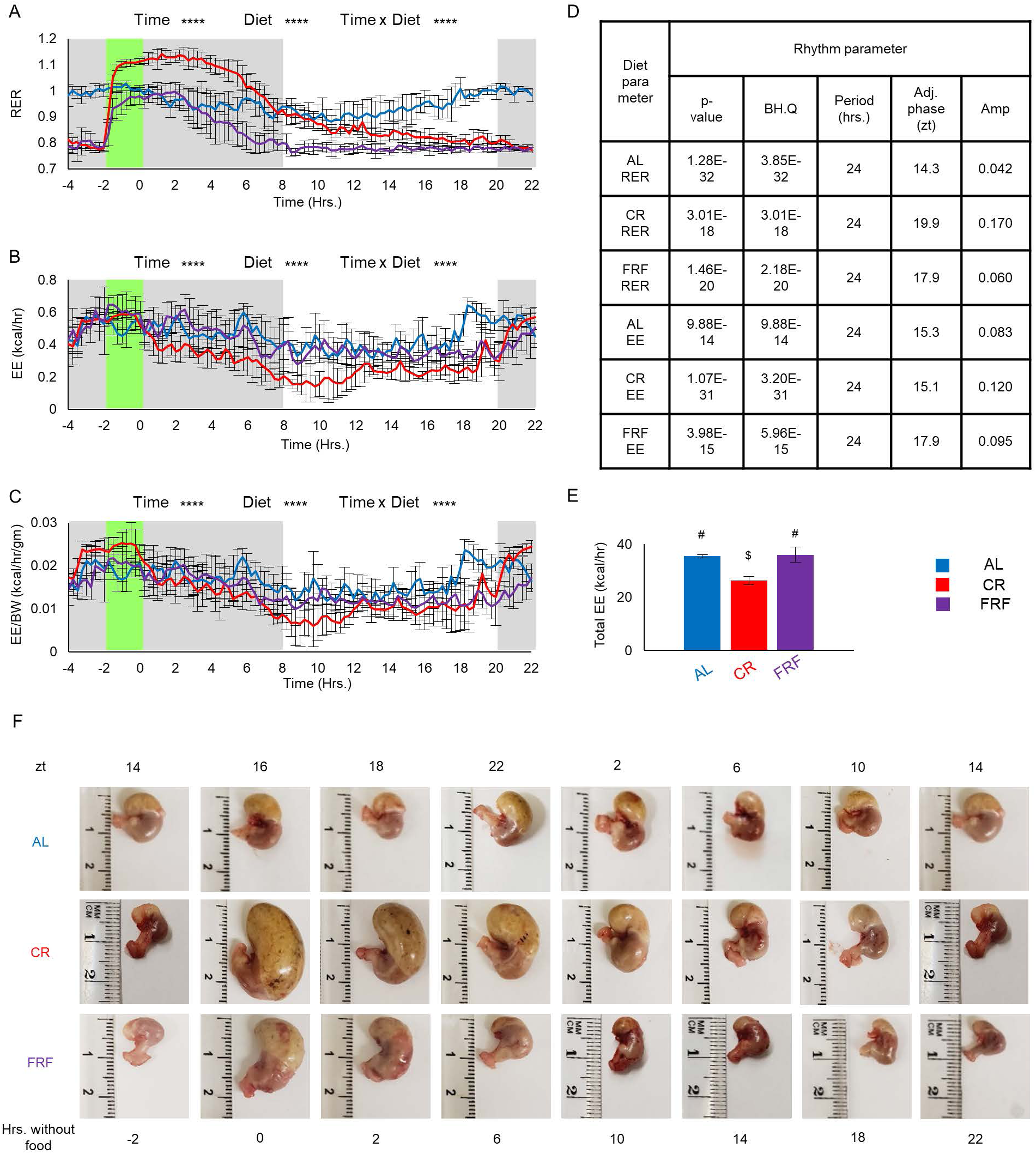
CR uncouples gastric emptying from systemic metabolic transitions. (A-C) Respiratory exchange ratio (RER) and energy expenditure (EE) measured by indirect calorimetry (CLAMS-HC). EE is displayed as (B) EE (kcal/hr), and (C) mass-specific EE (kcal/hr/gm), calculated as EE divided by body weight. For AL group, n=4; CR group, n=6; FRF group, n=6. Data are plotted at 15-minute intervals (X-axis shown in hours). −2 hours indicate time of food presentation to CR and FRF mice (ZT14). Feeding period for CR and FRF mice is indicated by the green bar; grey bars indicate the dark (active) phase. Mice were acclimated in metabolic cages for 3 days prior to data collection. Two-way ANOVA with Tukey’s correction for multiple comparisons was performed; significance set at p<0.05. (D) Rhythm analysis of RER and EE in AL, CR, and FRF regimen. Analysis was performed using the JTK_Cycle parameter in the MetaCycle package in R studio software (version 4.5.1). Statistical significance was set at p<0.05. (E) Total Energy expenditure of mice on AL, CR, and FRF diet (AL, n=4; CR, n=6; FRF, n=6). (F) Stomach weight of mice maintained on AL, CR, and FRF regimen (n=3 per diet per time point). Feeding period for CR and FRF mice is indicated by green bar. All data represented as Mean ± SD. For Fig 7E, different symbols indicate statistically significant effect of the diet, whereas the same symbols indicate no significant effect of the diet. Statistical significance was set at p<0.05. Light was turned on at ZT0 and light was turned off at ZT12. Light and Dark bars indicate light and dark phases of the day. AL is represented by blue bars and blue solid lines, CR is represented by red bars and red solid lines, and FRF is represented by purple bars and purple solid lines.

Energy expenditure (EE) in AL mice was higher during the dark (active) phase than during the light (resting) phase as expected (Figure 7B). FRF did not significantly alter EE compared with AL mice; both the dark/light pattern and absolute EE values were comparable between the two groups (Figure 7B). CR mice displayed a distinct EE profile. EE peaked during the 2-hour feeding window, declined rapidly until 8 hours after feeding, and subsequently began to increase again, coinciding with the onset of the light phase. Overall, EE was significantly lower in CR mice than in AL or FRF mice (Figure 7B). CR mice have reduced body weight (Figure 1) and after normalization of EE data to body weight, the absolute values were more comparable between the diets (Figure 7C). Interestingly, the EE reduction in CR mice was consistent with their decreased caloric intake, whereas EE in FRF mice was comparable with AL despite markedly reduced food consumption, suggesting that EE was not necessary directly related to the food intake. We also measured the circadian rhythmicity of RER and EE in all three feeding regimens using MetaCycle. Both RER and EE exhibited significant 24-hour rhythms under all three diets (Figure 7D). In AL mice, RER rhythms peaked at ZT14.3, whereas EE peaked at ZT15.3. FRF mice had their RER and EE rhythms peaked at ZT17.9 respectively. In CR mice, EE rhythms peaked at ZT15.1, which was similar to AL mice, but RER rhythms peaked at ZT19.9. Thus, FRF caused a phase-shift of both RER and EE by about 3 hours but the phase difference between RER and EE was about the same (<1 hour) in both diets. This close alignment of RER and EE phases in FRF mice suggests coordinated re-timing of substrate utilization and energy expenditure. In CR, the phase difference between RER and EE was almost 5 hours suggesting that, in CR, substrate utilization is more strongly influenced by the scheduled feeding window, whereas energy expenditure remains under the control of active/rest cycle.

The dynamics of stomach content and chyme weight over time are shown in Figure 7F. In AL mice, stomach size and chyme weight remained relatively constant throughout the day, consistent with their continuous feeding behavior.^21,22^ In FRF mice, stomach size and chyme content declined progressively during fasting, and by 6 hours without food, these mice already had significantly less chyme than AL mice. In CR mice, stomach size and chyme weight also decreased with fasting but with much slower kinetics. The stomach reached AL size only after 18 hours without food and became smaller than that of AL mice after 22 hours of fasting (Supplementary Figure 6D-E). Thus, stomach emptying occurred rapidly in FRF mice: even at 0 hours, immediately after consuming ∼ 1.7 g of dry food, the chyme weight was only ∼800 mg, suggesting that most food had already passed from the stomach. In contrast, digestion in CR mice was markedly slower, potentially reflecting metabolic adaptations to chronic calorie restriction.

## Discussion

The global rise in obesity and metabolic disease has increased interest in dietary interventions that incorporate fasting regimens to regulate body weight and metabolism. Caloric restriction (CR), time restricted feeding (TRF), and intermittent fasting (IF) all share a periodic fasting component, yet their physiological and metabolic consequences are not the same in experimental models^21,22,27,29,30,43–45^ and human studies.^17,18,46–50^ Fasting is an integral component of one-meal-per-day CR and contributes significantly to its longevity-associated benefits.^24,27,34^ Therefore, understanding how feeding patterns, reduced caloric intake, and fasting period interact to shape systemic metabolism is important. In this study, we directly compared chronic one-meal-a-day CR with an acute fasting-refeeding-fasting (FRF) and demonstrate that although the diets share a fasting component fasting cannot induce all metabolic effects of CR, and anticipation/entrainment component of CR in addition to reduced caloric intake, were key to metabolic adaptation.

CR improved glucose tolerance and fatty acid utilization at the systemic level, which is consistent with previous studies.^21–23,25–27,29,30^ In contrast, FRF mice showed impaired glucose tolerance and increased hepatic lipid accumulation despite having comparable fasting duration. Liver transcriptomic analyses further highlighted this distinction. Around half the genes that were differentially expressed in CR were also affected by FRF, thus indicating that the fasting component may be contributing substantially to the transcriptional reprogramming. Yet there were more genes that were uniquely regulated by each diet. Another notable distinction between the two diets was that CR increased the number of rhythmically expressed genes, which is consistent with earlier reports that CR robustly regulates circadian transcriptional output,^24,39,42^ whereas FRF disrupted rhythmic gene expression. CR enhanced the number of rhythmically expressed transcripts, whereas FRF disrupted the circadian transcriptome. Circadian rhythms play a critical role in maintaining metabolic homeostasis by temporally coordinating gene expression, signaling pathways, and physiological processes. Disruption of circadian rhythms is associated with adverse metabolic outcomes, whereas strengthening these rhythms enhances metabolic health. The metabolic changes induced by CR and FRF were consistent with that model. The circadian clock, which governs rhythmic physiological processes throughout the body, likely contributed to these differences but cannot fully account for the metabolic improvements observed under CR or the impairments observed under FRF. Indeed, certain metabolic benefits of time-restricted feeding and CR have been documented even in circadian clock–deficient mutants. Interestingly, while fasting disrupted circadian rhythms in the liver, it did not abolish circadian rhythms in energy expenditure: FRF mice still displayed elevated energy expenditure at the onset of the next dark phase, similar to AL-fed mice, despite the absence of feeding.

A striking feature of mechanistic distinction between CR and FRF was the temporal coordination of metabolic transition from fed to fasting and kinetics of gastric emptying. Metabolic responses to refeeding differed between the two dietary diets. In both CR and FRF mice, refeeding induced insulin secretion, upregulated hepatic mTOR signaling, and reduced plasma NEFAs. However, blood glucose increased upon refeeding in FRF mice but not in CR mice. Interestingly, in both dietary regimens, blood glucose levels did not correlate with changes in other key energy metabolites such as NEFAs and βOHB. Notably, RER in FRF mice shifted from approximately 0.7 (indicative of fat oxidation) to 1.0 (indicative of carbohydrate oxidation) within 15 minutes of food reintroduction. In contrast, RER in CR mice rose to approximately 1.1, signifying not only a shift to carbohydrate oxidation but also activation of lipogenic pathways. In FRF mice, the stomach was almost completely emptied within six hours of food withdrawal. Gastric emptying was so rapid that even at the 0-hour time point, the residual stomach content in FRF mice was less than 25% of that in CR mice, despite comparable food intake during the preceding two hours. In FRF mice, changes in circulating insulin and β-hydroxybutyrate (βOHB), hepatic mTOR signaling, whole-body respiratory exchange ratio (RER), and expression of fasting-responsive hepatic genes occurred within the first ten hours of fasting. The close correspondence between stomach emptiness and these physiological parameters suggests that gastric emptying served as one of the key triggers initiating metabolic adaptation in FRF mice. In contrast, gastric emptying in CR mice was markedly slower. Importantly, the metabolic alterations in CR mice were not triggered by gastric emptiness. After ten hours without food, stomach weight in CR mice was comparable to that of FRF mice after only two hours of fasting. Even after fourteen hours, stomach content in CR mice remained slightly greater than that of ad libitum (AL) controls and only after eighteen hours did CR mice exhibit stomach weights lower than AL mice and comparable to those of FRF animals (Figure 6 and Supplementary Figure 7). Despite this slow gastric emptying, plasma insulin levels in CR mice decreased within two hours of food withdrawal, mirroring the kinetics observed in FRF mice. Hepatic mTOR signaling, a well-established marker of nutrient sensing and the fed and fasted states, was downregulated after six hours, again paralleling changes in FRF liver. Circulating non-esterified fatty acids (NEFAs) began to rise in both groups after several hours of fasting. The RER in CR mice started to decrease after approximately 4 hours indicating a transition and after 10 hours it was predominant fatty acid oxidation, which was at least 8 hours before the stomach of CR mice became empty.

Collectively, these data indicate that while in FRF mice both fasting and refeeding responses were tightly linked to the physical presence or absence of food in the stomach and followed predictable physiological patterns, CR mice entered a fasted metabolic state after only a few hours without food, even their stomach still contained food. Thus, in CR mice, fasting-related signaling and metabolic reprogramming were not triggered by gastric emptiness. The metabolic responses in CR mice were decoupled from gastric content, suggesting anticipatory and entrained regulatory mechanisms. During fasting, organisms transition from reliance on exogenous nutrient supply to endogenous energy reserves, including hepatic glycogen and adipose-derived fatty acids, necessitating coordinated suppression of anabolic pathways and activation of catabolic programs.^51–55^ Thus, the fasting response necessitates precise coordination among multiple physiological systems, spanning cellular processes, and inter-organ communication. Insulin is a principal regulator of glucose and fatty acid metabolism. During fasting, low circulating insulin levels promote lipolysis of triacylglycerols in adipose tissue resulting in mobilization and release of NEFAs. Fatty acids subsequently become the main source for energy production in fasting. Low insulin also leads to suppression of mTOR activity and switching from anabolic to catabolic processes.^37,52,56–58^ The temporal changes in both CR and FRF mice consistent with this model and transition to fasting state might be driven by the rapid reduction of insulin. The difference between the diets was that in FRF blood insulin followed feeding and stomach emptying. In CR mice, blood insulin increase might be driven by the food in the stomach, but the decrease was independent from blood glucose and stomach content. These results suggested that, in addition to well documented food anticipation, mice on the CR diet also anticipated the prolonged fasting, which is in agreement with recent reports on entrainment of lipid metabolism in CR.^25,29,30^ Consistent with this framework, previous work has demonstrated that fasting and refeeding induce coordinated hepatic adaptations in glycogen utilization, lipid metabolism, and nutrient signaling pathways,^55^ which aligns with our observation that fasting alone does not fully recapitulate the regulated metabolic state observed under CR.

### Limitations of the study

The mechanistic analyses were confined to the liver. Future studies should determine whether similar anticipation/entrainment-dependent mechanisms occur in other metabolic organs such as skeletal muscle, adipose tissue, kidneys, and heart. Although sex-dependent clustering was observed in transcriptomic analyses, our study primarily focused on diet-dependent effects. Further investigation should be directed towards studying sex-specific mechanisms. Gastric emptying might be regulated differently between fed and fasted states.^59–62^ If the same is true for regulation of stomach emptying in CR, it will be a subject of future investigation.

### Concluding remarks

The success of calorie restriction (CR), time-restricted feeding (TRF), and intermittent fasting in treating metabolic syndrome and obesity is well documented. Periodic fasting without intentional caloric reduction has been proposed as an attractive alternative to traditional CR due to its perceived feasibility and adherence in humans.^8,9,47,49,50,63^ However, recent controlled trials have challenged this notion, suggesting that total caloric intake remains a primary determinant of metabolic outcomes.^64–66^ In animal models, both reduced caloric intake and duration of fasting have each been shown to independently contribute to metabolic benefits and lifespan extension.^21,22,24,45^ Consistent with this distinction, FRF, despite aligning feeding windows and extended fasting, did not improve metabolic regulation, enhance rhythmic transcriptome, or improved glucose tolerance, as observed under CR regimen. These findings underscore the importance of considering the synergistic contributions of temporal coordination, reduced caloric intake, and anticipatory/entrainment mechanisms in shaping metabolic adaptations to dietary interventions.

## STAR METHODS

### KEY RESOURCES TABLE

| REAGENT OR RESOURCE | SOURCE | IDENTIFIER |
| --- | --- | --- |
| **Antibodies** |  |  |
| Phospho-S6 Ribosomal Protein (Ser235/236) | Cell Signaling Technology | Cat# 4858S |
| Ribosomal Protein S6 Antibody (C-8) | Santa Cruz | Cat# sc-74459 |
| β-actin Monoclonal Antibody | Sigma | Cat# A5441 |
| Anti-rabbit IgG, HRP-linked Antibody | Cell Signaling Technology | Cat# 7074 |
| Anti-mouse IgG, HRP-linked Antibody | Cell Signaling Technology | Cat# 7076 |
| **Experimental Models: Organisms/Strains** |  |  |
| C57BL/6J mice | Jackson Laboratory | 000664 |
| **Chemicals, Peptides and Recombinant Proteins** |  |  |
| D-Glucose | Fisher | Cat# D16-1 |
| Methanol | Fisher | Cat# A412-4 |
| Tris Base | Fisher | Cat# BP152-5 |
| Glycine | Fisher | Cat# BP381-5 |
| EDTA | BioBasic | Cat# EB0436 |
| SDS | BioBasic | Cat# SB0485 |
| MOPS Free Acid | BioBasic | Cat# MB0360 |
| Ponceau S Solution | Sigma | Cat# P7170-1L |
| Bovine Serum Albumin | RPI Corp | Cat# A30075 |
| Great Value Instant Nonfat Dry Milk powder | Walmart | N/A |
| Clarity Western ECL Substrate | Bio-Rad | Cat# 1705061 |
| Bovine Gamma globulin, Standard, 2mg/ml | Thermo Scientific | Cat# 23212 |
| Sodium Chloride | Fisher | Cat# S671-3 |
| PageRuler™ Prestained protein ladder | Thermo Scientific | Cat# 26616 |
| NuPAGE™ Bis-Tris Mini Protein Gels 4-12% | Thermo Scientific | Cat# NP0323BOX |
| Bio-Rad Protein Assay Dye | Bio-Rad | Cat# 5000006 |
| Oil Red | Sigma-Aldrich | Cat# O0625 |
| **Critical Commercial Assays** |  |  |
| Ultra-Sensitive Mouse Insulin ELISA Kit | Crystal Chem | Cat# 90080 |
| Free Fatty Acid Assay Kit | Sigma-Aldrich | Cat# MAK466 |
| RNeasy Mini Kit | Qiagen | Cat# 74104 |
| **Software and Algorithms** |  |  |
| GraphPad Prism 8 | GraphPad | N/A |
| Image Studio Lite for western blots | LI-COR Biosciences | N/A |
| CI-Link (Version 1.13.0) | Columbus Instruments | Oxymax-CLAMS-HC |
| **Deposited Data** |  |  |
| RNA Sequencing data | Gene Expression Omnibus | GSE319761 |
| **Other** |  |  |
| Formulab Diet | Lab Diet | Cat# 5008 |
| Blood Glucose Meter | CVS | Item# 968574 |
| Blood Glucose Test strips | CVS | Item# 749202 |
| Precision Xtra Blood Glucose and Ketone Meter | Abbott | Item# 98814-65 |
| Precision Xtra Blood Ketone Test Strips | Abbott | Lot# 75001 |
| Comfort EZTM Insulin syringes | Med-Vet International | Cat# PH-98302013920 |
| Odyssey® Fc Imaging System | LI-COR Biosciences | N/A |
| XCell SureLock Mini-Cell Electrophoresis System | Invitrogen™ | Cat# EI0001 |
| PowerPac Universal Power Supply | Bio-Rad | Cat# 1645070 |
| 5200 Fragment Analyzer | Agilent | Cat# M5310AA |
| Nanodrop 2000 | Thermo Scientific | N/A |

### CONTACT FOR REAGENT AND RESOURCE SHARING

Requests for further information or reagents should be directed to the Lead Contact, Roman V. Kondratov.

### EXPERIMENTAL MODEL AND SUBJECT DETAILS

#### Mice

All experiments involving animals were conducted in accordance with Federal and University guidelines. All procedures were approved by IACUC at Cleveland State University. C57BL/6J mice were bred in-house at Cleveland State University. Mice were maintained on 12 h light: 12 h dark cycle (LD12:12) with lights on at 7am and lights off at 7pm. Animals were housed in groups of three-four animals per cage (Micro-VENT System Caging, #PC7115HT; Overall dimension: 7 3/4”W x 12”D x 6 1/2”H, Allentown, NJ) throughout the experiment. The temperature for the animal rooms was maintained at 20 ± 5^0^C and humidity between 30-70%. All mice were fed 5008 LabDiet (St. Louis, MO), consisting of 26.5% of energy from protein, 16.9% from fat and 56.5% from carbohydrate. Mice used in the experiments were 12-13 weeks of age at the start of the experiments.

### METHODS

#### Study design

Prior to the start of the experiment, all mice were fed ad libitum. At 12-13 weeks of age, mice were randomly assigned to two groups. One group of mice continued on ad libitum diet (AL), while the second group of mice were subjected to 30% caloric restriction (CR). In the CR group, food restriction was introduced gradually: a 10% reduction was implemented during the first week, followed by a 20% reduction during the second week, and finally a 30% reduction until the end of experiment. The CR group received food once per day at ZT14. For the unanticipated Fasting (F) experiment, AL mice continued to receive ad libitum amount of food until 24 weeks of age, after which they were split into two groups. One group continued on the AL diet, the second group had food access unexpectedly withdrawn at ZT16 (0 hours), and fasted until ZT14 (22 hours). For Fast-refeed-fast (FRF) experiments, mice were fed an AL diet until 24 weeks of age, after which mice were split into two groups: one group of mice remained on AL diet, the second group of mice had their food access removed at ZT16 and fasted for 22 hours. At ZT14, mice in the second group were fed AL amount of food for 2 hours until ZT16, after which, access to food was removed again for 22 hours. Mice in all experiments had unlimited access to water. Body weight and other physiologic parameters (blood glucose, ketone, NEFA, insulin) were collected at ZT16 (0 hours) and ZT14 (22 hours) for Fasting (F) experiments, and every four hours across the day for Fast-refeed-fast (FRF) experiments. Liver tissues were also collected at times mentioned above for both experiments and immediately frozen and stored at -80°C.

#### Blood glucose and β-hydroxybutyrate measurement

Blood for the glucose and β-hydroxybutyrate measurement was collected through the tail vein nick. Blood Glucose was measured using CVS Advanced Health Blood Glucometer (CVS Pharmacy, Woonsocket, RI) with CVS Health Advanced Glucose Meter Test Strips (CVS Pharmacy, Woonsocket, RI). Blood ketones were measured as β-hydroxybutyrate level using Precision Xtra Blood Glucose and Ketone meter with Precision Xtra Blood Ketone Test Strips (Abbott Laboratories, IL).

#### Intraperitoneal glucose tolerance tests

After 7-8 weeks on the diets, CR mice were subjected to an intraperitoneal glucose tolerance test (GTT). CR mice received their last meal at ZT14 the previous day and were fasted for 18 hours (ZT10). AL mice were used as non-fasted controls. The FRF group of mice were also fed at ZT14 the previous day, had their food access withdrawn after 2 hours at ZT16, and were fasted for the same duration as CR mice. At ZT10, mice in all three groups were intraperitoneally injected with glucose (1.2 mg/kg body weight). Blood glucose was measured via tail vein at 0, 15, 30-, 60-, 90- and 120-minute post-injection.

#### Plasma insulin assay

Blood was collected through the tail vein nick and immediately mixed with EDTA to prevent coagulation. Samples were centrifuged at 6800 rpm for 20 minutes at 4^0^C, and supernatant was used for the plasma insulin analysis. Plasma insulin was measured using commercially available mouse ultra-sensitive ELISA kit (Crystal Chem Inc., Downers Grove, IL) as per manufacturer’s instructions.

#### Serum NEFA assay

Serum NEFA was measured by colorimetric assay using the Free Fatty Acid Assay kit (Millipore Sigma, Burlington, MA) according to manufacturer protocol.

#### Oil Red O Staining

Mouse liver was perfused with 1× PBS and fixed in 10% buffered formalin. Samples were embedded in OCT and flash frozen. Samples were cut onto Superfrost plus microscopic slides (Fisher Scientific, Hampton, NH) using a cryostat at 10μm thickness. Sample slides were immersed in several solutions as follows: H2O (30 s), H2O (30 s), 60% isopropanol (30 s), Oil Red O for 18 min (Oil Red O powder was purchased from Sigma-Aldrich, prepared in 250 mL of isopropanol, after stirring for 10 min, 150 mL of H2O was added, left to stand for 10 min, then filtered), 60% isopropanol (30 s), 2× wash in H2O (30 s), hematoxylin (2 min) (Sigma-Aldrich, Burlington, MA), ammonium hydroxide (10 dips) (Fisher Scientific, Hampton, NH), and 3× wash with H2O (30 s). Coverslips were placed on the slides using aqua mount (Epredia), and the stained sections were imaged and quantified by assessing the percentage of red stain to the total area of the image using ImageJ software.

#### Whole body metabolism by indirect calorimetry

Respiratory exchange ratio (RER), Energy expenditure (EE), VO2, and VCO2 were determined using Comprehensive Laboratory Animal Monitoring System (CLAMS-HC, Columbus Instruments, Columbus, OH). Prior to metabolic measurements, mice on all three feeding regimens were acclimated in metabolic cages for 3 days, after which data was recorded using CI-Link software (version 1.13.0). AL mice had constant access to food in these cages; CR mice were fed daily at ZT14. FRF mice had their food access removed at ZT16, fasted for 22 hours, fed between ZT14-16, and food removed again for 22 hours. Throughout the experiment, mice in all three groups had constant access to water.

#### RNA-sequencing

Total RNA was extracted from 30 mg of liver tissue using RNeasy Mini Kit (Qiagen, Germantown, MD) according to manufacturer protocol. Concentration was determined by NanoDrop 2000 (Thermo Fisher Scientific, Waltham, MA). Four biologic samples per time point for AL, CR, and FRF groups were sent for mRNA-sequencing to Novogene (Sacramento, CA, USA). RNA integrity was verified by Novogene on 1% agarose gel, purity – on NanoPhotometer (IMPLEN, CA, USA), quantity and integrity - using RNA Nano 6000 Assay Kit of the Bioanalyzer 5400 (Agilent Technologies, CA, USA). 1 μg of RNA per sample was used as an input and sequencing library was prepared using NEBNext Ultra™ II RNA Library Prep Kit for Illumina (New England BioLabs, Ipswich, MA). mRNA was poly-A selected from total RNA using poly-T oligo-attached magnetic beads. Fragmentation was performed using divalent cations under high temperature in NEBNext First Strand Synthesis Reaction Buffer (5X). First strand cDNA was synthesized using random hexamer primer and M-MuLV Reverse Transcriptase (RNase H). Second strand cDNA synthesis was subsequently performed using DNA Polymerase I and RNase H. Remaining overhangs were converted into blunt ends by exonuclease/polymerase reaction. After adenylation of 3’ ends of DNA fragments, NEBNext Adaptor with hairpin loop structure was ligated to prepare for hybridization. The library fragments were purified with AMPure XP system (Beckman Coulter, Beverly, USA) in order to select cDNA fragments of preferentially 150∼200 bp in length. 3 μl USER Enzyme (New England BioLabs, Ipswich, MA) was incubated with size-selected, adaptor ligated cDNA at 37 °C for 15 min followed by 5 min at 95 °C before PCR. Then PCR was performed with Phusion High-Fidelity DNA polymerase, Universal PCR primers, and Index (X) Primer. PCR products were purified using AMPure XP system. Library quality was assessed on the Agilent Bioanalyzer 5400. The clustering of the index-coded samples was performed on a cBot Cluster Generation System using PE Cluster Kit cBot-HS (Illumina) according to the manufacturer’s instructions. After cluster generation, the library preparations were sequenced on an Illumina NovaSeq Xplus platform, and 50M paired-end reads (∼150bp) per sample were generated.

Subsequent processing was performed in-house using connection to Ohio Supercomputer Owens Cluster Server (Columbus, OH). RNA-Seq reads were mapped to the mouse protein coding genes (Ensembl: Mus_musculus; GRCm38 (mm10)) using Bowtie78 allowing up to 2-mismatches. The gene expected read counts and Transcripts Per Million (TPM) were estimated by RSEM (v1.2.3)79. The TPMs were further normalized by the EBSeq80 R package to correct potential batch effect. The transcriptomics data for this study has been uploaded to the Gene Expression Omnibus Repository (GEO): GSE319761.

#### Bioinformatics analysis

##### Differential Gene Expression Analysis

Differential gene expression (DEG) analysis was performed to compare three experimental conditions: CR vs AL, FRF vs AL, and CR vs FRF. RNA-seq count data for each condition were analyzed using the EBSeq package. Genes were considered differentially expressed if they satisfied the following thresholds: fold change (FC) > 2 for upregulated genes, FC < 0.5 for downregulated genes, and false discovery rate (FDR) < 0.05. EBSeq employs an empirical Bayesian hierarchical model to estimate posterior probabilities of differential expression, allowing robust identification of DEGs across multiple conditions. Volcano plots were generated for each comparison to visualize the distribution of DEGs, with log₂ fold change plotted against the –log_10_ (FDR), highlighting significantly upregulated and downregulated genes.

##### Gene Ontology Pathway Enrichment Analysis

Gene Ontology (GO) enrichment analysis was performed on differentially expressed genes (DEGs) to identify overrepresented biological processes, molecular functions, and cellular components. DEGs from each experimental comparison were analyzed separately. Statistical significance of enriched terms was assessed using false discovery rate (FDR)–adjusted p-values. Enriched GO terms were visualized and interpreted to elucidate the functional implications of gene expression changes under each condition. GO enrichment analysis was performed using the clusterProfiler R package. An adjusted p-value (FDR) threshold of < 0.05 was used to define statistical significance for enriched GO terms and pathways.

##### Visualization of DEG Overlap and Temporal Dynamics

All analyses were performed using R (version 4.1.0) and associated Bioconductor packages. Circos plots were generated in R using the circlize package to visualize temporal expression patterns of differentially expressed genes across multiple time points. Venn diagrams were generated to visualize overlaps among differentially expressed gene sets across experimental conditions using the Venn Diagram R package.

##### Western Blotting

Liver lysates were prepared with Cell Signaling Lysis Buffer (1M Tris Base pH 7.5, 5M NaCl, 0.5M EGTA, 0.5M EDTA, Triton-X, 0.1M Na_4_P_2_O_7_, 1M β-glycerophosphate, 1M Na_3_VO_4_) containing protease and phosphatase inhibitor cocktails (Sigma-Aldrich, Burlington, MA). The homogenates complete with SDS loading mix were loaded in 4-12% Bis-Tris NUPAGE gels (Thermo Fisher Scientific, Waltham, MA). After electrophoretic run, proteins were transferred onto PVDF membrane (Thermo Fisher Scientific, Waltham, MA) and blocked in 5% Milk prepared in TBS with 0.1% Tween-20 for 1 hr. Incubation with primary antibodies was done overnight with gentle shaking at 4^0^C. For the list of antibodies, refer to the Key Resources Table. β-actin was used as an internal control. Quantification of images was done using Image Studio Lite software.

##### Rhythmic analysis

The rhythmic analysis was performed in R software using DryR package. The groups of samples across the time scale of the experiment with n=4 biological replicates per diet per time point were directly compared for rhythmicity between all 3 diets and sorted into rhythmic models. The genes with 0 counts across the samples were omitted from the analysis and BIC weight > 0.4 was used as the cutoff for model fit.

##### Statistical Analysis

Statistical analysis was performed as follows: For Fasting (F) experiments, unpaired Student’s t-test with Welch’s correction was used for within a diet or between two diets. One-way ANOVA with Tukey’s correction for post-hoc analysis was used for comparisons across three diets (AL, CR, and F). For Fast-refeed-fast (FRF) experiments, Two-way ANOVA with Tukey’s correction for post-hoc analysis was performed across three diets (AL, CR, and FRF). All statistical analysis was performed using GraphPad Prism 8.0 (San Diego, CA). Number of biological replicates per time point per diet have been mentioned in the Figure legends. Data are shown as Mean ± SD. A p-value of ≤ 0.05 was considered statistically significant. Different symbols indicate statistically significant effect of the diet, whereas the same symbols indicate no statistical significance. Where indicated, letters (a,b,c) signify statistically significant effect of the diets (a – AL vs CR, b – AL vs FRF, c – CR vs FRF).

## Resource Availability

### Data code Availability

Raw RNA-Sequencing data has been deposited in the Gene Expression Omnibus: GSE319761 and will be available from the lead contact upon request.

This study did not generate any original code.

Any additional information required to re-analyze the data reported in this manuscript will be available from the lead contact upon request.

## Acknowledgments

This work was supported by U.S. National Institutes of Health/National Institute on Aging Grant R01AG039547 and funds from the Center for Gene Regulation in Health and Disease (Cleveland State University) to R.V.K. We thank Dr. Mahesh Ramamoorthy for his valuable insights for our study.

## Supplementary Figures

**Supplementary Figure 1.**
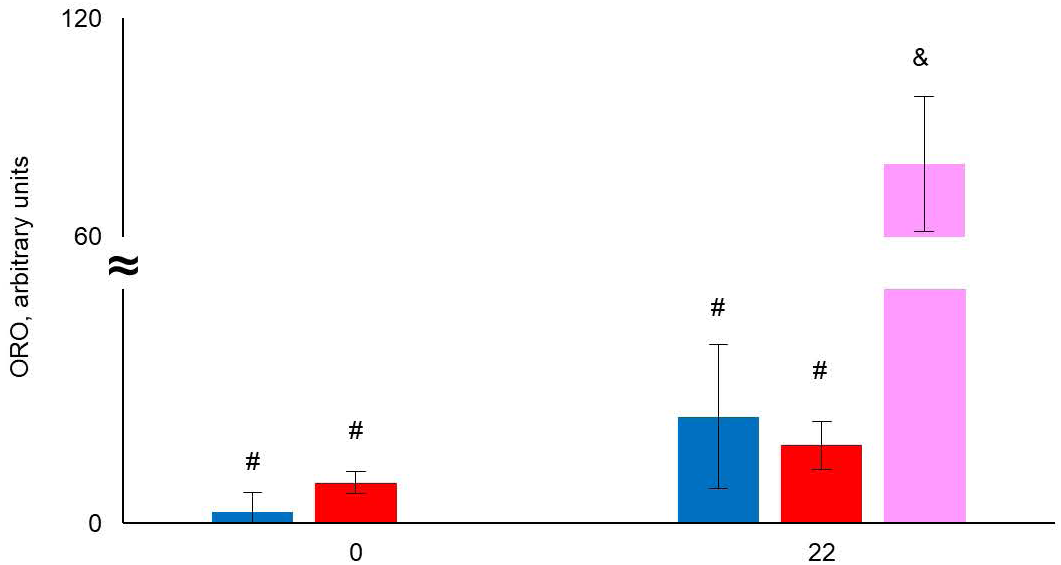
Effect of acute Fasting on lipid droplet accumulation. Quantification of Oil Red O staining from mice on AL, CR and Fasting at 0 hours (ZT16) and 22 hours (ZT14) of time without food. N=3 per diet per time point. All data represented as Mean ± SD. For comparison between diets at 0 hours without food, unpaired t-test with Welch’s correction was used. For comparison between diets at 22 hours without food, One-way ANOVA with Tukey’s correction for post-hoc analysis was used. For comparison within diets at 0 hours and 22 hours without food, unpaired t-test with Welch’s correction was used. Different symbols indicate statistically significant effect of the diet, whereas the same symbols indicate no significant effect of the diet. Statistical significance was set at p<0.05. Light was turned on at ZT0 and light was turned off at ZT12. Light and Dark bars indicate light and dark phases of the day. AL is represented by blue bar, CR is represented by red bar, and Fasting is represented by pink bar.

**Supplementary Figure 2.**
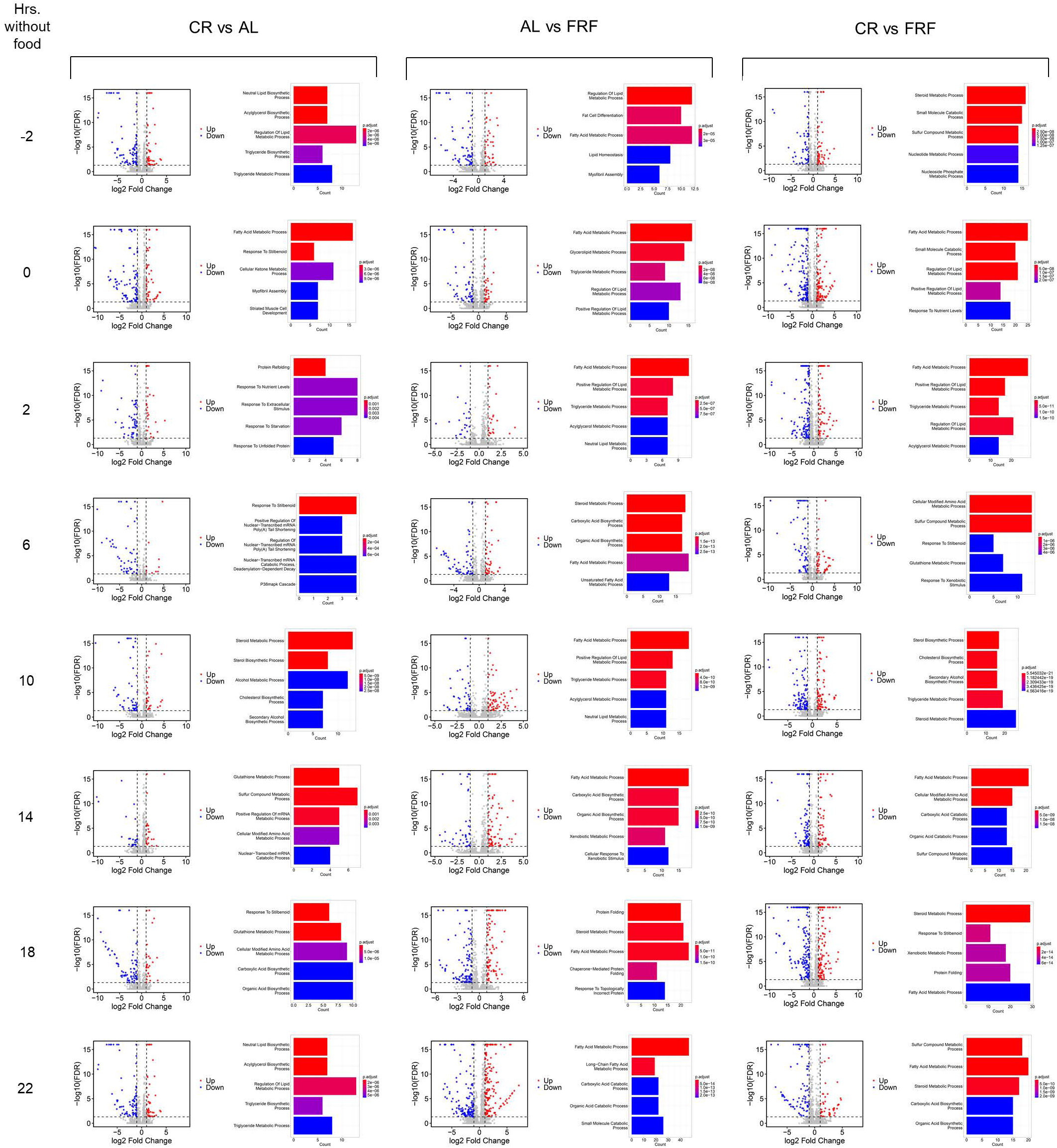
Temporal dynamics of pairwise differential expression under AL, CR, and FRF. Volcano plot and pathway enrichment analysis (using Gene Ontology) for pairwise comparison of CR and AL, FRF and AL, and CR and FRF at each time without food. Blue dots indicate differentially expressed genes that are upregulated whereas red dots indicate genes that are downregulated. Histogram indicates top 5 pathways for the genes in each pairwise comparison.

**Supplementary Figure 3A-C.**
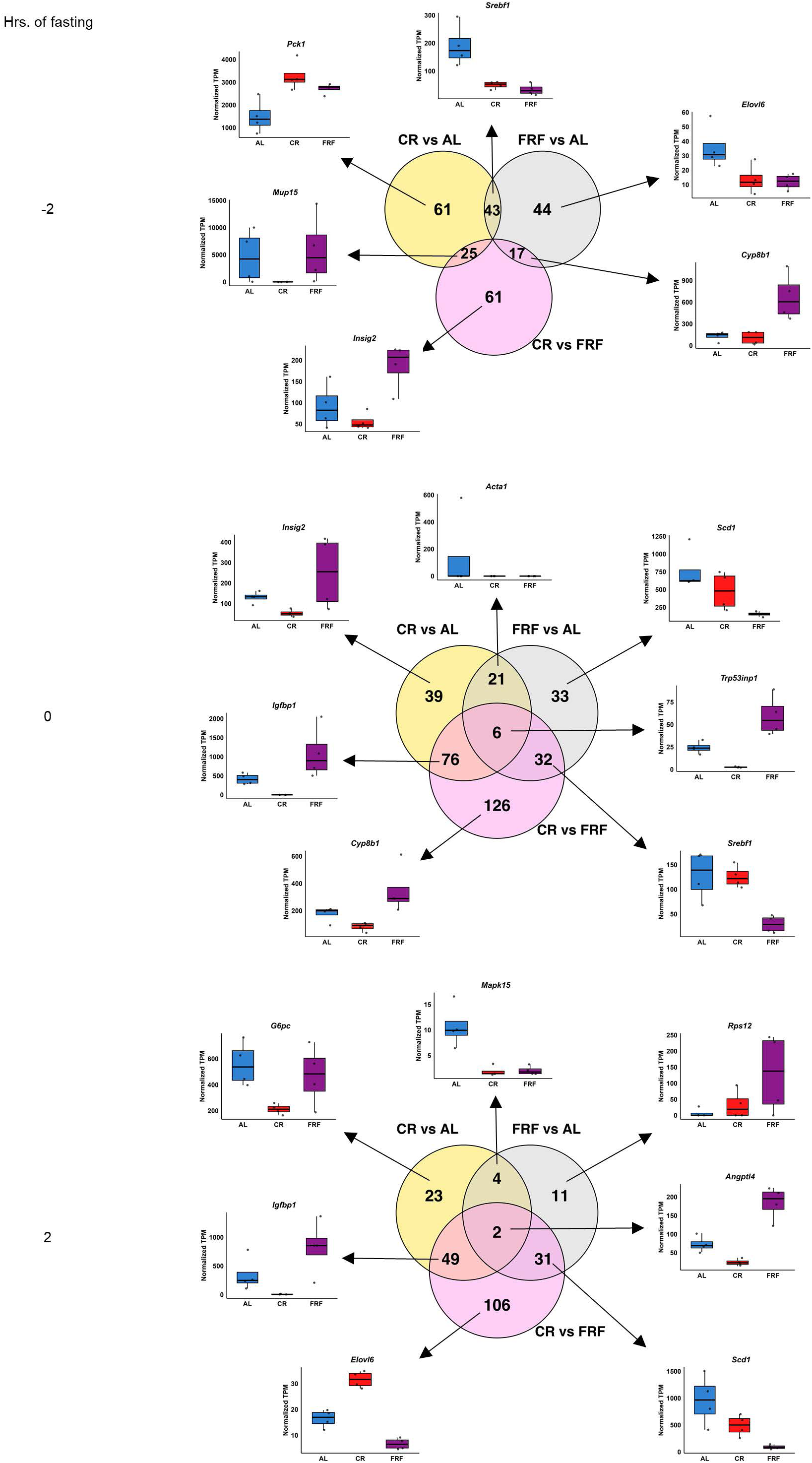
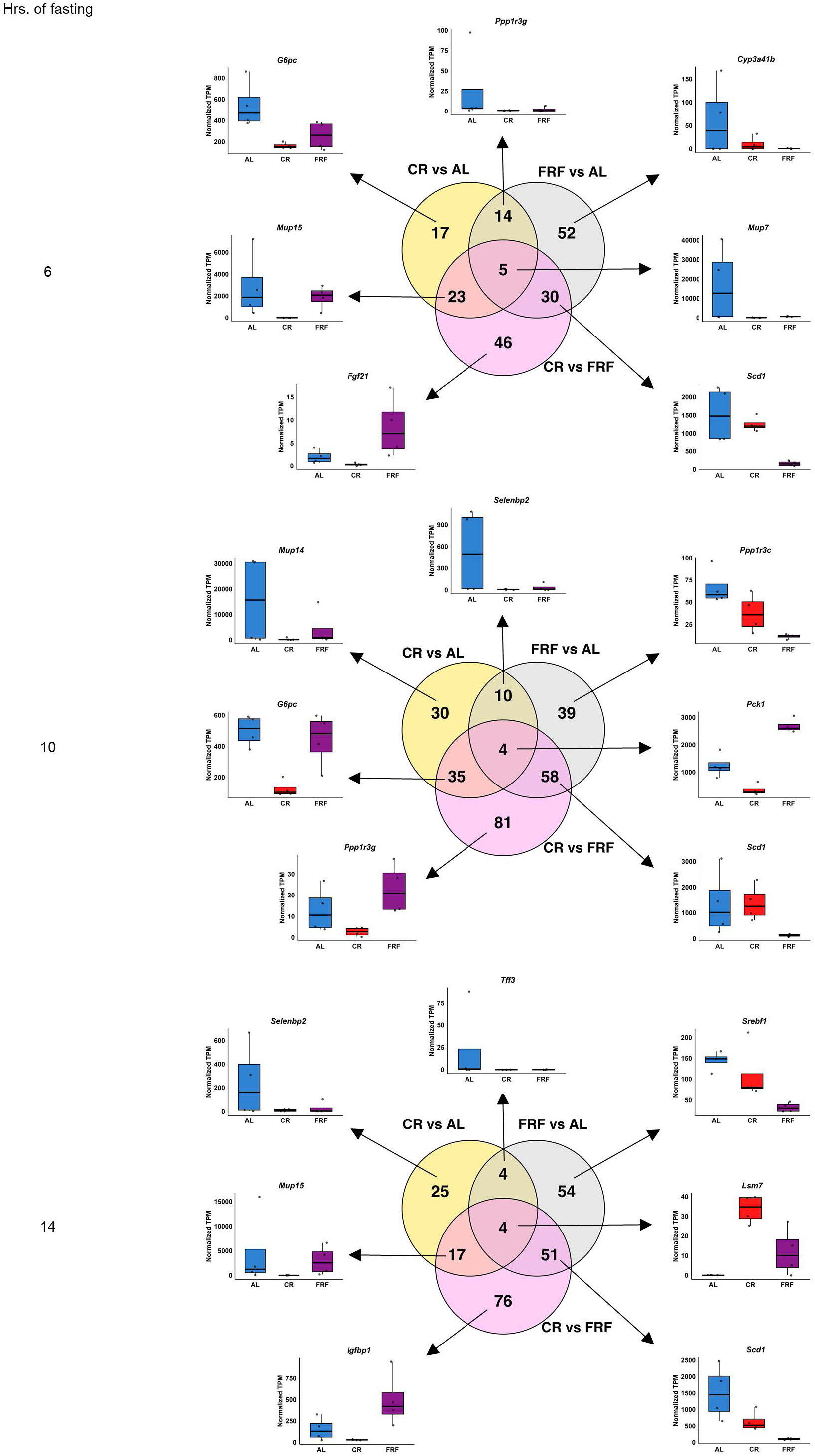
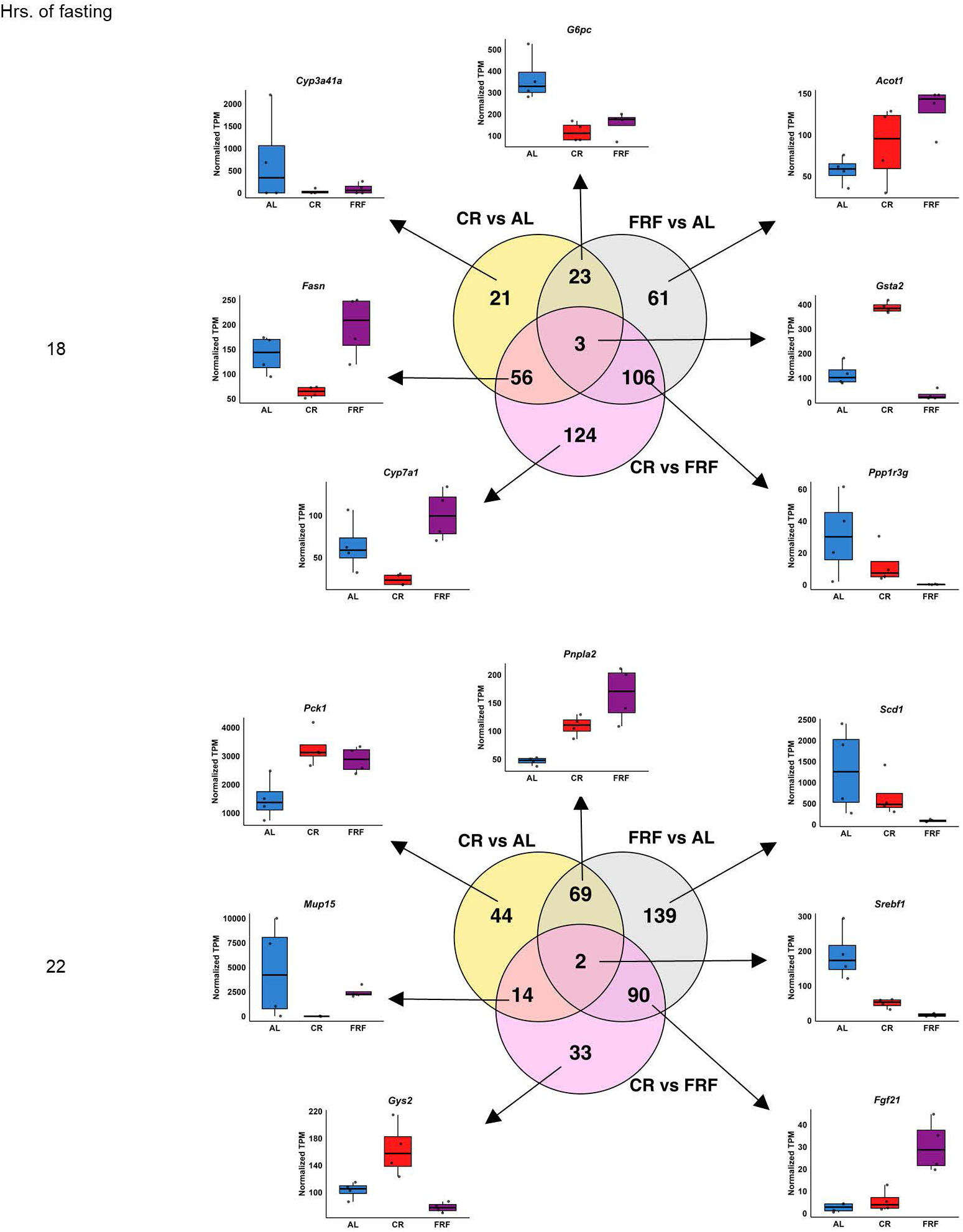
Temporal distribution of shared and diet-specific DEGs across fasting duration. Venn diagram analysis shows distribution of differentially expressed genes when all three diets (AL, CR, FRF) are compared. N=4 per diet per time point. Representative examples for each group are also illustrated at every time point in the Supplementary Figures 3A-C.

**Supplementary Figure 4.**
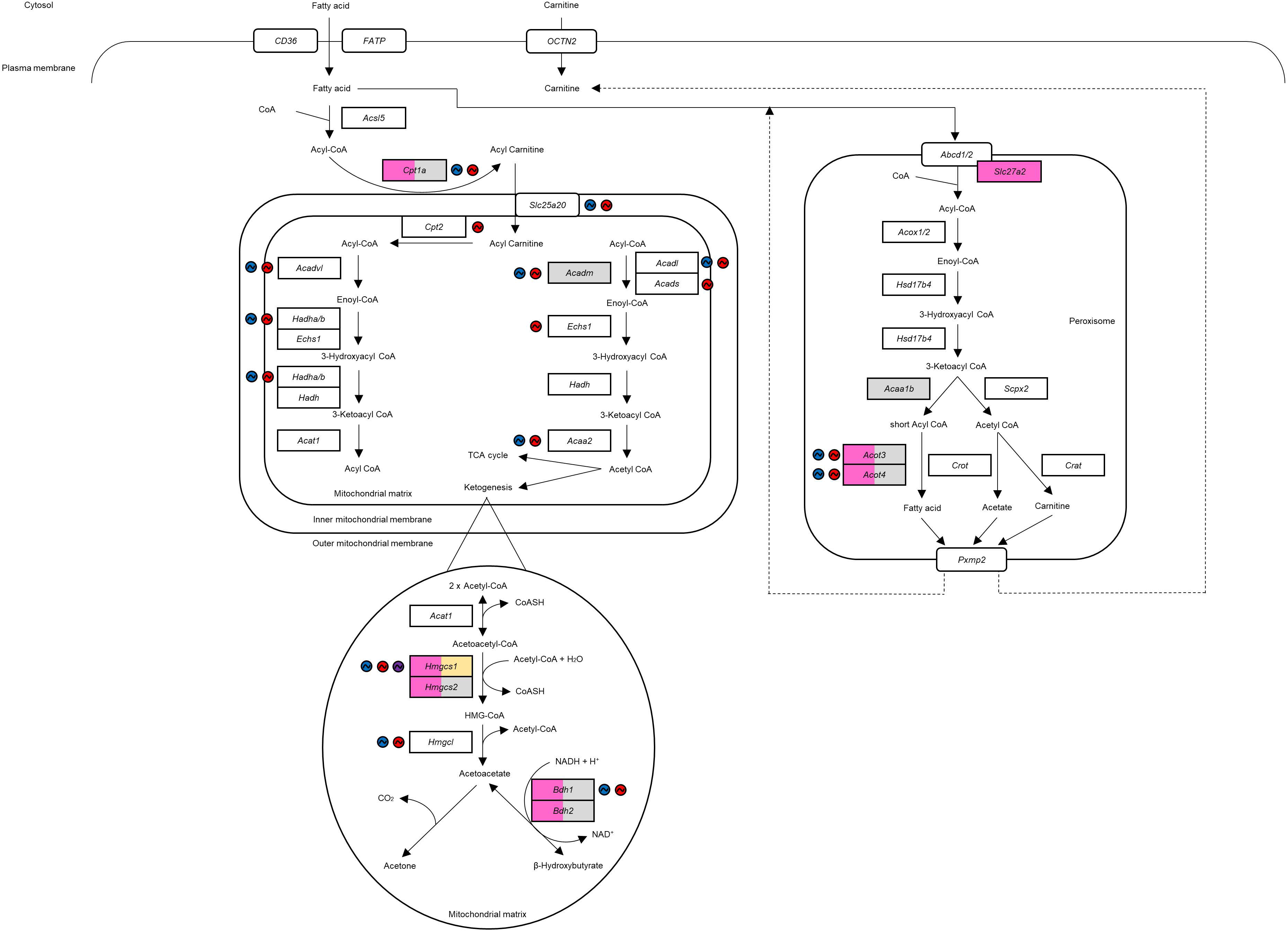
Differential and rhythmic regulation of key lipid metabolism enzymes under CR and FRF. Schematic representation of lipid metabolism pathways. Colored boxes indicate differential expression of enzymes in each pairwise comparison. Colored circles with sinusoid wave represent the rhythmicity of enzymes in respective diets.

**Supplementary Figure 5.**
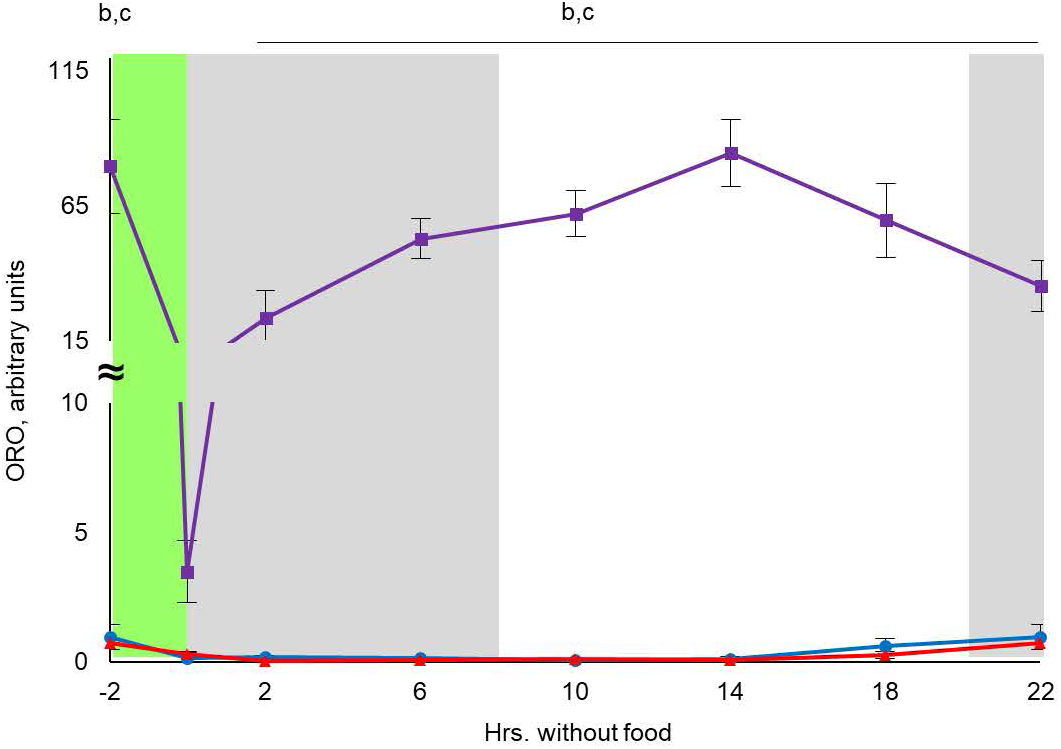
Temporal dynamics of hepatic lipid accumulation between CR and FRF. Quantification of Oil Red O staining from mice on AL, CR and FRF across time without food. N=3 per diet per time without food. The time of the day when the food was provided for CR and FRF mice is indicated by green bar. All data represented as Mean ± SD. Two-way ANOVA with Tukey’s correction for multiple comparison was performed. Letters indicate statistically significant effect of the diet; a – AL versus CR, b – AL versus FRF, c – CR versus FRF. Statistical significance was set at p<0.05. Light was turned on at ZT0 and light was turned off at ZT12. Grey bars indicate dark phase of the day. AL is represented by blue solid lines, CR is represented by red solid lines, and FRF is represented by purple solid lines.

**Supplementary Figure 6.**
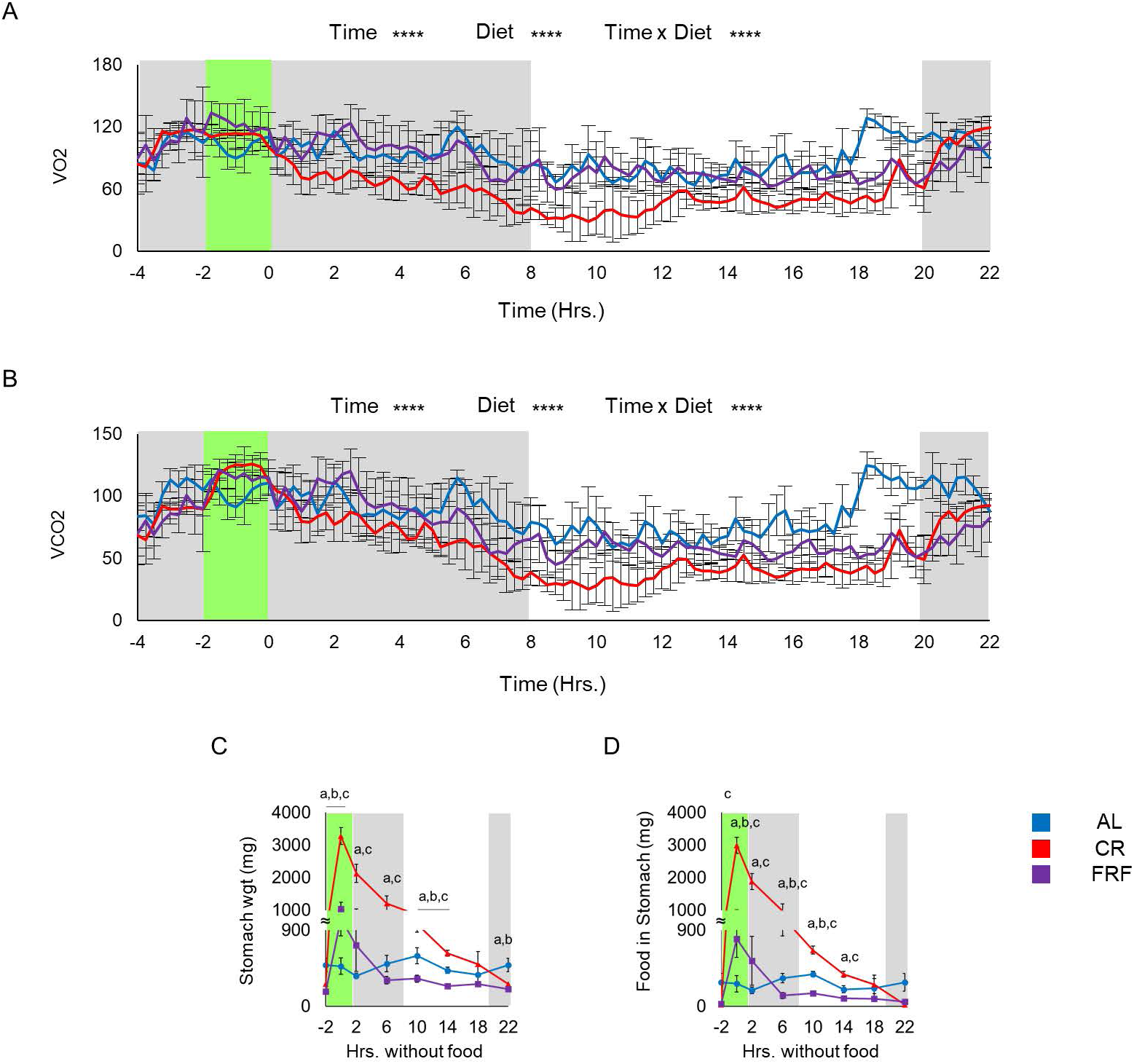
Diet-dependent differences in VO_2_, VCO_2_ and gastric emptying kinetics. (A-B) Volume of O_2_ and CO_2_ for each feeding group measured by indirect calorimetry. For AL group, n=4; CR group, n=6; FRF group, n=6. Data are plotted at 15-minute intervals (X-axis shown in hours). −2 hours indicate time of food presentation to CR and FRF mice (ZT14). Feeding period for CR and FRF mice is indicated by the green bar; grey bars indicate the dark (active) phase. Mice were acclimated in metabolic cages for 3 days prior to data collection. Two-way ANOVA with Tukey’s correction for multiple comparisons was performed; significance set at p<0.05. (C-D) Quantification of stomach weight and amount of food retained in the stomach (n=3 per diet per time point) of mice subjected to AL, CR, and FRF diet. Feeding period for CR and FRF mice is indicated by green bar. All data represented as Mean ± SD. Two-way ANOVA with Tukey’s correction for multiple comparisons was performed. Letters indicate statistically significant effect of the diet; a – AL versus CR, b – AL versus FRF, c – CR versus FRF. Statistical significance was set at p<0.05. Light was turned on at ZT0 and light was turned off at ZT12. Grey bars indicate dark phase of the day. AL is represented by blue solid lines, CR is represented by red solid lines, and FRF is represented by purple solid lines.

## Supplementary Tables

**Supplementary Table 1.** List of differentially expressed genes for time-independent pairwise comparisons between diets.

**Supplementary Table 2.** List of differentially expressed genes present in the intersection of pairwise comparison between CR-AL and FRF-AL. Genes in this subset, that are either upregulated, downregulated, or regulated in opposite direction by CR and FRF in comparison to AL are highlighted in green.

**Supplementary Tables 3-10. Related to Supplementary Figure 3A-C**. List of differentially expressed genes for pairwise comparisons between diets at each time point without food.

## Notes

### Competing Interest Statement

The authors have declared no competing interest.

