## Supplementary Table 1 for "Anticipatory metabolic reprogramming distinguishes caloric restriction from fasting-refeeding cycles"

Supplementary Table 1 for Fig 2G-J
Differentially expressed genes (FDR < 0.05)

| CRvsAL Exclusive DEGs | | |
|---|---|---|
| Gene.ID | FDR | FC |
| Bhlhe40 | 5.27578E-13 | 0.474924823 |
| Btg2 | 0 | 0.322231263 |
| C6 | 1.39754E-05 | 0.358375538 |
| C8b | 0.006272673 | 0.481382539 |
| Cela1 | 0.002888977 | 0.404186183 |
| Ces3a | 2.98601E-05 | 0.456136504 |
| Clstn3 | 0.001296143 | 0.453384421 |
| Col5a2 | 6.32029E-05 | 0.438272936 |
| Crygn | 0.000111388 | 0.490982753 |
| Csrnp1 | 4.26581E-09 | 0.476399963 |
| Cyp4a12b | 6.13559E-06 | 0.321864022 |
| Cyp7b1 | 4.93516E-07 | 0.314665942 |
| Cyr61 | 1.62524E-07 | 0.493440018 |
| Egfr | 4.23325E-06 | 0.388621421 |
| Elovl3 | 0.000106941 | 0.305352199 |
| Fitm1 | 0.009407554 | 0.474310122 |
| Fst | 4.22821E-08 | 0.457821439 |
| Il1b | 2.66079E-11 | 0.379481971 |
| Jun | 2.40614E-08 | 0.461413111 |
| Junb | 0.001480579 | 0.469859657 |
| Mup3 | 0.000357064 | 0.323035531 |
| Saa1 | 0.001955842 | 0.421190236 |
| Saa3 | 1.05992E-07 | 0.48287944 |
| Slc25a25 | 9.4309E-07 | 0.463830988 |
| Slc25a30 | 0.009118096 | 0.483327997 |
| Slco1a1 | 7.2659E-06 | 0.327280652 |
| Sln | 3.16671E-05 | 0.491679326 |
| Srd5a1 | 5.79085E-05 | 0.460833489 |
| Tnnt1 | 0.008134894 | 0.46980501 |
| Zfp36 | 0 | 0.439593214 |
| Zfp982 | 0.023584214 | 0.473255411 |
| Aldoc | 2.92211E-13 | 2.048148882 |
| Btbd19 | 0.002825416 | 2.079394916 |
| Cyp3a59 | 4.04183E-05 | 2.11078397 |
| Gstt2 | 0 | 2.026483322 |
| Hspa1b | 0.001236815 | 2.708530358 |
| Lgals4 | 0.000529366 | 2.293699908 |
| Mmp7 | 9.48261E-05 | 2.127188193 |
| Orm3 | 2.36469E-05 | 2.36841855 |

Supplementary Table 1 for Fig 2G-J
Differentially expressed genes (FDR < 0.05)

| CRvsAL Exclusive DEGs | | |
| --- | --- | --- |
| Gene.ID | FDR | FC |
| Pde6c | 0.007945836 | 2.482466031 |
| Rbm3 | 3.42837E-12 | 2.276146244 |
| Reg1 | 4.82886E-05 | 2.476462017 |
| Tceal8 | 1.55373E-09 | 2.368787303 |
| Trib3 | 0.018829307 | 2.118742212 |
| Tuba8 | 0.049940469 | 2.113252604 |

Supplementary Table 1 for Fig 2G-J

Differentially expressed genes (FDR < 0.05)

| FRFvsAL Exclusive DEGs | | |
|---|---|---|
| Gene.ID | FDR | FC |
| Adam11 | 5.89648E-05 | 0.478246696 |
| Adgrf1 | 1.68822E-10 | 0.377196464 |
| Asns | 1.46823E-05 | 0.345272297 |
| Bdh2 | 2.37088E-12 | 0.471563492 |
| Casq1 | 2.43259E-06 | 0.445635037 |
| Cd74 | 0.001495324 | 0.484465495 |
| Cryab | 0.000145559 | 0.359497147 |
| Cxcl1 | 3.66104E-08 | 0.334441596 |
| Cxcl10 | 7.64771E-07 | 0.49000392 |
| Cyp26a1 | 5.90416E-06 | 0.39047342 |
| Des | 0.001997079 | 0.439272739 |
| Dpy19l3 | 4.57877E-05 | 0.434966704 |
| Fam25c | 0 | 0.483862483 |
| G0s2 | 0.000263516 | 0.467522612 |
| Gadd45a | 2.07653E-10 | 0.353685193 |
| Gnat1 | 8.89783E-05 | 0.453543059 |
| H2-Aa | 0.002217588 | 0.498606701 |
| H2-Ab1 | 2.43237E-05 | 0.463939078 |
| H2-Eb1 | 1.08156E-05 | 0.427228782 |
| H2-Q1 | 0.000383619 | 0.382797047 |
| Hectd2 | 5.36459E-05 | 0.406615122 |
| Hes6 | 1.06926E-11 | 0.435114886 |
| Hsd3b2 | 1.63603E-06 | 0.480631037 |
| Hspb7 | 6.86527E-07 | 0.457006449 |
| Ifi47 | 6.68341E-10 | 0.494466424 |
| Insc | 1.96014E-09 | 0.485530938 |
| Jchain | 0.000181694 | 0.360417783 |
| Ldhb | 0.002960836 | 0.430289328 |
| Ly6d | 5.07898E-05 | 0.329949265 |
| Mapk15 | 7.41578E-10 | 0.371717315 |
| Mcm10 | 0.00430653 | 0.460959668 |
| Mgp | 0.000222944 | 0.427673896 |
| Mmd2 | 0.000112923 | 0.361532175 |
| Nat8 | 0.009625056 | 0.468363524 |
| Nat8f1 | 0 | 0.464951243 |
| Rpl3l | 2.30298E-06 | 0.492501717 |
| Rtn2 | 0.000344796 | 0.462520308 |
| Serpinb1a | 0.036230972 | 0.46098252 |
| Slc41a2 | 6.73296E-05 | 0.434154627 |

Supplementary Table 1 for Fig 2G-J
Differentially expressed genes (FDR < 0.05)

| FRFvsAL Exclusive DEGs | | |
|---|---|---|
| Gene.ID | FDR | FC |
| Spsb4 | 2.68893E-11 | 0.3756298 |
| Thrsp | 0.002815884 | 0.466476508 |
| Tmsb10 | 1.6187E-06 | 0.472182421 |
| Tpm2 | 5.52299E-07 | 0.281917207 |
| Zbp1 | 3.11555E-10 | 0.396794922 |
| Abcd4 | 3.75451E-09 | 2.128693375 |
| Acot1 | 9.00027E-07 | 2.659769365 |
| Cdip1 | 0 | 2.222108053 |
| Chd9 | 2.37688E-07 | 2.623639674 |
| Cpsf4l | 2.44393E-11 | 2.362976116 |
| Cyp2b10 | 0.002408369 | 2.649346356 |
| Defb1 | 6.25178E-09 | 3.169230505 |
| Fam134b | 2.25343E-07 | 2.612549043 |
| Fbxo31 | 1.95399E-14 | 2.802601127 |
| Glrx | 0 | 2.213328022 |
| Gpcpd1 | 0.020465913 | 2.172987922 |
| Idi1 | 0.038428714 | 2.083610916 |
| Il1r1 | 8.61556E-09 | 2.230057412 |
| Il6ra | 2.22045E-16 | 2.598990388 |
| Ip6k2 | 0 | 2.399545532 |
| Lpin2 | 4.62345E-06 | 2.089447805 |
| Mfsd2a | 2.38179E-06 | 2.320857533 |
| Pex11a | 1.54575E-09 | 2.140627094 |
| Plin5 | 1.58873E-13 | 2.815534222 |
| Pnpla2 | 2.83107E-13 | 2.131537061 |
| Retsat | 0 | 2.612270253 |
| Slc37a4 | 0 | 2.094275127 |
| Tango2 | 0 | 2.069924057 |
| Zbtb16 | 2.13794E-10 | 2.801246324 |

Supplementary Table 1 for Fig 2G-J
Differentially expressed genes (FDR < 0.05)

| CRvsFRF Exclusive DEGs | | |
|---|---|---|
| Gene.ID | FDR | FC |
| Acss3 | 1.0705E-09 | 0.472171971 |
| Arl4d | 8.86628E-06 | 0.46341284 |
| Atf5 | 0 | 0.484427386 |
| Ccnf | 8.88178E-16 | 0.448734119 |
| Cidec | 8.87981E-05 | 0.407741156 |
| Csrp3 | 2.5256E-10 | 0.472552551 |
| Fabp2 | 0 | 0.492776529 |
| Igfbp1 | 2.88658E-15 | 0.192075976 |
| Mthfr | 7.59802E-07 | 0.489090085 |
| Myom3 | 4.6709E-07 | 0.342987831 |
| Pnliprp1 | 1.07753E-07 | 0.37497653 |
| Serpina3k | 0 | 0.43240696 |
| Slc27a1 | 7.11701E-10 | 0.431397125 |
| Slc45a3 | 0 | 0.422851192 |
| Sult3a2 | 0.000134914 | 0.446585567 |
| Trp53inp1 | 1.00364E-13 | 0.328958355 |
| Tsc22d3 | 1.30227E-10 | 0.499096506 |
| Vnn1 | 2.20179E-10 | 0.434434472 |
| 5830473C10Rik | 0 | 2.242709612 |
| Actg2 | 0.000548804 | 2.095645341 |
| Adssl1 | 0 | 2.016053442 |
| Aqp8 | 0.00022966 | 2.725837881 |
| C330021F23Rik | 0.000750714 | 2.095713416 |
| Ces2a | 8.74856E-14 | 2.111472937 |
| Cldn5 | 3.76008E-09 | 2.220187567 |
| Crip1 | 0.01558729 | 2.105579901 |
| Cyp3a11 | 2.12306E-08 | 2.114909985 |
| Dbp | 0.003342863 | 2.399814948 |
| Ethe1 | 0.004768629 | 2.103907941 |
| Fam129b | 3.36978E-06 | 2.150645193 |
| Fam83f | 8.25778E-05 | 2.137056317 |
| Gm3839 | 0.006784917 | 2.040637366 |
| Gsta4 | 0 | 2.634986961 |
| Gstm1 | 0 | 2.503424989 |
| Gstm6 | 0 | 2.339857167 |
| Hsd17b6 | 1.39146E-05 | 2.150483021 |
| Mgst3 | 0 | 2.208058118 |
| Myl9 | 0.000747874 | 2.287285075 |
| Nanp | 0.00128717 | 2.064854455 |

Supplementary Table 1 for Fig 2G-J
Differentially expressed genes (FDR < 0.05)

| CRvsFRF Exclusive DEGs | | |
|---|---|---|
| Gene.ID | FDR | FC |
| Nr1d1 | 0.004283262 | 2.286966439 |
| Pcolce | 0.000156988 | 2.048985155 |
| Pdzk1ip1 | 4.40547E-05 | 2.447959872 |
| Pltp | 1.39914E-06 | 2.348268776 |
| Prss8 | 9.24889E-09 | 2.420989695 |
| Raet1e | 1.26363E-08 | 2.44684632 |
| S100a6 | 0.006180857 | 2.147345975 |
| S100g | 0.000253492 | 2.138635146 |
| Spp1 | 9.96014E-12 | 2.281877144 |
| Sprr2a2 | 1.20833E-05 | 2.07876985 |
| Sult2a2 | 0.000161015 | 2.820700003 |
| Tagln | 0.010042479 | 2.074496719 |
| Tmem51 | 7.90595E-11 | 2.549855692 |
| Tstd1 | 0 | 2.437173631 |
| Vegfb | 0 | 2.367946121 |

Supplementary Table 1 for Fig 2G-J
Differentially expressed genes (FDR < 0.05)

| CRvsAL Shared DEGs | | |
|---|---|---|
| Gene.ID | FDR | FC |
| Acta1 | 0 | 0.083876673 |
| Ckm | 0 | 0.077898462 |
| Cyp17a1 | 7.48158E-05 | 0.392896684 |
| Mb | 0 | 0.078575126 |
| Selenbp2 | 0 | 0.025833827 |
| Serpina1e | 0 | 0.055526867 |
| Tnnc2 | 0 | 0.083581845 |
| Tnni2 | 2.22045E-15 | 0.125417545 |
| Tnnt3 | 0 | 0.095863441 |
| Cib3 | 0.00103644 | 2.415263678 |
| Cyp4a10 | 1.65167E-10 | 2.009907611 |
| Gm3776 | 8.01306E-06 | 2.728449318 |
| Gsta1 | 8.09216E-06 | 2.769277841 |
| Gsta2 | 1.06026E-12 | 2.479751114 |
| Lpin1 | 0.03115805 | 2.119449999 |
| Sult2a3 | 1.00944E-11 | 5.020145168 |
| Sult2a5 | 3.90598E-07 | 3.288769507 |

| FRFvsAL Shared DEGs | | |
|---|---|---|
| Gene.ID | FDR | FC |
| Acta1 | 0 | 0.019740456 |
| Cib3 | 0.000651393 | 0.437517703 |
| Ckm | 0 | 0.036606473 |
| Gm3776 | 3.10862E-15 | 0.184471598 |
| Gsta1 | 3.14193E-14 | 0.196821754 |
| Gsta2 | 1.1957E-07 | 0.445361467 |
| Mb | 0 | 0.028679619 |
| Selenbp2 | 2.28817E-13 | 0.166708173 |
| Serpina1e | 0.00390842 | 0.366744636 |
| Sult2a3 | 2.00094E-07 | 0.279612243 |
| Sult2a5 | 7.16861E-05 | 0.367786047 |
| Tnnc2 | 0 | 0.037895842 |
| Tnni2 | 0 | 0.06157967 |
| Tnnt3 | 0 | 0.034324445 |
| Cyp17a1 | 2.62913E-09 | 3.514224773 |
| Cyp4a10 | 0 | 5.018063262 |
| Lpin1 | 1.75004E-12 | 5.484133448 |

| CRvsFRF Shared DEGs | | |
|---|---|---|
| Gene.ID | FDR | FC |
| Cyp17a1 | 0 | 0.111801808 |
| Cyp4a10 | 2.10942E-15 | 0.40053453 |
| Lpin1 | 0.002939248 | 0.386469443 |
| Selenbp2 | 2.24287E-12 | 0.154964373 |
| Serpina1e | 1.42941E-12 | 0.151404715 |
| Acta1 | 9.59233E-14 | 4.248973396 |
| Cib3 | 1.94289E-14 | 5.520379314 |
| Ckm | 5.8961E-07 | 2.127996914 |
| Gm3776 | 0 | 14.79062004 |
| Gsta1 | 0 | 14.06997848 |
| Gsta2 | 0 | 5.567951649 |
| Mb | 1.16993E-09 | 2.739754904 |
| Sult2a3 | 0 | 17.95395333 |
| Sult2a5 | 0 | 8.942072534 |
| Tnnc2 | 2.8824E-06 | 2.205567676 |
| Tnni2 | 0.00100766 | 2.036671262 |
| Tnnt3 | 2.00465E-09 | 2.792862069 |

Supplementary Table 1 for Fig 2G-J
Differentially expressed genes (FDR < 0.05)

| CRvsAL Shared DEGs | | |
|---|---|---|
| Gene.ID | FDR | FC |
| A1bg | 0.00151568 | 0.334701424 |
| Actn2 | 1.44857E-06 | 0.413776224 |
| Ak1 | 0.000133745 | 0.453781273 |
| Alas2 | 0.028928955 | 0.484106874 |
| Ankrd23 | 5.01124E-05 | 0.375329223 |
| Atp1a2 | 0.001413553 | 0.470780623 |
| Atp2a1 | 0 | 0.126532185 |
| Atp4a | 0.000234723 | 0.47110136 |
| C8a | 0.008265836 | 0.419464349 |
| C9 | 3.24333E-11 | 0.314884355 |
| Ces2c | 0.045906495 | 0.479617208 |
| Cfd | 1.31179E-08 | 0.246512136 |
| Ckmt2 | 0 | 0.148415668 |
| Cox6a2 | 9.39859E-12 | 0.211339311 |
| Cox8b | 7.79043E-12 | 0.208232941 |
| Cyp2u1 | 0.001500544 | 0.365595336 |
| Cyp3a44 | 3.08586E-07 | 0.252561806 |
| Cyp4a12a | 1.28222E-06 | 0.290368386 |
| Eef1a2 | 3.69732E-09 | 0.30907426 |
| Fabp3 | 0 | 0.125257662 |
| Gdf15 | 5.31583E-06 | 0.386298222 |
| Gstp2 | 3.07236E-05 | 0.307634376 |
| Hrc | 1.51468E-05 | 0.491529456 |
| Hsd3b5 | 2.14036E-10 | 0.207250608 |
| Irf5 | 8.9595E-13 | 0.475849408 |
| Keg1 | 3.36759E-08 | 0.390837264 |
| Ldb3 | 3.04045E-06 | 0.446534986 |
| Lifr | 0 | 0.316654398 |
| Moxd1 | 3.45313E-12 | 0.307849022 |
| Mup21 | 1.58846E-11 | 0.207886344 |
| Mybpc1 | 4.35187E-07 | 0.375071361 |
| Myh1 | 2.69784E-14 | 0.197271431 |
| Myh2 | 2.47347E-12 | 0.233645352 |
| Myl1 | 1.44329E-15 | 0.150091311 |
| Myl2 | 4.05598E-10 | 0.239387106 |
| Myl3 | 5.50553E-07 | 0.3693506 |
| Mylpf | 6.53472E-10 | 0.199748373 |
| Myot | 7.65473E-07 | 0.389895145 |
| Nat8f5 | 0.005379273 | 0.439069157 |

| FRFvsAL Shared DEGs | | |
|---|---|---|
| Gene.ID | FDR | FC |
| A1bg | 4.50598E-10 | 0.182987566 |
| Actn2 | 9.87999E-10 | 0.375907186 |
| Ak1 | 8.79315E-09 | 0.378163147 |
| Alas2 | 0.047257786 | 0.488713085 |
| Ankrd23 | 8.05008E-06 | 0.359279502 |
| Atp1a2 | 3.17344E-06 | 0.399203349 |
| Atp2a1 | 0 | 0.079635742 |
| Atp4a | 4.84526E-06 | 0.428967594 |
| C8a | 0.008417671 | 0.453904208 |
| C9 | 1.21423E-05 | 0.449987446 |
| Ces2c | 0.000409261 | 0.356608849 |
| Cfd | 1.0498E-08 | 0.24815986 |
| Ckmt2 | 0 | 0.117717391 |
| Cox6a2 | 0 | 0.152037992 |
| Cox8b | 2.22045E-16 | 0.16946122 |
| Cyp2u1 | 8.1271E-06 | 0.385084822 |
| Cyp3a44 | 9.37525E-05 | 0.329954762 |
| Cyp4a12a | 0.046540031 | 0.473458262 |
| Eef1a2 | 7.58282E-14 | 0.266530824 |
| Fabp3 | 0 | 0.083207161 |
| Gdf15 | 1.07779E-08 | 0.313364473 |
| Gstp2 | 5.01144E-12 | 0.17193627 |
| Hrc | 2.90949E-07 | 0.47080292 |
| Hsd3b5 | 1.36202E-11 | 0.191750317 |
| Irf5 | 7.10543E-15 | 0.451833441 |
| Keg1 | 4.59196E-06 | 0.470238834 |
| Ldb3 | 1.69323E-07 | 0.431287708 |
| Lifr | 0 | 0.375531416 |
| Moxd1 | 3.83775E-08 | 0.355363933 |
| Mup21 | 6.32587E-06 | 0.326399492 |
| Mybpc1 | 1.01876E-11 | 0.32326176 |
| Myh1 | 0 | 0.165279406 |
| Myh2 | 0 | 0.197448957 |
| Myl1 | 0 | 0.099424147 |
| Myl2 | 0 | 0.161931248 |
| Myl3 | 5.46774E-12 | 0.308769044 |
| Mylpf | 1.0697E-12 | 0.165138846 |
| Myot | 5.87088E-11 | 0.342273096 |
| Nat8f5 | 0.006429168 | 0.453276408 |

Supplementary Table 1 for Fig 2G-J

Differentially expressed genes (FDR < 0.05)

| CRvsAL Shared DEGs | | |
|---|---|---|
| Gene.ID | FDR | FC |
| Neb | 0.006628626 | 0.49724091 |
| Nr4a1 | 2.21045E-13 | 0.215201536 |
| Nudt7 | 3.00105E-06 | 0.297746118 |
| Pgam2 | 1.07859E-07 | 0.328367092 |
| Pvalb | 2.95208E-05 | 0.40107741 |
| Pygm | 0.002761798 | 0.445138017 |
| Slc10a2 | 1.84297E-14 | 0.289231484 |
| Slc22a28 | 1.27209E-12 | 0.240617356 |
| Slc25a4 | 0.032547473 | 0.459611035 |
| Tcap | 0 | 0.118995454 |
| Tff3 | 0 | 0.090081701 |
| Ugt2b37 | 2.17015E-07 | 0.335953837 |
| Ugt2b38 | 1.79011E-07 | 0.320610159 |
| Acot2 | 2.71862E-07 | 2.166020935 |
| Cyp2c39 | 1.12718E-05 | 2.268900606 |
| Gm45531 | 2.62771E-10 | 3.222344287 |
| Plin4 | 0.001144366 | 2.18480194 |
| Rgs16 | 0.001100314 | 2.365365319 |
| Sult1e1 | 9.91282E-11 | 4.114395433 |

| FRFvsAL Shared DEGs | | |
|---|---|---|
| Gene.ID | FDR | FC |
| Neb | 0.003319387 | 0.495326691 |
| Nr4a1 | 0.00296579 | 0.39471527 |
| Nudt7 | 1.84618E-05 | 0.31560912 |
| Pgam2 | 1.98075E-12 | 0.271771497 |
| Pvalb | 1.3043E-11 | 0.304421845 |
| Pygm | 3.1858E-06 | 0.352158518 |
| Slc10a2 | 1.59966E-08 | 0.424736989 |
| Slc22a28 | 5.90349E-09 | 0.30701492 |
| Slc25a4 | 8.49367E-06 | 0.312075478 |
| Tcap | 0 | 0.081621174 |
| Tff3 | 0 | 0.096537795 |
| Ugt2b37 | 0.001454006 | 0.488407708 |
| Ugt2b38 | 0.000277867 | 0.40994226 |
| Acot2 | 9.11968E-05 | 2.158893008 |
| Cyp2c39 | 2.06895E-11 | 3.245274129 |
| Gm45531 | 4.9657E-07 | 2.545390123 |
| Plin4 | 4.17066E-07 | 2.11842087 |
| Rgs16 | 7.21309E-11 | 4.561105097 |
| Sult1e1 | 5.09592E-14 | 4.182974608 |

Supplementary Table 1 for Fig 2G-J

Differentially expressed genes (FDR < 0.05)

| CRvsAL Shared DEGs | | |
|---|---|---|
| Gene.ID | FDR | FC |
| Aatk | 2.27264E-05 | 0.444051379 |
| Derl3 | 2.9529E-06 | 0.344047467 |
| Dio1 | 2.12053E-14 | 0.316450587 |
| Dusp1 | 0 | 0.242545446 |
| Fos | 3.14809E-11 | 0.301222056 |
| Gadd45g | 0.006404114 | 0.432425116 |
| Lad1 | 5.17372E-06 | 0.477169071 |
| Mup1 | 0 | 0.067627015 |
| Mup10 | 6.95244E-08 | 0.44216582 |
| Mup11 | 0 | 0.020807491 |
| Mup12 | 0 | 0.080700481 |
| Mup13 | 0 | 0.16280147 |
| Mup14 | 0 | 0.036437768 |
| Mup15 | 0 | 0.001251336 |
| Mup16 | 3.34116E-10 | 0.167154252 |
| Mup17 | 0 | 0.001208035 |
| Mup18 | 2.22045E-16 | 0.20894047 |
| Mup19 | 0 | 0.103706139 |
| Mup2 | 0 | 0.038488637 |
| Mup20 | 5.32863E-12 | 0.17773778 |
| Mup4 | 1.58697E-06 | 0.416818899 |
| Mup5 | 3.33765E-05 | 0.465988467 |
| Mup6 | 1.30851E-12 | 0.248572231 |
| Mup7 | 0 | 0.041307846 |
| Mup8 | 0 | 0.143658353 |
| Mup9 | 0 | 0.045248675 |
| Obp2a | 1.23726E-09 | 0.293190065 |
| Rarres1 | 2.71242E-06 | 0.265863697 |
| Rgs1 | 8.90322E-11 | 0.359815082 |
| RP24-346O11.2 | 0 | 0.209102883 |
| Serpina12 | 1.81847E-05 | 0.365804044 |
| Slc22a7 | 1.46495E-06 | 0.330381882 |
| Sult3a1 | 0.049317209 | 0.472270472 |
| Sult5a1 | 0 | 0.272105353 |
| Ugt3a1 | 0 | 0.453661355 |
| Akr1b7 | 0.002260086 | 2.407943032 |
| Car2 | 9.75297E-07 | 2.30142591 |
| Cbr3 | 0.000488135 | 2.45754448 |

| CRvsFRF Shared DEGs | | |
|---|---|---|
| Gene.ID | FDR | FC |
| Aatk | 1.04679E-05 | 0.425961795 |
| Derl3 | 2.5574E-12 | 0.208548355 |
| Dio1 | 0 | 0.254707234 |
| Dusp1 | 0.000210589 | 0.431392179 |
| Fos | 5.97125E-08 | 0.309374835 |
| Gadd45g | 0.001954184 | 0.347975065 |
| Lad1 | 9.99348E-08 | 0.375905411 |
| Mup1 | 0 | 0.101398968 |
| Mup10 | 8.05725E-08 | 0.485436284 |
| Mup11 | 0 | 0.026151123 |
| Mup12 | 3.28704E-12 | 0.164499557 |
| Mup13 | 1.0536E-13 | 0.20715241 |
| Mup14 | 0 | 0.061741411 |
| Mup15 | 0 | 0.001548788 |
| Mup16 | 0 | 0.182797973 |
| Mup17 | 0 | 0.002356411 |
| Mup18 | 0 | 0.206637236 |
| Mup19 | 0 | 0.081785208 |
| Mup2 | 0 | 0.058652744 |
| Mup20 | 3.86318E-08 | 0.207771643 |
| Mup4 | 2.8789E-07 | 0.444006451 |
| Mup5 | 3.02045E-08 | 0.483779257 |
| Mup6 | 5.13223E-12 | 0.316908514 |
| Mup7 | 0 | 0.065348485 |
| Mup8 | 1.1207E-09 | 0.274537863 |
| Mup9 | 0 | 0.060420802 |
| Obp2a | 3.69481E-06 | 0.404073571 |
| Rarres1 | 6.32738E-12 | 0.27125128 |
| Rgs1 | 3.91217E-08 | 0.373095298 |
| RP24-346O11.2 | 0 | 0.178303289 |
| Serpina12 | 0.005639979 | 0.47986207 |
| Slc22a7 | 0.010087846 | 0.487087857 |
| Sult3a1 | 0.000151349 | 0.291271926 |
| Sult5a1 | 4.67422E-09 | 0.428692469 |
| Ugt3a1 | 0 | 0.476651183 |
| Akr1b7 | 0.00027254 | 2.063976972 |
| Car2 | 2.53015E-07 | 2.344295988 |
| Cbr3 | 2.22448E-06 | 2.96245424 |

Supplementary Table 1 for Fig 2G-J
Differentially expressed genes (FDR < 0.05)

| CRvsAL Shared DEGs | | |
|---|---|---|
| Gene.ID | FDR | FC |
| Cd24a | 0.01075699 | 2.063029842 |
| Cyp2a5 | 2.26196E-06 | 2.504369767 |
| Cyp2g1 | 3.19286E-06 | 2.426559881 |
| Dmbt1 | 0 | 7.944036073 |
| Dsg1c | 0.002140063 | 2.031480893 |
| Fmo3 | 0 | 7.676858134 |
| Gm10642 | 0.000925496 | 2.009230208 |
| Gpx6 | 6.76188E-05 | 2.453376704 |
| Krt19 | 0.002251705 | 2.390049448 |
| Muc1 | 0.000281649 | 2.335947951 |
| Orm2 | 9.31621E-12 | 7.166736752 |
| Rdh9 | 1.31117E-12 | 2.802142771 |
| Sult2a1 | 0.029225605 | 2.329360699 |
| Tgtp1 | 0.021034826 | 2.270367552 |
| Ugt1a5 | 0.000338896 | 2.051239992 |

| CRvsFRF Shared DEGs | | |
|---|---|---|
| Gene.ID | FDR | FC |
| Cd24a | 0.001181821 | 2.259501222 |
| Cyp2a5 | 1.30601E-07 | 2.505562187 |
| Cyp2g1 | 1.41631E-11 | 3.517730926 |
| Dmbt1 | 0 | 9.540461307 |
| Dsg1c | 5.89167E-07 | 2.870496781 |
| Fmo3 | 1.09699E-10 | 4.635336743 |
| Gm10642 | 7.77657E-05 | 2.15658371 |
| Gpx6 | 1.21839E-09 | 3.686938591 |
| Krt19 | 2.11243E-05 | 2.896816189 |
| Muc1 | 9.74108E-08 | 2.933918287 |
| Orm2 | 1.16696E-12 | 7.794459649 |
| Rdh9 | 2.46367E-07 | 2.084025901 |
| Sult2a1 | 0.048327358 | 2.222802589 |
| Tgtp1 | 9.57677E-05 | 2.950673321 |
| Ugt1a5 | 1.07484E-08 | 2.689431926 |

Supplementary Table 1 for Fig 2G-J
Differentially expressed genes (FDR < 0.05)

| FRFvsAL Shared DEGs | | |
|---|---|---|
| Gene.ID | FDR | FC |
| Abca8a | 0 | 0.468629517 |
| Acta2 | 0.002010457 | 0.418713806 |
| Angptl8 | 1.17109E-06 | 0.288927335 |
| BC048546 | 1.85903E-10 | 0.471820134 |
| Car3 | 0 | 0.209210994 |
| Clec2h | 0.000176093 | 0.455780513 |
| Col3a1 | 1.50842E-08 | 0.414159548 |
| Cox7a1 | 0.00643221 | 0.441985882 |
| Cyp2c44 | 0 | 0.407018406 |
| Cyp2c55 | 6.39488E-14 | 0.215831734 |
| Cyp3a16 | 1.61464E-06 | 0.265921401 |
| Cyp3a41b | 5.93808E-09 | 0.216458601 |
| Fabp5 | 9.17599E-13 | 0.29097849 |
| Gas6 | 7.95549E-10 | 0.446579563 |
| Gstm3 | 3.51858E-05 | 0.35938635 |
| Hamp2 | 4.86827E-09 | 0.232488284 |
| Hao2 | 4.13247E-07 | 0.311450958 |
| Inmt | 5.21805E-15 | 0.451239977 |
| Klk1b4 | 3.40906E-08 | 0.424343438 |
| Lgals1 | 0 | 0.332571798 |
| Lpl | 2.65368E-05 | 0.358674377 |
| Nrep | 7.11347E-07 | 0.317950022 |
| Prom1 | 9.31447E-05 | 0.413165519 |
| Raet1d | 8.24649E-07 | 0.371691674 |
| Rnf186 | 1.62028E-11 | 0.364013254 |
| Scd1 | 0 | 0.100451474 |
| Serpina6 | 6.66134E-16 | 0.316614039 |
| Smim22 | 9.34653E-10 | 0.46450789 |
| Srebf1 | 0 | 0.253401818 |
| Sucnr1 | 2.76279E-12 | 0.353841111 |
| Sult2a7 | 0.021462335 | 0.458980308 |
| 1810055G02Rik | 2.68796E-12 | 2.51872783 |
| 2010003K11Rik | 1.44329E-15 | 2.755562446 |
| 2210010C04Rik | 0.001263165 | 2.092454774 |
| Acnat2 | 0.000132319 | 2.266672808 |
| Amy2a5 | 0 | 11.04069827 |
| Angptl4 | 7.58166E-10 | 2.559645122 |
| Apoa4 | 0 | 7.051300283 |
| Bhmt | 8.67084E-14 | 2.680011296 |

| CRvsFRF Shared DEGs | | |
|---|---|---|
| Gene.ID | FDR | FC |
| 1810055G02Rik | 3.47096E-10 | 0.403112964 |
| 2010003K11Rik | 3.42264E-09 | 0.431561438 |
| 2210010C04Rik | 6.65558E-06 | 0.404793774 |
| Acnat2 | 8.03024E-13 | 0.282418575 |
| Amy2a5 | 0 | 0.09363096 |
| Angptl4 | 0.000682775 | 0.483364809 |
| Apoa4 | 0 | 0.217798552 |
| Bhmt | 5.42899E-14 | 0.362226954 |
| Cdkn1a | 0 | 0.134025736 |
| Cebpd | 1.11022E-16 | 0.382195075 |
| Cel | 6.17545E-06 | 0.429464147 |
| Cela2a | 2.69129E-09 | 0.207008789 |
| Cela3b | 9.10383E-15 | 0.166129496 |
| Clps | 4.37236E-11 | 0.218219312 |
| Cpa1 | 1.03504E-10 | 0.25139185 |
| Cpb1 | 1.29633E-07 | 0.385345419 |
| Ctgf | 5.151E-05 | 0.484879129 |
| Ctrb1 | 8.11552E-11 | 0.16937972 |
| Ctrl | 6.36052E-08 | 0.296608082 |
| Cyp39a1 | 3.67537E-06 | 0.461043526 |
| Cyp4a14 | 0 | 0.356797822 |
| Cyp4a31 | 0 | 0.319349449 |
| Cyp8b1 | 8.26152E-08 | 0.376934865 |
| Ddit4 | 2.4899E-12 | 0.22554208 |
| E030018B13Rik | 1.11022E-16 | 0.205791168 |
| Eif4ebp3 | 2.73537E-12 | 0.299364025 |
| Fam35a | 0 | 0.405957892 |
| Fgf21 | 1.54746E-07 | 0.249883683 |
| Fkbp5 | 0.000428207 | 0.475199045 |
| Fmo2 | 0.001375068 | 0.477573911 |
| Gadd45b | 0.000178138 | 0.475527473 |
| Gfra1 | 0 | 0.457907546 |
| Insig2 | 9.355E-08 | 0.395365913 |
| Lcn2 | 0.00229257 | 0.469414371 |
| Lepr | 0.011229922 | 0.446962849 |
| Mat1a | 0 | 0.499147401 |
| Mt1 | 0.006821232 | 0.415612509 |
| Mt2 | 1.93423E-11 | 0.18378338 |
| Nlrp12 | 1.12698E-09 | 0.410265801 |

Supplementary Table 1 for Fig 2G-J
Differentially expressed genes (FDR < 0.05)

| FRFvsAL Shared DEGs | | |
|---|---|---|
| Gene.ID | FDR | FC |
| Cdkn1a | 3.76366E-14 | 5.556601228 |
| Cebpd | 1.3843E-11 | 2.205107854 |
| Cel | 0.000886592 | 2.028041792 |
| Cela2a | 0.003340332 | 2.590505854 |
| Cela3b | 1.06195E-08 | 4.127269354 |
| Clps | 2.41743E-08 | 3.820432548 |
| Cpa1 | 4.2384E-07 | 3.198574253 |
| Cpb1 | 0.000248465 | 2.161737368 |
| Ctgf | 1.72587E-11 | 2.629878267 |
| Ctrb1 | 5.93282E-09 | 5.008811989 |
| Ctrl | 0.000123808 | 2.607417379 |
| Cyp39a1 | 2.55351E-15 | 3.886519397 |
| Cyp4a14 | 0 | 5.255290467 |
| Cyp4a31 | 0 | 3.604994241 |
| Cyp8b1 | 2.22053E-07 | 2.522557108 |
| Ddit4 | 1.8928E-10 | 3.876601413 |
| E030018B13Rik | 1.59952E-11 | 3.701540076 |
| Eif4ebp3 | 0 | 6.898793291 |
| Fam35a | 0 | 2.212663729 |
| Fgf21 | 6.49637E-07 | 2.969894284 |
| Fkbp5 | 1.82188E-13 | 3.830466804 |
| Fmo2 | 7.08711E-05 | 2.554571395 |
| Gadd45b | 4.37836E-07 | 2.555489682 |
| Gfra1 | 0 | 2.283965417 |
| Insig2 | 3.94851E-06 | 2.262263082 |
| Lcn2 | 1.01999E-06 | 2.615707137 |
| Lepr | 0.003000924 | 2.451542049 |
| Mat1a | 0 | 2.2092763 |
| Mt1 | 5.14838E-08 | 3.70755198 |
| Mt2 | 7.47613E-12 | 5.494909803 |
| Nlrp12 | 8.32667E-15 | 3.22372386 |
| Pnlip | 1.93252E-10 | 4.696121248 |
| Prss2 | 1.56763E-09 | 4.611704186 |
| Rab30 | 8.98391E-08 | 2.881686232 |
| Rnase1 | 0.000123532 | 2.385565461 |
| Serpina3n | 0 | 2.072426734 |
| Slc17a8 | 6.10623E-15 | 3.372837487 |
| St3gal5 | 4.90274E-13 | 3.595602737 |
| Sult1d1 | 1.68128E-11 | 2.517566445 |

| CRvsFRF Shared DEGs | | |
|---|---|---|
| Gene.ID | FDR | FC |
| Pnlip | 0 | 0.148967615 |
| Prss2 | 9.99201E-16 | 0.14619585 |
| Rab30 | 0.000641564 | 0.457797629 |
| Rnase1 | 1.3431E-08 | 0.328294443 |
| Serpina3n | 0 | 0.461393131 |
| Slc17a8 | 2.27272E-08 | 0.423713742 |
| St3gal5 | 7.2734E-06 | 0.428695324 |
| Sult1d1 | 1.08685E-07 | 0.473472802 |
| Sycn | 2.42703E-11 | 0.230083731 |
| Try4 | 0 | 0.128467217 |
| Try5 | 2.42029E-14 | 0.177609675 |
| Zg16 | 9.86958E-07 | 0.346058161 |
| Abca8a | 2.95352E-09 | 2.034285601 |
| Acta2 | 3.88981E-06 | 2.860240668 |
| Angptl8 | 9.08717E-06 | 3.32410588 |
| BC048546 | 1.11022E-16 | 2.118506152 |
| Car3 | 2.89468E-07 | 2.956467775 |
| Clec2h | 2.59444E-09 | 3.244075016 |
| Col3a1 | 0.010998986 | 2.202113276 |
| Cox7a1 | 9.06298E-09 | 2.351357867 |
| Cyp2c44 | 0 | 2.000226121 |
| Cyp2c55 | 1.9984E-15 | 4.9937726 |
| Cyp3a16 | 0.000616037 | 2.775996854 |
| Cyp3a41b | 3.49526E-06 | 3.451939122 |
| Fabp5 | 0.000258252 | 2.465931601 |
| Gas6 | 3.64101E-07 | 2.181178738 |
| Gstm3 | 4.56281E-10 | 6.36683767 |
| Hamp2 | 3.02758E-13 | 6.379174038 |
| Hao2 | 2.27004E-08 | 3.386672794 |
| Inmt | 0 | 3.050326318 |
| Klk1b4 | 1.16912E-06 | 2.118601821 |
| Lgals1 | 0 | 2.607204491 |
| Lpl | 1.43574E-07 | 2.928396233 |
| Nrep | 0.000108935 | 2.904732756 |
| Prom1 | 0.000321195 | 2.249020757 |
| Raet1d | 2.49134E-13 | 4.089197752 |
| Rnf186 | 7.39853E-13 | 2.166867646 |
| Scd1 | 1.11022E-16 | 6.32515511 |
| Serpina6 | 3.05311E-14 | 2.919549482 |

Supplementary Table 1 for Fig 2G-J

Differentially expressed genes (FDR < 0.05)

| FRFvsAL Shared DEGs | | |
|---|---|---|
| Gene.ID | FDR | FC |
| Sycn | 4.27854E-06 | 3.072939989 |
| Try4 | 8.74721E-10 | 4.669376786 |
| Try5 | 1.99304E-09 | 4.223964283 |
| Zg16 | 4.18197E-06 | 2.755868032 |

| CRvsFRF Shared DEGs | | |
|---|---|---|
| Gene.ID | FDR | FC |
| Smim22 | 1.12577E-12 | 2.348859517 |
| Srebf1 | 1.68321E-10 | 2.657729656 |
| Sucnr1 | 5.51942E-10 | 2.549851095 |
| Sult2a7 | 1.14598E-06 | 2.997576327 |
