## Supplementary Table 2 for "Anticipatory metabolic reprogramming distinguishes caloric restriction from fasting-refeeding cycles"

Supplementary Table 2 for Fig 3D-G

| CR v AL up | FRF v AL up |
|---|---|
| 1810055G02Rik | 1810055G02Rik |
| 2010003K11Rik | 2010003K11Rik |
| Aass | Aass |
| Acot1 | Acot1 |
| Acot2 | Acot2 |
| Acot4 | Acot4 |
| Aifm2 | Aifm2 |
| Amy2a5 | Amy2a5 |
| Apoa4 | Apoa4 |
| Aprt | Aprt |
| Atl2 | Atl2 |
| Bnip3 | Bnip3 |
| Cdkn1a | Cdkn1a |
| Ces1g | Ces1g |
| Chd9 | Chd9 |
| Cpt1a | Cpt1a |
| Crat | Crat |
| Creld2 | Creld2 |
| Ctgf | Ctgf |
| Cyp39a1 | Cyp39a1 |
| Cyp3a59 | Cyp3a59 |
| Cyp4a10 | Cyp4a10 |
| Cyp4a14 | Cyp4a14 |
| Cyp51 | Cyp51 |
| Dnajb9 | Dnajb9 |
| Dynll1 | Dynll1 |
| Ethe1 | Ethe1 |
| Fam134b | Fam134b |
| Fbxo21 | Fbxo21 |
| Fbxo31 | Fbxo31 |
| Fkbp5 | Fkbp5 |
| Gabarapl1 | Gabarapl1 |
| Gadd45b | Gadd45b |
| Gde1 | Gde1 |
| Glrx | Glrx |
| Gm45531 | Gm45531 |
| Grpel2 | Grpel2 |
| Hbb-bs | Hbb-bs |
| Hist2h3c2 | Hist2h3c2 |
| Il6ra | Il6ra |
| Lepr | Lepr |
| Lsm7 | Lsm7 |
| Lss | Lss |
| Map1lc3b | Map1lc3b |
| Mat1a | Mat1a |

| CR v AL down | FRF v AL down |
|---|---|
| A1bg | A1bg |
| Abcg8 | Abcg8 |
| Acly | Acly |
| Adh4 | Adh4 |
| Ankrd23 | Ankrd23 |
| Apol9a | Apol9a |
| Atp2a1 | Atp2a1 |
| Bhlhe40 | Bhlhe40 |
| Btg2 | Btg2 |
| Car3 | Car3 |
| Cebpb | Cebpb |
| Cfd | Cfd |
| Cish | Cish |
| Ckm | Ckm |
| Ckmt2 | Ckmt2 |
| Cox6a2 | Cox6a2 |
| Cox8b | Cox8b |
| Csrnp1 | Csrnp1 |
| Cxcl1 | Cxcl1 |
| Cyp2c70 | Cyp2c70 |
| Cyp3a44 | Cyp3a44 |
| Cyp4a12a | Cyp4a12a |
| Cyp4a12b | Cyp4a12b |
| Cyp4f14 | Cyp4f14 |
| Cyr61 | Cyr61 |
| Dusp1 | Dusp1 |
| Dusp6 | Dusp6 |
| Eef1a2 | Eef1a2 |
| Elovl3 | Elovl3 |
| Elovl6 | Elovl6 |
| Fabp3 | Fabp3 |
| Fabp5 | Fabp5 |
| Fads2 | Fads2 |
| Fam25c | Fam25c |
| Fam47e | Fam47e |
| Fasn | Fasn |
| G0s2 | G0s2 |
| G6pc | G6pc |
| Gadd45a | Gadd45a |
| Gale | Gale |
| Gdf15 | Gdf15 |
| Gstp2 | Gstp2 |
| H2afv | H2afv |
| Hdhd3 | Hdhd3 |
| Hes6 | Hes6 |

| CR v AL up | FRF v AL down |
|---|---|
| Cyp2a5 | Cyp2a5 |
| Fmo3 | Fmo3 |
| Gm3776 | Gm3776 |
| Gsta1 | Gsta1 |
| Gsta2 | Gsta2 |
| Gstm1 | Gstm1 |
| Gstm3 | Gstm3 |
| Gys2 | Gys2 |
| Raet1d | Raet1d |
| Srsf3 | Srsf3 |
| Sult2a3 | Sult2a3 |
| Sult2a5 | Sult2a5 |
| Rdh9 | Cyp2c44 |
| Orm2 | Srebf1 |
| Acot2 | Scd1 |
| Gm45531 | Lifr |
| Hbb-bs | Serpina6 |
| Rbm3 | Car3 |
| Ctgf | Gale |
| Fbxo31 | Mid1ip1 |
| Fkbp5 | Nrep |
| Gstt2 | Hsd3b5 |
| Hba-a2 | Nat8f1 |
| Hbb-bt | Angptl8 |
| Lgals4 | Ces2a |
| Pnpla2 | Cfd |
| Rab30 | Inmt |
| Txnip | Lgals1 |
| Bnip3 | Aqp8 |
| Ces1g | Gstp2 |
| Fbxo21 | Acta1 |
| Lpin1 | Atp2a1 |
| Pcsk4 | Ckm |
| Por | Ckmt2 |
| Slc16a7 | Cox6a2 |
| Dmbt1 | Cox8b |
| Plin5 | Fabp3 |
| Slc16a5 | Gdf15 |
| Cyp3a59 | Insc |
| Aprt | Mb |
| Ethe1 | Myh1 |
| Tceal8 | Myl1 |
| Ak4 | Tcap |
| Aldoc | Tmem150a |
| Btbd19 | Tnnc2 |

| CR v AL down | FRF v AL up |
|---|---|
| Arl4a | Arl4a |
| Car14 | Car14 |
| Csrp3 | Csrp3 |
| D230025D16Rik | D230025D16Rik |
| Gadd45g | Gadd45g |
| Hfe2 | Hfe2 |
| Hmgcr | Hmgcr |
| Hopx | Hopx |
| Igfbp1 | Igfbp1 |
| Ip6k2 | Ip6k2 |
| Leap2 | Leap2 |
| Mvd | Mvd |
| Nsdhl | Nsdhl |
| Pdia4 | Pdia4 |
| Rapgef4 | Rapgef4 |
| Rnf125 | Rnf125 |
| Slc45a3 | Slc45a3 |
| Sult3a2 | Sult3a2 |
| Syvn1 | Syvn1 |
| Trp53inp1 | Trp53inp1 |
| Tsku | Tsku |
| Tubb4b | Tubb4b |
| Ubc | Ubc |
| Dusp1 | Eif4ebp3 |
| Lifr | Apoa4 |
| Mup15 | Cyp4a31 |
| Mup17 | Cdip1 |
| Selenbp2 | Cyp4a10 |
| Mup11 | Cyp4a14 |
| Mup2 | Rgs16 |
| Mup14 | Glrx |
| Mup9 | Slc37a4 |
| Sult5a1 | Nlrp12 |
| Cyp2c70 | Pnpla2 |
| Nr4a1 | Serpina3n |
| Sik1 | Bnip3 |
| Ugt3a1 | Fbxo31 |
| Nrep | St3gal5 |
| Slc22a30 | Cyp39a1 |
| Tff3 | Angptl4 |
| Thrsp | 1810055G02Rik |
| G6pc | Fkbp5 |
| Hsd3b5 | Gfra1 |
| Mup1 | Il6ra |
| Mup12 | Plin5 |

Supplementary Table 2 for Fig 3D-G

| CR v AL up | FRF v AL up |
|---|---|
| Mfsd2a | Mfsd2a |
| Mt1 | Mt1 |
| Nceh1 | Nceh1 |
| Pex11a | Pex11a |
| Phospho1 | Phospho1 |
| Plin5 | Plin5 |
| Pnpla2 | Pnpla2 |
| Por | Por |
| Rab30 | Rab30 |
| Rbm3 | Rbm3 |
| Rdh9 | Rdh9 |
| Reg1 | Reg1 |
| Retsat | Retsat |
| Rgs16 | Rgs16 |
| Serinc3 | Serinc3 |
| Serpina3m | Serpina3m |
| Slc16a7 | Slc16a7 |
| Slc20a1 | Slc20a1 |
| Slc22a5 | Slc22a5 |
| Slc8b1 | Slc8b1 |
| Slco1a4 | Slco1a4 |
| Sorbs3 | Sorbs3 |
| St3gal5 | St3gal5 |
| Sult1e1 | Sult1e1 |
| Tdo2 | Tdo2 |
| Tmem120a | Tmem120a |
| Tmem254c | Tmem254c |
| Tmem56 | Tmem56 |
| Try4 | Try4 |
| Tuba8 | Tuba8 |
| Txnip | Txnip |
| Fmo3 | Eif4ebp3 |
| Cyp2a5 | Cyp4a31 |
| Orm2 | Cdip1 |
| Sult2a3 | Slc37a4 |
| Gstt2 | Nlrp12 |
| Hba-a2 | Serpina3n |
| Hbb-bt | Angptl4 |
| Lgals4 | Gfra1 |
| Gsta2 | Ip6k2 |
| Lpin1 | Cyp8b1 |
| Pcsk4 | Mt2 |
| Dmbt1 | Cyp2c39 |
| Gm3776 | Tango2 |
| Gsta1 | Atf5 |

| CR v AL down | FRF v AL down |
|---|---|
| Hsd3b2 | Hsd3b2 |
| Hsd3b5 | Hsd3b5 |
| Inhbc | Inhbc |
| Insc | Insc |
| Keg1 | Keg1 |
| Kyat1 | Kyat1 |
| Lasp1 | Lasp1 |
| Lifr | Lifr |
| Ly6d | Ly6d |
| Mafb | Mafb |
| Mapk15 | Mapk15 |
| Mug2 | Mug2 |
| Mup1 | Mup1 |
| Mup12 | Mup12 |
| Mup20 | Mup20 |
| Mup21 | Mup21 |
| Mup7 | Mup7 |
| Myh1 | Myh1 |
| Myh2 | Myh2 |
| Myl1 | Myl1 |
| Myl2 | Myl2 |
| Mylpf | Mylpf |
| Nat8f1 | Nat8f1 |
| Nat8f5 | Nat8f5 |
| Nfil3 | Nfil3 |
| Noct | Noct |
| Nr0b2 | Nr0b2 |
| Nr4a1 | Nr4a1 |
| Nrep | Nrep |
| Oasl1 | Oasl1 |
| Onecut1 | Onecut1 |
| Pcsk9 | Pcsk9 |
| Pdia6 | Pdia6 |
| Pgam2 | Pgam2 |
| Phlda1 | Phlda1 |
| Plk3 | Plk3 |
| Pnrc1 | Pnrc1 |
| Ppp1r3g | Ppp1r3g |
| Saa1 | Saa1 |
| Saa2 | Saa2 |
| Saa3 | Saa3 |
| Selenbp2 | Selenbp2 |
| Serpina1e | Serpina1e |
| Serpina7 | Serpina7 |
| Sik1 | Sik1 |

| CR v AL up | FRF v AL down |
|---|---|
| Clec2h | Tnni2 |
| Cnppd1 | Tnnt3 |
| Eif4b | Acly |
| Gabarapl1 | Fabp5 |
| Il6ra | Selenbp2 |
| Lurap1l | Kyat1 |
| Mfsd2a | Tmsb10 |
| Pla2g12a | Acaa1b |
| Ppp1r10 | Cyp26a1 |
| Rpl22l1 | Dusp1 |
| Slc8b1 | Efna1 |
| Slco1a4 | Elovl6 |
| Soat2 | Myh2 |
| Sorbs3 | Mylpf |
| Sult1e1 | Thrsp |
| Zpr1 | Cish |
| Car2 | Eef1a2 |
| Cbr1 | Id3 |
| Chd9 | Sucnr1 |
| Cib3 | Tgm2 |
| Dnaja1 | A1bg |
| Dynll1 | Lasp1 |
| Gm20594 | Mapk15 |
| Odc1 | Cd74 |
| Reg1 | Fam25c |
| Tuba8 | Gstp3 |
| Orm3 | H2-Aa |
| Pmvk | H2-Eb1 |
| Cirbp | Ifi47 |
| Gstt3 | Adh4 |
| 2010003K11Rik | Cyp2c55 |
| Acot1 | Vegfb |
| Acot4 | BC048546 |
| Agpat9 | Hao2 |
| Crat | Rnf186 |
| Cyp39a1 | Ankrd23 |
| Cyp4a14 | G0s2 |
| Ehhadh | Gadd45a |
| Gadd45b | Hes6 |
| Hba-a1 | Myl2 |
| Pdk4 | Slc9a3r1 |
| Pex11a | A230050P20Rik |
| Plin2 | Aacs |
| Retsat | Fasn |
| Rgs16 | Gas6 |

| CR v AL down | FRF v AL up |
|---|---|
| Mup18 | Retsat |
| Mup19 | Cdkn1a |
| Mup7 | Cyp8b1 |
| Zfp36 | Fam134b |
| Bhlhe40 | Gm45531 |
| Jun | Rab30 |
| Slc25a25 | Rbm3 |
| Noct | Atl2 |
| A1bg | Pex11a |
| Ankrd23 | Por |
| Apol9a | Chd9 |
| Atp2a1 | Mt2 |
| Calr | 2010003K11Rik |
| Ckm | Cyp2c39 |
| Ckmt2 | Serpina3m |
| Cox6a2 | Tango2 |
| Fabp3 | Atf5 |
| Gale | Bhmt |
| Hdhd3 | Cpt1a |
| Myh1 | Got1 |
| Myl1 | Slc10a1 |
| Mylpf | Il1r1 |
| Pdia6 | Mfsd2a |
| Sdf2l1 | Rdh9 |
| Srebf1 | Slc17a8 |
| Tcap | Slc22a5 |
| Tnnc2 | Ctgf |
| Enho | Herpud1 |
| Mup13 | Sult1d1 |
| Mup8 | Acnat2 |
| Tob1 | Cyp17a1 |
| Dio1 | Ahcy |
| Pim3 | Ddit4 |
| Btg2 | Mat1a |
| Actb | Tmem120a |
| Cox8b | S100a10 |
| Derl3 | Abca6 |
| Hes6 | Cdo1 |
| Hsd3b7 | Gabarapl1 |
| Lasp1 | Gm4952 |
| Paqr9 | Inca1 |
| Saa1 | Lbp |
| Slc10a2 | Npc1 |
| Slc35b1 | Tymp |
| Tmem254a | Rps12 |

Supplementary Table 2 for Fig 3D-G

| CR v AL up | FRF v AL up |
|---|---|
| Slc16a5 | Bhmt |
| Tceal8 | Csrp3 |
| Gstm3 | Got1 |
| Ak4 | Slc10a1 |
| Aldoc | Il1r1 |
| Btbd19 | Slc17a8 |
| Clec2h | Herpud1 |
| Cnppd1 | Sult1d1 |
| Eif4b | Acnat2 |
| Lurap1l | Cyp17a1 |
| Pla2g12a | Ahcy |
| Ppp1r10 | Ddit4 |
| Rpl22l1 | S100a10 |
| Soat2 | Abca6 |
| Zpr1 | Cdo1 |
| Car2 | Gm4952 |
| Cbr1 | Inca1 |
| Cib3 | Lbp |
| Dnaja1 | Npc1 |
| Gm20594 | Tymp |
| Odc1 | Trp53inp1 |
| Srsf3 | Rps12 |
| Orm3 | Fam35a |
| Pmvk | Hsd17b10 |
| Cirbp | Hsp90ab1 |
| Gstt3 | Tsku |
| Agpat9 | 1600002H07Rik |
| Ehhadh | As3mt |
| Hba-a1 | Fam234b |
| Pdk4 | Fgf21 |
| Plin2 | Nr1i3 |
| Alas1 | Rapgef4 |
| Bag3 | Sult1a1 |
| Cbr3 | Arl4a |
| Gys2 | Ccng2 |
| Hsd17b13 | Fmo2 |
| Id1 | Lrg1 |
| Krt19 | Slc45a3 |
| Pex26 | Tsc22d3 |
| Osgin1 | Ttpa |
| Srm | Gldc |
| Acat2 | Insig2 |
| Gm2000 | Sec14l4 |
| Hmgcs1 | Mvk |
| Idi1 | Agtr1a |

| CR v AL down | FRF v AL down |
|---|---|
| Slc10a2 | Slc10a2 |
| Slc17a3 | Slc17a3 |
| Slc22a28 | Slc22a28 |
| Slc22a30 | Slc22a30 |
| Slc25a25 | Slc25a25 |
| Slco1a1 | Slco1a1 |
| Socs2 | Socs2 |
| Srebf1 | Srebf1 |
| Sucnr1 | Sucnr1 |
| Tcap | Tcap |
| Tcea3 | Tcea3 |
| Thrsp | Thrsp |
| Tmem150a | Tmem150a |
| Tnnc2 | Tnnc2 |
| Tob1 | Tob1 |
| Ugt2b38 | Ugt2b38 |
| Zfp36 | Zfp36 |
| Mup15 | Cyp2c44 |
| Mup17 | Scd1 |
| Mup11 | Serpina6 |
| Mup2 | Mid1ip1 |
| Mup14 | Angptl8 |
| Mup9 | Ces2a |
| Sult5a1 | Gm3776 |
| Ugt3a1 | Inmt |
| Tff3 | Lgals1 |
| Mup18 | Aqp8 |
| Mup19 | Acta1 |
| Jun | Mb |
| Calr | Tnni2 |
| Sdf2l1 | Tnnt3 |
| Enho | Tmsb10 |
| Mup13 | Acaa1b |
| Mup8 | Cyp26a1 |
| Dio1 | Efna1 |
| Pim3 | Id3 |
| Actb | Sult2a5 |
| Derl3 | Tgm2 |
| Hsd3b7 | Cd74 |
| Paqr9 | Gstp3 |
| Slc35b1 | H2-Aa |
| Tmem254a | H2-Eb1 |
| Tmem254b | Ifi47 |
| Cyp2d40 | Cyp2c55 |
| Gadd45g | Raet1d |

| CR v AL up | FRF v AL down |
|---|---|
| Slc22a5 | Onecut1 |
| St3gal5 | Pgam2 |
| Alas1 | Phlda1 |
| Bag3 | Slc41a2 |
| Cbr3 | Tgm1 |
| Dnajb9 | Tmsb4x |
| Hist2h3c2 | Agxt |
| Hsd17b13 | Btg2 |
| Id1 | C6 |
| Krt19 | Cyp2u1 |
| Pex26 | Cyp3a44 |
| Slc20a1 | Cyp4a12a |
| Creld2 | Cyp4a12b |
| Osgin1 | Eif2s3y |
| Srm | Gm3839 |
| Tmem254c | H2-Ab1 |
| Acat2 | Id2 |
| Cyp51 | Mmd2 |
| Gm2000 | Mup1 |
| Hmgcs1 | Mup12 |
| Idi1 | Mup20 |
| Lss | Mup21 |
| Ppp1r1b | Mup7 |
| Rdh11 | Nat8 |
| Serpina3m | Nat8f5 |
| Clpx | Ppp1r3g |
| Gm10273 | Psmb10 |
| Lsm7 | Saa3 |
| Trib3 | Serpina1e |
| Akr1b7 | Smim22 |
| Cd36 | Spsb4 |
| Gpihbp1 | Acy3 |
| Mgst3 | Bhlhe40 |
| Tgtp1 | Cyr61 |
| Apoa4 | Gclc |
| Cyp4a10 | Jchain |
| Phospho1 | Keg1 |
| Slc25a33 | Nr4a1 |
| 1110034G24Rik | Gstm6 |
| 1810055G02Rik | Mug2 |
| Aass | Serpina7 |
| Aifm2 | Bdh2 |
| Amy2a5 | Fabp1 |
| Atl2 | Hamp2 |
| Cdkn1a | Mcm10 |

| CR v AL down | FRF v AL up |
|---|---|
| Tmem254b | Creld2 |
| Adh4 | Cyp51 |
| Car3 | Fam35a |
| Cfd | Hsd17b10 |
| Cyp2d40 | Hsp90ab1 |
| Ppp1r3g | Slc16a7 |
| Serpina1e | Sult1e1 |
| Slc22a7 | 1600002H07Rik |
| Tlcd2 | As3mt |
| Kyat1 | Fam234b |
| Mafb | Fgf21 |
| Plk3 | Nr1i3 |
| Slc25a30 | Slco1a4 |
| Cyp3a44 | Sorbs3 |
| RP24-346O11.2 | Sult1a1 |
| Csrnp1 | Ccng2 |
| Onecut1 | Fmo2 |
| Fasn | Lrg1 |
| Apon | Serinc3 |
| Cldn1 | Tsc22d3 |
| Ddc | Ttpa |
| Dexi | Acot1 |
| Dnajb11 | Ethe1 |
| Eef1a2 | Gldc |
| Gadd45a | Insig2 |
| Gck | Sec14l4 |
| Gtf3c6 | Slc8b1 |
| Insc | Mvk |
| Ly6d | Agtr1a |
| Mlec | Cpsf4l |
| Myh2 | Crot |
| Myl2 | Fbxo21 |
| Nat8f2 | Fus |
| Nsmf | Gm5096 |
| Oasl1 | Tcp11l2 |
| Psen2 | A930011G23Rik |
| Slc17a3 | Ciart |
| Suox | Nrg4 |
| Tubb2a | Usp50 |
| Ugt2b37 | Chordc1 |
| Cebpb | Cyp2b13 |
| Dusp6 | Dhcr24 |
| Fabp2 | Mgll |
| Mup16 | Slc35g1 |
| N4bp2l1 | Tmem254c |

Supplementary Table 2 for Fig 3D-G

| CR v AL up | FRF v AL up |
|---|---|
| Ppp1r1b | Cpsf4l |
| Rdh11 | Crot |
| Clpx | Fus |
| Gm10273 | Gm5096 |
| Trib3 | Tcp11l2 |
| Akr1b7 | A930011G23Rik |
| Cd36 | Ciart |
| Gpihbp1 | Nrg4 |
| Gstm1 | Usp50 |
| Mgst3 | Chordc1 |
| Raet1d | Cyp2b13 |
| Tgtp1 | Dhcr24 |
| Slc25a33 | Hmgcr |
| 1110034G24Rik | Mgll |
| Cidec | Slc35g1 |
| Ddhd2 | Apoa5 |
| Fitm2 | Arg1 |
| Gpcpd1 | E030018B13Rik |
| Hsdl2 | Gpt2 |
| Plpp1 | Igfbp1 |
| Ppp2r2d | Pck1 |
| Scyl2 | Per1 |
| Sgk2 | Amy1 |
| Sult2a5 | Asl |
| Ten1 | Coa6 |
| Tmed5 | Fgf1 |
| | Fmo5 |
| | Ivns1abp |
| | Nampt |
| | Parp16 |
| | Pfkfb1 |
| | Ppp1r3b |
| | Rhbg |
| | Zbtb16 |
| | Ahsa1 |
| | Arhgef19 |
| | Atp2a2 |
| | Caprin1 |
| | Car14 |
| | Crip2 |
| | Ddit3 |
| | Dnajb1 |
| | Dnajb2 |
| | Hfe2 |
| | Hsp90b1 |

| CR v AL down | FRF v AL down |
|---|---|
| Slc22a7 | Vegfb |
| Tlcd2 | BC048546 |
| Slc25a30 | Hao2 |
| RP24-346O11.2 | Rnf186 |
| Apon | Slc9a3r1 |
| Car14 | A230050P20Rik |
| Cldn1 | Aacs |
| Ddc | Gas6 |
| Dexi | Slc41a2 |
| Dnajb11 | Tgm1 |
| Gck | Tmsb4x |
| Gtf3c6 | Agxt |
| Hfe2 | C6 |
| Leap2 | Cyp2u1 |
| Mlec | Eif2s3y |
| Nat8f2 | Gm3839 |
| Nsmf | H2-Ab1 |
| Psen2 | Id2 |
| Suox | Mmd2 |
| Tubb2a | Nat8 |
| Tubb4b | Psmb10 |
| Ugt2b37 | Smim22 |
| Csrp3 | Spsb4 |
| Fabp2 | Acy3 |
| Igfbp1 | Gclc |
| Mup16 | Gsta2 |
| N4bp2l1 | Jchain |
| Npr2 | Gsta1 |
| Slc13a3 | Gstm6 |
| Slc38a2 | Bdh2 |
| Tat | Fabp1 |
| Trp53inp1 | Hamp2 |
| Upp2 | Mcm10 |
| Fos | Sult2a3 |
| Jund | Steap4 |
| Mup10 | Tk1 |
| Rarres1 | Tspan4 |
| Tsku | Acss2 |
| Mup6 | Erdr1 |
| Hsd17b2 | Asns |
| Insig1 | Atp4a |
| Pdia4 | C1qa |
| Sc5d | Cd52 |
| Arl4a | Ctss |
| Cmah | Cxcl9 |

| CR v AL up | FRF v AL down |
|---|---|
| Cidec | Saa1 |
| Cpt1a | Inhbc |
| Ddhd2 | Slc10a2 |
| Fam134b | Slc17a3 |
| Fitm2 | Steap4 |
| Gde1 | Tk1 |
| Glrx | Tspan4 |
| Gpcpd1 | Zfp36 |
| Grpel2 | Acss2 |
| Hsdl2 | Erdr1 |
| Lepr | Pcsk9 |
| Map1lc3b | Socs2 |
| Mat1a | Cyp4f14 |
| Mt1 | Asns |
| Nceh1 | Atp4a |
| Plpp1 | C1qa |
| Ppp2r2d | Cd52 |
| Scyl2 | Ctss |
| Serinc3 | Cxcl9 |
| Sgk2 | Elovl3 |
| Tdo2 | Pde4b |
| Ten1 | Sik1 |
| Tmed5 | BC029214 |
| Tmem120a | Dusp6 |
| Tmem56 | Nme6 |
| Try4 | Nr0b2 |
| | Paox |
| | Pdcd4 |
| | 5830473C10Rik |
| | Amdhd1 |
| | Arrdc3 |
| | Cbs |
| | Cxcl1 |
| | Gstm4 |
| | Klf10 |
| | St6gal1 |
| | Abcc2 |
| | Abcg5 |
| | Abcg8 |
| | Bdh1 |
| | Cebpb |
| | Ces1e |
| | Csrnp1 |
| | Cyp2g1 |
| | Fdx1 |

| CR v AL down | FRF v AL up |
|---|---|
| Npr2 | Apoa5 |
| Pgam2 | Arg1 |
| Slc13a3 | E030018B13Rik |
| Slc38a2 | Gpt2 |
| Tat | Hbb-bs |
| Upp2 | Pck1 |
| Mapk15 | Per1 |
| Nat8f1 | Txnip |
| Cyp4a12a | Aifm2 |
| Cyp4a12b | Amy1 |
| Elovl3 | Asl |
| Fos | Coa6 |
| Jund | Fgf1 |
| Mup10 | Fmo5 |
| Mup20 | Gde1 |
| Rarres1 | Ivns1abp |
| Slco1a1 | Lsm7 |
| Cyr61 | Nampt |
| G0s2 | Parp16 |
| Keg1 | Pfkfb1 |
| Inhbc | Ppp1r3b |
| Cish | Rhbg |
| Mup6 | Zbtb16 |
| Slc22a28 | Ahsa1 |
| Acly | Arhgef19 |
| Cyp4f14 | Atp2a2 |
| Fam25c | Caprin1 |
| Hsd17b2 | Crip2 |
| Insig1 | Ddit3 |
| Sc5d | Dnajb1 |
| Cmah | Dnajb2 |
| Cyp2d37-ps | Dnajb9 |
| Fermt2 | Dynll1 |
| Grb7 | Hist2h3c2 |
| Iigp1 | Hsp90b1 |
| Irs2 | Hspa5 |
| Klf15 | Hyou1 |
| Nfil3 | Lrrc59 |
| Nr0b2 | Mal2 |
| Pnrc1 | Manf |
| Rsrp1 | Morf4l2 |
| Sdc4 | Pnkd |
| Serpinb1a | Rtp3 |
| Ubb | Slc20a1 |
| Wsb1 | Snrpa |

Supplementary Table 2 for Fig 3D-G

| CR v AL up | FRF v AL up |
|---|---|
| | Hspa5 |
| | Hyou1 |
| | Leap2 |
| | Lrrc59 |
| | Mal2 |
| | Manf |
| | Morf4l2 |
| | Mvd |
| | Pdia4 |
| | Pnkd |
| | Rtp3 |
| | Snrpa |
| | Sqle |
| | Srxn1 |
| | Syvn1 |
| | Tardbp |
| | Tubb4b |
| | Ubc |
| | Ubxn4 |
| | 2210010C04Rik |
| | A1cf |
| | Aaed1 |
| | Abcb4 |
| | Acacb |
| | Acadm |
| | Acot12 |
| | Acot3 |
| | Acss3 |
| | Agpat3 |
| | Cd163 |
| | Cel |
| | Cela3b |
| | Ces1b |
| | Chic2 |
| | Cpa1 |
| | Cpa2 |
| | Cpb1 |
| | Cxcl13 |
| | Cyp2b10 |
| | D230025D16Rik |
| | Defb1 |
| | Dnmbp |
| | Ei24 |
| | Gadd45g |
| | Hacl1 |

| CR v AL down | FRF v AL down |
|---|---|
| Cyp2d37-ps | Pde4b |
| D230025D16Rik | BC029214 |
| Fermt2 | Nme6 |
| Grb7 | Paox |
| Iigp1 | Pdcd4 |
| Ip6k2 | 5830473C10Rik |
| Irs2 | Amdhd1 |
| Klf15 | Arrdc3 |
| Rnf125 | Cbs |
| Rsrp1 | Gstm1 |
| Sdc4 | Gstm3 |
| Serpinb1a | Gstm4 |
| Slc45a3 | Klf10 |
| Ubb | St6gal1 |
| Ubc | Abcc2 |
| Wsb1 | Abcg5 |
| Fst | Bdh1 |
| Me1 | Ces1e |
| Rapgef4 | Cyp2a5 |
| Trp53inp2 | Cyp2g1 |
| Hopx | Fdx1 |
| Trib1 | Foxq1 |
| Errfi1 | Gamt |
| Inhbe | Gnat1 |
| Obp2a | Gpr146 |
| Stip1 | Gsta4 |
| Cela2a | Gys2 |
| Ces3b | Junb |
| Cyp3a41b | Pigp |
| Rgs3 | Pltp |
| Saa4 | Rbp1 |
| Serpina1c | Rpa3 |
| Cldn2 | S100a11 |
| Cyp7a1 | Slc22a29 |
| Esr1 | Slc38a3 |
| Fdps | Slc7a2 |
| Hmgcr | Spp1 |
| Igfals | Sult1c2 |
| Igtp | Sult2a1 |
| Msmo1 | Sult2a2 |
| Mvd | Tef |
| Nsdhl | Tmem50a |
| Sult3a2 | Tstd1 |
| Urad | Uox |
| Acat3 | Ypel3 |

| CR v AL up | FRF v AL down |
|---|---|
| | Foxq1 |
| | G6pc |
| | Gamt |
| | Gnat1 |
| | Gpr146 |
| | Gsta4 |
| | H2afv |
| | Junb |
| | Mafb |
| | Pigp |
| | Plk3 |
| | Pltp |
| | Rbp1 |
| | Rpa3 |
| | S100a11 |
| | Saa2 |
| | Slc22a29 |
| | Slc25a25 |
| | Slc38a3 |
| | Slc7a2 |
| | Spp1 |
| | Sult1c2 |
| | Sult2a1 |
| | Sult2a2 |
| | Tef |
| | Tmem50a |
| | Tob1 |
| | Tstd1 |
| | Uox |
| | Ypel3 |
| | Ak1 |
| | Amd1 |
| | Apol9a |
| | Casq1 |
| | Ces2c |
| | Comtd1 |
| | Cxx1b |
| | Cyp1a2 |
| | Cyp2c67 |
| | Cyp2c70 |
| | Cyp7b1 |
| | Fam47e |
| | Hdhd3 |
| | Hsd3b2 |
| | Hspb7 |

| CR v AL down | FRF v AL up |
|---|---|
| Abcg8 | Sqle |
| Fst | Srxn1 |
| H2afv | Tardbp |
| Me1 | Tuba8 |
| Saa3 | Ubxn4 |
| Trp53inp2 | 2210010C04Rik |
| Trib1 | A1cf |
| Errfi1 | Aaed1 |
| Inhbe | Aass |
| Obp2a | Abcb4 |
| Ugt2b38 | Acacb |
| Cxcl1 | Acadm |
| Gdf15 | Acot12 |
| Stip1 | Acot2 |
| Cela2a | Acot3 |
| Ces3b | Acot4 |
| Cyp3a41b | Acss3 |
| Fads2 | Agpat3 |
| Gstp2 | Amy2a5 |
| Mug2 | Cd163 |
| Mup21 | Cel |
| Nat8f5 | Cela3b |
| Rgs3 | Ces1b |
| Saa4 | Ces1g |
| Serpina1c | Chic2 |
| Cldn2 | Cpa1 |
| Cyp7a1 | Cpa2 |
| Elovl6 | Cpb1 |
| Esr1 | Crat |
| Fabp5 | Cxcl13 |
| Fdps | Cyp2b10 |
| Igfals | Cyp3a59 |
| Igtp | Defb1 |
| Msmo1 | Dnmbp |
| Phlda1 | Ei24 |
| Saa2 | Gadd45b |
| Socs2 | Grpel2 |
| Urad | Hacl1 |
| Acat3 | Hmgcs2 |
| Akr1c6 | Klf9 |
| Apol7a | Lcn2 |
| Apol9b | Lepr |
| Fam47e | Map1lc3b |
| Gm4756 | Mt1 |
| Hsd3b2 | Nceh1 |

Supplementary Table 2 for Fig 3D-G

| CR v AL up | FRF v AL up |
|---|---|
| | Hmgcs2 |
| | Klf9 |
| | Lcn2 |
| | Ncoa4 |
| | Pcx |
| | Plscr2 |
| | Pnliprp1 |
| | Ppara |
| | Prss2 |
| | Rabgef1 |
| | Rhobtb1 |
| | Rnase1 |
| | Sco2 |
| | Slc25a22 |
| | Sult3a2 |
| | Sun2 |
| | Sycn |
| | Tm4sf4 |
| | Tmem106b |
| | Vnn1 |
| | Vnn3 |
| | Wbp1l |
| | Zg16 |
| | Cd9 |
| | Chil3 |
| | Csad |
| | Fdft1 |
| | Htatip2 |
| | Rnf125 |
| | Bri3 |
| | Hopx |
| | Hsd17b7 |
| | Lhpp |
| | Nsdhl |

| CR v AL down | FRF v AL down |
|---|---|
| Akr1c6 | Ak1 |
| Apol7a | Amd1 |
| Apol9b | Casq1 |
| Gm4756 | Ces2c |
| Hsd3b3 | Comtd1 |
| Irf5 | Cxx1b |
| Nit2 | Cyp1a2 |
| Nudt7 | Cyp2c67 |
| Pde9a | Cyp7b1 |
| Ppp1r3c | Hspb7 |
| Pter | Ociad2 |
| Syvn1 | Pygm |
| Tmem86b | Qprt |
| Tsc22d1 | Sec23b |
| | Srsf3 |
| | Tbcb |
| | Tuba1b |
| | Tuba4a |
| | Tubb5 |
| | Txndc5 |
| | Ube2l6 |
| | Zbp1 |
| | Fmo3 |
| | Myl3 |
| | Pvalb |
| | Acmsd |
| | Coq8a |
| | Crem |
| | Mme |
| | Zfp36l1 |

| CR v AL up | FRF v AL down |
|---|---|
| | Ly6d |
| | Oasl1 |
| | Ociad2 |
| | Pdia6 |
| | Pygm |
| | Qprt |
| | Sec23b |
| | Slc22a28 |
| | Slc22a30 |
| | Slco1a1 |
| | Tbcb |
| | Tcea3 |
| | Tuba1b |
| | Tuba4a |
| | Tubb5 |
| | Txndc5 |
| | Ube2l6 |
| | Ugt2b38 |
| | Zbp1 |
| | Fads2 |
| | Myl3 |
| | Noct |
| | Pvalb |
| | Acmsd |
| | Coq8a |
| | Crem |
| | Mme |
| | Nfil3 |
| | Pnrc1 |
| | Zfp36l1 |

| CR v AL down | FRF v AL up |
|---|---|
| Hsd3b3 | Ncoa4 |
| Irf5 | Pcx |
| Nit2 | Plscr2 |
| Nudt7 | Pnliprp1 |
| Pcsk9 | Ppara |
| Pde9a | Prss2 |
| Ppp1r3c | Rabgef1 |
| Pter | Reg1 |
| Serpina7 | Rhobtb1 |
| Sucnr1 | Rnase1 |
| Tcea3 | Sco2 |
| Tmem150a | Slc25a22 |
| Tmem86b | Sun2 |
| Tsc22d1 | Sycn |
| | Tdo2 |
| | Tm4sf4 |
| | Tmem106b |
| | Tmem56 |
| | Try4 |
| | Vnn1 |
| | Vnn3 |
| | Wbp1l |
| | Zg16 |
| | Cd9 |
| | Chil3 |
| | Csad |
| | Fdft1 |
| | Htatip2 |
| | Lss |
| | Phospho1 |
| | Aprt |
| | Bri3 |
| | Hsd17b7 |
| | Lhpp |
