## Supplementary Table 3 for "Anticipatory metabolic reprogramming distinguishes caloric restriction from fasting-refeeding cycles"

Supplementary Table 3 DEG_Categories_- 2 hrs for Suppl Fig 3A
Differentially expressed genes (FDR < 0.05)

| CRvsAL Exclusive DEGs | | |
|---|---|---|
| Gene.ID | FDR | FC |
| A1bg | 0.000161468 | 0.022494117 |
| Actb | 2.1123E-06 | 0.462234901 |
| Actg1 | 0.000312774 | 0.256448608 |
| Apol9a | 3.39328E-05 | 0.361855082 |
| Cldn1 | 3.86996E-05 | 0.356644807 |
| Cyp2c70 | 6.64135E-13 | 0.45547095 |
| Ddc | 0.008872595 | 0.4298744 |
| Dexi | 0.000232498 | 0.476455319 |
| Dnajb11 | 0.02927422 | 0.440632056 |
| Eef1a2 | 0.012722121 | 0.12783209 |
| Gtf3c6 | 0.000515649 | 0.457003141 |
| Hdhd3 | 2.29736E-05 | 0.277377563 |
| Hist1h1c | 0.021091703 | 0.302694247 |
| Lasp1 | 9.68636E-05 | 0.485501816 |
| Ly6d | 0.015948297 | 0.090509239 |
| Mlec | 0.006929145 | 0.456945448 |
| Nsmf | 3.10193E-06 | 0.434075827 |
| Oasl1 | 0.018491822 | 0.300625183 |
| Paqr9 | 0.000211454 | 0.443709606 |
| Psen2 | 0 | 0.462170401 |
| Saa1 | 0.01042045 | 0.338947616 |
| Slc22a30 | 1.4426E-05 | 0.468230013 |
| Slc35b1 | 0.018531248 | 0.448292155 |
| Sult5a1 | 0.002386219 | 0.28118253 |
| Suox | 2.02061E-14 | 0.471062146 |
| Tff3 | 0.009953766 | 0.101286331 |
| Tmem254a | 3.41443E-08 | 0.46157796 |
| Tmem254b | 3.41443E-08 | 0.46157796 |
| Tuba1c | 8.90557E-06 | 0.371654328 |
| Tubb4b | 8.91422E-07 | 0.454756233 |
| Ugt2b37 | 0.031303043 | 0.196850359 |
| Aldoc | 0.03449837 | 2.468244311 |
| Bnip3 | 0 | 2.040963693 |
| Btbd19 | 0.034198831 | 6.11296469 |
| Cnppd1 | 0.044049427 | 2.049069969 |
| Ctgf | 0.023117743 | 2.762446575 |
| Eif4b | 6.03878E-08 | 2.036661649 |
| Fbxo31 | 0.00229881 | 2.809205215 |
| Fkbp5 | 0.034471522 | 2.933677653 |

Supplementary Table 3 DEG_Categories_- 2 hrs for Suppl Fig 3A
Differentially expressed genes (FDR < 0.05)

| CRvsAL Exclusive DEGs | | |
|---|---|---|
| Gene.ID | FDR | FC |
| Fmo3 | 0.004214822 | 14.92548587 |
| Gstt2 | 0 | 2.537481604 |
| Hba-a2 | 0.039178615 | 4.600116094 |
| Hbb-bs | 0.006178635 | 4.606085079 |
| Hbb-bt | 0.037179066 | 3.749696997 |
| Il6ra | 0.003358306 | 2.456574411 |
| Lpin2 | 0 | 3.958765011 |
| Lurap1l | 0.005711281 | 2.019847262 |
| Mfsd2a | 4.17264E-05 | 2.568720052 |
| Pck1 | 0.018059189 | 2.194503828 |
| Pcsk4 | 0.005070984 | 3.544167656 |
| Pla2g12a | 0.046725244 | 2.105916719 |
| Por | 4.97783E-06 | 2.233938769 |
| Ppp1r10 | 5.60885E-05 | 3.073917385 |
| Rpl22l1 | 2.67509E-10 | 2.225147651 |
| Slc16a7 | 0.006054518 | 2.217238353 |
| Slc8b1 | 1.6406E-06 | 2.646000484 |
| Slco1a4 | 0.002569096 | 2.332368301 |
| Soat2 | 0.00562905 | 2.82479667 |
| Sorbs3 | 0.007890249 | 2.667211321 |
| Sult1e1 | 0.023935219 | 7.813292957 |
| Txnip | 0.001040081 | 2.617121157 |

Supplementary Table 3 DEG_Categories_- 2 hrs for Suppl Fig 3A
Differentially expressed genes (FDR < 0.05)

| FRFvsAL Exclusive DEGs | | |
|---|---|---|
| Gene.ID | FDR | FC |
| Acaa1b | 1.1073E-05 | 0.473738197 |
| Bag3 | 0.003814494 | 0.459911213 |
| Ces2a | 3.23075E-13 | 0.496973058 |
| Cfd | 0.013702161 | 0.11679973 |
| Cry1 | 1.64863E-05 | 0.269149616 |
| Cyp26a1 | 0.006114472 | 0.104260966 |
| Cyp2c44 | 0.000170437 | 0.480099592 |
| Dnaja1 | 0.013883616 | 0.424184935 |
| Efna1 | 0.000921411 | 0.416100227 |
| Elovl6 | 0.044019027 | 0.353320364 |
| Fkbp4 | 0 | 0.394565996 |
| Hspa8 | 3.13235E-08 | 0.396975371 |
| Hsph1 | 0.00056109 | 0.225123905 |
| Inhbc | 0.009604386 | 0.382992237 |
| Palld | 0.026755655 | 0.430433335 |
| Ppp1r3c | 0.02771732 | 0.380840988 |
| Slc9a3r1 | 4.03166E-12 | 0.444377122 |
| Steap4 | 7.34615E-07 | 0.451211893 |
| Stip1 | 1.21935E-09 | 0.401526147 |
| Tk1 | 2.93913E-05 | 0.482910781 |
| Tmem150a | 0.000396882 | 0.498936693 |
| Zfp36 | 0.019214479 | 0.464682698 |
| Abca6 | 8.75696E-07 | 2.089594264 |
| Agtr1a | 8.25078E-07 | 2.312696918 |
| Cdip1 | 9.33631E-12 | 2.41247293 |
| Cdo1 | 0 | 2.376222301 |
| Cpsf4l | 0.001949451 | 3.26825245 |
| Cpt1a | 9.49604E-05 | 2.041189274 |
| Cyp4a10 | 0.029016693 | 3.830200421 |
| Cyp4a14 | 0.013386647 | 4.726332309 |
| Eif4ebp3 | 0.001476512 | 4.867962874 |
| Fam134b | 0 | 3.963990886 |
| Fus | 1.04706E-05 | 2.009767336 |
| Got1 | 3.04201E-14 | 2.946162709 |
| Gpcpd1 | 0.001884775 | 4.618087486 |
| Inca1 | 9.78493E-07 | 4.731990658 |
| Lbp | 0.022231823 | 2.066888131 |
| Nlrp12 | 5.92515E-06 | 5.08055396 |
| Npc1 | 0.000198371 | 2.027295931 |

Supplementary Table 3 DEG_Categories_- 2 hrs for Suppl Fig 3A
Differentially expressed genes (FDR < 0.05)

| FRFvsAL Exclusive DEGs | | |
|---|---|---|
| Gene.ID | FDR | FC |
| Rgs16 | 3.90671E-10 | 8.239430288 |
| Serpina3n | 0.000202565 | 2.133737389 |
| Slc10a1 | 0.047152797 | 2.394304527 |
| Slc25a47 | 4.08665E-06 | 2.770745845 |
| Tcp11l2 | 0.015286227 | 2.651725822 |

Supplementary Table 3 DEG_Categories_- 2 hrs for Suppl Fig 3A
Differentially expressed genes (FDR < 0.05)

| CRvsFRF Exclusive DEGs | | |
|---|---|---|
| Gene.ID | FDR | FC |
| 1600002H07Rik | 0.018559604 | 0.452347685 |
| Abcb11 | 0.003201232 | 0.486712993 |
| Acox2 | 4.77688E-08 | 0.495885391 |
| Apof | 2.71426E-08 | 0.479727278 |
| BC004004 | 0.000119277 | 0.490574797 |
| Camk1d | 0.030763156 | 0.369702662 |
| Cbs | 7.26816E-05 | 0.355830088 |
| Cth | 0 | 0.475728863 |
| Cyp4a31 | 0.039317182 | 0.309032649 |
| E030018B13Rik | 0.023600393 | 0.174696313 |
| Enho | 0.00556961 | 0.474193052 |
| Ethe1 | 1.65751E-05 | 0.376176254 |
| Gpd1 | 0.003889146 | 0.444121994 |
| Ifi47 | 0.006412339 | 0.335222836 |
| Iigp1 | 0.001160705 | 0.499905943 |
| Insig2 | 4.44529E-06 | 0.29997351 |
| Kynu | 1.33017E-11 | 0.415084015 |
| Lgals9 | 3.0726E-10 | 0.470219376 |
| Mup1 | 0.041749814 | 0.076064648 |
| Mup18 | 5.32898E-05 | 0.201370728 |
| Mup7 | 0.013755367 | 0.046550037 |
| Phospho1 | 0.000147929 | 0.199202586 |
| Prkd3 | 1.82925E-09 | 0.470386882 |
| RP24-346O11.2 | 0.045639573 | 0.19796309 |
| S100a10 | 2.02717E-06 | 0.276069336 |
| Sdr42e1 | 0.000524645 | 0.441309766 |
| Slc22a7 | 0.006771986 | 0.370665882 |
| Stra6l | 1.15424E-07 | 0.389295123 |
| Tmem86b | 1.6035E-12 | 0.482955953 |
| Urad | 1.07899E-07 | 0.482273966 |
| Abcb1a | 0.038306453 | 3.294462594 |
| Acot1 | 0.038122976 | 3.12105924 |
| Acot4 | 0.003312447 | 2.057046756 |
| Adh1 | 0 | 2.017093528 |
| Akr1c19 | 0.005286235 | 2.09422977 |
| Alas1 | 1.11241E-05 | 2.75154419 |
| Bud13 | 0.040436944 | 4.825060336 |
| Ces1d | 0 | 3.363580808 |
| Efhd2 | 0.00014272 | 2.085679817 |

Supplementary Table 3 DEG_Categories_- 2 hrs for Suppl Fig 3A
Differentially expressed genes (FDR < 0.05)

| CRvsFRF Exclusive DEGs | | |
|---|---|---|
| Gene.ID | FDR | FC |
| Etfbkmt | 0.007271915 | 2.41425428 |
| Fam195a | 1.89719E-09 | 2.028162067 |
| Fcgr2b | 0.000737644 | 2.017272587 |
| Galk1 | 1.51562E-08 | 2.126641945 |
| Gm20594 | 1.46673E-08 | 4.490513981 |
| Gsta2 | 0.000101493 | 3.514994695 |
| Gstm3 | 0.048893279 | 2.250595214 |
| Id1 | 0.000519679 | 2.62850646 |
| Id3 | 0.000319344 | 2.068180071 |
| Nmrk1 | 0.000343369 | 2.328113396 |
| Nrg4 | 2.39889E-06 | 3.815830156 |
| Nutf2 | 0.000683667 | 6.032097061 |
| Reln | 0.009118074 | 2.538385583 |
| Slc43a3 | 0.034333094 | 2.109643543 |
| Slc6a9 | 0.016142826 | 2.643142255 |
| Smim22 | 1.1115E-05 | 3.177529575 |
| Sult2a5 | 0.003468962 | 10.96574496 |
| Tstd1 | 0.001177527 | 2.638285277 |
| Ugt1a5 | 0.001621243 | 3.965055718 |
| Vpreb3 | 0.002109947 | 5.327495852 |
| Zfp36l2 | 0.021352665 | 2.144043091 |
| Zfp970 | 0.001453281 | 2.23301753 |

Supplementary Table 3 DEG_Categories_- 2 hrs for Suppl Fig 3A
Differentially expressed genes (FDR < 0.05)

| CRvsAL Shared DEGs | | |
| --- | --- | --- |
| Gene.ID | FDR | FC |

| FRFvsAL Shared DEGs | | |
| --- | --- | --- |
| Gene.ID | FDR | FC |

| CRvsFRF Shared DEGs | | |
| --- | --- | --- |
| Gene.ID | FDR | FC |

Supplementary Table 3 DEG_Categories_- 2 hrs for Suppl Fig 3A
Differentially expressed genes (FDR < 0.05)

| CRvsAL Shared DEGs | | |
|---|---|---|
| Gene.ID | FDR | FC |
| Acta1 | 0 | 0.007112479 |
| Angptl8 | 0.019887877 | 0.174875998 |
| Ankrd23 | 0.033019047 | 0.139115508 |
| Atp2a1 | 0 | 0.030606404 |
| Ckm | 1.09464E-06 | 0.015119127 |
| Ckmt2 | 1.44329E-15 | 0.043521007 |
| Cox6a2 | 0.000436964 | 0.058978239 |
| Cox8b | 1.84666E-07 | 0.062097443 |
| Dusp1 | 0.009074032 | 0.480545347 |
| Fabp3 | 3.64921E-05 | 0.031758607 |
| Gadd45a | 0.008267785 | 0.276466072 |
| Gale | 4.59787E-05 | 0.183769259 |
| Gck | 0.001752063 | 0.430604005 |
| Hes6 | 0.019363784 | 0.454088223 |
| Hsp90aa1 | 0.000697248 | 0.475461824 |
| Insc | 0.000120306 | 0.288287297 |
| Lifr | 0.000493267 | 0.302207225 |
| Mb | 0 | 0.009809615 |
| Mid1ip1 | 1.69715E-05 | 0.314977717 |
| Myh1 | 0.00038161 | 0.075334998 |
| Myh2 | 0.001837762 | 0.078809804 |
| Myl1 | 0 | 0.038215964 |
| Myl2 | 6.78106E-10 | 0.055026425 |
| Mylpf | 0.000481439 | 0.046507667 |
| Nrep | 0.000233615 | 0.355524411 |
| Rcan1 | 4.30415E-06 | 0.381332338 |
| Slc10a2 | 0.000173244 | 0.197677963 |
| Slc17a3 | 0.041565037 | 0.491317906 |
| Srebf1 | 0.000314199 | 0.265132787 |
| Tcap | 2.13938E-05 | 0.030526018 |
| Thrsp | 0.031312332 | 0.462200809 |
| Tnnc2 | 0 | 0.013324462 |
| Tnni2 | 0 | 0.01856975 |
| Tnnt3 | 0 | 0.012516599 |
| Tubb2a | 2.40418E-10 | 0.346535616 |
| Angptl4 | 8.47803E-05 | 3.012691692 |
| Fbxo21 | 0.004262808 | 2.145852168 |
| Gabarapl1 | 0.000135676 | 2.043413157 |
| Gm45531 | 2.83497E-10 | 6.151280272 |

| FRFvsAL Shared DEGs | | |
|---|---|---|
| Gene.ID | FDR | FC |
| Acta1 | 1.66895E-07 | 0.008395566 |
| Angptl8 | 0.002797083 | 0.123075498 |
| Ankrd23 | 0.045375909 | 0.135794881 |
| Atp2a1 | 4.52864E-05 | 0.031739565 |
| Ckm | 0 | 0.014101846 |
| Ckmt2 | 1.88738E-15 | 0.043521007 |
| Cox6a2 | 5.73258E-09 | 0.057239703 |
| Cox8b | 2.29927E-07 | 0.062097443 |
| Dusp1 | 9.88762E-08 | 0.316582384 |
| Fabp3 | 0 | 0.028665514 |
| Gadd45a | 0.000689356 | 0.188157743 |
| Gale | 0.00415704 | 0.285510673 |
| Gck | 0.001012565 | 0.380227599 |
| Hes6 | 0.000476906 | 0.320867144 |
| Hsp90aa1 | 5.77467E-11 | 0.332673149 |
| Insc | 0.000394922 | 0.344926719 |
| Lifr | 0.005646792 | 0.330666986 |
| Mb | 0 | 0.009809615 |
| Mid1ip1 | 0.037609381 | 0.49523516 |
| Myh1 | 0.000466047 | 0.075334998 |
| Myh2 | 0.002234822 | 0.078809804 |
| Myl1 | 0 | 0.038215964 |
| Myl2 | 8.5704E-10 | 0.055026425 |
| Mylpf | 0.003651198 | 0.060367553 |
| Nrep | 1.33606E-09 | 0.175634531 |
| Rcan1 | 0.000786201 | 0.466350647 |
| Slc10a2 | 0.002614387 | 0.245881147 |
| Slc17a3 | 0.013807243 | 0.48567331 |
| Srebf1 | 1.3874E-05 | 0.181329043 |
| Tcap | 0 | 0.030102349 |
| Thrsp | 0.001050405 | 0.305227559 |
| Tnnc2 | 3.92076E-06 | 0.016271696 |
| Tnni2 | 4.80162E-05 | 0.026050579 |
| Tnnt3 | 0 | 0.012516599 |
| Tubb2a | 0.000585632 | 0.38431988 |
| Angptl4 | 3.24393E-07 | 2.691177251 |
| Fbxo21 | 5.50671E-09 | 3.02520238 |
| Gabarapl1 | 8.43274E-11 | 2.295809614 |
| Gm45531 | 1.24979E-11 | 7.452045088 |

Supplementary Table 3 DEG_Categories_- 2 hrs for Suppl Fig 3A
Differentially expressed genes (FDR < 0.05)

| CRvsAL Shared DEGs | | |
|---|---|---|
| Gene.ID | FDR | FC |
| Lpin1 | 2.149E-06 | 7.04833915 |
| Pnpla2 | 4.2866E-10 | 2.306619815 |
| Rab30 | 0.036352634 | 3.099032407 |
| Rbm3 | 0 | 3.005465789 |

| FRFvsAL Shared DEGs | | |
|---|---|---|
| Gene.ID | FDR | FC |
| Lpin1 | 0.037279033 | 6.550686774 |
| Pnpla2 | 4.24255E-08 | 2.7105205 |
| Rab30 | 0.000961856 | 2.858851491 |
| Rbm3 | 1.32567E-11 | 2.379826134 |

Supplementary Table 3 DEG_Categories_- 2 hrs for Suppl Fig 3A
Differentially expressed genes (FDR < 0.05)

| FRFvsAL Shared DEGs | | |
|---|---|---|
| Gene.ID | FDR | FC |
| G0s2 | 1.14187E-05 | 0.161673716 |
| Gdf15 | 3.33787E-07 | 0.074793001 |
| Gm3776 | 0.007206761 | 0.116052279 |
| Inmt | 0 | 0.369750292 |
| Lgals1 | 5.9601E-05 | 0.309137698 |
| Rorc | 0 | 0.273157778 |
| Serpina6 | 0.044602576 | 0.273915879 |
| Tspan4 | 2.947E-08 | 0.377292913 |
| Atf5 | 0.020140939 | 2.142552672 |
| Bhmt | 0.000322214 | 4.724734268 |
| Crot | 0.00066611 | 2.162418316 |
| Csrp3 | 1.49701E-08 | 3.425883798 |
| Cyp8b1 | 0.012949105 | 5.245016025 |
| Gm4952 | 3.04777E-11 | 3.515880027 |
| Gm5096 | 0.009266496 | 3.305931603 |
| Nr1d1 | 7.27281E-08 | 5.095043262 |
| Tymp | 9.47252E-07 | 3.825266411 |

| CRvsFRF Shared DEGs | | |
|---|---|---|
| Gene.ID | FDR | FC |
| Atf5 | 1.5963E-06 | 0.291149779 |
| Bhmt | 9.89374E-08 | 0.135314568 |
| Crot | 0.001549501 | 0.486039898 |
| Csrp3 | 7.42739E-14 | 0.277136972 |
| Cyp8b1 | 0.002579943 | 0.158093646 |
| Gm4952 | 1.07975E-11 | 0.417494159 |
| Gm5096 | 0.003850164 | 0.292065921 |
| Nr1d1 | 4.27175E-05 | 0.306545588 |
| Tymp | 8.71796E-08 | 0.250091102 |
| G0s2 | 0.004492047 | 5.716810318 |
| Gdf15 | 6.39962E-05 | 8.900194334 |
| Gm3776 | 0.000335354 | 13.59668385 |
| Inmt | 0 | 4.95181558 |
| Lgals1 | 0.015967246 | 2.444806667 |
| Rorc | 0 | 3.536768421 |
| Serpina6 | 0.000611041 | 3.964197584 |
| Tspan4 | 1.02218E-09 | 2.451312819 |

Supplementary Table 3 DEG_Categories_- 2 hrs for Suppl Fig 3A
Differentially expressed genes (FDR < 0.05)

| CRvsFRF Shared DEGs | | |
|---|---|---|
| Gene.ID | FDR | FC |
| Apon | 0 | 0.397703005 |
| Calr | 3.02914E-08 | 0.42199271 |
| Car14 | 3.08542E-08 | 0.237848688 |
| Derl3 | 1.88559E-08 | 0.081689433 |
| Hfe2 | 0.007310529 | 0.470473186 |
| Hsd3b7 | 0 | 0.259689916 |
| Leap2 | 8.66195E-09 | 0.331142365 |
| Mup11 | 2.14673E-05 | 0.02105072 |
| Mup14 | 0.002068948 | 0.041427304 |
| Mup15 | 3.78142E-13 | 0.001760648 |
| Mup17 | 1.80389E-12 | 0.002229172 |
| Mup2 | 0.000190223 | 0.054784341 |
| Mup9 | 0.001264855 | 0.063774894 |
| Nat8f2 | 0.014409676 | 0.460315202 |
| Pdia6 | 4.24503E-05 | 0.430478166 |
| Sdf2l1 | 5.58538E-05 | 0.194275795 |
| Selenbp2 | 0.031518937 | 0.12404344 |
| Acot2 | 0.002828264 | 2.6364132 |
| Ak4 | 0.000812053 | 2.440967119 |
| Ces1g | 1.53372E-06 | 2.485681061 |
| Clec2h | 0.018658605 | 4.613423998 |
| Cyp2a5 | 0 | 5.172062028 |
| Lgals4 | 1.39507E-10 | 3.859536469 |
| Sult2a3 | 3.38826E-05 | 26.82370479 |
| Zpr1 | 1.48629E-05 | 2.306866962 |

| CRvsAL Shared DEGs | | |
|---|---|---|
| Gene.ID | FDR | FC |
| Apon | 0.002740541 | 0.492686732 |
| Calr | 0.000201905 | 0.446967353 |
| Car14 | 3.38516E-05 | 0.335754164 |
| Derl3 | 0.005034792 | 0.122701987 |
| Hfe2 | 8.9493E-05 | 0.494058947 |
| Hsd3b7 | 5.47505E-08 | 0.47378237 |
| Leap2 | 0.007038164 | 0.307275336 |
| Mup11 | 0.001298062 | 0.029745716 |
| Mup14 | 0.022102387 | 0.051322963 |
| Mup15 | 1.22191E-12 | 0.002231482 |
| Mup17 | 1.04601E-11 | 0.002032802 |
| Mup2 | 0.000534978 | 0.045864712 |
| Mup9 | 0.009828442 | 0.06478358 |
| Nat8f2 | 1.73107E-07 | 0.439516265 |
| Pdia6 | 0.012903498 | 0.408921903 |
| Sdf2l1 | 0.01085324 | 0.118309897 |
| Selenbp2 | 2.21781E-05 | 0.033694678 |
| Acot2 | 7.58608E-05 | 3.353889184 |
| Ak4 | 0.002895199 | 2.010564998 |
| Ces1g | 0.001807636 | 2.384660636 |
| Clec2h | 0.007026118 | 5.477260673 |
| Cyp2a5 | 0 | 3.206588652 |
| Lgals4 | 1.38556E-13 | 4.895683461 |
| Sult2a3 | 0.002691416 | 13.27224087 |
| Zpr1 | 1.16134E-08 | 3.044291561 |
