## Supplementary Table 4 for "Anticipatory metabolic reprogramming distinguishes caloric restriction from fasting-refeeding cycles"

Supplementary Table 4 DEG_Categories_0 hrs for Suppl Fig 3A

Differentially expressed genes (FDR < 0.05)

| CRvsAL Exclusive DEGs | | |
|---|---|---|
| Gene.ID | FDR | FC |
| Adh4 | 0.01237448 | 0.461090901 |
| Ankrd23 | 0.042771781 | 0.172040241 |
| Cmah | 0.018100949 | 0.421916757 |
| Coq8a | 0.000434704 | 0.483923166 |
| Cyp2c70 | 0.000298803 | 0.420181191 |
| Cyp2d37-ps | 0.010444778 | 0.277780939 |
| Cyp2d40 | 0.018821577 | 0.337468818 |
| Dusp1 | 6.08857E-07 | 0.335968326 |
| Dusp6 | 0.000615213 | 0.455728085 |
| Enho | 0.001115397 | 0.352224973 |
| Fermt2 | 0.045200373 | 0.459823008 |
| Iigp1 | 2.67137E-07 | 0.480847863 |
| Insig2 | 1.3026E-05 | 0.411849273 |
| Lifr | 2.68829E-08 | 0.289684436 |
| Nfil3 | 8.59457E-07 | 0.328460611 |
| Nr0b2 | 9.85623E-06 | 0.154020993 |
| Rsrp1 | 1.53941E-07 | 0.477637657 |
| Serpina1e | 0.036956803 | 0.058236956 |
| Serpinb1a | 0.028611931 | 0.196471965 |
| Slc13a3 | 5.27862E-05 | 0.286886301 |
| Tat | 0.011552034 | 0.489267322 |
| Tlcd2 | 1.38172E-07 | 0.377155163 |
| Tob1 | 2.37943E-12 | 0.4994851 |
| Ubc | 0.005311395 | 0.442722099 |
| Wsb1 | 0.03278819 | 0.342170654 |
| Zfp36 | 0.001488971 | 0.393500735 |
| Acot2 | 0.034477305 | 2.301150785 |
| Cbr1 | 0.003572411 | 2.071010074 |
| Chd9 | 4.28103E-05 | 2.109095412 |
| Dnaja1 | 2.67418E-05 | 2.036406266 |
| Dnajb9 | 0.000104143 | 2.036886679 |
| Gm20594 | 4.21482E-06 | 2.705837029 |
| Gstt2 | 3.67656E-09 | 2.276539898 |
| Gys2 | 2.03614E-06 | 2.038772807 |
| Hsd17b13 | 9.89465E-07 | 2.052459447 |
| Hspa8 | 0 | 2.138298229 |
| Pcsk4 | 0.000545328 | 2.81761319 |
| Plin5 | 6.7797E-09 | 2.051872217 |
| Slc20a1 | 9.87016E-08 | 3.124658283 |

Supplementary Table 4 DEG_Categories_0 hrs for Suppl Fig 3A
Differentially expressed genes (FDR < 0.05)

| FRFvsAL Exclusive DEGs | | |
|---|---|---|
| Gene.ID | FDR | FC |
| A230050P20Rik | 0.04320964 | 0.484080747 |
| Acly | 3.45608E-10 | 0.39161691 |
| Acss2 | 1.63368E-05 | 0.460146142 |
| Cldn1 | 0.000687842 | 0.342677175 |
| Eef1a2 | 0.008976562 | 0.118672877 |
| Elovl6 | 0.010129783 | 0.48090313 |
| Erdr1 | 6.08662E-06 | 0.286725638 |
| Insc | 0.012995 | 0.450102448 |
| Nat8f1 | 0 | 0.460287573 |
| Pcsk9 | 0.000123435 | 0.429422551 |
| Phlda1 | 3.97615E-06 | 0.300815168 |
| Ppp1r3c | 0.004213439 | 0.307390858 |
| Scd1 | 1.59477E-05 | 0.202878678 |
| Serpina6 | 0.00354648 | 0.415991639 |
| Slc41a2 | 0.03764833 | 0.285546362 |
| Socs2 | 0.038649836 | 0.439315048 |
| Sucnr1 | 0.037900977 | 0.425146989 |
| Apoa4 | 5.40728E-08 | 6.503581468 |
| Atl2 | 0.03068531 | 2.01291318 |
| Bnip3 | 0 | 2.230101051 |
| Cdip1 | 0.000281615 | 2.058003964 |
| Ciart | 0.000429884 | 4.775031279 |
| Eif4ebp3 | 0.010318147 | 3.541403009 |
| Fbxo31 | 8.78771E-07 | 3.106871329 |
| Glrx | 3.54077E-06 | 2.009348789 |
| Nr1d2 | 0.007651306 | 2.341061857 |
| Nrg4 | 0.048733633 | 2.528310909 |
| Pex11a | 6.32677E-06 | 2.374665071 |
| Pnpla2 | 0 | 2.409569864 |
| Slc17a8 | 0.017904015 | 2.92222894 |
| Slc22a5 | 0.001407774 | 2.494418136 |
| Slc37a4 | 2.64694E-09 | 2.092243302 |
| Usp50 | 0.01157888 | 2.886606418 |

Supplementary Table 4 DEG_Categories_0 hrs for Suppl Fig 3A
Differentially expressed genes (FDR < 0.05)

| CRvsFRF Exclusive DEGs | | |
|---|---|---|
| Gene.ID | FDR | FC |
| Abcb4 | 2.22045E-16 | 0.45655783 |
| Abcg8 | 0.001301705 | 0.439056914 |
| Acss3 | 0.000264124 | 0.287629312 |
| Agtr1a | 2.45155E-06 | 0.421897659 |
| Angptl4 | 0.000219676 | 0.20096322 |
| Arl4d | 0.009861665 | 0.271117597 |
| Bhmt | 8.04173E-05 | 0.195986569 |
| Brap | 2.65121E-13 | 0.415076683 |
| Btg1 | 1.36528E-07 | 0.288435997 |
| Btg2 | 0.022610527 | 0.377894617 |
| Ccng2 | 0.002645446 | 0.143937473 |
| Cdkn1a | 0.003959723 | 0.109051312 |
| Cdo1 | 0.000299986 | 0.494919835 |
| Chic2 | 0.047140264 | 0.393000776 |
| Coq10b | 1.44591E-05 | 0.487701994 |
| Cpt1a | 0 | 0.478949981 |
| Cyp17a1 | 0.001044578 | 0.127889712 |
| Cyp8b1 | 0.0157767 | 0.237609654 |
| Dio1 | 0.000301696 | 0.304430191 |
| E030018B13 Rik | 0.019808775 | 0.162555154 |
| Ell2 | 0.002205114 | 0.420325328 |
| Erbb3 | 8.56537E-05 | 0.173640996 |
| Errfi1 | 2.09832E-14 | 0.446878495 |
| Etfbkmt | 2.67386E-05 | 0.453486308 |
| Fam35a | 0.000133651 | 0.331471593 |
| Fbxo21 | 0.003590203 | 0.389819375 |
| Glul | 1.65989E-05 | 0.489469151 |
| Gm4756 | 0.000683347 | 0.472193742 |
| Gm4952 | 0.007386337 | 0.434823767 |
| Gne | 1.20145E-10 | 0.482409648 |
| Gpcpd1 | 0.031440785 | 0.231363822 |
| Herpud1 | 1.55077E-07 | 0.427279242 |
| Irf2bp2 | 9.5133E-07 | 0.296087007 |
| Junb | 0.000353733 | 0.421101216 |
| Klf13 | 0.010009709 | 0.429605934 |
| Klf9 | 3.76331E-10 | 0.204701296 |
| Kynu | 0.037425634 | 0.47842401 |
| Lims2 | 4.64578E-11 | 0.424728845 |

Supplementary Table 4 DEG_Categories_0 hrs for Suppl Fig 3A
Differentially expressed genes (FDR < 0.05)

| CRvsFRF Exclusive DEGs | | |
|---|---|---|
| Gene.ID | FDR | FC |
| Lurap1l | 9.7204E-05 | 0.439379498 |
| Mafb | 2.74763E-05 | 0.375989292 |
| Mat1a | 0.003137002 | 0.438937138 |
| Mt1 | 0.024283398 | 0.182673502 |
| Mup10 | 0.000192482 | 0.329002067 |
| Myo1b | 0.002231096 | 0.438292647 |
| Npc1 | 0.002402304 | 0.336792276 |
| Oat | 0.001441529 | 0.477220197 |
| Pfkfb1 | 0.004452005 | 0.457238495 |
| Pim3 | 0.021446547 | 0.453119813 |
| Ppara | 2.64178E-05 | 0.467876938 |
| Prkd3 | 0.000867918 | 0.363051452 |
| Rapgef4 | 0 | 0.335971705 |
| Rarres1 | 3.28125E-10 | 0.197528535 |
| Rcl1 | 1.12133E-14 | 0.393047545 |
| Rhbg | 0.000237815 | 0.272110101 |
| RP24-346O11.2 | 0 | 0.114689745 |
| Serpina3n | 2.22045E-16 | 0.384497885 |
| Slc19a2 | 5.27646E-05 | 0.340072361 |
| Slc25a25 | 8.02677E-06 | 0.467842056 |
| Slc27a1 | 0.02958691 | 0.370104537 |
| Slc35g1 | 0.000156822 | 0.31342514 |
| Slc7a2 | 3.86802E-12 | 0.351783472 |
| Slc8b1 | 0.013596982 | 0.220548288 |
| Slco1b2 | 0 | 0.43456644 |
| Sult3a1 | 0.031003027 | 0.123207957 |
| Susd6 | 0.014236397 | 0.448400165 |
| Tcp11l2 | 4.62318E-08 | 0.17439882 |
| Them4 | 0.004305591 | 0.415490565 |
| Tle1 | 0.03719443 | 0.342412212 |
| Ttpa | 1.46901E-06 | 0.443840131 |
| Txnip | 0.041498601 | 0.33402953 |
| Tymp | 0.040812075 | 0.460618101 |
| Zfp36l2 | 0.025394941 | 0.490599525 |
| Abca8a | 0.041141575 | 2.139616268 |
| Actb | 4.10976E-11 | 2.644611649 |
| Aldh1b1 | 6.56328E-05 | 2.551259084 |
| Aldoa | 8.60327E-08 | 2.308433549 |

Supplementary Table 4 DEG_Categories_0 hrs for Suppl Fig 3A

Differentially expressed genes (FDR < 0.05)

| CRvsFRF Exclusive DEGs | | |
|---|---|---|
| Gene.ID | FDR | FC |
| Aldoc | 0.014303513 | 2.604575564 |
| Arhgdia | 0.000941102 | 2.15151428 |
| BC029214 | 0.007049417 | 2.046279207 |
| Bcl3 | 0.027019664 | 2.971869011 |
| Bdh2 | 0.008905605 | 3.494507241 |
| Ccnd1 | 0.004892457 | 4.994444443 |
| Cd63 | 4.60246E-05 | 2.757378993 |
| Cd74 | 5.39723E-10 | 2.586447755 |
| Ces2a | 6.08273E-05 | 2.094774175 |
| Creld2 | 0.035522085 | 3.935630294 |
| Cryl1 | 1.74044E-05 | 2.259844502 |
| Cyp26a1 | 9.93286E-07 | 3.558623392 |
| Ddit3 | 6.74655E-06 | 4.829158339 |
| Emilin1 | 0.039193798 | 2.513501419 |
| Fam84b | 0.045702793 | 3.204115883 |
| Foxa2 | 0.003458181 | 3.454823455 |
| G0s2 | 0.008736928 | 3.199973983 |
| Gclc | 0.003517098 | 2.493877107 |
| Gm8186 | 0.016682586 | 2.173030578 |
| Gsta4 | 2.92223E-10 | 3.183160472 |
| Gstm1 | 3.83138E-13 | 2.504045254 |
| Gstm3 | 0.004776158 | 4.376156232 |
| H2afx | 0.007596554 | 3.582806686 |
| Hamp2 | 0.037063867 | 6.171114884 |
| Hdhd3 | 1.33227E-15 | 3.908594585 |
| Hspa5 | 0.001571915 | 2.480788784 |
| Hyou1 | 5.93535E-05 | 2.437469525 |
| Manf | 1.11367E-07 | 3.512018555 |
| Mgst3 | 2.71913E-06 | 2.167190963 |
| Muc1 | 0.026449873 | 5.569668997 |
| Osgin1 | 1.25692E-05 | 2.924665968 |
| Psmb10 | 9.16372E-08 | 2.106932779 |
| S100a11 | 0.000711133 | 2.642075651 |
| S1pr1 | 2.18633E-11 | 3.27939325 |
| Samd1 | 7.11539E-06 | 2.743176688 |
| Sh3bp1 | 0.02263523 | 4.264403916 |
| Smim22 | 0.001788548 | 3.474481598 |
| Spp1 | 8.91742E-06 | 2.494608753 |
| Sult2a3 | 0.000557772 | 14.02615166 |

Supplementary Table 4 DEG_Categories_0 hrs for Suppl Fig 3A
Differentially expressed genes (FDR < 0.05)

| CRvsFRF Exclusive DEGs | | |
|---|---|---|
| Gene.ID | FDR | FC |
| Tceal8 | 0.03373943 | 2.310870253 |
| Tkt | 0.003619461 | 2.038424736 |
| Tmsb10 | 1.47346E-07 | 2.557813029 |
| Tstd1 | 2.27329E-08 | 3.308928422 |
| Tuba1b | 5.94603E-05 | 2.534427271 |
| Tuba4a | 0 | 2.282768298 |
| Tubb5 | 2.10231E-07 | 2.463282237 |
| Ugt1a5 | 0.047927154 | 2.050485635 |
| Xbp1 | 0.017830792 | 2.046681407 |
| Zfp966 | 0.032585459 | 2.051610946 |
| Zfp970 | 0.016611361 | 2.033533502 |

Supplementary Table 4 DEG_Categories_0 hrs for Suppl Fig 3A
Differentially expressed genes (FDR < 0.05)

| CRvsAL Shared DEGs | | |
|---|---|---|
| Gene.ID | FDR | FC |
| Lpin2 | 6.26376E-05 | 0.401890459 |
| Slc25a47 | 0.001952583 | 0.439483085 |
| Trp53inp1 | 5.52336E-13 | 0.139331842 |
| Aacs | 0.000452767 | 3.342323202 |
| Id3 | 0 | 3.278569906 |
| Mid1ip1 | 1.11144E-05 | 2.077426381 |

| FRFvsAL Shared DEGs | | |
|---|---|---|
| Gene.ID | FDR | FC |
| Aacs | 0.002910431 | 0.373139606 |
| Id3 | 5.17278E-08 | 0.458939866 |
| Mid1ip1 | 5.39116E-05 | 0.432447046 |
| Lpin2 | 0.000526804 | 2.610937835 |
| Slc25a47 | 9.07946E-11 | 2.323241524 |
| Trp53inp1 | 0.027192122 | 2.398118037 |

| CRvsFRF Shared DEGs | | |
|---|---|---|
| Gene.ID | FDR | FC |
| Lpin2 | 4.13657E-11 | 0.15392571 |
| Slc25a47 | 0 | 0.189168057 |
| Trp53inp1 | 2.12608E-13 | 0.058100494 |
| Aacs | 5.34082E-10 | 8.957299479 |
| Id3 | 0 | 7.143789733 |
| Mid1ip1 | 2.22045E-16 | 4.803886159 |

Supplementary Table 4 DEG_Categories_0 hrs for Suppl Fig 3A
Differentially expressed genes (FDR < 0.05)

| CRvsAL Shared DEGs | | |
|---|---|---|
| Gene.ID | FDR | FC |
| Acta1 | 2.44864E-08 | 0.007256136 |
| Atp2a1 | 1.04039E-05 | 0.028212725 |
| Car3 | 4.30688E-07 | 0.338517688 |
| Cfd | 0.004914755 | 0.08006415 |
| Ckm | 0 | 0.013682403 |
| Ckmt2 | 2.28954E-09 | 0.055856683 |
| Cox6a2 | 1.8698E-05 | 0.068596941 |
| Cox8b | 0.002417499 | 0.078429517 |
| Fabp3 | 0 | 0.040457835 |
| Hsd3b5 | 0.00235772 | 0.059605619 |
| Mb | 0 | 0.01356825 |
| Myh1 | 3.66281E-08 | 0.059237191 |
| Myl1 | 2.22045E-16 | 0.041860397 |
| Mylpf | 0.002509872 | 0.066444427 |
| Pgam2 | 0.035533101 | 0.148601927 |
| Selenbp2 | 1.70851E-05 | 0.019872448 |
| Tcap | 0 | 0.036770757 |
| Tnnc2 | 1.22876E-06 | 0.016207145 |
| Tnni2 | 3.66863E-06 | 0.020360904 |
| Tnnt3 | 1.17804E-06 | 0.015484092 |
| Rdh9 | 0.003505131 | 2.70567026 |

| FRFvsAL Shared DEGs | | |
|---|---|---|
| Gene.ID | FDR | FC |
| Acta1 | 0 | 0.006867145 |
| Atp2a1 | 1.89939E-05 | 0.028867525 |
| Car3 | 5.17364E-14 | 0.156971505 |
| Cfd | 0.003162003 | 0.068911359 |
| Ckm | 0 | 0.013682403 |
| Ckmt2 | 3.74096E-09 | 0.055856683 |
| Cox6a2 | 2.86633E-05 | 0.068596941 |
| Cox8b | 0.000995319 | 0.065095774 |
| Fabp3 | 0.000168567 | 0.044472627 |
| Hsd3b5 | 0.013154222 | 0.076749706 |
| Mb | 0 | 0.01356825 |
| Myh1 | 5.86896E-08 | 0.059237191 |
| Myl1 | 3.33067E-16 | 0.041860397 |
| Mylpf | 0.001511942 | 0.056367967 |
| Pgam2 | 0.01983695 | 0.125991931 |
| Selenbp2 | 0.003678836 | 0.048738303 |
| Tcap | 0 | 0.036770757 |
| Tnnc2 | 0 | 0.014559569 |
| Tnni2 | 8.47928E-06 | 0.021497598 |
| Tnnt3 | 0 | 0.013192669 |
| Rdh9 | 0.001977125 | 2.392779769 |

Supplementary Table 4 DEG_Categories_0 hrs for Suppl Fig 3A

Differentially expressed genes (FDR < 0.05)

| FRFvsAL Shared DEGs | | |
|---|---|---|
| Gene.ID | FDR | FC |
| Aqp8 | 1.15587E-05 | 0.297534914 |
| Cish | 0.003734302 | 0.290175216 |
| Fabp5 | 0.004103897 | 0.19758801 |
| Fasn | 0.025756757 | 0.434606652 |
| Gale | 0.000165319 | 0.214631941 |
| Gas6 | 0.001308832 | 0.326324529 |
| Inmt | 0 | 0.405371064 |
| Lgals1 | 4.71015E-07 | 0.337045259 |
| Nrep | 4.08661E-09 | 0.161919716 |
| Onecut1 | 0.008514289 | 0.225547558 |
| Sdf2l1 | 0.005683657 | 0.372789553 |
| Srebf1 | 0.000123551 | 0.23457352 |
| Sult2a5 | 0.026333657 | 0.148948282 |
| Tgm1 | 0.04220113 | 0.425187524 |
| Tgm2 | 0.000348061 | 0.464163687 |
| Tmem150a | 0 | 0.355542371 |
| Tmsb4x | 0.003193822 | 0.445737296 |
| Tubb2a | 0.01696962 | 0.402963559 |
| 1810055G02Rik | 0.000284751 | 2.666384094 |
| A930011G23Rik | 0.000197308 | 2.751061148 |
| Cyp4a31 | 0.000214054 | 3.391021261 |
| Fkbp5 | 0.004278252 | 3.563814111 |
| Gfra1 | 1.91147E-06 | 2.516651214 |
| Il1r1 | 0.039840353 | 2.24094055 |
| Il6ra | 5.31919E-05 | 2.700361055 |
| Lpin1 | 1.15052E-05 | 5.159372631 |
| Mfsd2a | 0.000128914 | 2.284728866 |
| Nlrp12 | 0.015598684 | 2.429283547 |
| Por | 9.6532E-08 | 2.000769063 |
| Rab30 | 2.57088E-06 | 3.251085674 |
| Rgs16 | 0 | 8.690539709 |
| St3gal5 | 0.00107829 | 3.385750918 |

| CRvsFRF Shared DEGs | | |
|---|---|---|
| Gene.ID | FDR | FC |
| 1810055G02Rik | 0 | 0.175551721 |
| A930011G23Rik | 4.05321E-08 | 0.256597251 |
| Cyp4a31 | 6.00154E-05 | 0.276380931 |
| Fkbp5 | 0.013679984 | 0.357886058 |
| Gfra1 | 0.000152663 | 0.449652648 |
| Il1r1 | 0.006111518 | 0.413617147 |
| Il6ra | 2.41585E-13 | 0.193235026 |
| Lpin1 | 5.07372E-14 | 0.069930646 |
| Mfsd2a | 2.34332E-06 | 0.438675418 |
| Nlrp12 | 0.001819243 | 0.371588651 |
| Por | 2.46646E-10 | 0.443282529 |
| Rab30 | 3.88383E-09 | 0.217583254 |
| Rgs16 | 0 | 0.074891615 |
| St3gal5 | 2.97819E-05 | 0.290612225 |
| Aqp8 | 0.000141868 | 2.565801132 |
| Cish | 4.79019E-05 | 3.321449811 |
| Fabp5 | 0.04353666 | 3.307345767 |
| Fasn | 4.1032E-06 | 2.861277462 |
| Gale | 0.000131288 | 4.359885366 |
| Gas6 | 1.53024E-06 | 3.073868349 |
| Inmt | 0.00567638 | 3.301228662 |
| Lgals1 | 0.000285485 | 2.276032714 |
| Nrep | 1.17317E-07 | 15.56585918 |
| Onecut1 | 5.22799E-05 | 8.710690777 |
| Sdf2l1 | 0.00977837 | 4.979809886 |
| Srebf1 | 6.55032E-15 | 4.154876108 |
| Sult2a5 | 8.00247E-05 | 15.91434554 |
| Tgm1 | 4.81606E-05 | 4.372353636 |
| Tgm2 | 0.000114357 | 2.816300664 |
| Tmem150a | 2.55351E-15 | 2.559421878 |
| Tmsb4x | 2.05562E-05 | 2.028689871 |
| Tubb2a | 0.007312315 | 3.051812877 |

Supplementary Table 4 DEG_Categories_0 hrs for Suppl Fig 3A
Differentially expressed genes (FDR < 0.05)

| CRvsFRF Shared DEGs | | |
|---|---|---|
| Gene.ID | FDR | FC |
| 8430408G22Rik | 1.91035E-09 | 0.044053463 |
| Arl4a | 0.00027706 | 0.257143113 |
| Cebpb | 1.11022E-16 | 0.410710701 |
| Csrp3 | 0 | 0.251092571 |
| D230025D16Rik | 9.12834E-06 | 0.359243774 |
| Fabp2 | 0.000271152 | 0.341746211 |
| G6pc | 0 | 0.155794419 |
| Gadd45g | 3.58713E-13 | 0.236318488 |
| Grb7 | 0.000117152 | 0.457042314 |
| Hsd3b7 | 1.13165E-12 | 0.369595814 |
| Igfbp1 | 1.86628E-13 | 0.002916058 |
| Ip6k2 | 3.66961E-05 | 0.149716662 |
| Irs2 | 5.12623E-08 | 0.132532971 |
| Klf15 | 1.02919E-08 | 0.375103624 |
| Mt2 | 0.00014274 | 0.027030451 |
| Mup1 | 1.74174E-05 | 0.016413464 |
| Mup11 | 5.16645E-08 | 0.009347168 |
| Mup12 | 0.000668536 | 0.037955233 |
| Mup13 | 4.81316E-10 | 0.12143437 |
| Mup14 | 0.000499174 | 0.032790058 |
| Mup15 | 0 | 0.001129717 |
| Mup16 | 0 | 0.125653248 |
| Mup17 | 2.3348E-11 | 0.00171951 |
| Mup18 | 9.4732E-12 | 0.227570548 |
| Mup19 | 3.29029E-08 | 0.033435123 |
| Mup2 | 2.43439E-09 | 0.022129977 |
| Mup7 | 2.75959E-05 | 0.013231616 |
| Mup8 | 0.032308502 | 0.200340153 |
| Mup9 | 1.85978E-11 | 0.028623918 |
| N4bp2l1 | 0 | 0.284058966 |
| Npr2 | 1.24148E-11 | 0.360195135 |
| Nr4a1 | 1.60506E-10 | 0.057214285 |
| Pck1 | 0 | 0.086379566 |
| Pnrc1 | 9.92218E-09 | 0.216199719 |
| Ppp1r3g | 5.75207E-13 | 0.036897406 |
| Rnf125 | 4.95604E-05 | 0.143247116 |
| Rps12 | 0 | 0.022350382 |
| Sdc4 | 9.19638E-08 | 0.238610706 |
| Sik1 | 0.001182431 | 0.365637379 |

| CRvsAL Shared DEGs | | |
|---|---|---|
| Gene.ID | FDR | FC |
| 8430408G22Rik | 3.05123E-06 | 0.040340571 |
| Arl4a | 2.25925E-08 | 0.398511896 |
| Cebpb | 7.55208E-08 | 0.470488046 |
| Csrp3 | 0.042370059 | 0.366904144 |
| D230025D16Rik | 0.006957866 | 0.425993857 |
| Fabp2 | 1.92669E-06 | 0.483468861 |
| G6pc | 0 | 0.167164878 |
| Gadd45g | 0.020377266 | 0.340078796 |
| Grb7 | 0.000179954 | 0.378632358 |
| Hsd3b7 | 0 | 0.300718801 |
| Igfbp1 | 0 | 0.007573699 |
| Ip6k2 | 2.25153E-09 | 0.265006254 |
| Irs2 | 7.12431E-05 | 0.225414689 |
| Klf15 | 7.15539E-13 | 0.481358715 |
| Mt2 | 0.013695941 | 0.095828548 |
| Mup1 | 1.22436E-06 | 0.007423756 |
| Mup11 | 3.91952E-07 | 0.005246925 |
| Mup12 | 6.96462E-05 | 0.018872516 |
| Mup13 | 0.00259238 | 0.084724358 |
| Mup14 | 3.88375E-05 | 0.01251661 |
| Mup15 | 5.50338E-13 | 0.000733746 |
| Mup16 | 0.047588828 | 0.105908063 |
| Mup17 | 4.34541E-13 | 0.000623802 |
| Mup18 | 0.027193105 | 0.208223373 |
| Mup19 | 0.010196181 | 0.047677926 |
| Mup2 | 5.06809E-06 | 0.016728742 |
| Mup7 | 2.00694E-06 | 0.006162909 |
| Mup8 | 0.002105597 | 0.081494532 |
| Mup9 | 0.000123829 | 0.020869797 |
| N4bp2l1 | 8.42071E-06 | 0.468320123 |
| Npr2 | 0.00783037 | 0.491079715 |
| Nr4a1 | 0.000919122 | 0.04897815 |
| Pck1 | 0 | 0.138185244 |
| Pnrc1 | 0 | 0.172104368 |
| Ppp1r3g | 0.007876555 | 0.101251669 |
| Rnf125 | 1.57365E-06 | 0.152143203 |
| Rps12 | 0 | 0.028799495 |
| Sdc4 | 0 | 0.340029948 |
| Sik1 | 0.031774913 | 0.400159858 |

Supplementary Table 4 DEG_Categories_0 hrs for Suppl Fig 3A
Differentially expressed genes (FDR < 0.05)

| CRvsFRF Shared DEGs | | |
|---|---|---|
| Gene.ID | FDR | FC |
| Slc22a7 | 6.01105E-06 | 0.178196828 |
| Slc38a2 | 0 | 0.103763977 |
| Slc45a3 | 0.019720023 | 0.361588621 |
| Sult5a1 | 0.002480793 | 0.253048986 |
| Ubb | 0 | 0.416821775 |
| Ugt3a1 | 0 | 0.286423787 |
| Upp2 | 4.7796E-10 | 0.266553383 |
| Actg1 | 0 | 5.549069909 |
| Alas1 | 1.22885E-05 | 2.815878279 |
| Angptl8 | 0 | 39.73254584 |
| Bag3 | 1.85323E-08 | 2.288063414 |
| Car2 | 8.11085E-05 | 3.48423136 |
| Cbr3 | 0.001805608 | 11.79023718 |
| Cib3 | 0.005152894 | 9.317465814 |
| Cyp2a5 | 1.56175E-07 | 2.287089565 |
| Dmbt1 | 5.14448E-05 | 24.19818641 |
| Dynll1 | 0 | 5.824545258 |
| Fmo3 | 5.175E-09 | 6.747261178 |
| Gm3776 | 0.000167431 | 18.22480173 |
| Gsta1 | 4.72084E-06 | 31.85552449 |
| Gsta2 | 0 | 6.450040284 |
| Hist1h1c | 0.006207007 | 6.896652014 |
| Hist2h3c2 | 0.000267789 | 5.87117729 |
| Hsp90aa1 | 0.001380132 | 2.48715371 |
| Hspa1b | 0 | 8.569321608 |
| Hsph1 | 0.024286201 | 2.252285865 |
| Id1 | 9.97312E-05 | 3.318554035 |
| Krt19 | 0.00984682 | 8.33667018 |
| Odc1 | 3.944E-11 | 2.114423685 |
| Orm2 | 3.28524E-05 | 21.38265405 |
| Pex26 | 0.006016022 | 2.403442156 |
| Rcan1 | 0 | 4.385766782 |
| Reg1 | 0.00886393 | 7.323454678 |
| Slc16a5 | 0.000756626 | 2.290415487 |
| Srsf3 | 1.44551E-13 | 2.311524116 |
| Tuba1c | 0 | 3.26803854 |
| Tuba8 | 5.89491E-06 | 4.926852307 |

| CRvsAL Shared DEGs | | |
|---|---|---|
| Gene.ID | FDR | FC |
| Slc22a7 | 1.42218E-05 | 0.09450653 |
| Slc38a2 | 2.22045E-16 | 0.20315097 |
| Slc45a3 | 0.012447284 | 0.494742269 |
| Sult5a1 | 5.19029E-13 | 0.164197271 |
| Ubb | 4.44711E-12 | 0.494859547 |
| Ugt3a1 | 9.3604E-12 | 0.298542815 |
| Upp2 | 0.000114339 | 0.442276366 |
| Actg1 | 0.033560745 | 2.186071295 |
| Alas1 | 0.001046508 | 2.267586114 |
| Angptl8 | 0 | 10.58996399 |
| Bag3 | 1.64547E-06 | 2.08238713 |
| Car2 | 0.001812335 | 2.769386772 |
| Cbr3 | 0.009124639 | 9.403250755 |
| Cib3 | 0.004709818 | 9.933294794 |
| Cyp2a5 | 0 | 3.246855412 |
| Dmbt1 | 0.000595555 | 17.88525009 |
| Dynll1 | 1.08802E-14 | 3.252589366 |
| Fmo3 | 1.27731E-06 | 4.590243481 |
| Gm3776 | 0.003386507 | 11.34435232 |
| Gsta1 | 0.000510476 | 15.68627133 |
| Gsta2 | 1.50908E-09 | 2.888788327 |
| Hist1h1c | 0.015655471 | 5.937234458 |
| Hist2h3c2 | 0.017062747 | 3.586665467 |
| Hsp90aa1 | 5.65029E-09 | 2.104596245 |
| Hspa1b | 0 | 9.886348307 |
| Hsph1 | 1.6731E-09 | 2.340782167 |
| Id1 | 2.20235E-08 | 4.387674302 |
| Krt19 | 0.033806765 | 6.920924002 |
| Odc1 | 1.75193E-12 | 2.075005369 |
| Orm2 | 0.001948369 | 11.8168031 |
| Pex26 | 0.020050929 | 2.340465098 |
| Rcan1 | 4.39537E-13 | 3.282153247 |
| Reg1 | 0.011039065 | 7.400120132 |
| Slc16a5 | 0.012974869 | 2.316671338 |
| Srsf3 | 1.62548E-12 | 2.700072189 |
| Tuba1c | 7.1676E-13 | 2.341880851 |
| Tuba8 | 5.6318E-06 | 5.723106872 |
