## Supplementary Table 5 for "Anticipatory metabolic reprogramming distinguishes caloric restriction from fasting-refeeding cycles"

Supplementary Table 5 DEG_Categories_2 hrs for Suppl Fig 3A
Differentially expressed genes (FDR < 0.05)

| CRvsAL Exclusive DEGs | | |
|---|---|---|
| Gene.ID | FDR | FC |
| Aacs | 0.016010707 | 0.467793921 |
| Adh4 | 1.7733E-05 | 0.45218296 |
| Bhlhe40 | 9.03276E-05 | 0.340807945 |
| Cyp2d40 | 3.02975E-09 | 0.422997305 |
| Enho | 7.34977E-07 | 0.323260818 |
| Fst | 0.032011055 | 0.250324514 |
| G6pc | 0.000411316 | 0.38011479 |
| H2afv | 0.007644236 | 0.475082585 |
| Me1 | 0.000222346 | 0.471607381 |
| Mup18 | 0.003979065 | 0.277806932 |
| Nat8f1 | 1.01215E-09 | 0.42846977 |
| Sik1 | 1.01076E-05 | 0.242083754 |
| Slc25a30 | 0.004147272 | 0.226141642 |
| Zfp36 | 6.65707E-06 | 0.294504498 |
| Ctrb1 | 0.002576387 | 12.01349936 |
| Cyp2a5 | 1.52705E-10 | 2.622233108 |
| Dnaja1 | 0 | 2.080629905 |
| Hspa1b | 0.005808955 | 4.053752062 |
| Hspa8 | 0 | 3.101878131 |
| Orm3 | 0.027377452 | 4.342514262 |
| Osgin1 | 0.00139879 | 2.477173678 |
| Rdh9 | 5.78094E-06 | 2.685937558 |
| Reg1 | 0.034893131 | 6.272926162 |

Supplementary Table 5 DEG_Categories_2 hrs for Suppl Fig 3A
Differentially expressed genes (FDR < 0.05)

| FRFvsAL Exclusive DEGs | | |
|---|---|---|
| Gene.ID | FDR | FC |
| Cfd | 0.008033306 | 0.222643448 |
| Cyp4f14 | 5.74765E-08 | 0.493867055 |
| Gale | 0.002128776 | 0.393175238 |
| Sucnr1 | 0.000126008 | 0.374970035 |
| Tgm1 | 0.030262917 | 0.320795861 |
| Glrx | 1.38518E-06 | 2.020358208 |
| Plin5 | 0.000450824 | 2.123344014 |
| Rbm3 | 2.05817E-06 | 2.973683774 |
| Rps12 | 0.000337303 | 16.54341335 |
| Slc22a5 | 0.001249365 | 2.082988253 |
| Slc37a4 | 4.65183E-13 | 2.062220806 |

Supplementary Table 5 DEG_Categories_2 hrs for Suppl Fig 3A
Differentially expressed genes (FDR < 0.05)

| CRvsFRF Exclusive DEGs | | |
|---|---|---|
| Gene.ID | FDR | FC |
| 1810055G02Rik | 0.000499274 | 0.196383017 |
| 8430408G22Rik | 2.84832E-06 | 0.106196538 |
| A930011G23Rik | 0.025220725 | 0.308380979 |
| Aatk | 0.008549194 | 0.225411955 |
| Abcb4 | 3.33067E-16 | 0.435934276 |
| Acot12 | 0 | 0.371370114 |
| Apoa1 | 0 | 0.464907116 |
| Apoa5 | 6.89593E-09 | 0.497745504 |
| Arl4a | 0.003728273 | 0.288511684 |
| Arl4d | 0.00618493 | 0.288549144 |
| Brap | 3.44169E-15 | 0.410528752 |
| Bri3 | 7.77156E-16 | 0.474301912 |
| Ccng2 | 0.000252464 | 0.268604886 |
| Cdkn1a | 0.039624415 | 0.235391991 |
| Cpt1a | 4.70068E-13 | 0.391959052 |
| Csad | 0.014998266 | 0.311188831 |
| Ddit4 | 0.028520828 | 0.185703393 |
| Dirc2 | 0.002565351 | 0.313388066 |
| Eif4b | 6.95698E-05 | 0.492726879 |
| Errfi1 | 1.55431E-15 | 0.390167974 |
| Etfbkmt | 5.40884E-06 | 0.353426344 |
| Fam134b | 0.001361009 | 0.374580123 |
| Gch1 | 0.000182872 | 0.413236569 |
| Gfra1 | 9.32871E-06 | 0.490150897 |
| Gne | 0 | 0.319291761 |
| Hmga1 | 0.030885956 | 0.459814882 |
| Hmgcs2 | 9.80327E-13 | 0.46176778 |
| Lipg | 0.031772856 | 0.355481729 |
| Lpin1 | 5.27813E-10 | 0.050168607 |
| Lrg1 | 0.009284108 | 0.48113617 |
| Mfsd2a | 1.9115E-11 | 0.139971606 |
| Mup1 | 7.07943E-05 | 0.045612841 |
| Mup12 | 0.032424668 | 0.067141284 |
| Mup19 | 0.014604039 | 0.056053157 |
| Mup6 | 0.002151887 | 0.201778312 |
| Mup7 | 0.007087638 | 0.032421405 |
| Mup8 | 0.004857935 | 0.149413672 |
| Nfkbia | 0.000117013 | 0.459254155 |
| Obp2a | 0.010358937 | 0.160618792 |

Supplementary Table 5 DEG_Categories_2 hrs for Suppl Fig 3A
Differentially expressed genes (FDR < 0.05)

| CRvsFRF Exclusive DEGs | | |
|---|---|---|
| Gene.ID | FDR | FC |
| Pank1 | 1.31006E-14 | 0.435565806 |
| Ppp1r15a | 0.015433239 | 0.301867569 |
| Rab30 | 0.000557825 | 0.221021388 |
| Rcl1 | 1.44329E-15 | 0.440056899 |
| Retsat | 0.02913294 | 0.467494893 |
| Rgs16 | 0.000194861 | 0.077683618 |
| Sdc4 | 0 | 0.452260985 |
| Sds | 5.20362E-13 | 0.473908347 |
| Serpina3n | 5.32907E-15 | 0.42281462 |
| Slc17a8 | 0.04786658 | 0.285908556 |
| Slc25a33 | 0.000204257 | 0.350001963 |
| Slc27a1 | 0.048055933 | 0.408972136 |
| Slc27a2 | 0 | 0.468816308 |
| Slc38a2 | 0 | 0.274745072 |
| Slc45a3 | 0.014111381 | 0.409608032 |
| Slc8b1 | 2.29932E-09 | 0.249984876 |
| Slco1b2 | 2.55351E-15 | 0.499520253 |
| Socs2 | 4.61955E-06 | 0.230667446 |
| Sun2 | 0.00103029 | 0.341897926 |
| Tat | 0.02945967 | 0.408557282 |
| Tdo2 | 4.34025E-08 | 0.410983279 |
| Tmem120a | 2.83515E-11 | 0.489902308 |
| Tmem56 | 3.87634E-09 | 0.42460226 |
| Tnfaip8l1 | 0.003946113 | 0.46007701 |
| Trp53inp1 | 2.76072E-06 | 0.156374404 |
| Ttpa | 0.000106431 | 0.425671375 |
| Txnip | 0.007223789 | 0.330854974 |
| Ubb | 0 | 0.467652735 |
| Vnn1 | 5.15586E-06 | 0.386986427 |
| Ypel3 | 2.66169E-07 | 0.441601686 |
| Acmsd | 0.001793731 | 2.904771555 |
| Actb | 0 | 2.325424647 |
| Aldh1b1 | 8.00539E-05 | 2.888483657 |
| Apol7a | 5.09393E-12 | 2.316520362 |
| Arhgef19 | 7.22324E-06 | 2.08661387 |
| Ccnd1 | 0.001748457 | 5.103480857 |
| Cyp2c55 | 0.006554253 | 4.418450682 |
| Cyp7a1 | 0.04839633 | 2.565170492 |
| Ddc | 0.001324354 | 2.074197079 |

Supplementary Table 5 DEG_Categories_2 hrs for Suppl Fig 3A
Differentially expressed genes (FDR < 0.05)

| CRvsFRF Exclusive DEGs | | |
|---|---|---|
| Gene.ID | FDR | FC |
| Elovl1 | 0.000285257 | 2.414793479 |
| Elovl6 | 4.31449E-05 | 4.207834376 |
| Ethe1 | 5.54545E-07 | 3.094081368 |
| Fam47e | 0.007668404 | 2.108100891 |
| Fmo3 | 1.60405E-05 | 22.20850307 |
| Gsta1 | 8.66778E-06 | 9.950094111 |
| Gsta2 | 3.10862E-14 | 3.688472253 |
| Gsta4 | 8.97823E-09 | 2.060515066 |
| Gstm3 | 5.55406E-05 | 7.59888796 |
| Hamp2 | 0.0188916 | 8.911853698 |
| Hdhd3 | 0 | 4.135905846 |
| Hspb1 | 0.001862557 | 2.374957246 |
| Igtp | 0.008245197 | 2.107482966 |
| Inmt | 6.49075E-07 | 2.276414775 |
| Krt19 | 0.021176632 | 7.73676182 |
| Rcan1 | 0 | 3.090173841 |
| Sdc1 | 0 | 2.38595305 |
| Sdf2l1 | 2.24427E-07 | 3.696452033 |
| Sec23b | 0.006514951 | 2.563171573 |
| Sept9 | 0.016687542 | 2.573135974 |
| Spp1 | 8.28135E-05 | 2.926489311 |
| Sult2a3 | 0.000795077 | 12.69088798 |
| Sult2a7 | 3.67317E-07 | 5.264543894 |
| Syvn1 | 4.34365E-09 | 2.153579049 |
| Tmem150a | 0 | 2.303522573 |
| Tuba4a | 0 | 2.4156845 |
| Tubb2a | 5.67751E-08 | 2.430053994 |
| Ugt1a5 | 0 | 3.458162537 |

Supplementary Table 5 DEG_Categories_2 hrs for Suppl Fig 3A

Differentially expressed genes (FDR < 0.05)

| CRvsAL Shared DEGs | | |
|---|---|---|
| Gene.ID | FDR | FC |
| Angptl4 | 4.60511E-05 | 0.338232668 |
| Angptl8 | 3.71871E-07 | 2.580793463 |

| FRFvsAL Shared DEGs | | |
|---|---|---|
| Gene.ID | FDR | FC |
| Angptl8 | 0.007793701 | 0.242144397 |
| Angptl4 | 3.78292E-05 | 2.498315741 |

| CRvsFRF Shared DEGs | | |
|---|---|---|
| Gene.ID | FDR | FC |
| Angptl4 | 0 | 0.135384276 |
| Angptl8 | 0 | 10.65807633 |

Supplementary Table 5 DEG_Categories_2 hrs for Suppl Fig 3A
Differentially expressed genes (FDR < 0.05)

| CRvsAL Shared DEGs | | |
|---|---|---|
| Gene.ID | FDR | FC |
| Kyat1 | 0.000129037 | 0.429994925 |
| Lifr | 0 | 0.349490515 |
| Mapk15 | 0.00032538 | 0.245972896 |
| Chd9 | 0 | 2.324130086 |

| FRFvsAL Shared DEGs | | |
|---|---|---|
| Gene.ID | FDR | FC |
| Kyat1 | 3.9883E-06 | 0.372482764 |
| Lifr | 4.32263E-06 | 0.452824698 |
| Mapk15 | 0.001068763 | 0.261414675 |
| Chd9 | 7.65208E-06 | 2.689287838 |

Supplementary Table 5 DEG_Categories_2 hrs for Suppl Fig 3A
Differentially expressed genes (FDR < 0.05)

| FRFvsAL Shared DEGs | | |
|---|---|---|
| Gene.ID | FDR | FC |
| A1bg | 5.40926E-05 | 0.034050842 |
| Acly | 9.49416E-05 | 0.329160817 |
| Aqp8 | 0 | 0.235589929 |
| Car3 | 0.015420926 | 0.163170912 |
| Fasn | 0.000395858 | 0.327449446 |
| Gm3776 | 0.04308721 | 0.167438656 |
| Lasp1 | 0.000227984 | 0.464973234 |
| Mid1ip1 | 3.75743E-09 | 0.350604799 |
| Nrep | 1.10638E-05 | 0.210728658 |
| Ppp1r3c | 5.02535E-07 | 0.303599042 |
| Scd1 | 2.41168E-05 | 0.095701958 |
| Srebf1 | 2.24869E-10 | 0.187289959 |
| Tgm2 | 2.23828E-07 | 0.453016328 |
| Apoa4 | 0 | 4.250373766 |
| Bnip3 | 0 | 2.387768765 |
| Cdip1 | 0.003771963 | 2.217010458 |
| Ctgf | 1.33242E-05 | 4.126876517 |
| Cyp4a31 | 0.004924767 | 2.483616988 |
| Fbxo31 | 1.12857E-06 | 3.055681435 |
| Fkbp5 | 5.02585E-11 | 3.780775994 |
| Herpud1 | 0.014700044 | 2.172867678 |
| Il1r1 | 0.003044059 | 2.924066144 |
| Ip6k2 | 2.06604E-08 | 2.198831173 |
| Lpin2 | 1.68965E-12 | 2.261598204 |
| Mt2 | 0.005907744 | 10.91725426 |
| Nlrp12 | 0.00448268 | 3.107936192 |
| Pex11a | 4.79776E-06 | 2.298949486 |
| Pnpla2 | 2.72005E-14 | 2.320480811 |
| Por | 0 | 2.172033749 |
| St3gal5 | 0.000810565 | 3.009449798 |
| Sult1d1 | 0.004994719 | 2.369943348 |

| CRvsFRF Shared DEGs | | |
|---|---|---|
| Gene.ID | FDR | FC |
| Apoa4 | 0 | 0.174539417 |
| Bnip3 | 0 | 0.449058258 |
| Cdip1 | 2.4213E-05 | 0.397361831 |
| Ctgf | 3.35133E-09 | 0.15056536 |
| Cyp4a31 | 0.001098764 | 0.354206802 |
| Fbxo31 | 8.39329E-14 | 0.248590097 |
| Fkbp5 | 0 | 0.234542181 |
| Herpud1 | 4.07445E-07 | 0.332090513 |
| Il1r1 | 7.14971E-05 | 0.27489697 |
| Ip6k2 | 0 | 0.247069311 |
| Lpin2 | 0 | 0.185846894 |
| Mt2 | 2.90539E-05 | 0.048149434 |
| Nlrp12 | 1.8619E-05 | 0.211358487 |
| Pex11a | 4.00347E-06 | 0.450406461 |
| Pnpla2 | 3.21965E-13 | 0.431909486 |
| Por | 0 | 0.315862736 |
| St3gal5 | 2.9259E-09 | 0.295218896 |
| Sult1d1 | 0.000928039 | 0.438033987 |
| A1bg | 0.000120666 | 18.85117535 |
| Acly | 4.21885E-15 | 2.7474738 |
| Aqp8 | 0 | 4.352202323 |
| Car3 | 2.413E-05 | 3.452442493 |
| Fasn | 0 | 3.302224811 |
| Gm3776 | 4.78881E-07 | 12.8506504 |
| Lasp1 | 5.032E-08 | 2.756631373 |
| Mid1ip1 | 7.5004E-10 | 2.68639683 |
| Nrep | 0 | 8.738949911 |
| Ppp1r3c | 0 | 4.596258874 |
| Scd1 | 5.41954E-05 | 5.338416554 |
| Srebf1 | 0 | 5.342800878 |
| Tgm2 | 0 | 3.474156722 |

Supplementary Table 5 DEG_Categories_2 hrs for Suppl Fig 3A
Differentially expressed genes (FDR < 0.05)

| CRvsFRF Shared DEGs | | |
|---|---|---|
| Gene.ID | FDR | FC |
| Abcg8 | 5.72564E-12 | 0.277865955 |
| Cebpb | 4.45901E-10 | 0.237383389 |
| Csrp3 | 0.000302195 | 0.211211469 |
| Dio1 | 0 | 0.18156544 |
| Dusp1 | 0.001014836 | 0.287127902 |
| Fabp2 | 0 | 0.313962103 |
| Fgf21 | 5.86198E-14 | 0.062794039 |
| Gadd45g | 0.003943019 | 0.068690618 |
| Igfbp1 | 3.49889E-11 | 0.009593308 |
| Jun | 7.48568E-12 | 0.232594686 |
| Mafb | 0.000143899 | 0.226127749 |
| Mup11 | 4.95964E-06 | 0.006626283 |
| Mup13 | 0.010976883 | 0.142394177 |
| Mup14 | 0.005029085 | 0.034449358 |
| Mup15 | 2.0639E-13 | 0.001347726 |
| Mup17 | 7.76379E-13 | 0.001307715 |
| Mup2 | 0.002000398 | 0.035963305 |
| Mup9 | 0.001308852 | 0.041524029 |
| N4bp2l1 | 2.70894E-13 | 0.320513886 |
| Npr2 | 0.000431121 | 0.481629121 |
| Nr4a1 | 0.000922377 | 0.111192962 |
| Pck1 | 0 | 0.154822303 |
| Pim3 | 0.002370913 | 0.251692487 |
| Plk3 | 0 | 0.192816854 |
| Ppp1r3g | 1.71463E-05 | 0.029494439 |
| Rapgef4 | 1.26837E-06 | 0.375638273 |
| Saa3 | 0.009771881 | 0.318801377 |
| Selenbp2 | 0.001318814 | 0.051923478 |
| Slc25a25 | 0.000755307 | 0.195068924 |
| Slc25a47 | 0 | 0.194655791 |
| Trp53inp2 | 1.05986E-05 | 0.390772332 |
| Ugt3a1 | 1.26687E-07 | 0.369177622 |
| Upp2 | 5.25135E-14 | 0.255627504 |
| Actg1 | 0 | 4.326207952 |
| Creld2 | 4.21539E-05 | 2.950410303 |
| Dmbt1 | 1.37022E-05 | 34.71318445 |
| Dynll1 | 0.003896042 | 2.399449671 |
| Hist1h1c | 2.43317E-07 | 4.078788884 |
| Hsp90aa1 | 0 | 2.417966404 |

| CRvsAL Shared DEGs | | |
|---|---|---|
| Gene.ID | FDR | FC |
| Abcg8 | 0.000151696 | 0.379203124 |
| Cebpb | 7.42358E-09 | 0.301348138 |
| Csrp3 | 2.29814E-05 | 0.340917195 |
| Dio1 | 1.20841E-09 | 0.228987847 |
| Dusp1 | 1.80178E-06 | 0.145199561 |
| Fabp2 | 5.38392E-09 | 0.402675252 |
| Fgf21 | 0.000401165 | 0.140510577 |
| Gadd45g | 0.012983538 | 0.12165908 |
| Igfbp1 | 6.96733E-08 | 0.022682192 |
| Jun | 0.000300688 | 0.297784824 |
| Mafb | 0.025946567 | 0.203228431 |
| Mup11 | 3.05759E-05 | 0.016711148 |
| Mup13 | 0.009847275 | 0.21465154 |
| Mup14 | 0.028651686 | 0.060429089 |
| Mup15 | 9.14824E-14 | 0.002225951 |
| Mup17 | 1.12668E-11 | 0.001735605 |
| Mup2 | 0.000378178 | 0.040725754 |
| Mup9 | 0.006953288 | 0.066797175 |
| N4bp2l1 | 7.2127E-07 | 0.4514704 |
| Npr2 | 4.12569E-05 | 0.456149985 |
| Nr4a1 | 3.56871E-10 | 0.041654975 |
| Pck1 | 3.23161E-06 | 0.202414221 |
| Pim3 | 1.94798E-08 | 0.380912043 |
| Plk3 | 0.006574765 | 0.262372998 |
| Ppp1r3g | 3.94703E-05 | 0.044915916 |
| Rapgef4 | 0.007279268 | 0.474831888 |
| Saa3 | 9.3804E-06 | 0.362839756 |
| Selenbp2 | 0.000280968 | 0.055305902 |
| Slc25a25 | 0 | 0.198240683 |
| Slc25a47 | 1.21601E-09 | 0.377341042 |
| Trp53inp2 | 3.39111E-08 | 0.472120832 |
| Ugt3a1 | 4.88498E-15 | 0.373777975 |
| Upp2 | 6.788E-09 | 0.348939142 |
| Actg1 | 3.03346E-12 | 2.635056366 |
| Creld2 | 6.80414E-11 | 3.592845533 |
| Dmbt1 | 1.30672E-05 | 38.06518996 |
| Dynll1 | 0.00025385 | 2.944527272 |
| Hist1h1c | 0.048220261 | 2.122261407 |
| Hsp90aa1 | 0 | 2.266663401 |

Supplementary Table 5 DEG_Categories_2 hrs for Suppl Fig 3A
Differentially expressed genes (FDR < 0.05)

| CRvsFRF Shared DEGs | | |
|---|---|---|
| Gene.ID | FDR | FC |
| Hspa5 | 7.14541E-11 | 2.243320245 |
| Hyou1 | 0 | 2.835069992 |
| Manf | 1.54022E-08 | 2.759873721 |
| Odc1 | 2.56483E-05 | 2.021169022 |
| Orm2 | 2.40698E-06 | 11.32922775 |
| Slc16a5 | 8.68456E-06 | 2.563330877 |
| Srm | 1.1472E-10 | 2.386076896 |
| Srsf3 | 0.003003785 | 2.084002711 |
| Tuba1c | 0 | 4.048563534 |
| Tuba8 | 4.11694E-08 | 5.110818333 |

| CRvsAL Shared DEGs | | |
|---|---|---|
| Gene.ID | FDR | FC |
| Hspa5 | 8.24928E-09 | 2.193924103 |
| Hyou1 | 1.1096E-07 | 2.151470051 |
| Manf | 0 | 2.500208345 |
| Odc1 | 2.71813E-08 | 2.456988669 |
| Orm2 | 2.71009E-05 | 9.142695889 |
| Slc16a5 | 9.05349E-07 | 2.963996294 |
| Srm | 1.56795E-06 | 2.451680998 |
| Srsf3 | 4.8482E-09 | 2.110837084 |
| Tuba1c | 0 | 2.488425447 |
| Tuba8 | 1.08198E-10 | 6.39164523 |
