## Supplementary Table 6 for "Anticipatory metabolic reprogramming distinguishes caloric restriction from fasting-refeeding cycles"

Supplementary Table 6 DEG_Categories_6 hrs for Suppl Fig 3B
Differentially expressed genes (FDR < 0.05)

| CRvsAL Exclusive DEGs | | |
|---|---|---|
| Gene.ID | FDR | FC |
| Bhlhe40 | 5.60552E-07 | 0.289437092 |
| Cyp2c70 | 3.90664E-06 | 0.473625723 |
| G6pc | 0.019648975 | 0.298739188 |
| Gadd45g | 0.018152564 | 0.060441334 |
| Jund | 2.19935E-13 | 0.464303963 |
| Mup8 | 0.004282262 | 0.101180595 |
| Noct | 0.00021463 | 0.201890151 |
| Nr4a1 | 0.000304424 | 0.066503 |
| Slco1a1 | 0.000119387 | 0.040697865 |
| Tob1 | 0.000740928 | 0.486635366 |
| Trib1 | 0.031698799 | 0.443314547 |
| Zfp36 | 5.19545E-05 | 0.369015564 |
| Cyp2a5 | 0.001575528 | 2.097328972 |
| Cyp3a59 | 0.027960209 | 2.082377835 |
| Fmo3 | 0.031007146 | 5.07510997 |
| Rdh9 | 2.10477E-08 | 3.005918442 |
| Rps12 | 0.002051645 | 15.09595226 |

Supplementary Table 6 DEG_Categories_6 hrs for Suppl Fig 3B
Differentially expressed genes (FDR < 0.05)

| FRFvsAL Exclusive DEGs | | |
|---|---|---|
| Gene.ID | FDR | FC |
| Agxt | 0 | 0.477588998 |
| Asns | 0.006112002 | 0.098598233 |
| Atp4a | 0.042720772 | 0.188931357 |
| C1qa | 0.029054085 | 0.452012045 |
| C6 | 0.003223899 | 0.082159671 |
| Cd52 | 0.036004342 | 0.368471978 |
| Ctss | 0.043856134 | 0.467202062 |
| Cxcl9 | 0.025991933 | 0.288551756 |
| Cyp26a1 | 0.000699379 | 0.400833522 |
| Cyp2u1 | 0.03158137 | 0.170732074 |
| Cyp3a16 | 7.75719E-05 | 0.035437052 |
| Cyp3a41b | 9.31295E-06 | 0.027567657 |
| Efna1 | 0.027323103 | 0.47598708 |
| Fabp5 | 0.001236178 | 0.184725938 |
| Fam25c | 4.52408E-11 | 0.303355525 |
| Gstp2 | 0.018170313 | 0.115373169 |
| Gstp3 | 0.032881343 | 0.480493505 |
| H2-Eb1 | 1.36304E-06 | 0.254726195 |
| Id2 | 2.22045E-16 | 0.448741922 |
| Ifi47 | 0.036655392 | 0.436084096 |
| Mmd2 | 0.002441902 | 0.264950138 |
| Mup21 | 0.001477322 | 0.056592944 |
| Nat8 | 0.000483489 | 0.067241965 |
| Nat8f1 | 1.01096E-05 | 0.404571947 |
| Nat8f5 | 0.017648065 | 0.132875859 |
| Pde4b | 0.011685664 | 0.314192181 |
| Psmb10 | 6.21774E-07 | 0.49270114 |
| Saa3 | 0.03401964 | 0.265436008 |
| Smim22 | 0.01046358 | 0.304651622 |
| Spsb4 | 0.03659851 | 0.268180438 |
| Tmsb4x | 0.002070877 | 0.46226053 |
| Cyp2b13 | 0.011849939 | 2.995564103 |
| Cyp2c39 | 0.000125015 | 3.533812825 |
| Cyp4a31 | 3.03261E-05 | 2.763961296 |
| Cyp51 | 0.025778815 | 2.963905537 |
| Dhcr24 | 2.22045E-16 | 2.593836321 |
| Fam35a | 0.000442536 | 2.578356099 |
| Glrx | 1.44873E-07 | 2.377692857 |
| Hmgcr | 0.016173955 | 4.045927812 |

Supplementary Table 6 DEG_Categories_6 hrs for Suppl Fig 3B
Differentially expressed genes (FDR < 0.05)

| FRFvsAL Exclusive DEGs | | |
|---|---|---|
| Gene.ID | FDR | FC |
| Hsd17b10 | 2.58957E-06 | 2.145205544 |
| Hsp90ab1 | 0 | 2.234416188 |
| Hspb1 | 1.90753E-08 | 2.962443755 |
| Il6ra | 0.000275828 | 3.68597978 |
| Ip6k2 | 0.02881573 | 2.616244298 |
| Lpin1 | 0.01276511 | 7.8661214 |
| Mgll | 3.18328E-06 | 2.247241416 |
| Retsat | 0.000553309 | 3.167657087 |
| Serpina3m | 9.98756E-09 | 2.091660557 |
| Slc16a7 | 0.020581613 | 2.022087336 |
| Slc35g1 | 0.037677312 | 2.347902953 |
| Sult1e1 | 0.029471765 | 4.945955727 |
| Tango2 | 0.024857603 | 2.248442633 |

Supplementary Table 6 DEG_Categories_6 hrs for Suppl Fig 3B
Differentially expressed genes (FDR < 0.05)

| CRvsFRF Exclusive DEGs | | |
|---|---|---|
| Gene.ID | FDR | FC |
| 1810055G02Rik | 0.003770426 | 0.262842679 |
| Atp2a2 | 0.007883122 | 0.476751577 |
| Bhmt | 0.047081127 | 0.402804512 |
| Cdkn1a | 0.000107074 | 0.145718288 |
| Ddit3 | 0.027816137 | 0.385313507 |
| Dio1 | 0 | 0.219081919 |
| Dnaja1 | 0 | 0.26598635 |
| Dnajb1 | 2.31149E-07 | 0.300167738 |
| Ero1lb | 0.045495787 | 0.401183825 |
| Fabp2 | 1.77636E-15 | 0.476500136 |
| Fgf21 | 0.01452384 | 0.139067833 |
| Gck | 0.038658509 | 0.361928122 |
| Lcn2 | 0.002404503 | 0.334515086 |
| Mup3 | 0.018308611 | 0.480229871 |
| Pim3 | 0.012559693 | 0.348664433 |
| Plk3 | 0.000144157 | 0.308012272 |
| Serpina3k | 4.80024E-09 | 0.4406007 |
| Sult3a1 | 0 | 0.168865403 |
| Tsc22d3 | 4.52079E-05 | 0.386095521 |
| Vnn1 | 6.42925E-06 | 0.384395558 |
| BC048546 | 0.006356568 | 2.337836622 |
| Cib3 | 0.015429456 | 7.657375049 |
| Ckm | 0.020717776 | 7.045924261 |
| Cox7a1 | 0.047838547 | 3.568170294 |
| Cyp2c55 | 0.031912499 | 6.527568138 |
| Ethe1 | 0.048747735 | 2.020246375 |
| Gm3776 | 0.000330488 | 8.147739869 |
| Gsta1 | 4.11847E-05 | 12.1797829 |
| Gsta2 | 2.1497E-06 | 5.275078012 |
| Gsta4 | 7.91762E-11 | 2.541726123 |
| Gstm1 | 5.84799E-06 | 2.188185355 |
| Gstm3 | 2.11285E-05 | 6.457532011 |
| Gstm6 | 3.99352E-06 | 2.587410615 |
| Inmt | 9.88829E-06 | 2.218736034 |
| Lgals1 | 9.19356E-06 | 3.049846647 |
| Mgst3 | 0.001956256 | 2.207924383 |
| Raet1d | 0.044969357 | 6.244706548 |
| Rnf125 | 0.001104413 | 2.014377294 |
| Spp1 | 0.017881791 | 2.705056179 |

Supplementary Table 6 DEG_Categories_6 hrs for Suppl Fig 3B
Differentially expressed genes (FDR < 0.05)

| CRvsFRF Exclusive DEGs | | |
|---|---|---|
| Gene.ID | FDR | FC |
| Srsf5 | 0.000514215 | 2.199404267 |
| Sucnr1 | 8.20638E-05 | 2.65470472 |
| Sult2a3 | 0.001756517 | 16.35464622 |
| Sult2a5 | 0.011059256 | 10.12406809 |
| Tnnc2 | 0.006109364 | 9.586116142 |
| Tstd1 | 0.001811774 | 2.652899071 |
| Tuba1b | 0.017373934 | 2.099767981 |

Supplementary Table 6 DEG_Categories_6 hrs for Suppl Fig 3B
Differentially expressed genes (FDR < 0.05)

| CRvsAL Shared DEGs | | |
|---|---|---|
| Gene.ID | FDR | FC |
| Hspa8 | 1.03906E-12 | 0.329719601 |
| Hsph1 | 0.006839159 | 0.477064142 |
| Mup1 | 6.18087E-09 | 0.010097401 |
| Mup12 | 4.17358E-08 | 0.010737376 |
| Mup7 | 3.05447E-09 | 0.00407594 |

| FRFvsAL Shared DEGs | | |
|---|---|---|
| Gene.ID | FDR | FC |
| Mup1 | 0.011366454 | 0.087201791 |
| Mup12 | 0.010945952 | 0.080747911 |
| Mup7 | 0.002623612 | 0.037616075 |
| Hspa8 | 4.96907E-07 | 2.114656795 |
| Hsph1 | 0.000290865 | 2.220334561 |

| CRvsFRF Shared DEGs | | |
|---|---|---|
| Gene.ID | FDR | FC |
| Hspa8 | 0 | 0.155921094 |
| Hsph1 | 5.28089E-10 | 0.214861377 |
| Mup1 | 1.37867E-06 | 0.115793507 |
| Mup12 | 2.29426E-06 | 0.132974038 |
| Mup7 | 2.55351E-15 | 0.108356324 |

Supplementary Table 6 DEG_Categories_6 hrs for Suppl Fig 3B
Differentially expressed genes (FDR < 0.05)

| CRvsAL Shared DEGs | | |
|---|---|---|
| Gene.ID | FDR | FC |
| Btg2 | 0.00495161 | 0.206426777 |
| Cyp3a44 | 0.019039249 | 0.094504518 |
| Cyp4a12a | 0.034974566 | 0.097600827 |
| Cyp4a12b | 0.014820192 | 0.088701609 |
| Elovl3 | 0.001280773 | 0.038029918 |
| Hsd3b5 | 2.19351E-07 | 0.017800487 |
| Lifr | 0 | 0.34568742 |
| Mup20 | 0.007198196 | 0.042843659 |
| Ppp1r3g | 0.003863052 | 0.064394163 |
| Selenbp2 | 1.60929E-05 | 0.023803594 |
| Serpina1e | 1.24437E-07 | 0.008296322 |
| Sik1 | 0.000163134 | 0.349703638 |
| Tff3 | 3.48482E-05 | 0.029570646 |
| Tmem254c | 0.011862695 | 2.076932554 |

| FRFvsAL Shared DEGs | | |
|---|---|---|
| Gene.ID | FDR | FC |
| Btg2 | 0.021433049 | 0.267561397 |
| Cyp3a44 | 0.014510401 | 0.101021801 |
| Cyp4a12a | 2.42294E-06 | 0.022186839 |
| Cyp4a12b | 1.65596E-05 | 0.033594648 |
| Elovl3 | 2.31875E-07 | 0.00950548 |
| Hsd3b5 | 2.68941E-06 | 0.026216225 |
| Lifr | 0 | 0.333691086 |
| Mup20 | 0.003966061 | 0.042800836 |
| Ppp1r3g | 0.035362414 | 0.11133218 |
| Selenbp2 | 4.77134E-06 | 0.021396041 |
| Serpina1e | 5.1374E-06 | 0.015785097 |
| Sik1 | 0.01211584 | 0.399190212 |
| Tff3 | 1.01883E-06 | 0.020154729 |
| Tmem254c | 0.009650546 | 2.670148919 |

Supplementary Table 6 DEG_Categories_6 hrs for Suppl Fig 3B
Differentially expressed genes (FDR < 0.05)

| FRFvsAL Shared DEGs | | |
|---|---|---|
| Gene.ID | FDR | FC |
| Car3 | 8.92839E-11 | 0.342294062 |
| Cd74 | 5.54278E-06 | 0.283947022 |
| Cyp2c44 | 1.31249E-08 | 0.3400248 |
| Eif2s3y | 4.46866E-05 | 0.06742173 |
| Gm3839 | 0.004890308 | 0.123964105 |
| H2-Aa | 0.000139745 | 0.254006613 |
| H2-Ab1 | 8.42058E-08 | 0.257073866 |
| Hist1h1c | 0.000309578 | 0.365546451 |
| Scd1 | 0.000204699 | 0.105018443 |
| Serpina6 | 9.98904E-07 | 0.298908448 |
| Srebf1 | 4.41701E-05 | 0.425526834 |
| Tmsb10 | 3.46077E-10 | 0.338083846 |
| 2010003K11Rik | 4.6974E-08 | 4.043170278 |
| Acnat2 | 0.007178589 | 2.473451423 |
| Apoa4 | 0 | 6.545413795 |
| Chordc1 | 0.004960929 | 2.081756682 |
| Creld2 | 0.011082042 | 7.86808613 |
| Cyp17a1 | 1.15015E-06 | 3.368624654 |
| Cyp39a1 | 5.19101E-11 | 3.985225336 |
| Cyp4a10 | 0 | 6.397057353 |
| Cyp4a14 | 0 | 6.528646106 |
| Derl3 | 0.002081432 | 9.532463499 |
| Eif4ebp3 | 6.62193E-09 | 8.178156626 |
| Gfra1 | 0.000302427 | 2.102948454 |
| Herpud1 | 0.030338464 | 2.214491097 |
| Hsp90aa1 | 0.000581582 | 3.39670408 |
| Serpina3n | 2.34579E-12 | 2.070584864 |
| Slc37a4 | 9.00391E-14 | 2.503206758 |
| Stip1 | 0.003161281 | 2.184062514 |
| Tsku | 0.00324637 | 3.011050714 |

| CRvsFRF Shared DEGs | | |
|---|---|---|
| Gene.ID | FDR | FC |
| 2010003K11Rik | 3.10001E-05 | 0.341795397 |
| Acnat2 | 3.7903E-13 | 0.309718767 |
| Apoa4 | 2.20647E-06 | 0.29883668 |
| Chordc1 | 3.383E-07 | 0.298617995 |
| Creld2 | 0.014630042 | 0.127568139 |
| Cyp17a1 | 0 | 0.124964658 |
| Cyp39a1 | 4.26126E-11 | 0.337820631 |
| Cyp4a10 | 6.45052E-07 | 0.373837441 |
| Cyp4a14 | 0 | 0.321037135 |
| Derl3 | 0.035171585 | 0.167123092 |
| Eif4ebp3 | 4.54872E-07 | 0.162237122 |
| Gfra1 | 0.0001126 | 0.438903419 |
| Herpud1 | 4.75608E-09 | 0.289613653 |
| Hsp90aa1 | 0.015521799 | 0.362028605 |
| Serpina3n | 0 | 0.420111889 |
| Slc37a4 | 4.74887E-12 | 0.499425907 |
| Stip1 | 0.001397211 | 0.418692405 |
| Tsku | 1.65467E-07 | 0.165589879 |
| Car3 | 2.95498E-05 | 2.849758306 |
| Cd74 | 0.000206279 | 2.435128533 |
| Cyp2c44 | 5.47883E-09 | 2.383965496 |
| Eif2s3y | 0.001528396 | 13.73634968 |
| Gm3839 | 0.010581277 | 7.092979136 |
| H2-Aa | 0.000869983 | 2.872728508 |
| H2-Ab1 | 0.000139225 | 2.64848315 |
| Hist1h1c | 0.002335057 | 2.160681912 |
| Scd1 | 0 | 7.870097827 |
| Serpina6 | 4.78284E-13 | 2.85833428 |
| Srebf1 | 0.039800679 | 2.302984585 |
| Tmsb10 | 0.000619398 | 2.141808106 |

Supplementary Table 6 DEG_Categories_6 hrs for Suppl Fig 3B
Differentially expressed genes (FDR < 0.05)

| CRvsFRF Shared DEGs | | |
|---|---|---|
| Gene.ID | FDR | FC |
| Dusp1 | 1.22023E-05 | 0.189264555 |
| Fos | 0.027560367 | 0.132198631 |
| Hopx | 7.66623E-05 | 0.348430211 |
| Hspa1b | 0.002464969 | 0.236399644 |
| Jun | 0.006108175 | 0.329578177 |
| Mup10 | 6.49147E-13 | 0.240470445 |
| Mup11 | 2.79909E-12 | 0.035176222 |
| Mup14 | 2.60902E-13 | 0.096473103 |
| Mup15 | 0 | 0.001413698 |
| Mup16 | 0 | 0.122753387 |
| Mup17 | 4.49976E-10 | 0.0039804 |
| Mup18 | 3.33067E-16 | 0.113999206 |
| Mup19 | 9.15677E-07 | 0.071913881 |
| Mup2 | 7.48366E-08 | 0.05906963 |
| Mup9 | 7.21201E-13 | 0.053553029 |
| Rarres1 | 5.43612E-06 | 0.212057294 |
| RP24-346O11.2 | 0 | 0.093517426 |
| Slc25a25 | 0.044455679 | 0.331462874 |
| Sult5a1 | 0.002047342 | 0.275657757 |
| Acta1 | 0.00010867 | 25.10634396 |
| Mb | 4.95839E-05 | 15.02016412 |
| Tnni2 | 0.010831697 | 9.102436623 |
| Tnnt3 | 0.002505476 | 12.22548874 |

| CRvsAL Shared DEGs | | |
|---|---|---|
| Gene.ID | FDR | FC |
| Dusp1 | 1.01839E-11 | 0.115597447 |
| Fos | 0.005857439 | 0.174320551 |
| Hopx | 0.006736729 | 0.499378394 |
| Hspa1b | 0.039212583 | 0.235726313 |
| Jun | 2.22951E-06 | 0.169021647 |
| Mup10 | 2.66454E-15 | 0.185958579 |
| Mup11 | 1.23665E-08 | 0.006997331 |
| Mup14 | 1.63811E-07 | 0.012886355 |
| Mup15 | 3.55271E-15 | 0.000934355 |
| Mup16 | 0 | 0.092246372 |
| Mup17 | 2.14122E-08 | 0.001918107 |
| Mup18 | 0.012879066 | 0.119842277 |
| Mup19 | 3.05356E-07 | 0.087403598 |
| Mup2 | 0 | 0.028062668 |
| Mup9 | 0 | 0.039822341 |
| Rarres1 | 3.48303E-06 | 0.214420869 |
| RP24-346O11.2 | 0 | 0.096949371 |
| Slc25a25 | 0.002815452 | 0.157106953 |
| Sult5a1 | 0.000115781 | 0.124988325 |
| Acta1 | 0 | 26.44159529 |
| Mb | 5.25983E-05 | 15.02016412 |
| Tnni2 | 0.016301641 | 9.351957568 |
| Tnnt3 | 0.016394069 | 12.87568696 |
