## Supplementary Table 7 for "Anticipatory metabolic reprogramming distinguishes caloric restriction from fasting-refeeding cycles"

Supplementary Table 7 DEG_Categories_10 hrs for Suppl Fig 3B
Differentially expressed genes (FDR < 0.05)

| CRvsAL Exclusive DEGs | | |
|---|---|---|
| Gene.ID | FDR | FC |
| Btg2 | 0.018074361 | 0.275419977 |
| Csrnp1 | 0.021857721 | 0.314219241 |
| G0s2 | 1.41813E-07 | 0.1351881 |
| Hsph1 | 0.047844656 | 0.405983733 |
| Hyou1 | 0.044710864 | 0.325354522 |
| Inhbe | 0.016146232 | 0.251918769 |
| Jund | 8.36786E-06 | 0.481557938 |
| Mup1 | 0.007987429 | 0.046786739 |
| Mup12 | 0.009395984 | 0.054042027 |
| Mup14 | 6.12333E-05 | 0.019909746 |
| Mup19 | 0.023206847 | 0.057727283 |
| Mup7 | 0.003695779 | 0.033082777 |
| Mup8 | 0.000470191 | 0.061240089 |
| Noct | 0.045937376 | 0.194574639 |
| Obp2a | 0.015510665 | 0.128833144 |
| Onecut1 | 0.000951733 | 0.230567577 |
| Sik1 | 0.004007467 | 0.236293337 |
| Sult5a1 | 0.004059938 | 0.328908477 |
| Tob1 | 4.44089E-16 | 0.359206063 |
| Ugt2b38 | 0.043767903 | 0.145180872 |
| Ugt3a1 | 0.028150868 | 0.459283709 |
| Zfp36 | 3.73943E-09 | 0.437497816 |
| 8430408G22Rik | 0.029443405 | 2.560806384 |
| Aprt | 0.005128067 | 2.079141535 |
| Cyp3a41a | 0.00579725 | 17.40147625 |
| Cyp51 | 0.000391324 | 2.272736689 |
| Fmo3 | 0.000758436 | 19.82539796 |
| Gm2000 | 0.019237038 | 2.841055584 |
| Slc16a7 | 0.000540592 | 2.137218621 |
| Tceal8 | 0.004589637 | 3.986185892 |

Supplementary Table 7 DEG_Categories_10 hrs for Suppl Fig 3B
Differentially expressed genes (FDR < 0.05)

| FRFvsAL Exclusive DEGs | | |
|---|---|---|
| Gene.ID | FDR | FC |
| Acy3 | 0.001706853 | 0.493370622 |
| Ces2a | 0.000304856 | 0.473320996 |
| Cyp2c44 | 0.001704568 | 0.370454585 |
| Cyp2c55 | 0.000120974 | 0.197766564 |
| Gdf15 | 0.024869969 | 0.171245215 |
| Gsta2 | 0.024370908 | 0.413622711 |
| Gstp2 | 0.009819122 | 0.084090631 |
| Hsd3b5 | 0.010755565 | 0.084449439 |
| Ifi47 | 0.00490783 | 0.412318632 |
| Insc | 0.043241505 | 0.458750518 |
| Jchain | 0.032312734 | 0.099776268 |
| Lasp1 | 0.000200266 | 0.467099661 |
| Mid1ip1 | 0.022016176 | 0.367760542 |
| Nme6 | 0.020416673 | 0.477566028 |
| Pdcd4 | 0.006579932 | 0.47377349 |
| Ppp1r3c | 4.09953E-10 | 0.184325216 |
| Vegfb | 0.048461842 | 0.387712096 |
| Ahcy | 0.048446982 | 2.390152141 |
| Arg1 | 7.06443E-09 | 2.007901709 |
| As3mt | 0.004337359 | 2.36500727 |
| Cyp2c39 | 0.003098366 | 4.321076828 |
| Cyp39a1 | 0.035189664 | 6.840483539 |
| Cyp4a10 | 0.005902887 | 7.949043529 |
| Cyp4a14 | 0.00120481 | 8.97462745 |
| Ddit4 | 0.018615574 | 3.812365268 |
| Fam134b | 0.00210129 | 2.859328681 |
| Fam234b | 0.004519759 | 2.372614199 |
| Hbb-bs | 0.044841923 | 2.010233046 |
| Il6ra | 0.006185253 | 4.035345451 |
| Ip6k2 | 4.97449E-05 | 2.741172199 |
| Mat1a | 0.016532609 | 3.659839784 |
| Nr1i3 | 0.002958318 | 2.422094228 |
| Retsat | 0.021515362 | 3.112872088 |
| Rgs16 | 0.007970218 | 16.89553431 |
| Slc17a8 | 0.007707766 | 4.807119045 |
| Slco1a4 | 0.010437294 | 2.736317403 |
| Sorbs3 | 0.002530659 | 2.382543139 |
| Sult1a1 | 0.00019834 | 2.097352778 |
| Tango2 | 0.044672369 | 2.145641783 |

Supplementary Table 7 DEG_Categories_10 hrs for Suppl Fig 3B

Differentially expressed genes (FDR < 0.05)

| CRvsFRF Exclusive DEGs | | |
|---|---|---|
| Gene.ID | FDR | FC |
| A930011G23Rik | 0.001007345 | 0.183552107 |
| Abcb4 | 0.027790341 | 0.481903319 |
| Agpat9 | 0.005117595 | 0.331564722 |
| Ap3m1 | 0.001391936 | 0.346228539 |
| Arl4a | 0.003769442 | 0.474874474 |
| Cebpb | 6.02008E-06 | 0.368662395 |
| Cidec | 0.008810019 | 0.093640135 |
| Coq10b | 0.000767389 | 0.209808758 |
| Creb3l3 | 5.30332E-06 | 0.432403774 |
| Ctgf | 0.017559327 | 0.185463135 |
| Cyp7a1 | 0.020618962 | 0.380801885 |
| Elmod3 | 0.022729638 | 0.49138887 |
| Fitm2 | 0.024595397 | 0.320175943 |
| Gadd45g | 6.4726E-14 | 0.174541897 |
| Grpel2 | 0.020577119 | 0.418994325 |
| Hmgcs2 | 0.048878597 | 0.454510527 |
| Hopx | 4.60743E-14 | 0.332427515 |
| Hspa1b | 0.005165184 | 0.112801328 |
| Hspb8 | 0.00088072 | 0.424392494 |
| Klf9 | 0.00437277 | 0.417081735 |
| Lipg | 0.00276867 | 0.190078565 |
| Mt2 | 0.006204278 | 0.068264041 |
| Nceh1 | 0.036676174 | 0.374727997 |
| Pex11a | 0.035578478 | 0.220110862 |
| Plin2 | 0.001080728 | 0.435108547 |
| Pole4 | 0.002168244 | 0.487104828 |
| Ppara | 0.009834536 | 0.331976825 |
| Ppp1r3g | 0.000358481 | 0.149831237 |
| Slc22a5 | 0.041307102 | 0.204447306 |
| Slc25a51 | 0.008133415 | 0.436601357 |
| Slc27a1 | 0.044936381 | 0.350462979 |
| Slc27a2 | 0.005194646 | 0.484886068 |
| Slc35g1 | 0.041615389 | 0.399245912 |
| Slc39a14 | 6.88029E-07 | 0.424607708 |
| Tbc1d8 | 0.020110248 | 0.331784445 |
| Tdo2 | 0.031129546 | 0.452989468 |
| Them4 | 0.005360421 | 0.340813782 |
| Tmed5 | 0.025049815 | 0.4453781 |
| Tmem56 | 0.007076959 | 0.434422752 |

Supplementary Table 7 DEG_Categories_10 hrs for Suppl Fig 3B
Differentially expressed genes (FDR < 0.05)

| CRvsFRF Exclusive DEGs | | |
|---|---|---|
| Gene.ID | FDR | FC |
| Trp53inp2 | 5.13569E-07 | 0.323211194 |
| Tsku | 0.032116908 | 0.161880274 |
| Acsl5 | 9.75268E-05 | 2.182315311 |
| Aldoc | 0.012090227 | 3.200976849 |
| Bbc3 | 0.004971926 | 2.948513936 |
| BC048546 | 0.047525161 | 2.559480881 |
| C330021F23Rik | 0.001002741 | 5.630703221 |
| Ces2e | 0.038888854 | 2.695892165 |
| Coq8a | 1.07248E-13 | 2.849523076 |
| Cox7a1 | 4.26904E-05 | 4.455410948 |
| Cyb5b | 0 | 2.342385821 |
| Cyp1a2 | 1.08359E-11 | 2.540754631 |
| Cyp2g1 | 0.033351907 | 3.081479768 |
| Fdps | 9.14334E-05 | 2.428447513 |
| Gcat | 1.27634E-05 | 2.150591861 |
| Gm10642 | 0.001388705 | 6.040786498 |
| Gsta1 | 0.000190646 | 23.0044258 |
| Gstm6 | 8.35475E-05 | 3.091720637 |
| Insig1 | 1.8977E-12 | 2.90654903 |
| Lgals1 | 0.004563922 | 3.706505529 |
| Ly6a | 0.014247446 | 2.303961587 |
| Msmo1 | 1.42905E-05 | 2.157539579 |
| Muc1 | 0.020795287 | 7.007032235 |
| Mvd | 0.00445251 | 3.740244721 |
| Orm3 | 0.031844758 | 2.927548103 |
| Osgin1 | 4.36619E-08 | 5.120847934 |
| Pdzk1ip1 | 0.0163805 | 4.359308232 |
| Psen2 | 0.000733243 | 2.071578736 |
| Pxdc1 | 0.002263661 | 2.24103653 |
| Ropn1l | 0.000348178 | 2.462240749 |
| S1pr1 | 0.004731088 | 2.342828943 |
| Sc5d | 1.66644E-13 | 3.005494732 |
| Sigmar1 | 4.2861E-05 | 2.120776396 |
| Siva1 | 0.003549012 | 2.985741484 |
| Smim1 | 0.001560404 | 3.367820311 |
| Sucnr1 | 0.016027135 | 4.184259488 |
| Thrsp | 0.013752761 | 3.688792698 |
| Tkt | 3.81112E-08 | 2.294293887 |
| Tstd1 | 0.000854875 | 2.6113004 |

Supplementary Table 7 DEG_Categories_10 hrs for Suppl Fig 3B
Differentially expressed genes (FDR < 0.05)

| CRvsFRF Exclusive DEGs | | |
| --- | --- | --- |
| Gene.ID | FDR | FC |
| Tuba1b | 2.29444E-05 | 2.354467994 |
| Ugdh | 0.027533708 | 2.237162833 |
| Wfdc2 | 0.045787325 | 3.102444038 |

Supplementary Table 7 DEG_Categories_10 hrs for Suppl Fig 3B
Differentially expressed genes (FDR < 0.05)

| CRvsAL Shared DEGs | | |
|---|---|---|
| Gene.ID | FDR | FC |
| Bhlhe40 | 0 | 0.179069711 |
| Dusp1 | 0 | 0.107379022 |
| Nr4a1 | 6.66134E-16 | 0.027050954 |
| Pck1 | 0.003938237 | 0.28193835 |

| FRFvsAL Shared DEGs | | |
|---|---|---|
| Gene.ID | FDR | FC |
| Bhlhe40 | 6.87233E-05 | 0.448387085 |
| Dusp1 | 0 | 0.33630968 |
| Nr4a1 | 0.000197095 | 0.218105436 |
| Pck1 | 2.23816E-09 | 2.175733671 |

| CRvsFRF Shared DEGs | | |
|---|---|---|
| Gene.ID | FDR | FC |
| Bhlhe40 | 3.04574E-06 | 0.399364114 |
| Dusp1 | 5.96039E-05 | 0.319286148 |
| Nr4a1 | 0.0310293 | 0.12402696 |
| Pck1 | 0 | 0.129583117 |

Supplementary Table 7 DEG_Categories_10 hrs for Suppl Fig 3B

Differentially expressed genes (FDR < 0.05)

| CRvsAL Shared DEGs | | |
|---|---|---|
| Gene.ID | FDR | FC |
| Cyr61 | 0.000204466 | 0.266684674 |
| Dusp6 | 1.15068E-10 | 0.254579489 |
| Id3 | 0.004149076 | 0.411892017 |
| Keg1 | 0.016783251 | 0.392584795 |
| Lifr | 0 | 0.364373576 |
| Selenbp2 | 1.00969E-06 | 0.015436898 |
| Cyp3a16 | 0.010564367 | 11.96980906 |
| Gm45531 | 0.006361191 | 3.358350692 |
| Rdh9 | 4.41869E-14 | 3.114868483 |
| Serpina3m | 0.001419526 | 2.070521019 |

| FRFvsAL Shared DEGs | | |
|---|---|---|
| Gene.ID | FDR | FC |
| Cyr61 | 0.014708634 | 0.406348634 |
| Dusp6 | 0.019141296 | 0.491422411 |
| Id3 | 6.98329E-05 | 0.322617893 |
| Keg1 | 1.24039E-05 | 0.359326116 |
| Lifr | 4.53388E-11 | 0.410652814 |
| Selenbp2 | 0.004242797 | 0.06373172 |
| Cyp3a16 | 0.019041011 | 9.294789528 |
| Gm45531 | 1.2575E-05 | 3.490774658 |
| Rdh9 | 0.013123237 | 2.089666984 |
| Serpina3m | 0.001175985 | 2.740920644 |

Supplementary Table 7 DEG_Categories_10 hrs for Suppl Fig 3B
Differentially expressed genes (FDR < 0.05)

| FRFvsAL Shared DEGs | | | CRvsFRF Shared DEGs | | |
|---|---|---|---|---|---|
| Gene.ID | FDR | FC | Gene.ID | FDR | FC |
| Acly | 0.04364039 | 0.303217011 | 1600002H07Rik | 7.2875E-13 | 0.213913904 |
| Adh4 | 0.014363238 | 0.410331547 | 1810055G02Rik | 5.0601E-07 | 0.120647834 |
| Angptl8 | 1.29435E-06 | 0.108695515 | 2010003K11Rik | 0.000116703 | 0.242359705 |
| Aqp8 | 2.15265E-08 | 0.272605726 | Alas1 | 1.72734E-05 | 0.147989194 |
| BC029214 | 0.020085239 | 0.38396057 | Angptl4 | 2.9629E-10 | 0.110734943 |
| Car3 | 6.14948E-05 | 0.13689699 | Apoa4 | 0.007441547 | 0.183364557 |
| Cyp26a1 | 1.56333E-08 | 0.102920382 | Apoa5 | 0 | 0.442838487 |
| Efna1 | 0 | 0.35507563 | Atl2 | 0.012717424 | 0.341261513 |
| Gale | 7.19808E-05 | 0.140852682 | Bnip3 | 0.025703469 | 0.447860601 |
| Gclc | 4.93279E-07 | 0.474899565 | Cdip1 | 0.000309097 | 0.383290078 |
| Gm3776 | 0.038819291 | 0.163704701 | Cdkn1a | 3.68234E-07 | 0.077929089 |
| Kyat1 | 1.30408E-07 | 0.434248237 | Cpt1a | 3.16018E-08 | 0.315337091 |
| Nat8f1 | 1.31501E-06 | 0.345050774 | Cyp17a1 | 0.018879976 | 0.127706242 |
| Nr0b2 | 2.5198E-10 | 0.225132061 | Cyp4a31 | 0.000275218 | 0.150386397 |
| Paox | 0.030061255 | 0.494928916 | E030018B13Rik | 0.010876961 | 0.17692584 |
| Raet1d | 0.007838474 | 0.234720215 | Eif4ebp3 | 7.55792E-05 | 0.083969306 |
| Scd1 | 0.023765843 | 0.095248142 | Fam35a | 0.003982336 | 0.268859165 |
| Srebf1 | 0.001085776 | 0.212724931 | Fbxo31 | 0.000235185 | 0.220467117 |
| Tubb2a | 2.90875E-05 | 0.427783949 | Fgf21 | 4.09162E-12 | 0.043625908 |
| 1600002H07Rik | 0.000302441 | 2.439706232 | Gfra1 | 0.000164012 | 0.265400503 |
| 1810055G02Rik | 2.24208E-05 | 5.707463928 | Glrx | 0.035889335 | 0.478156591 |
| 2010003K11Rik | 0.007697362 | 3.127722813 | Got1 | 4.62828E-10 | 0.328359334 |
| Alas1 | 0.029047252 | 3.125637739 | Gpcpd1 | 2.93669E-05 | 0.296331364 |
| Angptl4 | 0.000786678 | 3.4452091 | Gpt2 | 0.000986451 | 0.479263865 |
| Apoa4 | 5.52703E-06 | 11.74480823 | Igfbp1 | 0 | 0.09410833 |
| Apoa5 | 0 | 2.077727325 | Mfsd2a | 1.96774E-09 | 0.065050183 |
| Atl2 | 0.036661681 | 2.685058769 | Nlrp12 | 0.000272836 | 0.312097301 |
| Bnip3 | 0.000480735 | 3.060683858 | Per1 | 0.000678418 | 0.217936701 |
| Cdip1 | 3.78791E-05 | 3.058564306 | Plin5 | 0.012532316 | 0.239327464 |
| Cdkn1a | 6.53791E-05 | 7.463751448 | Pnpla2 | 0.003270388 | 0.442650366 |
| Cpt1a | 1.49425E-05 | 2.546078899 | Rab30 | 3.58432E-05 | 0.140617074 |
| Cyp17a1 | 0.005033938 | 9.813560534 | Rapgef4 | 0.00088131 | 0.356145138 |
| Cyp4a31 | 0.000334605 | 6.764714137 | Serpina3n | 5.0008E-07 | 0.461612755 |
| E030018B13Rik | 0.003201 | 7.165512526 | Slc25a47 | 0 | 0.270990606 |
| Eif4ebp3 | 2.42711E-06 | 18.84442915 | Slc37a4 | 4.7252E-07 | 0.434132373 |
| Fam35a | 0.028673453 | 2.990676762 | St3gal5 | 0.0005243 | 0.096295482 |
| Fbxo31 | 0.000117407 | 4.934066452 | Tmem120a | 0.003237935 | 0.471021765 |
| Fgf21 | 0.00062202 | 3.834702215 | Txnip | 2.21265E-10 | 0.378243606 |
| Gfra1 | 0.000307541 | 3.638248996 | Usp2 | 0.019650832 | 0.148935781 |

Supplementary Table 7 DEG_Categories_10 hrs for Suppl Fig 3B
Differentially expressed genes (FDR < 0.05)

| FRFvsAL Shared DEGs | | |
|---|---|---|
| Gene.ID | FDR | FC |
| Glrx | 0.00039653 | 3.01062987 |
| Got1 | 2.83398E-09 | 2.972949639 |
| Gpcpd1 | 2.06231E-05 | 3.2825194 |
| Gpt2 | 0.001553804 | 2.005570322 |
| Igfbp1 | 2.97667E-10 | 3.818077587 |
| Mfsd2a | 1.24117E-05 | 6.954023868 |
| Nlrp12 | 0.000353739 | 3.130749294 |
| Per1 | 0.043719312 | 3.137383695 |
| Plin5 | 0.015329904 | 4.217531663 |
| Pnpla2 | 9.72319E-06 | 2.960266725 |
| Rab30 | 0.001077678 | 5.505938083 |
| Rapgef4 | 0.039194417 | 2.107552333 |
| Serpina3n | 6.73635E-06 | 2.432654034 |
| Slc25a47 | 2.19298E-10 | 2.670709961 |
| Slc37a4 | 2.65641E-11 | 2.942574183 |
| St3gal5 | 0.00013433 | 13.17118133 |
| Tmem120a | 0.007979569 | 2.108159673 |
| Txnip | 9.34808E-14 | 2.381250242 |
| Usp2 | 0.012177552 | 7.702304032 |

| CRvsFRF Shared DEGs | | |
|---|---|---|
| Gene.ID | FDR | FC |
| Acly | 0 | 4.837481343 |
| Adh4 | 0.009221182 | 2.520616367 |
| Angptl8 | 0.01131172 | 2.845823297 |
| Aqp8 | 0.04108393 | 10.16667828 |
| BC029214 | 4.18138E-05 | 4.003450059 |
| Car3 | 0.016557828 | 6.213641972 |
| Cyp26a1 | 0.043608481 | 10.40313455 |
| Efna1 | 4.06856E-08 | 2.03651117 |
| Gale | 1.47142E-07 | 4.724000606 |
| Gclc | 2.09166E-12 | 2.062808907 |
| Gm3776 | 6.23528E-05 | 27.1169993 |
| Kyat1 | 1.62568E-09 | 2.030419286 |
| Nat8f1 | 2.82791E-06 | 2.820371451 |
| Nr0b2 | 0.000736542 | 3.747548101 |
| Paox | 0.037249592 | 2.190614184 |
| Raet1d | 1.04001E-06 | 5.397034002 |
| Scd1 | 4.0315E-05 | 10.70678171 |
| Srebf1 | 7.79148E-10 | 5.379739214 |
| Tubb2a | 0.000685596 | 2.561512372 |

Supplementary Table 7 DEG_Categories_10 hrs for Suppl Fig 3B
Differentially expressed genes (FDR < 0.05)

| CRvsFRF Shared DEGs | | |
|---|---|---|
| Gene.ID | FDR | FC |
| Dio1 | 6.80604E-05 | 0.149231118 |
| Errfi1 | 0 | 0.156951367 |
| G6pc | 0.00225372 | 0.28494423 |
| Jun | 0.00398169 | 0.1792761 |
| Mafb | 0.001105908 | 0.142696193 |
| Mup11 | 0.005975479 | 0.037855968 |
| Mup15 | 3.6593E-12 | 0.001979352 |
| Mup17 | 1.23974E-10 | 0.002294504 |
| Mup2 | 0.004915404 | 0.09789139 |
| Mup9 | 0.002035678 | 0.074429254 |
| Pim3 | 1.42694E-08 | 0.276405668 |
| Plk3 | 5.9047E-05 | 0.074030985 |
| Slc25a22 | 7.2274E-10 | 0.253599106 |
| Slc25a25 | 6.05917E-08 | 0.114257401 |
| Slc25a30 | 0.000556615 | 0.182399799 |
| Slc38a2 | 4.87717E-09 | 0.236544851 |
| Tat | 0.042431593 | 0.195517196 |
| Trp53inp1 | 3.42647E-05 | 0.222935818 |
| Acat2 | 0 | 2.823723378 |
| Acss2 | 4.7624E-10 | 4.798324534 |
| Car2 | 0.048200409 | 2.812805498 |
| Ces1g | 8.42261E-11 | 2.630564588 |
| Dhcr7 | 0.000107753 | 2.703414266 |
| Dmbt1 | 0.00015671 | 21.5174812 |
| Ethe1 | 4.44089E-16 | 4.090209402 |
| Fdft1 | 5.19499E-08 | 2.505184211 |
| Hmgcs1 | 0 | 3.674674216 |
| Idi1 | 0.023619879 | 2.057913626 |
| Lss | 0.000278819 | 3.569989049 |
| Orm2 | 9.5024E-13 | 7.412383418 |
| Pmvk | 2.43838E-12 | 3.399847452 |
| Ppp1r1b | 0.000357388 | 3.61956929 |
| Rdh11 | 5.32371E-05 | 2.924118503 |
| Sqle | 6.3219E-06 | 2.008677317 |
| Sult2a3 | 0.000335492 | 21.6395993 |

| CRvsAL Shared DEGs | | |
|---|---|---|
| Gene.ID | FDR | FC |
| Dio1 | 0.000448458 | 0.198062988 |
| Errfi1 | 6.66134E-15 | 0.265867139 |
| G6pc | 4.996E-15 | 0.252565254 |
| Jun | 8.82072E-13 | 0.165036889 |
| Mafb | 0 | 0.13480028 |
| Mup11 | 3.47183E-06 | 0.012291175 |
| Mup15 | 8.88178E-16 | 0.000707649 |
| Mup17 | 1.11022E-16 | 0.000442382 |
| Mup2 | 2.78097E-05 | 0.026001682 |
| Mup9 | 2.96037E-05 | 0.023669167 |
| Pim3 | 0.009270801 | 0.431165838 |
| Plk3 | 0.000132264 | 0.114584832 |
| Slc25a22 | 0.002336807 | 0.448494067 |
| Slc25a25 | 0 | 0.101147395 |
| Slc25a30 | 5.23486E-07 | 0.205722763 |
| Slc38a2 | 0.037011633 | 0.297856396 |
| Tat | 0.019092602 | 0.334095245 |
| Trp53inp1 | 0.001138599 | 0.388977527 |
| Acat2 | 4.43518E-10 | 2.102291261 |
| Acss2 | 0.000103262 | 2.872021984 |
| Car2 | 0.020409354 | 3.531400703 |
| Ces1g | 1.87781E-05 | 2.447662751 |
| Dhcr7 | 0.001205284 | 2.109956807 |
| Dmbt1 | 1.47327E-13 | 21.6229654 |
| Ethe1 | 8.06521E-08 | 2.306246414 |
| Fdft1 | 1.12939E-06 | 2.252318487 |
| Hmgcs1 | 3.2363E-13 | 2.514744126 |
| Idi1 | 0.000579848 | 2.626687647 |
| Lss | 7.18669E-09 | 2.796801379 |
| Orm2 | 1.33227E-15 | 9.061756427 |
| Pmvk | 9.04519E-11 | 3.108445329 |
| Ppp1r1b | 0.015236262 | 3.043678364 |
| Rdh11 | 0.000165256 | 2.111882665 |
| Sqle | 1.03927E-05 | 2.314754894 |
| Sult2a3 | 0.000368492 | 23.88118856 |
