## Supplementary Table 8 for "Anticipatory metabolic reprogramming distinguishes caloric restriction from fasting-refeeding cycles"

Supplementary Table 8 DEG_Categories_14 hrs for Suppl Fig 3B
Differentially expressed genes (FDR < 0.05)

| CRvsAL Exclusive DEGs | | |
|---|---|---|
| Gene.ID | FDR | FC |
| Btg2 | 0.020628905 | 0.345566871 |
| G0s2 | 0.001292948 | 0.162423862 |
| Gdf15 | 0.00142184 | 0.136201126 |
| Hsp90aa1 | 0.002062573 | 0.469595745 |
| Hyou1 | 0.022505635 | 0.391344507 |
| Inhbc | 1.32393E-07 | 0.36890585 |
| Jun | 0.030900605 | 0.250835494 |
| Pim3 | 0.002760151 | 0.369924872 |
| Plk3 | 0.04999262 | 0.403246869 |
| Selenbp2 | 0.000578303 | 0.043525911 |
| Slc25a25 | 0.020583042 | 0.273704342 |
| Stip1 | 0.000403501 | 0.494889399 |
| Ugt3a1 | 0.001977724 | 0.491295308 |
| Zfp36 | 0.007298211 | 0.352174953 |
| Aprt | 6.93001E-13 | 2.346075761 |
| Clpx | 0 | 2.296153286 |
| Cyp2a5 | 3.5032E-06 | 2.116683885 |
| Fmo3 | 0.038831919 | 10.35851438 |
| Gm10273 | 0.038187069 | 4.956357514 |
| Gstt2 | 0.010705171 | 2.146404745 |
| Gstt3 | 0.012572436 | 2.110977289 |
| Hbb-bs | 0.004598804 | 2.019371244 |
| Orm3 | 0.001918854 | 4.820189929 |
| Pmvk | 0.000165039 | 2.059035396 |
| Rbm3 | 3.49078E-06 | 2.044923491 |

Supplementary Table 8 DEG_Categories_14 hrs for Suppl Fig 3B
Differentially expressed genes (FDR < 0.05)

| FRFvsAL Exclusive DEGs | | |
|---|---|---|
| Gene.ID | FDR | FC |
| 5830473C10Rik | 0.025729688 | 0.471177326 |
| Adh4 | 2.68864E-07 | 0.481221809 |
| BC048546 | 0.005941396 | 0.40416292 |
| Cbs | 0.015482399 | 0.496967066 |
| Ces2a | 2.19935E-13 | 0.417968837 |
| Cyp2c44 | 0.000345816 | 0.351050548 |
| Gstp2 | 0.005034589 | 0.054604865 |
| Hao2 | 0.044294877 | 0.148071785 |
| Hsd3b5 | 0.003563222 | 0.069954244 |
| Id2 | 0.00016351 | 0.475039923 |
| Ifi47 | 0.000594833 | 0.350948374 |
| Klf10 | 0.04464656 | 0.399565773 |
| Kyat1 | 1.2812E-13 | 0.429827948 |
| Mapk15 | 0.00249862 | 0.233596393 |
| Mid1ip1 | 0.000128391 | 0.458788419 |
| Mug2 | 0.029860608 | 0.448090548 |
| Rnf186 | 0.00024923 | 0.275782446 |
| Serpina7 | 0.006027604 | 0.381037783 |
| Smim22 | 0.011345641 | 0.292148511 |
| Srebf1 | 0 | 0.226381442 |
| St6gal1 | 0.042299751 | 0.489189331 |
| Tmsb10 | 0.049207002 | 0.361428792 |
| 1600002H07Rik | 0.010790999 | 2.219038671 |
| 1810055G02Rik | 0.023838499 | 2.774951216 |
| Ahcy | 7.24265E-10 | 2.638011827 |
| Amy1 | 0 | 2.059421204 |
| Angptl4 | 0.026152607 | 2.569692651 |
| Apoa4 | 0.001653831 | 13.00902663 |
| Asl | 0 | 2.322347719 |
| Atl2 | 0.00011863 | 2.014446649 |
| Bnip3 | 5.00965E-09 | 2.153730886 |
| Cdip1 | 0.001567311 | 2.407302354 |
| Coa6 | 3.2311E-06 | 2.212206018 |
| Cyp2c39 | 0.03957352 | 4.65035248 |
| Cyp39a1 | 0.001577605 | 5.849358552 |
| Cyp4a31 | 0.000155979 | 4.385782733 |
| Fmo2 | 0.036789011 | 5.200810381 |
| Fmo5 | 9.19642E-12 | 2.218931667 |
| Glrx | 0 | 3.369649354 |

Supplementary Table 8 DEG_Categories_14 hrs for Suppl Fig 3B
Differentially expressed genes (FDR < 0.05)

| FRFvsAL Exclusive DEGs | | |
|---|---|---|
| Gene.ID | FDR | FC |
| Il1r1 | 0.016707964 | 2.427407522 |
| Mt2 | 0.022723013 | 8.865075729 |
| Nlrp12 | 0.018074945 | 3.480329542 |
| Nr1i3 | 0.003510061 | 2.115933402 |
| Pfkfb1 | 0.012320904 | 2.747144105 |
| Retsat | 0.000521702 | 2.614109532 |
| Rgs16 | 0.011801666 | 3.681978011 |
| Rhbg | 0.037317117 | 2.934028577 |
| Serpina3m | 4.83213E-07 | 2.67437228 |
| Slc17a8 | 0.027373064 | 5.218707326 |
| Slc37a4 | 0 | 2.898122903 |
| Sult1a1 | 0.00188464 | 2.218504276 |
| Tango2 | 0.011184146 | 2.692681294 |
| Tmem120a | 4.6229E-05 | 2.04822988 |
| Ttpa | 2.55155E-08 | 2.061878083 |

Supplementary Table 8 DEG_Categories_14 hrs for Suppl Fig 3B
Differentially expressed genes (FDR < 0.05)

| CRvsFRF Exclusive DEGs | | |
|---|---|---|
| Gene.ID | FDR | FC |
| A1cf | 0.004871402 | 0.401659252 |
| Abhd6 | 4.00037E-05 | 0.390806527 |
| Acot4 | 0.048036711 | 0.466097242 |
| Ahsa1 | 0.045890894 | 0.460036151 |
| Cacybp | 0.003561833 | 0.456017843 |
| Cebpb | 0.002418713 | 0.445137986 |
| Cebpd | 0.0375835 | 0.303214639 |
| Crat | 4.31744E-05 | 0.340606614 |
| Cyp17a1 | 0.01055798 | 0.091992399 |
| Dio1 | 0 | 0.153957647 |
| Dnaja1 | 0 | 0.354136483 |
| Dynll1 | 0.013701481 | 0.467132466 |
| Etfbkmt | 1.22624E-09 | 0.251754649 |
| Fabp2 | 0.012025685 | 0.490163231 |
| Fam35a | 0.014335629 | 0.235088228 |
| Fgf21 | 9.90489E-08 | 0.126719832 |
| Foxa3 | 0.002527939 | 0.436899341 |
| Gadd45b | 8.87703E-05 | 0.267517435 |
| Gck | 0.009335952 | 0.384447794 |
| Herpud1 | 2.66669E-09 | 0.340758128 |
| Hist2h3c1 | 0.012044045 | 0.234715438 |
| Hp | 0 | 0.443976204 |
| Hsdl2 | 2.67598E-11 | 0.445121134 |
| Hsp90ab1 | 0.013681802 | 0.37503547 |
| Hspa1b | 0.003787229 | 0.187806058 |
| Igfbp1 | 0.000617171 | 0.060108423 |
| Lcn2 | 0.042837442 | 0.221695292 |
| Lpin2 | 0.031053685 | 0.45847545 |
| Mfsd2a | 0.002311192 | 0.260575632 |
| Mgll | 1.11022E-16 | 0.379140675 |
| Mup18 | 0.000730212 | 0.305422537 |
| Ncoa4 | 2.45359E-14 | 0.469321742 |
| Nfil3 | 2.42234E-08 | 0.240424781 |
| Nmrk1 | 0.021106692 | 0.452547522 |
| Nnmt | 0 | 0.415341882 |
| Pnrc1 | 9.19373E-07 | 0.35484395 |
| Rsrp1 | 0.000356531 | 0.339831511 |
| Rtp3 | 0.009069521 | 0.466177932 |
| Slc17a3 | 0.004131384 | 0.377027966 |

Supplementary Table 8 DEG_Categories_14 hrs for Suppl Fig 3B
Differentially expressed genes (FDR < 0.05)

| CRvsFRF Exclusive DEGs | | |
|---|---|---|
| Gene.ID | FDR | FC |
| Slc22a5 | 0.012694058 | 0.419412212 |
| Tat | 0.000215985 | 0.307169946 |
| Trp53inp1 | 0.01146439 | 0.286995544 |
| Trp53inp2 | 0.000521608 | 0.390937404 |
| Ubb | 1.62986E-10 | 0.426257473 |
| Usp2 | 3.5102E-05 | 0.283659704 |
| Vnn1 | 1.55169E-08 | 0.319689265 |
| Zfp970 | 0.001109598 | 0.319587127 |
| 3010026O09 Rik | 0.001982996 | 4.987933309 |
| Abca8a | 0.028936185 | 2.269578074 |
| Abcg5 | 6.22203E-07 | 2.071892208 |
| Adssl1 | 0.00668495 | 2.127129027 |
| Agxt | 1.06676E-09 | 2.733383303 |
| BC029214 | 7.83427E-08 | 3.090945844 |
| Camk1d | 0.007800773 | 2.445641994 |
| Car3 | 0 | 4.507712785 |
| Cyp1a2 | 0.001901818 | 3.321369933 |
| Cyp2g1 | 0.025362745 | 10.58836471 |
| Fam129b | 0.020456163 | 3.88558216 |
| Fam213b | 0.027496611 | 2.371358044 |
| Fbxo21 | 5.12956E-12 | 2.714424458 |
| Fus | 2.42042E-06 | 2.844736701 |
| Galt | 9.34217E-09 | 2.073116199 |
| H2-Q6 | 0.015530741 | 2.067104109 |
| Hal | 0 | 3.110666792 |
| Id3 | 9.45206E-06 | 2.767253552 |
| Inca1 | 0.000201734 | 2.304755822 |
| Lpl | 0.001211751 | 4.205786769 |
| Mgst3 | 0.048975293 | 2.07954178 |
| Pltp | 0.001414884 | 2.067924076 |
| Raet1d | 2.42446E-09 | 4.978950888 |
| Raet1e | 0.000260849 | 3.712821903 |
| Rassf3 | 0.034973617 | 2.177294493 |
| Sgk2 | 0.014989915 | 2.192160028 |
| Sucnr1 | 7.30787E-05 | 4.159320734 |
| Sult2a3 | 0.019043824 | 8.980018126 |
| Tstd1 | 2.86638E-06 | 2.19229116 |

Supplementary Table 8 DEG_Categories_14 hrs for Suppl Fig 3B
Differentially expressed genes (FDR < 0.05)

| CRvsAL Shared DEGs | | |
|---|---|---|
| Gene.ID | FDR | FC |
| Hsph1 | 0.029425579 | 0.244008047 |
| Tsku | 0.001978047 | 0.254856684 |
| Gstm3 | 0.00396749 | 2.060009823 |
| Lsm7 | 0 | 34.64780163 |

| FRFvsAL Shared DEGs | | |
|---|---|---|
| Gene.ID | FDR | FC |
| Gstm3 | 1.06581E-14 | 0.245599686 |
| Hsph1 | 0.00187517 | 2.520365401 |
| Lsm7 | 0.006970163 | 12.84734028 |
| Tsku | 1.87594E-12 | 3.276717952 |

| CRvsFRF Shared DEGs | | |
|---|---|---|
| Gene.ID | FDR | FC |
| Hsph1 | 0 | 0.096814552 |
| Tsku | 0 | 0.077778035 |
| Gstm3 | 0 | 8.387672884 |
| Lsm7 | 0.022751137 | 2.696885181 |

Supplementary Table 8 DEG_Categories_14 hrs for Suppl Fig 3B
Differentially expressed genes (FDR < 0.05)

| CRvsAL Shared DEGs | | |
|---|---|---|
| Gene.ID | FDR | FC |
| Cxcl1 | 0.022265361 | 0.229017157 |
| Cyr61 | 0.00297455 | 0.375262697 |
| Lifr | 1.27864E-10 | 0.303935784 |
| Tff3 | 2.22045E-15 | 0.042816414 |

| FRFvsAL Shared DEGs | | |
|---|---|---|
| Gene.ID | FDR | FC |
| Cxcl1 | 0.01505286 | 0.214009393 |
| Cyr61 | 0.000238683 | 0.346785245 |
| Lifr | 4.09125E-07 | 0.356392612 |
| Tff3 | 0.000252244 | 0.046159476 |

Supplementary Table 8 DEG_Categories_14 hrs for Suppl Fig 3B
Differentially expressed genes (FDR < 0.05)

| FRFvsAL Shared DEGs | | |
|---|---|---|
| Gene.ID | FDR | FC |
| Amdhd1 | 5.57372E-06 | 0.487987024 |
| Arrdc3 | 2.68599E-05 | 0.231401742 |
| Cyp2c55 | 0.001658716 | 0.129297271 |
| Dbp | 0.000110543 | 0.164544799 |
| Gm3776 | 0.007723816 | 0.101285175 |
| Gsta1 | 0.016258844 | 0.123625829 |
| Gstm1 | 0 | 0.430234471 |
| Gstm4 | 3.73787E-06 | 0.435750351 |
| Gstm6 | 3.33067E-16 | 0.396886601 |
| Lgals1 | 0.001735812 | 0.369438393 |
| Nat8f1 | 0 | 0.457133952 |
| Nr1d1 | 0 | 0.057618428 |
| Scd1 | 5.64936E-05 | 0.069532347 |
| Aifm2 | 0.000191404 | 2.196031644 |
| Arl4a | 0.046220987 | 2.222607527 |
| As3mt | 1.88423E-07 | 2.376443032 |
| Ccng2 | 0.00275 | 2.616706811 |
| Cdkn1a | 0.000153445 | 17.28975424 |
| Ces1d | 2.71136E-05 | 2.46629416 |
| Chd9 | 0.010569413 | 4.620925002 |
| Cry1 | 0.001698358 | 5.843971712 |
| Ctgf | 0.000628982 | 3.283161729 |
| Cyp4a10 | 9.18607E-10 | 5.930438765 |
| Cyp4a14 | 1.13282E-10 | 4.622591125 |
| Ddit4 | 0.001417047 | 10.69451644 |
| Eif4ebp3 | 2.67286E-11 | 14.17644158 |
| Fbxo31 | 0.000653774 | 3.278183194 |
| Fgf1 | 4.68636E-12 | 2.895405815 |
| Fkbp5 | 2.28963E-06 | 5.383674095 |
| Gde1 | 1.9929E-05 | 2.459267439 |
| Gfra1 | 9.28813E-12 | 2.869587849 |
| Hspa8 | 0.000117043 | 2.078336637 |
| Il6ra | 9.60761E-08 | 3.886163439 |
| Ip6k2 | 2.76125E-10 | 2.901187257 |
| Ivns1abp | 2.05747E-05 | 2.163347386 |
| Lpin1 | 5.784E-08 | 9.289259167 |
| Lrg1 | 2.66454E-14 | 2.3000976 |
| Mat1a | 4.58673E-06 | 2.849948828 |
| Nampt | 0.038290906 | 2.269902024 |

| CRvsFRF Shared DEGs | | |
|---|---|---|
| Gene.ID | FDR | FC |
| Aifm2 | 0.000294257 | 0.49856018 |
| Arl4a | 0.032091365 | 0.447026458 |
| As3mt | 0.000225865 | 0.490627686 |
| Ccng2 | 0.000627983 | 0.331461323 |
| Cdkn1a | 0.000101703 | 0.055061434 |
| Ces1d | 0.004072834 | 0.458669803 |
| Chd9 | 0.008231893 | 0.212710177 |
| Cry1 | 0.0070084 | 0.218247661 |
| Ctgf | 7.45976E-06 | 0.236001609 |
| Cyp4a10 | 0.021718358 | 0.409313891 |
| Cyp4a14 | 4.78826E-06 | 0.398592195 |
| Ddit4 | 0.00341749 | 0.110703741 |
| Eif4ebp3 | 0.02200211 | 0.316437915 |
| Fbxo31 | 0.01052426 | 0.39452836 |
| Fgf1 | 0 | 0.342059371 |
| Fkbp5 | 2.72295E-05 | 0.219495056 |
| Gde1 | 1.62422E-06 | 0.375714303 |
| Gfra1 | 2.71602E-07 | 0.440360711 |
| Hspa8 | 0 | 0.195580784 |
| Il6ra | 2.30994E-06 | 0.301625041 |
| Ip6k2 | 8.32667E-15 | 0.351056578 |
| Ivns1abp | 4.45565E-07 | 0.440364934 |
| Lpin1 | 1.3459E-06 | 0.144234118 |
| Lrg1 | 0 | 0.498102393 |
| Mat1a | 0.004740869 | 0.460254035 |
| Nampt | 0.026977795 | 0.44018822 |
| Parp16 | 0.013022636 | 0.390709358 |
| Pex11a | 0.005622518 | 0.440303055 |
| Plin5 | 1.01295E-05 | 0.359976086 |
| Ppp1r3b | 1.08887E-08 | 0.279539508 |
| Rorc | 6.11145E-07 | 0.279267805 |
| Serinc3 | 2.30217E-10 | 0.499564515 |
| Serpina3n | 0 | 0.452220644 |
| Slc45a3 | 0.001107802 | 0.310252795 |
| St3gal5 | 0.026893856 | 0.360490517 |
| Sult1d1 | 0.017115961 | 0.396722408 |
| Tsc22d3 | 0 | 0.202305715 |
| Zbtb16 | 0.00347502 | 0.229244574 |
| Amdhd1 | 2.44249E-15 | 2.646030103 |

Supplementary Table 8 DEG_Categories_14 hrs for Suppl Fig 3B
Differentially expressed genes (FDR < 0.05)

| FRFvsAL Shared DEGs | | |
|---|---|---|
| Gene.ID | FDR | FC |
| Parp16 | 0.012619249 | 2.647401735 |
| Pex11a | 0.037811906 | 2.1931508 |
| Plin5 | 1.10342E-08 | 4.177744505 |
| Ppp1r3b | 0.001617113 | 2.275121591 |
| Rorc | 7.7939E-10 | 4.883855614 |
| Serinc3 | 2.29972E-12 | 2.178291959 |
| Serpina3n | 0 | 2.745454518 |
| Slc45a3 | 0.000579941 | 3.544694051 |
| St3gal5 | 1.9936E-05 | 5.029112641 |
| Sult1d1 | 0.00651631 | 2.78904922 |
| Tsc22d3 | 0 | 3.646002753 |
| Zbtb16 | 0.001260653 | 5.018932932 |

| CRvsFRF Shared DEGs | | |
|---|---|---|
| Gene.ID | FDR | FC |
| Arrdc3 | 0.00122058 | 6.408619405 |
| Cyp2c55 | 0.002934194 | 4.892127804 |
| Dbp | 1.07395E-07 | 4.543284307 |
| Gm3776 | 3.0117E-12 | 16.07315966 |
| Gsta1 | 1.57074E-12 | 13.17254815 |
| Gstm1 | 0 | 2.80417652 |
| Gstm4 | 4.44089E-16 | 2.928512059 |
| Gstm6 | 5.24403E-12 | 3.248930564 |
| Lgals1 | 0.015051404 | 2.111812238 |
| Nat8f1 | 6.66134E-16 | 2.065597558 |
| Nr1d1 | 0 | 18.79643084 |
| Scd1 | 0.001061206 | 6.078213743 |

Supplementary Table 8 DEG_Categories_14 hrs for Suppl Fig 3B

Differentially expressed genes (FDR < 0.05)

| CRvsFRF Shared DEGs | | |
|---|---|---|
| Gene.ID | FDR | FC |
| Derl3 | 0.034900306 | 0.131441254 |
| Dusp1 | 0.025429789 | 0.12898598 |
| Mup15 | 5.44009E-15 | 0.001344303 |
| Mup17 | 2.19338E-11 | 0.003397492 |
| Noct | 0.000525453 | 0.264294154 |
| Paqr9 | 0.000134934 | 0.408966406 |
| Acss2 | 0.002029127 | 2.70486964 |
| Aqp8 | 0.012719464 | 3.587024254 |
| Cirbp | 4.99967E-06 | 4.40359119 |
| Coq8a | 5.04854E-05 | 2.439660259 |
| Ethe1 | 2.30385E-06 | 3.683039795 |
| Gm45531 | 0.000139273 | 2.658033794 |
| Gsta2 | 1.88738E-15 | 6.328011776 |
| Orm2 | 0.002704803 | 6.174955264 |
| Rdh9 | 1.72119E-08 | 2.524941514 |
| Tceal8 | 0.036436168 | 2.765468451 |
| Trib3 | 3.26913E-08 | 4.452261778 |

| CRvsAL Shared DEGs | | |
|---|---|---|
| Gene.ID | FDR | FC |
| Derl3 | 0.020736182 | 0.138000616 |
| Dusp1 | 4.47572E-08 | 0.086186749 |
| Mup15 | 4.45866E-12 | 0.000866887 |
| Mup17 | 3.98402E-11 | 0.001133665 |
| Noct | 0.049244831 | 0.199911939 |
| Paqr9 | 0.005922102 | 0.472115697 |
| Acss2 | 0.020328083 | 2.382644939 |
| Aqp8 | 0.00161787 | 2.522258502 |
| Cirbp | 5.67823E-05 | 3.566526043 |
| Coq8a | 1.29486E-06 | 2.362652219 |
| Ethe1 | 3.00032E-06 | 2.835632124 |
| Gm45531 | 0.00081554 | 2.811318622 |
| Gsta2 | 1.52508E-05 | 2.845651531 |
| Orm2 | 0.001741792 | 6.672596384 |
| Rdh9 | 2.48463E-06 | 2.42137742 |
| Tceal8 | 0.011601672 | 2.901930244 |
| Trib3 | 5.85821E-05 | 2.975900472 |
