## Supplementary Table 9 for "Anticipatory metabolic reprogramming distinguishes caloric restriction from fasting-refeeding cycles"

Supplementary Table 9 DEG_Categories_18 hrs for Suppl Fig 3C
Differentially expressed genes (FDR < 0.05)

| CRvsAL Exclusive DEGs | | |
|---|---|---|
| Gene.ID | FDR | FC |
| Angptl8 | 1.51512E-06 | 0.357823576 |
| Cyp2d40 | 2.84877E-10 | 0.496521176 |
| Cyp3a41a | 0.003643867 | 0.045390878 |
| Dusp1 | 0.000354562 | 0.190284043 |
| Fads2 | 0.000272929 | 0.418354082 |
| Fos | 0.031784845 | 0.228027545 |
| Hes6 | 0.00256989 | 0.366757616 |
| Lifr | 1.08091E-12 | 0.290351977 |
| Noct | 0.009752713 | 0.228022078 |
| Onecut1 | 0.00360172 | 0.190640857 |
| Rgs3 | 0.002616796 | 0.267905806 |
| Saa4 | 1.23673E-05 | 0.429248078 |
| Slc10a2 | 8.03847E-05 | 0.179834589 |
| Slc22a28 | 0.017885683 | 0.144676988 |
| Slc22a30 | 0.004776924 | 0.455803228 |
| Tmem254a | 0.008627342 | 0.487441714 |
| Tmem254b | 0.008627342 | 0.487441714 |
| Aprt | 1.11022E-16 | 2.224002674 |
| Cbr1 | 0.000626046 | 2.830319764 |
| Coq8a | 2.94144E-07 | 2.114385386 |
| Gpihbp1 | 0.000162191 | 2.071011317 |

Supplementary Table 9 DEG_Categories_18 hrs for Suppl Fig 3C
Differentially expressed genes (FDR < 0.05)

| FRFvsAL Exclusive DEGs | | |
|---|---|---|
| Gene.ID | FDR | FC |
| Acaa1b | 5.75409E-08 | 0.48923157 |
| Amy2a2 | 0.040883217 | 0.19965094 |
| Amy2a3 | 0.040883217 | 0.19965094 |
| Amy2a4 | 0.040883217 | 0.19965094 |
| BC048546 | 5.64738E-07 | 0.308289029 |
| Bdh2 | 0.001417559 | 0.43118113 |
| Car3 | 0.02759954 | 0.210729374 |
| Cd74 | 9.05244E-10 | 0.379274179 |
| Cebpb | 3.74511E-06 | 0.479690563 |
| Ces1e | 9.11271E-13 | 0.429869729 |
| Clps | 0.015050557 | 0.149593812 |
| Ctrl | 0.037145481 | 0.179990564 |
| Cyp2c44 | 2.88878E-07 | 0.394852234 |
| Fabp1 | 0.000856043 | 0.454594779 |
| Fabp5 | 0.000105178 | 0.281649448 |
| Fdx1 | 6.36032E-09 | 0.481569582 |
| Foxq1 | 0.003388383 | 0.155098292 |
| Gamt | 1.82729E-05 | 0.447556914 |
| Gnat1 | 0.005574144 | 0.287989143 |
| Gpr146 | 0.001510462 | 0.485539176 |
| Gstp3 | 0.011928415 | 0.306340467 |
| Gys2 | 4.79101E-10 | 0.453811214 |
| H2-Aa | 1.03761E-07 | 0.308576746 |
| H2-Ab1 | 1.29951E-09 | 0.295860539 |
| H2-Eb1 | 1.43609E-05 | 0.320319631 |
| Hamp2 | 0.007650052 | 0.089904315 |
| Jchain | 0.02786098 | 0.272367394 |
| Lpin1 | 0.046918284 | 0.356459932 |
| Mcm10 | 0.000901271 | 0.206903828 |
| Nat8f1 | 6.22038E-11 | 0.470781227 |
| Plk3 | 0.027117983 | 0.370231493 |
| Pltp | 0.008945444 | 0.375298216 |
| Pnlip | 0.014255547 | 0.152935309 |
| Rbp1 | 0.046618093 | 0.253705547 |
| Rnf186 | 0.00622475 | 0.21180726 |
| Rpa3 | 0.047305329 | 0.455845461 |
| S100a11 | 0.028894096 | 0.406672213 |
| Saa2 | 0.005648887 | 0.391574677 |
| Slc22a29 | 0.007531088 | 0.255336422 |

Supplementary Table 9 DEG_Categories_18 hrs for Suppl Fig 3C
Differentially expressed genes (FDR < 0.05)

| FRFvsAL Exclusive DEGs | | |
|---|---|---|
| Gene.ID | FDR | FC |
| Slc38a3 | 6.23621E-11 | 0.382494187 |
| Spp1 | 0.044104336 | 0.491833281 |
| Sult1c2 | 0.044253049 | 0.30771938 |
| Try5 | 0.009881963 | 0.141164083 |
| Uox | 4.94482E-12 | 0.437920642 |
| Vegfb | 0.008939718 | 0.443446411 |
| Ypel3 | 0.003151153 | 0.415341313 |
| Acot1 | 3.70695E-06 | 2.275620033 |
| Actb | 2.26262E-05 | 2.328808012 |
| Arhgef19 | 0.000150253 | 2.47671094 |
| Car14 | 3.64435E-07 | 2.232129026 |
| Cldn1 | 0.000246671 | 3.808480987 |
| Clpx | 9.53838E-11 | 2.721062217 |
| Cyp4a10 | 0.000216823 | 4.111403951 |
| Cyp4a14 | 0.001762615 | 5.007183769 |
| Cyp4a31 | 0.00014031 | 3.748843259 |
| Gldc | 0 | 3.37011926 |
| Hfe2 | 0.00083285 | 2.170082023 |
| Retsat | 3.24322E-05 | 2.150098243 |
| Slc20a1 | 0.000589411 | 2.250987756 |
| Srxn1 | 4.02153E-05 | 2.17810884 |
| Tuba8 | 2.09612E-05 | 4.940222932 |

Supplementary Table 9 DEG_Categories_18 hrs for Suppl Fig 3C
Differentially expressed genes (FDR < 0.05)

| CRvsFRF Exclusive DEGs | | |
|---|---|---|
| Gene.ID | FDR | FC |
| 1600002H07Rik | 5.54225E-05 | 0.284322801 |
| Acmsd | 0.003135689 | 0.430232259 |
| Agpat2 | 0 | 0.360504371 |
| C6 | 0.016586745 | 0.147287396 |
| C8b | 0.012133147 | 0.279440106 |
| Chordc1 | 1.5432E-05 | 0.225212056 |
| Cldn2 | 0.000104724 | 0.460014716 |
| Cmah | 0.000231532 | 0.275816044 |
| Creld1 | 8.32491E-10 | 0.445218415 |
| Crp | 0 | 0.458457147 |
| Cyp17a1 | 1.18645E-05 | 0.050735199 |
| Cyp4f14 | 6.64083E-07 | 0.491521324 |
| Cyp7a1 | 7.87955E-09 | 0.242290996 |
| Cyp7b1 | 0.04361062 | 0.156198811 |
| Ddx17 | 5.97462E-06 | 0.27706485 |
| Derl3 | 0.031645268 | 0.14098233 |
| Dnajc3 | 0 | 0.416589823 |
| Egfr | 3.60329E-05 | 0.177462177 |
| Ero1lb | 0.000226511 | 0.328686952 |
| Fdps | 0.011477499 | 0.33282867 |
| Fgf1 | 1.19157E-08 | 0.354428944 |
| Gfra1 | 2.47083E-07 | 0.478866986 |
| Gmppb | 4.09738E-07 | 0.282657917 |
| H2afx | 7.33884E-06 | 0.335933348 |
| Hdhd3 | 6.08119E-08 | 0.358072764 |
| Hhex | 0.015207124 | 0.432953907 |
| Hist2h3c1 | 2.4031E-05 | 0.087702099 |
| Hspb8 | 2.03766E-09 | 0.492489037 |
| Idi1 | 0.016084291 | 0.343196326 |
| Igfals | 0 | 0.431947464 |
| Keg1 | 0.000567498 | 0.252920806 |
| Lad1 | 7.97219E-06 | 0.083896912 |
| Lrrc3 | 2.01144E-05 | 0.301979315 |
| Lss | 0.001557115 | 0.3262352 |
| Mgll | 0 | 0.415721271 |
| Mlec | 1.08329E-06 | 0.323149405 |
| Mup16 | 0 | 0.060520754 |
| Mup3 | 0.046879904 | 0.152017386 |
| Nfil3 | 0.000146755 | 0.19672233 |

Supplementary Table 9 DEG_Categories_18 hrs for Suppl Fig 3C
Differentially expressed genes (FDR < 0.05)

| CRvsFRF Exclusive DEGs | | |
|---|---|---|
| Gene.ID | FDR | FC |
| Nnmt | 0.004967979 | 0.450390934 |
| Nutf2 | 0.017715268 | 0.352575185 |
| Oasl1 | 0.002298867 | 0.303761923 |
| Odc1 | 0 | 0.346676712 |
| Paqr9 | 0 | 0.265661413 |
| Pdia6 | 1.11022E-15 | 0.37083777 |
| Pdlim1 | 8.04999E-05 | 0.45399777 |
| Pnrc1 | 3.97311E-08 | 0.42848225 |
| Ppp1r3b | 0 | 0.273838301 |
| Raph1 | 0.004497042 | 0.388620061 |
| Rarres1 | 2.87146E-05 | 0.22129962 |
| Sdr9c7 | 0.000119123 | 0.224113098 |
| Serpina12 | 0.018110816 | 0.143016713 |
| Serpina3k | 0 | 0.21110319 |
| Slc17a3 | 1.2934E-11 | 0.215543352 |
| Slc22a7 | 0.012220772 | 0.269721634 |
| Slc35b1 | 4.18141E-07 | 0.405957868 |
| Stra6l | 1.41947E-06 | 0.489560876 |
| Tmem150a | 0 | 0.367471658 |
| Tsc22d1 | 2.11016E-06 | 0.247483301 |
| Ube2g2 | 0.020513083 | 0.384881606 |
| Uck1 | 4.92523E-07 | 0.418252129 |
| Urad | 3.97474E-08 | 0.413268441 |
| 1700001C19Rik | 0.006686273 | 2.773966626 |
| Abcd2 | 7.81426E-05 | 2.747522342 |
| Acot3 | 0.004508629 | 4.620475729 |
| Adssl1 | 0.002555928 | 2.994465089 |
| Amdhd1 | 8.92153E-05 | 2.017794832 |
| Angptl4 | 0.009074639 | 2.087338327 |
| Aqp9 | 0 | 2.363840115 |
| Arrdc3 | 0.00143423 | 3.612165825 |
| Car1 | 0.002592525 | 3.560822664 |
| Casp6 | 5.6207E-07 | 3.819063633 |
| Cblc | 4.8943E-05 | 2.61963449 |
| Cd63 | 0.002538965 | 2.539374928 |
| Ces1g | 0 | 2.87405281 |
| Ces2e | 8.65876E-09 | 2.257478985 |
| Ciart | 0.001345428 | 3.802642283 |
| Cpt1a | 4.82862E-10 | 2.033053552 |

Supplementary Table 9 DEG_Categories_18 hrs for Suppl Fig 3C

Differentially expressed genes (FDR < 0.05)

| CRvsFRF Exclusive DEGs | | |
|---|---|---|
| Gene.ID | FDR | FC |
| Cryl1 | 2.13013E-06 | 3.338889357 |
| Cyp2a22 | 2.81661E-05 | 6.896974819 |
| Cyp2c55 | 0.000266516 | 6.53209573 |
| Cyp3a11 | 1.77527E-05 | 2.982705975 |
| Flot1 | 0.000588148 | 2.300781596 |
| Galk1 | 2.5277E-06 | 2.04360914 |
| Gm2000 | 0.000602321 | 2.053202797 |
| Gm3839 | 0.045201861 | 4.871547174 |
| Gstm2 | 0.041239589 | 2.346052217 |
| Gstm4 | 5.82942E-06 | 2.406972133 |
| Gstm6 | 4.99279E-05 | 2.528417965 |
| Gstt2 | 0 | 2.699433886 |
| Haus7 | 5.43845E-05 | 3.189705939 |
| Hoga1 | 3.33067E-16 | 2.661016161 |
| Hsd17b6 | 0.025205531 | 3.40526832 |
| Idh2 | 0 | 2.080990756 |
| Il18 | 0.000557663 | 3.52917186 |
| Lgmn | 0.000209771 | 2.036172682 |
| Mfsd2a | 0.000170256 | 2.015793079 |
| Orm3 | 0.001710005 | 3.203163936 |
| Prom1 | 0.042461088 | 3.651721017 |
| Psme1 | 0.008422286 | 2.259713157 |
| Rasgrp2 | 3.70032E-05 | 2.229701837 |
| Rbm3 | 0.001572782 | 3.444563662 |
| Rgs16 | 1.2999E-09 | 5.910109892 |
| Rnf19b | 0.007648083 | 2.015796697 |
| Ropn1l | 0.015985289 | 2.397510691 |
| Rras | 0.002037873 | 2.017522499 |
| Rtn4 | 2.38249E-10 | 2.495143346 |
| S100a4 | 0.01560705 | 4.875143847 |
| Sds | 3.23115E-05 | 2.840848797 |
| Serpinb1a | 0.020333169 | 2.989742579 |
| Slc22a27 | 0.007819448 | 3.231947653 |
| Slc2a2 | 1.44329E-15 | 2.217530056 |
| Slc8b1 | 1.92424E-12 | 6.042416258 |
| Slco1a4 | 8.44104E-05 | 2.310907083 |
| Soat2 | 1.1714E-06 | 3.632730788 |
| Sorbs3 | 4.2175E-06 | 3.416035041 |
| Sqrdl | 0 | 2.058937399 |

Supplementary Table 9 DEG_Categories_18 hrs for Suppl Fig 3C
Differentially expressed genes (FDR < 0.05)

| CRvsFRF Exclusive DEGs | | |
|---|---|---|
| Gene.ID | FDR | FC |
| Sult2a7 | 2.31315E-05 | 12.17250162 |
| Tmem86a | 0.013720329 | 2.796542021 |
| Txnip | 0.00755624 | 4.663336479 |
| Ubd | 0.034338371 | 4.925323402 |
| Ugp2 | 0 | 2.037841129 |
| Ugt1a5 | 4.48048E-05 | 3.108287743 |
| Unc119 | 0.019606347 | 2.772755626 |

Supplementary Table 9 DEG_Categories_18 hrs for Suppl Fig 3C

Differentially expressed genes (FDR < 0.05)

| CRvsAL Shared DEGs | | |
|---|---|---|
| Gene.ID | FDR | FC |
| Rcan1 | 1.04996E-05 | 0.411060261 |
| Gsta2 | 0 | 3.329883715 |
| Raet1d | 1.24103E-05 | 2.355771217 |

| FRFvsAL Shared DEGs | | |
|---|---|---|
| Gene.ID | FDR | FC |
| Gsta2 | 0.002677521 | 0.265581803 |
| Raet1d | 0.000216494 | 0.26622687 |
| Rcan1 | 4.91889E-05 | 2.253436045 |

| CRvsFRF Shared DEGs | | |
|---|---|---|
| Gene.ID | FDR | FC |
| Rcan1 | 0 | 0.182414878 |
| Gsta2 | 0 | 12.53807182 |
| Raet1d | 0 | 8.848735733 |

Supplementary Table 9 DEG_Categories_18 hrs for Suppl Fig 3C

Differentially expressed genes (FDR < 0.05)

| CRvsAL Shared DEGs | | |
|---|---|---|
| Gene.ID | FDR | FC |
| Btg2 | 0.001172573 | 0.323207194 |
| Cela2a | 0.002250588 | 0.063553998 |
| Cfd | 0.021159124 | 0.123097338 |
| Cish | 0.000510605 | 0.286661739 |
| Csrnp1 | 0.020383261 | 0.315510753 |
| Ctrb1 | 0 | 0.033530036 |
| Cyp3a16 | 0.000234909 | 0.044265133 |
| Cyp3a41b | 2.31743E-05 | 0.026066394 |
| Cyp3a44 | 9.70241E-06 | 0.025719602 |
| G6pc | 8.57246E-05 | 0.325190152 |
| Gstp2 | 0.045686529 | 0.102872874 |
| Mafb | 0.005901722 | 0.313052811 |
| Mug2 | 0.003767801 | 0.468933155 |
| Nr4a1 | 0.002029837 | 0.082181525 |
| Pim3 | 0.008104349 | 0.399415647 |
| Slc25a25 | 6.98218E-05 | 0.295768592 |
| Slc25a30 | 0.048392041 | 0.304148404 |
| Tff3 | 0 | 0.028745544 |
| Thrsp | 6.12337E-06 | 0.247494825 |
| Tob1 | 1.02361E-05 | 0.442671419 |
| 2010003K11Rik | 0.011915504 | 2.235911582 |
| Ethe1 | 4.0426E-05 | 2.872152762 |
| Insig2 | 9.79327E-06 | 2.250651772 |

| FRFvsAL Shared DEGs | | |
|---|---|---|
| Gene.ID | FDR | FC |
| Btg2 | 0.003478761 | 0.374345166 |
| Cela2a | 0.006584384 | 0.08763657 |
| Cfd | 0.009126829 | 0.120845401 |
| Cish | 1.74098E-05 | 0.232778775 |
| Csrnp1 | 0.010173611 | 0.3142003 |
| Ctrb1 | 7.90188E-05 | 0.040225522 |
| Cyp3a16 | 1.33908E-06 | 0.023007618 |
| Cyp3a41b | 0 | 0.020516793 |
| Cyp3a44 | 0.007512745 | 0.088332984 |
| G6pc | 0.010552886 | 0.427818711 |
| Gstp2 | 0.046283473 | 0.120848991 |
| Mafb | 0.002712429 | 0.29735265 |
| Mug2 | 0.000948947 | 0.447188375 |
| Nr4a1 | 0.004236819 | 0.107813032 |
| Pim3 | 0.038765435 | 0.422652907 |
| Slc25a25 | 0.010072909 | 0.452474518 |
| Slc25a30 | 0.02379014 | 0.307844983 |
| Tff3 | 0.000119145 | 0.039753809 |
| Thrsp | 0.002393546 | 0.370654222 |
| Tob1 | 9.11859E-07 | 0.395566229 |
| 2010003K11Rik | 1.32962E-09 | 3.480497248 |
| Ethe1 | 0.002192042 | 2.03538992 |
| Insig2 | 0 | 3.459127706 |

Supplementary Table 9 DEG_Categories_18 hrs for Suppl Fig 3C
Differentially expressed genes (FDR < 0.05)

| FRFvsAL Shared DEGs | | |
|---|---|---|
| Gene.ID | FDR | FC |
| Abcc2 | 2.06495E-10 | 0.489320695 |
| Abcg5 | 0 | 0.310096702 |
| Abcg8 | 2.8314E-12 | 0.247017467 |
| Agxt | 0.018510013 | 0.470449119 |
| Bdh1 | 6.1901E-07 | 0.355412829 |
| Ces2a | 1.42912E-06 | 0.463288595 |
| Coq10b | 0 | 0.282448076 |
| Cyp2a5 | 1.97331E-08 | 0.337327707 |
| Cyp2g1 | 0.002138348 | 0.182247918 |
| Dbp | 0 | 0.033006355 |
| Gpcpd1 | 4.24771E-06 | 0.194211821 |
| Gsta4 | 2.46492E-05 | 0.473769827 |
| H2afv | 1.06954E-07 | 0.395492993 |
| Hao2 | 0.000545342 | 0.084146197 |
| Inmt | 0 | 0.267192718 |
| Junb | 3.66467E-09 | 0.23113763 |
| Lgals1 | 0.036683514 | 0.240901184 |
| Lpin2 | 1.65676E-06 | 0.344639849 |
| Nr1d1 | 1.20731E-11 | 0.105195946 |
| Nr1d2 | 0.001311509 | 0.204571254 |
| Pcsk4 | 4.85167E-14 | 0.1579907 |
| Pigp | 0.00625593 | 0.48583895 |
| Ppp1r3g | 0.000214748 | 0.039447952 |
| Scd1 | 0.008725791 | 0.115892258 |
| Serpina6 | 1.22605E-06 | 0.146812914 |
| Slc25a47 | 1.62093E-14 | 0.377702102 |
| Slc7a2 | 5.1481E-13 | 0.352695979 |
| Sucnr1 | 0.002347147 | 0.307783337 |
| Sult2a1 | 4.57884E-05 | 0.030464091 |
| Sult2a2 | 1.33181E-05 | 0.025294627 |
| Sult2a3 | 8.89271E-11 | 0.053215301 |
| Sult2a5 | 1.58463E-05 | 0.045164258 |
| Tef | 1.82464E-07 | 0.372553703 |
| Tmem50a | 8.69016E-12 | 0.467474916 |
| Tmsb10 | 0.000140912 | 0.343087976 |
| Tstd1 | 1.26522E-05 | 0.317097748 |
| Upp2 | 0 | 0.128905654 |
| Acnat2 | 9.05942E-05 | 6.727527535 |
| Actg1 | 0.000230561 | 3.070571065 |

| CRvsFRF Shared DEGs | | |
|---|---|---|
| Gene.ID | FDR | FC |
| Acnat2 | 2.13354E-05 | 0.154828078 |
| Actg1 | 4.6943E-10 | 0.198807997 |
| Ahsa1 | 0 | 0.342832667 |
| Apoa4 | 0.040093827 | 0.242420137 |
| Atf5 | 3.29162E-10 | 0.402884829 |
| Atp2a2 | 0 | 0.272132115 |
| Bag3 | 6.39544E-12 | 0.208372929 |
| Bcl6 | 1.35362E-05 | 0.164510754 |
| Bhmt | 0.0054964 | 0.433006784 |
| Calr | 0 | 0.297792233 |
| Caprin1 | 3.12238E-07 | 0.427954553 |
| Cdkn1a | 0 | 0.076967309 |
| Chd9 | 1.16454E-07 | 0.288528409 |
| Chka | 2.58438E-12 | 0.184545381 |
| Creld2 | 0 | 0.046301711 |
| Crip2 | 0.008883612 | 0.441424553 |
| Cyp39a1 | 0.03101415 | 0.452849499 |
| Cyp8b1 | 0.002905329 | 0.397998076 |
| Ddc | 3.88578E-15 | 0.329536598 |
| Ddit3 | 1.39522E-06 | 0.187797488 |
| Dnaja1 | 0 | 0.276672246 |
| Dnajb1 | 2.00581E-08 | 0.276518527 |
| Dnajb11 | 0 | 0.265961734 |
| Dnajb2 | 0 | 0.319406912 |
| Dnajb9 | 0 | 0.30461696 |
| Dynll1 | 6.83651E-11 | 0.292155592 |
| Eif4ebp3 | 0 | 0.264193723 |
| Fkbp4 | 0 | 0.284999444 |
| Gck | 0 | 0.18078419 |
| Herpud1 | 0.002149181 | 0.46048736 |
| Hist1h1c | 2.48177E-09 | 0.305543739 |
| Hist2h3c2 | 1.21443E-05 | 0.092903226 |
| Hsp90aa1 | 0 | 0.085927626 |
| Hsp90ab1 | 0 | 0.284744978 |
| Hsp90b1 | 6.21144E-11 | 0.276070198 |
| Hspa1b | 0 | 0.065368129 |
| Hspa5 | 0 | 0.112454229 |
| Hspa8 | 0 | 0.158911997 |
| Hspb1 | 9.47575E-13 | 0.090043508 |

Supplementary Table 9 DEG_Categories_18 hrs for Suppl Fig 3C
Differentially expressed genes (FDR < 0.05)

| FRFvsAL Shared DEGs | | | CRvsFRF Shared DEGs | | |
|---|---|---|---|---|---|
| Gene.ID | FDR | FC | Gene.ID | FDR | FC |
| Ahsa1 | 0 | 2.082707052 | Hsph1 | 0 | 0.086146977 |
| Apoa4 | 0.008096375 | 5.606905415 | Hyou1 | 0 | 0.114980283 |
| Atf5 | 5.20695E-14 | 2.746422515 | Leap2 | 7.32747E-15 | 0.428385992 |
| Atp2a2 | 0 | 3.39025207 | Lrrc59 | 6.951E-12 | 0.327129402 |
| Bag3 | 5.74831E-08 | 3.197539265 | Mal2 | 9.00959E-06 | 0.329043422 |
| Bcl6 | 0.00023741 | 4.664407015 | Manf | 0 | 0.12033308 |
| Bhmt | 1.33511E-06 | 2.79788673 | Morf4l2 | 1.33677E-08 | 0.453758313 |
| Calr | 7.77156E-16 | 2.425541751 | Mvd | 2.97578E-06 | 0.127186013 |
| Caprin1 | 0.002368904 | 2.083623294 | Palld | 1.92387E-11 | 0.252929018 |
| Cdkn1a | 0 | 11.39709162 | Pdia3 | 2.22045E-16 | 0.355193227 |
| Chd9 | 1.70674E-05 | 2.784580846 | Pdia4 | 0 | 0.17848046 |
| Chka | 7.99052E-09 | 4.002812141 | Plin5 | 0.000593897 | 0.465491305 |
| Creld2 | 2.06823E-12 | 10.62730425 | Pnkd | 3.27915E-11 | 0.462110016 |
| Crip2 | 0.017896779 | 2.425011472 | Ppp1r3c | 0.000662607 | 0.388958397 |
| Cyp39a1 | 0.000262041 | 3.89372659 | Rorc | 4.01043E-11 | 0.479039471 |
| Cyp8b1 | 3.03543E-06 | 2.595360346 | Rtp3 | 1.50069E-12 | 0.22314668 |
| Ddc | 9.40581E-13 | 2.645362553 | S100a10 | 1.26065E-06 | 0.45997168 |
| Ddit3 | 3.94347E-06 | 5.354577172 | Sdf2l1 | 0 | 0.040013543 |
| Dnaja1 | 0 | 2.557424207 | Sec14l4 | 0 | 0.400316409 |
| Dnajb1 | 1.88582E-05 | 2.659604511 | Snrpa | 4.519E-05 | 0.377000791 |
| Dnajb11 | 5.32907E-15 | 2.553648793 | Sqle | 7.14721E-06 | 0.208815771 |
| Dnajb2 | 2.97984E-12 | 2.229525008 | Stip1 | 0 | 0.223139798 |
| Dnajb9 | 5.29909E-13 | 2.116253891 | Syvn1 | 0 | 0.14781372 |
| Dynll1 | 3.50402E-11 | 3.317945217 | Tardbp | 2.21785E-11 | 0.396405556 |
| Eif4ebp3 | 0 | 5.407211569 | Tsc22d3 | 0 | 0.277329241 |
| Fkbp4 | 0 | 2.162876918 | Tuba1c | 0 | 0.232372872 |
| Gck | 8.54494E-12 | 2.315364166 | Tubb2a | 0 | 0.175419466 |
| Herpud1 | 0.00099656 | 2.128613073 | Tubb4b | 0 | 0.32972941 |
| Hist1h1c | 0.001081709 | 2.346900918 | Ubc | 0 | 0.388632556 |
| Hist2h3c2 | 0.000999116 | 8.396479132 | Ubxn4 | 0.000183933 | 0.40373515 |
| Hsp90aa1 | 0 | 7.04247941 | Abcc2 | 2.12674E-12 | 2.151454247 |
| Hsp90ab1 | 0 | 3.043043308 | Abcg5 | 0 | 3.932193884 |
| Hsp90b1 | 3.88413E-08 | 3.022213155 | Abcg8 | 1.76879E-11 | 3.979358314 |
| Hspa1b | 4.27993E-09 | 4.95653136 | Agxt | 4.80258E-08 | 3.236423642 |
| Hspa5 | 0 | 6.097472219 | Bdh1 | 0 | 3.019359733 |
| Hspa8 | 1.83187E-14 | 3.290439926 | Ces2a | 2.5192E-08 | 2.160427586 |
| Hspb1 | 2.53592E-10 | 8.353072595 | Coq10b | 2.18516E-06 | 2.468003092 |
| Hsph1 | 0 | 5.07243148 | Cyp2a5 | 1.23235E-14 | 4.411343768 |
| Hyou1 | 0 | 5.30686638 | Cyp2g1 | 8.82674E-06 | 9.441936681 |

Supplementary Table 9 DEG_Categories_18 hrs for Suppl Fig 3C
Differentially expressed genes (FDR < 0.05)

| FRFvsAL Shared DEGs | | |
|---|---|---|
| Gene.ID | FDR | FC |
| Leap2 | 0 | 3.745379297 |
| Lrrc59 | 4.88339E-05 | 2.412793506 |
| Mal2 | 4.98313E-06 | 2.158053155 |
| Manf | 0 | 5.549941763 |
| Morf4l2 | 1.91559E-05 | 2.043923334 |
| Mvd | 0.000862949 | 4.884801346 |
| Palld | 2.9526E-09 | 3.611677881 |
| Pdia3 | 1.18498E-09 | 2.153131071 |
| Pdia4 | 6.27276E-14 | 4.279677725 |
| Plin5 | 1.22125E-15 | 2.818328742 |
| Pnkd | 1.04565E-11 | 2.213248991 |
| Ppp1r3c | 0.015451167 | 2.17830598 |
| Rorc | 3.39704E-06 | 2.088217485 |
| Rtp3 | 1.22156E-06 | 2.414187063 |
| S100a10 | 8.82254E-09 | 2.678598292 |
| Sdf2l1 | 0 | 11.54081353 |
| Sec14l4 | 7.53574E-10 | 2.008933662 |
| Snrpa | 0.007279022 | 2.440154501 |
| Sqle | 0.039138321 | 2.848644907 |
| Stip1 | 0 | 3.454055517 |
| Syvn1 | 0 | 5.123576227 |
| Tardbp | 2.69451E-13 | 2.567699509 |
| Tsc22d3 | 0 | 2.374697504 |
| Tuba1c | 3.9968E-14 | 2.344387578 |
| Tubb2a | 0 | 8.630297588 |
| Tubb4b | 1.00697E-13 | 2.195642673 |
| Ubc | 0 | 3.133020088 |
| Ubxn4 | 0.009056935 | 2.338962326 |

| CRvsFRF Shared DEGs | | |
|---|---|---|
| Gene.ID | FDR | FC |
| Dbp | 0 | 32.1008205 |
| Gpcpd1 | 7.15599E-07 | 4.77287422 |
| Gsta4 | 0 | 3.661968112 |
| H2afv | 4.17851E-06 | 2.415526666 |
| Hao2 | 0.000267453 | 14.44012294 |
| Inmt | 0 | 4.265475432 |
| Junb | 0.000218114 | 2.759993073 |
| Lgals1 | 2.32832E-11 | 4.352670888 |
| Lpin2 | 0 | 4.715502028 |
| Nr1d1 | 1.44329E-14 | 12.1282117 |
| Nr1d2 | 9.97237E-05 | 5.124065615 |
| Pcsk4 | 2.75044E-11 | 5.543226502 |
| Pigp | 3.19636E-07 | 2.924367042 |
| Ppp1r3g | 0.000322648 | 10.57094708 |
| Scd1 | 8.91144E-11 | 5.064798618 |
| Serpina6 | 0 | 5.247965815 |
| Slc25a47 | 3.33067E-16 | 3.85684705 |
| Slc7a2 | 1.38778E-14 | 3.545166763 |
| Sucnr1 | 3.72788E-05 | 2.982814429 |
| Sult2a1 | 1.21663E-05 | 32.95320565 |
| Sult2a2 | 3.38325E-06 | 42.63961267 |
| Sult2a3 | 0 | 58.55314251 |
| Sult2a5 | 1.28972E-07 | 44.8167993 |
| Tef | 6.40408E-11 | 2.897772313 |
| Tmem50a | 1.26282E-08 | 2.164899391 |
| Tmsb10 | 0.001923089 | 3.452564651 |
| Tstd1 | 6.26832E-13 | 3.628352463 |
| Upp2 | 3.60382E-05 | 5.890972893 |

Supplementary Table 9 DEG_Categories_18 hrs for Suppl Fig 3C
Differentially expressed genes (FDR < 0.05)

| CRvsFRF Shared DEGs | | |
|---|---|---|
| Gene.ID | FDR | FC |
| Aacs | 2.11599E-05 | 0.281461289 |
| Ces3b | 9.63778E-09 | 0.220848372 |
| Cyp4a12a | 3.5939E-06 | 0.027014936 |
| Cyp4a12b | 0.000179938 | 0.063719965 |
| Dio1 | 0 | 0.288649083 |
| Elovl3 | 5.29394E-06 | 0.023188443 |
| Fasn | 1.43121E-05 | 0.327180098 |
| Hsd3b5 | 1.02334E-06 | 0.023172134 |
| Inhbc | 0.00023022 | 0.478152661 |
| Jun | 0.00622329 | 0.355907842 |
| Mid1ip1 | 2.80794E-10 | 0.408832112 |
| Mup1 | 5.14971E-11 | 0.004313148 |
| Mup10 | 1.15707E-12 | 0.279094506 |
| Mup11 | 1.64195E-07 | 0.002690428 |
| Mup12 | 1.64987E-09 | 0.009882968 |
| Mup13 | 3.5083E-14 | 0.090539635 |
| Mup14 | 2.85169E-09 | 0.008861613 |
| Mup15 | 0 | 0.000900101 |
| Mup17 | 1.4877E-14 | 0.001825171 |
| Mup18 | 0 | 0.104192798 |
| Mup19 | 0 | 0.029842962 |
| Mup2 | 0 | 0.015725549 |
| Mup20 | 0.000218458 | 0.033144702 |
| Mup21 | 0.007184688 | 0.128392419 |
| Mup6 | 0.019502442 | 0.215398758 |
| Mup7 | 2.35199E-11 | 0.002799954 |
| Mup8 | 0.005243281 | 0.114879764 |
| Mup9 | 5.12923E-14 | 0.018117058 |
| Nat8f5 | 2.51528E-05 | 0.054560929 |
| Nrep | 3.71723E-07 | 0.214853607 |
| RP24-346O11.2 | 0 | 0.084823285 |
| Selenbp2 | 2.35168E-05 | 0.028909209 |
| Serpina1c | 4.94785E-05 | 0.431183767 |
| Serpina1e | 6.82105E-11 | 0.004678485 |
| Slco1a1 | 6.43322E-08 | 0.023145017 |
| Sult5a1 | 4.31951E-06 | 0.185698122 |
| Tsku | 0.000333429 | 0.330517345 |
| Ugt3a1 | 1.51005E-06 | 0.453923395 |

| CRvsAL Shared DEGs | | |
|---|---|---|
| Gene.ID | FDR | FC |
| Aacs | 1.26246E-07 | 0.243295996 |
| Ces3b | 0.008351098 | 0.219478684 |
| Cyp4a12a | 6.75883E-06 | 0.022884555 |
| Cyp4a12b | 0.000378616 | 0.055390357 |
| Dio1 | 2.63055E-08 | 0.262220464 |
| Elovl3 | 0.018259487 | 0.12572071 |
| Fasn | 0.002267791 | 0.461660514 |
| Hsd3b5 | 7.96468E-08 | 0.013979347 |
| Inhbc | 3.32748E-06 | 0.471973823 |
| Jun | 0.002105576 | 0.393448883 |
| Mid1ip1 | 0.00056746 | 0.410360197 |
| Mup1 | 1.04119E-08 | 0.00496009 |
| Mup10 | 1.50031E-06 | 0.216271824 |
| Mup11 | 2.53239E-09 | 0.003439595 |
| Mup12 | 5.86921E-09 | 0.005730695 |
| Mup13 | 0.000478689 | 0.080467688 |
| Mup14 | 1.26171E-07 | 0.006883155 |
| Mup15 | 4.44089E-15 | 0.00101866 |
| Mup17 | 2.70817E-12 | 0.001609222 |
| Mup18 | 0.01492847 | 0.107736866 |
| Mup19 | 0.001680609 | 0.046975365 |
| Mup2 | 9.80103E-08 | 0.014711565 |
| Mup20 | 0.0020819 | 0.039313648 |
| Mup21 | 0.001665798 | 0.064358428 |
| Mup6 | 0.044141957 | 0.149022192 |
| Mup7 | 2.06152E-10 | 0.002410252 |
| Mup8 | 0.001373764 | 0.078689782 |
| Mup9 | 6.03429E-06 | 0.019441267 |
| Nat8f5 | 0.006852897 | 0.114360301 |
| Nrep | 0 | 0.138707814 |
| RP24-346O11.2 | 0.001311852 | 0.10246546 |
| Selenbp2 | 1.75055E-08 | 0.009511436 |
| Serpina1c | 0.011083723 | 0.44901291 |
| Serpina1e | 4.12513E-10 | 0.003543524 |
| Slco1a1 | 4.84808E-07 | 0.019781772 |
| Sult5a1 | 0.002429042 | 0.189713993 |
| Tsku | 1.46352E-06 | 0.435294201 |
| Ugt3a1 | 5.6397E-10 | 0.424451677 |

Supplementary Table 9 DEG_Categories_18 hrs for Suppl Fig 3C
Differentially expressed genes (FDR < 0.05)

| CRvsFRF Shared DEGs | | |
|---|---|---|
| Gene.ID | FDR | FC |
| 8430408G22 Rik | 6.93601E-09 | 5.097586363 |
| Akr1b7 | 0.000948671 | 4.24201019 |
| Cd36 | 5.50627E-05 | 2.2608536 |
| Cirbp | 5.05315E-09 | 5.182630331 |
| Cyp3a59 | 2.95844E-10 | 3.628516221 |
| Fbxo21 | 6.30607E-14 | 4.260386358 |
| Fmo3 | 0 | 11.35882558 |
| Gm3776 | 0 | 18.36663223 |
| Gm45531 | 1.9702E-07 | 7.286465529 |
| Gsta1 | 0 | 22.34129589 |
| Gstm1 | 0 | 3.432386652 |
| Gstm3 | 7.91867E-12 | 11.95831853 |
| Gstt3 | 3.35458E-06 | 2.018233651 |
| Mgst3 | 2.02956E-09 | 3.696715469 |
| Orm2 | 6.10707E-07 | 9.788982677 |
| Rdh9 | 0 | 3.870802552 |
| Tceal8 | 0.000451353 | 2.537005894 |
| Tgtp1 | 0.00246373 | 14.01918238 |

| CRvsAL Shared DEGs | | |
|---|---|---|
| Gene.ID | FDR | FC |
| 8430408G22 Rik | 0.000718148 | 2.818281126 |
| Akr1b7 | 0.020196369 | 3.856705835 |
| Cd36 | 4.70797E-05 | 2.187470353 |
| Cirbp | 0.002858865 | 2.315964773 |
| Cyp3a59 | 2.19078E-07 | 3.023725547 |
| Fbxo21 | 2.69032E-05 | 2.164264137 |
| Fmo3 | 4.44089E-15 | 7.05461394 |
| Gm3776 | 3.33067E-14 | 5.397915078 |
| Gm45531 | 0.002310121 | 3.508308998 |
| Gsta1 | 2.01482E-05 | 3.447999661 |
| Gstm1 | 4.39703E-05 | 2.035177058 |
| Gstm3 | 3.54461E-06 | 5.719638638 |
| Gstt3 | 8.82945E-10 | 2.311418754 |
| Mgst3 | 0.040893019 | 2.089777855 |
| Orm2 | 4.38058E-07 | 11.27248018 |
| Rdh9 | 1.04138E-07 | 2.106624652 |
| Tceal8 | 5.28477E-07 | 3.682350312 |
| Tgtp1 | 0.018080854 | 12.4705215 |
