## Supplementary Table 10 for "Anticipatory metabolic reprogramming distinguishes caloric restriction from fasting-refeeding cycles"

Supplementary Table 10 DEG_Categories_22 hrs for Suppl Fig 3C
Differentially expressed genes (FDR < 0.05)

| CRvsAL Exclusive DEGs | | |
|---|---|---|
| Gene.ID | FDR | FC |
| Apon | 0.002740541 | 0.492686732 |
| Car14 | 3.38516E-05 | 0.335754164 |
| Cldn1 | 3.86996E-05 | 0.356644807 |
| Ddc | 0.008872595 | 0.4298744 |
| Dexi | 0.000232498 | 0.476455319 |
| Gck | 0.001752063 | 0.430604005 |
| Gtf3c6 | 0.000515649 | 0.457003141 |
| Hsd3b7 | 5.47505E-08 | 0.47378237 |
| Hsp90aa1 | 0.000697248 | 0.475461824 |
| Lasp1 | 9.68636E-05 | 0.485501816 |
| Leap2 | 0.007038164 | 0.307275336 |
| Mlec | 0.006929145 | 0.456945448 |
| Mup14 | 0.022102387 | 0.051322963 |
| Nat8f2 | 1.73107E-07 | 0.439516265 |
| Nsmf | 3.10193E-06 | 0.434075827 |
| Paqr9 | 0.000211454 | 0.443709606 |
| Psen2 | 0 | 0.462170401 |
| Slc10a2 | 0.000173244 | 0.197677963 |
| Slc17a3 | 0.041565037 | 0.491317906 |
| Slc35b1 | 0.018531248 | 0.448292155 |
| Sult5a1 | 0.002386219 | 0.28118253 |
| Suox | 2.02061E-14 | 0.471062146 |
| Tmem254a | 3.41443E-08 | 0.46157796 |
| Tmem254b | 3.41443E-08 | 0.46157796 |
| Tubb2a | 2.40418E-10 | 0.346535616 |
| Tubb4b | 8.91422E-07 | 0.454756233 |
| Ugt2b37 | 0.031303043 | 0.196850359 |
| Ak4 | 0.002895199 | 2.010564998 |
| Aldoc | 0.03449837 | 2.468244311 |
| Cnppd1 | 0.044049427 | 2.049069969 |
| Eif4b | 6.03878E-08 | 2.036661649 |
| Fbxo21 | 0.004262808 | 2.145852168 |
| Fmo3 | 0.004214822 | 14.92548587 |
| Gstt2 | 0 | 2.537481604 |
| Hba-a2 | 0.039178615 | 4.600116094 |
| Hbb-bs | 0.006178635 | 4.606085079 |
| Lgals4 | 1.38556E-13 | 4.895683461 |
| Lurap1l | 0.005711281 | 2.019847262 |
| Pck1 | 0.018059189 | 2.194503828 |

Supplementary Table 10 DEG_Categories_22 hrs for Suppl Fig 3C

Differentially expressed genes (FDR < 0.05)

| CRvsAL Exclusive DEGs | | |
|---|---|---|
| Gene.ID | FDR | FC |
| Pla2g12a | 0.046725244 | 2.105916719 |
| Ppp1r10 | 5.60885E-05 | 3.073917385 |
| Rpl22l1 | 2.67509E-10 | 2.225147651 |
| Soat2 | 0.00562905 | 2.82479667 |
| Txnip | 0.001040081 | 2.617121157 |

Supplementary Table 10 DEG_Categories_22 hrs for Suppl Fig 3C
Differentially expressed genes (FDR < 0.05)

| FRFvsAL Exclusive DEGs | | |
|---|---|---|
| Gene.ID | FDR | FC |
| A230050P20Rik | 0.004588242 | 0.360988295 |
| Acy3 | 0.000165817 | 0.466760059 |
| Adh4 | 1.02801E-08 | 0.407288266 |
| Ak1 | 0.024622387 | 0.192613777 |
| Amd1 | 0.011845457 | 0.490884319 |
| Bcl6 | 0.006422905 | 0.24377971 |
| Bhlhe40 | 0.003308917 | 0.494031657 |
| C6 | 0.00389092 | 0.108384871 |
| Car3 | 2.55795E-13 | 0.096706853 |
| Casq1 | 0.042827174 | 0.218548301 |
| Ces2c | 0.005680097 | 0.227116813 |
| Comtd1 | 0.016774296 | 0.299637819 |
| Cxx1b | 0.028972216 | 0.401078064 |
| Cyp2c67 | 0.000586386 | 0.438788391 |
| Cyp2u1 | 0.001939511 | 0.126775055 |
| Cyp7b1 | 0.010445733 | 0.124093816 |
| Elovl6 | 0.001105522 | 0.304954583 |
| Fabp5 | 0.003608326 | 0.237333344 |
| Gdf15 | 0.001951914 | 0.24694442 |
| Gstp2 | 0.011088646 | 0.125610394 |
| Hsd3b2 | 0.013499649 | 0.33676691 |
| Hspb7 | 0.04099595 | 0.2198921 |
| Kyat1 | 0.015334861 | 0.439302519 |
| Mcm10 | 0.017577086 | 0.23902211 |
| Mmd2 | 0.008689235 | 0.14796541 |
| Mup1 | 0.00146218 | 0.033411322 |
| Mup12 | 0.008720644 | 0.058267296 |
| Mup20 | 0.004198181 | 0.053820614 |
| Mup21 | 9.78375E-06 | 0.035334151 |
| Mup7 | 0.011680955 | 0.040108285 |
| Nat8f1 | 0 | 0.440742896 |
| Nat8f5 | 4.47047E-05 | 0.055663756 |
| Ociad2 | 0.006182095 | 0.270281817 |
| Pdia3 | 0.000380056 | 0.440094155 |
| Pygm | 0.033259175 | 0.169681102 |
| Qprt | 0.001860528 | 0.457721875 |
| Raet1d | 0.035590303 | 0.398007229 |
| Saa3 | 0.000193961 | 0.396754273 |
| Scd1 | 0.041025421 | 0.070533689 |

Supplementary Table 10 DEG_Categories_22 hrs for Suppl Fig 3C
Differentially expressed genes (FDR < 0.05)

| FRFvsAL Exclusive DEGs | | |
|---|---|---|
| Gene.ID | FDR | FC |
| Sec23b | 0.02923564 | 0.461809202 |
| Serpina1e | 8.92975E-06 | 0.016939046 |
| Serpina7 | 0.010998123 | 0.275969295 |
| Slc22a28 | 0.000827611 | 0.117171825 |
| Slc9a3r1 | 2.39173E-10 | 0.484018049 |
| Slco1a1 | 0.001244904 | 0.097682124 |
| Spsb4 | 0.001265288 | 0.173322092 |
| Srsf3 | 0.000321217 | 0.423730368 |
| Tbcb | 8.3775E-05 | 0.468709798 |
| Tcea3 | 3.01822E-05 | 0.402815231 |
| Tgm2 | 0.007032684 | 0.409879032 |
| Tmem150a | 5.78426E-14 | 0.438737377 |
| Tmsb10 | 0.048073991 | 0.417249457 |
| Tuba1b | 8.99645E-08 | 0.332010936 |
| Tuba4a | 0.004382353 | 0.431186044 |
| Tubb5 | 0.002246281 | 0.499965303 |
| Txndc5 | 9.80158E-06 | 0.499663116 |
| Ube2l6 | 6.4504E-05 | 0.364320951 |
| Ugt2b38 | 0.013171741 | 0.162950825 |
| Vegfb | 0.004184978 | 0.426953304 |
| Zbp1 | 0.010269124 | 0.23799987 |
| 1810055G02Rik | 0.001347336 | 3.56543341 |
| A1cf | 3.35033E-05 | 2.00075905 |
| Aaed1 | 1.84143E-07 | 2.421932889 |
| Aass | 8.39917E-09 | 2.025930586 |
| Abca6 | 1.2192E-06 | 2.497941356 |
| Abcb4 | 7.20535E-14 | 2.148989511 |
| Acacb | 0.023685581 | 2.099878796 |
| Acadm | 0 | 2.018664579 |
| Acot1 | 3.73546E-12 | 3.665358233 |
| Acot12 | 1.82488E-10 | 2.250373255 |
| Acot3 | 0.006717841 | 2.665829778 |
| Acot4 | 2.05221E-05 | 2.002331468 |
| Acss3 | 2.32234E-05 | 2.784915741 |
| Agpat3 | 0 | 2.224109576 |
| Ahcy | 0 | 2.408610456 |
| Arl4a | 8.71303E-13 | 3.038418763 |
| Atl2 | 2.14808E-09 | 2.179089115 |
| Ccng2 | 0.041435291 | 3.748097676 |

Supplementary Table 10 DEG_Categories_22 hrs for Suppl Fig 3C
Differentially expressed genes (FDR < 0.05)

| FRFvsAL Exclusive DEGs | | |
|---|---|---|
| Gene.ID | FDR | FC |
| Cd163 | 0.037120025 | 3.053916926 |
| Cdip1 | 0 | 2.797559459 |
| Cdo1 | 4.32987E-15 | 2.354033594 |
| Ces1b | 1.09912E-14 | 2.360902967 |
| Chd9 | 8.10793E-06 | 2.817672769 |
| Chic2 | 0.01254075 | 2.57874868 |
| Cpa2 | 0.008905029 | 6.798839018 |
| Cpt1a | 9.91847E-07 | 2.295368615 |
| Crat | 1.01147E-11 | 2.252308092 |
| Cyp2b10 | 2.1264E-11 | 3.924670144 |
| Cyp3a59 | 8.46877E-05 | 2.272294105 |
| D230025D16Rik | 5.5454E-06 | 2.323971626 |
| Dnmbp | 0.002554422 | 2.738850927 |
| Ei24 | 0 | 2.060837559 |
| Fam234b | 0.001762258 | 2.212327429 |
| Glrx | 0 | 2.220896361 |
| Got1 | 0.003855104 | 3.099643272 |
| Gpcpd1 | 0.003747172 | 3.947022742 |
| Grpel2 | 6.53488E-06 | 2.77405903 |
| Hacl1 | 2.12317E-09 | 2.167957681 |
| Hmgcs2 | 8.99281E-15 | 2.319333576 |
| Ip6k2 | 0 | 3.744889043 |
| Klf9 | 4.19575E-12 | 3.355264025 |
| Lbp | 2.22045E-16 | 2.635011349 |
| Lcn2 | 0.002309956 | 2.480311173 |
| Lrg1 | 6.43929E-15 | 2.282268399 |
| Map1lc3b | 0 | 2.225123373 |
| Mt1 | 0.000700745 | 5.549134039 |
| Nceh1 | 0.018127496 | 2.32752062 |
| Ncoa4 | 1.31151E-12 | 2.032353394 |
| Npc1 | 4.18474E-06 | 2.200101045 |
| Pcx | 0 | 2.224134158 |
| Pex11a | 1.42109E-13 | 2.785969852 |
| Pim3 | 1.27165E-06 | 2.08140159 |
| Plscr2 | 0.000125373 | 2.085899118 |
| Ppara | 6.72326E-06 | 2.31721522 |
| Rabgef1 | 0.007198647 | 2.269371637 |
| Rapgef4 | 0.007164405 | 2.00011077 |
| Rgs16 | 7.54952E-15 | 7.939740289 |

Supplementary Table 10 DEG_Categories_22 hrs for Suppl Fig 3C
Differentially expressed genes (FDR < 0.05)

| FRFvsAL Exclusive DEGs | | |
|---|---|---|
| Gene.ID | FDR | FC |
| Rhobtb1 | 0.000200902 | 2.067649141 |
| Sec14l4 | 1.53512E-09 | 2.027617873 |
| Serinc3 | 0 | 2.280366406 |
| Serpina3m | 1.10507E-09 | 2.460388971 |
| Slc10a1 | 5.97406E-07 | 2.006523589 |
| Slc22a5 | 1.18394E-05 | 3.011912072 |
| Slc25a22 | 0.015769718 | 2.022168815 |
| Slc25a30 | 0.036397463 | 3.044066017 |
| Slc25a47 | 7.84602E-06 | 3.03894638 |
| Slc37a4 | 1.00586E-13 | 2.030373596 |
| St3gal5 | 2.46044E-11 | 4.5758674 |
| Sun2 | 2.27247E-07 | 2.702125941 |
| Tango2 | 0 | 2.771741948 |
| Tdo2 | 5.81541E-10 | 2.211360967 |
| Tm4sf4 | 0 | 2.258124181 |
| Tmem106b | 0.012813415 | 2.06545867 |
| Tmem120a | 4.77396E-15 | 2.011144227 |
| Tmem56 | 5.63736E-07 | 2.574997535 |
| Trp53inp1 | 0.000178878 | 2.455256652 |
| Ttpa | 0 | 2.349351731 |
| Vnn3 | 1.68141E-10 | 2.457381233 |
| Wbp1l | 1.16335E-10 | 2.040421445 |

Supplementary Table 10 DEG_Categories_22 hrs for Suppl Fig 3C
Differentially expressed genes (FDR < 0.05)

| CRvsFRF Exclusive DEGs | | |
|---|---|---|
| Gene.ID | FDR | FC |
| Arl4d | 1.15762E-07 | 0.478642794 |
| Atf5 | 0 | 0.331351512 |
| Dio1 | 0 | 0.293728863 |
| Fabp2 | 2.55351E-15 | 0.454330189 |
| Fos | 0.000874465 | 0.209665882 |
| Idi1 | 0.000412292 | 0.418530352 |
| Klk1 | 0.01849125 | 0.161937708 |
| Kynu | 0 | 0.433186222 |
| Mgll | 7.54619E-13 | 0.478333194 |
| Mup16 | 7.26646E-05 | 0.305368199 |
| Mup18 | 2.76446E-14 | 0.201300267 |
| Mup19 | 2.22045E-16 | 0.165809463 |
| Nnmt | 4.75204E-05 | 0.473704125 |
| Nt5e | 4.85269E-06 | 0.319614017 |
| Rarres1 | 4.04472E-05 | 0.305773733 |
| RP24-346O11.2 | 9.3337E-09 | 0.224820365 |
| Sdr42e1 | 0.000363062 | 0.439782443 |
| Sqle | 0.003490587 | 0.353741704 |
| Sult3a1 | 1.12579E-07 | 0.180996457 |
| Ucp2 | 0.000112008 | 0.483766948 |
| Ugt3a1 | 1.97198E-06 | 0.466663892 |
| Cib3 | 0.000811125 | 11.52772861 |
| Dbp | 0.007257569 | 6.223711415 |
| Ddx3y | 0.026633748 | 11.60753164 |
| Erdr1 | 0.037574426 | 2.88199519 |
| Gm20594 | 0.008025819 | 2.206681898 |
| Gsta2 | 4.65846E-06 | 3.754115666 |
| Gstm1 | 4.78284E-13 | 2.488817832 |
| Gstm3 | 5.51781E-14 | 5.899520626 |
| Gys2 | 0.000111847 | 2.085567529 |
| Rassf3 | 3.81535E-05 | 3.73800898 |
| Sult1c2 | 0.003856779 | 3.396987764 |
| Sult2a5 | 8.40741E-06 | 25.9627682 |

Supplementary Table 10 DEG_Categories_22 hrs for Suppl Fig 3C
Differentially expressed genes (FDR < 0.05)

| CRvsAL Shared DEGs | | |
|---|---|---|
| Gene.ID | FDR | FC |
| Hes6 | 0.019363784 | 0.454088223 |
| Srebf1 | 0.000314199 | 0.265132787 |

| FRFvsAL Shared DEGs | | |
|---|---|---|
| Gene.ID | FDR | FC |
| Hes6 | 1.11059E-08 | 0.187987447 |
| Srebf1 | 9.58008E-11 | 0.089996787 |

| CRvsFRF Shared DEGs | | |
|---|---|---|
| Gene.ID | FDR | FC |
| Hes6 | 0.039169934 | 2.415524172 |
| Srebf1 | 1.10127E-06 | 2.946025035 |

Supplementary Table 10 DEG_Categories_22 hrs for Suppl Fig 3C

Differentially expressed genes (FDR < 0.05)

| CRvsAL Shared DEGs | | |
|---|---|---|
| Gene.ID | FDR | FC |
| A1bg | 0.000161468 | 0.022494117 |
| Acta1 | 0 | 0.007112479 |
| Actb | 2.1123E-06 | 0.462234901 |
| Actg1 | 0.000312774 | 0.256448608 |
| Angptl8 | 0.019887877 | 0.174875998 |
| Ankrd23 | 0.033019047 | 0.139115508 |
| Apol9a | 3.39328E-05 | 0.361855082 |
| Atp2a1 | 0 | 0.030606404 |
| Calr | 0.000201905 | 0.446967353 |
| Ckm | 1.09464E-06 | 0.015119127 |
| Ckmt2 | 1.44329E-15 | 0.043521007 |
| Cox6a2 | 0.000436964 | 0.058978239 |
| Cox8b | 1.84666E-07 | 0.062097443 |
| Cyp2c70 | 6.64135E-13 | 0.45547095 |
| Derl3 | 0.005034792 | 0.122701987 |
| Dnajb11 | 0.02927422 | 0.440632056 |
| Eef1a2 | 0.012722121 | 0.12783209 |
| Fabp3 | 3.64921E-05 | 0.031758607 |
| Gadd45a | 0.008267785 | 0.276466072 |
| Gale | 4.59787E-05 | 0.183769259 |
| Hdhd3 | 2.29736E-05 | 0.277377563 |
| Hist1h1c | 0.021091703 | 0.302694247 |
| Insc | 0.000120306 | 0.288287297 |
| Lifr | 0.000493267 | 0.302207225 |
| Ly6d | 0.015948297 | 0.090509239 |
| Mb | 0 | 0.009809615 |
| Mid1ip1 | 1.69715E-05 | 0.314977717 |
| Myh1 | 0.00038161 | 0.075334998 |
| Myh2 | 0.001837762 | 0.078809804 |
| Myl1 | 0 | 0.038215964 |
| Myl2 | 6.78106E-10 | 0.055026425 |
| Mylpf | 0.000481439 | 0.046507667 |
| Nrep | 0.000233615 | 0.355524411 |
| Oasl1 | 0.018491822 | 0.300625183 |
| Pdia6 | 0.012903498 | 0.408921903 |
| Rcan1 | 4.30415E-06 | 0.381332338 |
| Saa1 | 0.01042045 | 0.338947616 |
| Sdf2l1 | 0.01085324 | 0.118309897 |
| Selenbp2 | 2.21781E-05 | 0.033694678 |

| FRFvsAL Shared DEGs | | |
|---|---|---|
| Gene.ID | FDR | FC |
| A1bg | 4.64778E-06 | 0.013575622 |
| Acta1 | 0 | 0.007112479 |
| Actb | 1.00169E-10 | 0.342104662 |
| Actg1 | 6.33981E-06 | 0.203865937 |
| Angptl8 | 2.51702E-05 | 0.073432933 |
| Ankrd23 | 0.023922754 | 0.152735067 |
| Apol9a | 7.73289E-05 | 0.39278156 |
| Atp2a1 | 0 | 0.030606404 |
| Calr | 0.000345847 | 0.467938794 |
| Ckm | 0 | 0.014101846 |
| Ckmt2 | 1.33227E-15 | 0.043521007 |
| Cox6a2 | 3.82991E-09 | 0.057239703 |
| Cox8b | 0.000876056 | 0.075176627 |
| Cyp2c70 | 0 | 0.387241973 |
| Derl3 | 0.007555478 | 0.150169467 |
| Dnajb11 | 0.012593383 | 0.43583227 |
| Eef1a2 | 0.003854362 | 0.121858657 |
| Fabp3 | 0 | 0.028665514 |
| Gadd45a | 0.000201479 | 0.194098854 |
| Gale | 0.000517833 | 0.26575103 |
| Hdhd3 | 8.39951E-06 | 0.271096277 |
| Hist1h1c | 0.003364561 | 0.268144745 |
| Insc | 3.367E-06 | 0.289777755 |
| Lifr | 9.24522E-06 | 0.234429031 |
| Ly6d | 0.016835126 | 0.10821178 |
| Mb | 1.2062E-07 | 0.010833492 |
| Mid1ip1 | 2.57049E-11 | 0.196451771 |
| Myh1 | 0.000305644 | 0.075334998 |
| Myh2 | 0.001462295 | 0.078809804 |
| Myl1 | 0 | 0.038215964 |
| Myl2 | 5.74218E-10 | 0.055026425 |
| Mylpf | 0.000823292 | 0.058511474 |
| Nrep | 1.31006E-14 | 0.087752864 |
| Oasl1 | 0.038511951 | 0.360964439 |
| Pdia6 | 0.000617567 | 0.345715726 |
| Rcan1 | 0.000158213 | 0.478208593 |
| Saa1 | 0.048069922 | 0.437179075 |
| Sdf2l1 | 0.012317609 | 0.140093366 |
| Selenbp2 | 9.40491E-07 | 0.023529007 |

Supplementary Table 10 DEG_Categories_22 hrs for Suppl Fig 3C
Differentially expressed genes (FDR < 0.05)

| CRvsAL Shared DEGs | | |
|---|---|---|
| Gene.ID | FDR | FC |
| Slc22a30 | 1.4426E-05 | 0.468230013 |
| Tcap | 2.13938E-05 | 0.030526018 |
| Tff3 | 0.009953766 | 0.101286331 |
| Thrsp | 0.031312332 | 0.462200809 |
| Tnnc2 | 0 | 0.013324462 |
| Tnni2 | 0 | 0.01856975 |
| Tnnt3 | 0 | 0.012516599 |
| Tuba1c | 8.90557E-06 | 0.371654328 |
| Acot2 | 7.58608E-05 | 3.353889184 |
| Angptl4 | 8.47803E-05 | 3.012691692 |
| Bnip3 | 0 | 2.040963693 |
| Ces1g | 0.001807636 | 2.384660636 |
| Ctgf | 0.023117743 | 2.762446575 |
| Fbxo31 | 0.00229881 | 2.809205215 |
| Fkbp5 | 0.034471522 | 2.933677653 |
| Gabarapl1 | 0.000135676 | 2.043413157 |
| Gm45531 | 2.83497E-10 | 6.151280272 |
| Il6ra | 0.003358306 | 2.456574411 |
| Lpin1 | 2.149E-06 | 7.04833915 |
| Lpin2 | 0 | 3.958765011 |
| Mfsd2a | 4.17264E-05 | 2.568720052 |
| Pnpla2 | 4.2866E-10 | 2.306619815 |
| Por | 4.97783E-06 | 2.233938769 |
| Rab30 | 0.036352634 | 3.099032407 |
| Rbm3 | 0 | 3.005465789 |
| Slc16a7 | 0.006054518 | 2.217238353 |
| Slc8b1 | 1.6406E-06 | 2.646000484 |
| Slco1a4 | 0.002569096 | 2.332368301 |
| Sorbs3 | 0.007890249 | 2.667211321 |
| Sult1e1 | 0.023935219 | 7.813292957 |

| FRFvsAL Shared DEGs | | |
|---|---|---|
| Gene.ID | FDR | FC |
| Slc22a30 | 9.87941E-09 | 0.40744626 |
| Tcap | 0 | 0.030102349 |
| Tff3 | 0.003341694 | 0.09708531 |
| Thrsp | 6.64842E-07 | 0.317227385 |
| Tnnc2 | 0 | 0.013324462 |
| Tnni2 | 3.63907E-05 | 0.030436936 |
| Tnnt3 | 0 | 0.012516599 |
| Tuba1c | 7.58067E-07 | 0.378502403 |
| Acot2 | 0.003289587 | 3.679152353 |
| Angptl4 | 0 | 4.135683187 |
| Bnip3 | 0 | 2.417277549 |
| Ces1g | 6.18609E-05 | 2.199517557 |
| Ctgf | 0.000223594 | 3.263300559 |
| Fbxo31 | 4.87388E-14 | 4.219375735 |
| Fkbp5 | 0 | 5.277351898 |
| Gabarapl1 | 3.32857E-11 | 2.246605337 |
| Gm45531 | 3.60607E-07 | 6.063228051 |
| Il6ra | 5.48677E-09 | 3.337380937 |
| Lpin1 | 4.45735E-09 | 10.5834441 |
| Lpin2 | 6.20842E-06 | 3.545548036 |
| Mfsd2a | 1.0705E-09 | 3.629791239 |
| Pnpla2 | 1.66064E-07 | 3.482393788 |
| Por | 1.04362E-07 | 2.83484284 |
| Rab30 | 9.61646E-08 | 5.100997691 |
| Rbm3 | 5.48531E-10 | 2.637323478 |
| Slc16a7 | 8.78962E-07 | 2.275920132 |
| Slc8b1 | 5.18052E-09 | 3.355120382 |
| Slco1a4 | 3.29566E-09 | 2.925260625 |
| Sorbs3 | 0.011028262 | 2.738345137 |
| Sult1e1 | 0.007782338 | 6.431837048 |

Supplementary Table 10 DEG_Categories_22 hrs for Suppl Fig 3C
Differentially expressed genes (FDR < 0.05)

| FRFvsAL Shared DEGs | | |
|---|---|---|
| Gene.ID | FDR | FC |
| Aqp8 | 1.6472E-08 | 0.372025104 |
| BC048546 | 0.001433467 | 0.26201291 |
| Bdh2 | 0.018288205 | 0.344827516 |
| Cyp1a2 | 0 | 0.413718762 |
| Cyp2c44 | 2.30749E-12 | 0.311024027 |
| Cyp2c55 | 0.013327382 | 0.141364026 |
| Cyp3a16 | 0.021129702 | 0.16804868 |
| Cyp3a41b | 0.008303037 | 0.159337097 |
| Cyp4a12a | 9.38186E-09 | 0.013017981 |
| Cyp4a12b | 4.1814E-07 | 0.023992328 |
| Eif2s3y | 4.03034E-07 | 0.063473258 |
| Fam25c | 0.001578344 | 0.367797134 |
| Fam47e | 0.005676731 | 0.303278246 |
| Gas6 | 0.006218384 | 0.34075295 |
| Gm3776 | 0.009775483 | 0.15437848 |
| Gm3839 | 0.027555964 | 0.284144978 |
| Gsta1 | 0.00243967 | 0.114555197 |
| Gstm6 | 1.3833E-06 | 0.411302562 |
| Gstp3 | 0.003942899 | 0.37831944 |
| Hao2 | 0.042689738 | 0.180551099 |
| Hsd3b5 | 4.36415E-08 | 0.017472419 |
| Inmt | 0 | 0.480075727 |
| Lgals1 | 7.14792E-08 | 0.24694017 |
| Nat8 | 0.000243058 | 0.079462621 |
| Psmb10 | 2.09567E-08 | 0.431924959 |
| Serpina6 | 0.000273083 | 0.211416685 |
| 2010003K11Rik | 0.012828477 | 2.647919905 |
| 2210010C04Rik | 0.000346485 | 15.17308137 |
| Acnat2 | 1.65618E-05 | 2.99196617 |
| Amy2a2 | 0.007243284 | 7.078133317 |
| Amy2a3 | 0.007243284 | 7.078133317 |
| Amy2a4 | 0.007243284 | 7.078133317 |
| Amy2a5 | 2.45761E-07 | 80.5898748 |
| Apoa4 | 7.31579E-07 | 7.821023084 |
| Bhmt | 3.28782E-09 | 4.350357441 |
| Cdkn1a | 0 | 14.70908858 |
| Cel | 0.000953363 | 12.23359011 |
| Cela2a | 7.46159E-05 | 32.40447702 |
| Cela3b | 2.67645E-06 | 48.11498484 |

| CRvsFRF Shared DEGs | | |
|---|---|---|
| Gene.ID | FDR | FC |
| 2010003K11Rik | 0.002338317 | 0.262818575 |
| 2210010C04Rik | 0.000420647 | 0.064606453 |
| Acnat2 | 0 | 0.122831543 |
| Amy2a2 | 0.009371398 | 0.139957605 |
| Amy2a3 | 0.009371398 | 0.139957605 |
| Amy2a4 | 0.009371398 | 0.139957605 |
| Amy2a5 | 3.11112E-06 | 0.016779983 |
| Apoa4 | 7.66794E-06 | 0.140013096 |
| Bhmt | 7.77156E-16 | 0.146959275 |
| Cdkn1a | 0 | 0.096049625 |
| Cel | 0.001234956 | 0.080772181 |
| Cela2a | 2.09323E-05 | 0.023140038 |
| Cela3b | 2.87261E-06 | 0.019963675 |
| Clps | 1.84644E-05 | 0.028141431 |
| Clpx | 0 | 0.305765748 |
| Cpa1 | 2.4006E-05 | 0.033492205 |
| Cpb1 | 0.000514007 | 0.069552145 |
| Csrp3 | 5.34431E-05 | 0.384037546 |
| Ctrb1 | 4.36204E-08 | 0.0073486 |
| Ctrl | 0.000405922 | 0.050986832 |
| Cxcl13 | 0.025385608 | 0.190733426 |
| Cyp17a1 | 0 | 0.070814257 |
| Cyp2c39 | 0.000427486 | 0.407498252 |
| Cyp39a1 | 2.81726E-08 | 0.385450918 |
| Cyp4a10 | 0 | 0.308492973 |
| Cyp4a14 | 0 | 0.298732003 |
| Cyp4a31 | 2.4576E-08 | 0.269976634 |
| Cyp8b1 | 0.02837858 | 0.245638254 |
| Ddit4 | 4.34117E-05 | 0.11699492 |
| Defb1 | 9.83931E-06 | 0.23030274 |
| Eif4ebp3 | 1.2182E-05 | 0.305023298 |
| Fam134b | 0.04721117 | 0.385348182 |
| Fgf21 | 3.1699E-05 | 0.210483819 |
| Fmo2 | 2.47791E-12 | 0.389935879 |
| Gadd45b | 0.000282045 | 0.416918732 |
| Gadd45g | 0.00284732 | 0.39976088 |
| Gfra1 | 3.33067E-16 | 0.469776718 |
| Gldc | 0 | 0.372072929 |
| Gm4952 | 9.68658E-12 | 0.37514922 |

Supplementary Table 10 DEG_Categories_22 hrs for Suppl Fig 3C
Differentially expressed genes (FDR < 0.05)

| FRFvsAL Shared DEGs | | |
|---|---|---|
| Gene.ID | FDR | FC |
| Clps | 0 | 44.88868458 |
| Clpx | 0 | 2.975348364 |
| Cpa1 | 1.36765E-05 | 30.55806889 |
| Cpb1 | 0.000405779 | 14.18202751 |
| Csrp3 | 6.34578E-07 | 2.47225583 |
| Ctrb1 | 2.08529E-08 | 142.2276814 |
| Ctrl | 3.12106E-05 | 26.89257825 |
| Cxcl13 | 0.027323207 | 4.846709011 |
| Cyp17a1 | 2.62793E-06 | 4.14463347 |
| Cyp2c39 | 1.4606E-07 | 6.336789154 |
| Cyp39a1 | 0 | 5.454240683 |
| Cyp4a10 | 0 | 7.098840491 |
| Cyp4a14 | 0 | 7.032732983 |
| Cyp4a31 | 1.1779E-07 | 3.513363675 |
| Cyp8b1 | 0.036664676 | 3.375710794 |
| Ddit4 | 7.98529E-05 | 7.539814594 |
| Defb1 | 4.5964E-11 | 8.429093343 |
| Eif4ebp3 | 0 | 8.367821776 |
| Fam134b | 3.3691E-06 | 5.7326733 |
| Fgf21 | 5.47306E-09 | 8.6313226 |
| Fmo2 | 8.18856E-12 | 4.60514469 |
| Gadd45b | 2.6333E-08 | 5.903167812 |
| Gadd45g | 0.002667863 | 3.115211539 |
| Gfra1 | 0 | 2.425185243 |
| Gldc | 1.64313E-14 | 2.354638175 |
| Gm4952 | 9.06275E-13 | 3.912734708 |
| Hsd17b10 | 0 | 2.614302646 |
| Insig2 | 0 | 4.937489652 |
| Lepr | 0.000945693 | 4.242301754 |
| Mat1a | 5.11841E-09 | 2.669881392 |
| Mt2 | 8.83331E-06 | 13.9684119 |
| Nlrp12 | 4.5125E-12 | 5.827051899 |
| Plin5 | 0 | 4.300667363 |
| Pnlip | 8.10365E-06 | 41.84583741 |
| Pnliprp1 | 0.019638135 | 7.748291279 |
| Prss2 | 7.0326E-07 | 63.20758888 |
| Reg1 | 0.000362351 | 14.21301888 |
| Retsat | 0 | 3.927902648 |
| Rnase1 | 8.95093E-05 | 19.65373431 |

| CRvsFRF Shared DEGs | | |
|---|---|---|
| Gene.ID | FDR | FC |
| Hsd17b10 | 0 | 0.493384301 |
| Insig2 | 0 | 0.123208948 |
| Lepr | 0.000978232 | 0.236930852 |
| Mat1a | 9.20097E-12 | 0.417624789 |
| Mt2 | 0.00034516 | 0.106906505 |
| Nlrp12 | 0.00024226 | 0.401167901 |
| Plin5 | 2.22045E-16 | 0.381356489 |
| Pnlip | 1.82148E-06 | 0.018535194 |
| Pnliprp1 | 0.000486477 | 0.066799103 |
| Prss2 | 1.56513E-06 | 0.016686829 |
| Reg1 | 1.51758E-05 | 0.069300622 |
| Retsat | 0 | 0.455763242 |
| Rnase1 | 3.25617E-12 | 0.049742441 |
| S100a10 | 0 | 0.203583066 |
| Sco2 | 1.51902E-10 | 0.414443676 |
| Serpina3n | 2.77012E-12 | 0.451646593 |
| Slc45a3 | 2.43685E-07 | 0.318811966 |
| Sult1d1 | 0 | 0.422566868 |
| Sult3a2 | 0.000149787 | 0.172524458 |
| Sycn | 0.000157639 | 0.041865877 |
| Try4 | 4.84192E-07 | 0.014553031 |
| Try5 | 4.90402E-06 | 0.022375837 |
| Tymp | 3.22142E-06 | 0.265538835 |
| Vnn1 | 1.11022E-16 | 0.368356449 |
| Zg16 | 0.000137277 | 0.047991822 |
| Aqp8 | 0.004870778 | 2.585027595 |
| BC048546 | 7.25311E-09 | 3.870085189 |
| Bdh2 | 0.041415234 | 2.150537871 |
| Cyp1a2 | 5.84385E-05 | 2.85552346 |
| Cyp2c44 | 0 | 2.869026384 |
| Cyp2c55 | 0.015147095 | 6.80733354 |
| Cyp3a16 | 0.001048701 | 16.67011171 |
| Cyp3a41b | 0.000157695 | 20.5750672 |
| Cyp4a12a | 0.000149144 | 20.23734092 |
| Cyp4a12b | 0.001300732 | 11.49111425 |
| Eif2s3y | 4.41449E-09 | 17.3942819 |
| Fam25c | 0.004234393 | 2.271237362 |
| Fam47e | 0.001536463 | 2.338756194 |
| Gas6 | 0.029030079 | 2.201916904 |

Supplementary Table 10 DEG_Categories_22 hrs for Suppl Fig 3C

Differentially expressed genes (FDR < 0.05)

| FRFvsAL Shared DEGs | | |
|---|---|---|
| Gene.ID | FDR | FC |
| S100a10 | 2.82709E-09 | 2.918105477 |
| Sco2 | 7.26013E-11 | 2.836782654 |
| Serpina3n | 6.1543E-12 | 2.650037212 |
| Slc45a3 | 0.003454022 | 2.552441498 |
| Sult1d1 | 0 | 3.988141473 |
| Sult3a2 | 0.027776077 | 3.410205667 |
| Sycn | 0.00016663 | 22.3118637 |
| Try4 | 9.82666E-07 | 60.06844425 |
| Try5 | 0 | 51.33903113 |
| Tymp | 8.79801E-06 | 3.602731368 |
| Vnn1 | 2.44896E-07 | 3.071217447 |
| Zg16 | 3.33067E-16 | 23.26291042 |

| CRvsFRF Shared DEGs | | |
|---|---|---|
| Gene.ID | FDR | FC |
| Gm3776 | 0.001237656 | 10.2211535 |
| Gm3839 | 0.034486865 | 6.724992825 |
| Gsta1 | 0.008453427 | 12.56257588 |
| Gstm6 | 1.10709E-06 | 2.787639677 |
| Gstp3 | 2.5524E-13 | 3.570599337 |
| Hao2 | 0.010060308 | 6.690984529 |
| Hsd3b5 | 0.000212489 | 18.37366236 |
| Inmt | 0 | 3.813846765 |
| Lgals1 | 0.00011438 | 3.060587125 |
| Nat8 | 0.002430511 | 10.80063736 |
| Psmb10 | 0.006052128 | 2.266096826 |
| Serpina6 | 1.11135E-06 | 5.136097301 |

Supplementary Table 10 DEG_Categories_22 hrs for Suppl Fig 3C

Differentially expressed genes (FDR < 0.05)

| CRvsFRF Shared DEGs | | |
|---|---|---|
| Gene.ID | FDR | FC |
| Dusp1 | 0.000479693 | 0.433414265 |
| Hfe2 | 1.87308E-07 | 0.396386368 |
| Mup11 | 0.000558371 | 0.162616973 |
| Mup15 | 0 | 0.004201549 |
| Mup17 | 0 | 0.009987453 |
| Mup2 | 8.40994E-13 | 0.113757957 |
| Mup9 | 0 | 0.145717049 |
| Btbd19 | 0.018726301 | 6.180225685 |
| Clec2h | 0.001558238 | 6.286309082 |
| Cyp2a5 | 0 | 2.795489861 |
| Hbb-bt | 0.047035704 | 3.509745344 |
| Pcsk4 | 0.000930375 | 3.809967249 |
| Sult2a3 | 1.70932E-05 | 27.45804412 |
| Zpr1 | 0.000108453 | 2.08828486 |

| CRvsAL Shared DEGs | | |
|---|---|---|
| Gene.ID | FDR | FC |
| Dusp1 | 0.009074032 | 0.480545347 |
| Hfe2 | 8.9493E-05 | 0.494058947 |
| Mup11 | 0.001298062 | 0.029745716 |
| Mup15 | 1.22191E-12 | 0.002231482 |
| Mup17 | 1.04601E-11 | 0.002032802 |
| Mup2 | 0.000534978 | 0.045864712 |
| Mup9 | 0.009828442 | 0.06478358 |
| Btbd19 | 0.034198831 | 6.11296469 |
| Clec2h | 0.007026118 | 5.477260673 |
| Cyp2a5 | 0 | 3.206588652 |
| Hbb-bt | 0.037179066 | 3.749696997 |
| Pcsk4 | 0.005070984 | 3.544167656 |
| Sult2a3 | 0.002691416 | 13.27224087 |
| Zpr1 | 1.16134E-08 | 3.044291561 |
